# Multilevel proteomics reveals host perturbations by SARS-CoV-2 and SARS-CoV

**DOI:** 10.1101/2020.06.17.156455

**Authors:** Alexey Stukalov, Virginie Girault, Vincent Grass, Ozge Karayel, Valter Bergant, Christian Urban, Darya A. Haas, Yiqi Huang, Lila Oubraham, Anqi Wang, M. Sabri Hamad, Antonio Piras, Fynn M. Hansen, Maria C. Tanzer, Igor Paron, Luca Zinzula, Thomas Enghleitner, Maria Reinecke, Teresa M. Lavacca, Rosina Ehmann, Roman Wölfel, Jörg Jores, Bernhard Kuster, Ulrike Protzer, Roland Rad, John Ziebuhr, Volker Thiel, Pietro Scaturro, Matthias Mann, Andreas Pichlmair

## Abstract

The global emergence of SARS-CoV-2 urgently requires an in-depth understanding of molecular functions of viral proteins and their interactions with the host proteome. Several individual omics studies have extended our knowledge of COVID-19 pathophysiology^1–10^. Integration of such datasets to obtain a holistic view of virus-host interactions and to define the pathogenic properties of SARS-CoV-2 is limited by the heterogeneity of the experimental systems. We therefore conducted a concurrent multi-omics study of SARS-CoV-2 and SARS-CoV. Using state-of-the-art proteomics, we profiled the interactome of both viruses, as well as their influence on transcriptome, proteome, ubiquitinome and phosphoproteome in a lung-derived human cell line. Projecting these data onto the global network of cellular interactions revealed crosstalk between the perturbations taking place upon SARS-CoV-2 and SARS-CoV infections at different layers and identified unique and common molecular mechanisms of these closely related coronaviruses. The TGF-β pathway, known for its involvement in tissue fibrosis, was specifically dysregulated by SARS-CoV-2 ORF8 and autophagy by SARS-CoV-2 ORF3. The extensive dataset (available at https://covinet.innatelab.org) highlights many hotspots that can be targeted by existing drugs and it can guide rational design of virus- and host-directed therapies, which we exemplify by identifying kinase and MMPs inhibitors with potent antiviral effects against SARS-CoV-2.

## Main text

### Comparative SARS-CoV-2 and SARS-CoV virus-host interactome and effectome

To identify interactions of SARS-CoV-2 and SARS-CoV with cellular proteins, we transduced A549 lung carcinoma cells with lentiviruses expressing individual HA-tagged viral proteins (Figure 1a; Extended data Fig. 1a; Supplementary Table 1). Affinity purification followed by mass spectrometry (AP-MS) analysis and statistical modelling of the quantitative data identified 1□801 interactions between 1□086 cellular proteins and 24 SARS-CoV-2 and 27 SARS-CoV bait proteins (Figure 1b; Extended data Fig. 1b; Supplementary Table 2), significantly expanding the currently reported interactions of SARS-CoV-2 and SARS-CoV (Supplementary Table 10)^1–11^. The resulting virus-host interaction network revealed a wide range of cellular activities intercepted by SARS-CoV-2 and SARS-CoV (Figure 1b; Extended data Table 1; Supplementary Table 2). In particular, we discovered that SARS-CoV-2 targets a number of key innate immunity regulators (ORF7b–MAVS, –UNC93B1), stress response components (N–HSPA1A) and DNA damage response mediators (ORF7a–ATM, –ATR) (Figure 1b; Extended data Fig. 1c-e). Additionally, SARS-CoV-2 proteins interact with molecular complexes involved in intracellular trafficking (*e*.*g*. ER Golgi trafficking) and transport (*e*.*g*. Solute carriers, Ion transport by ATPases) as well as cellular metabolism (*e*.*g*. Mitochondrial respiratory chain, Glycolysis) (Figure 1b, Extended data Table 1, Supplementary Table 2). Comparing the AP-MS data of homologous SARS-CoV-2 and SARS-CoV proteins identified differences in the enrichment of individual host targets, highlighting potential virus-specific interactions (Figure 1b (edge color); Figure 1c; Extended data Fig. 1f, 2a-b; Supplementary Table 2). For instance, we recapitulated the known interaction between SARS-CoV NSP2 and prohibitins (PHB, PHB2)^12^ but this was not conserved in SARS-CoV-2 NSP2, suggesting that the two viruses differ in their ability to modulate mitochondrial function and homeostasis through NSP2 (Extended data Fig. 2a). The exclusive interaction of SARS-CoV-2 ORF8 with the TGFB1-LTBP1 complex is another interaction potentially explaining the differences in pathogenicity of the two viruses (Extended data Fig. 1f, 2b). Notably, disbalanced TGF-β signaling has been linked to lung fibrosis and oedema, a common complication of severe pulmonary diseases including COVID-19^13–16^.

**Figure 1.**
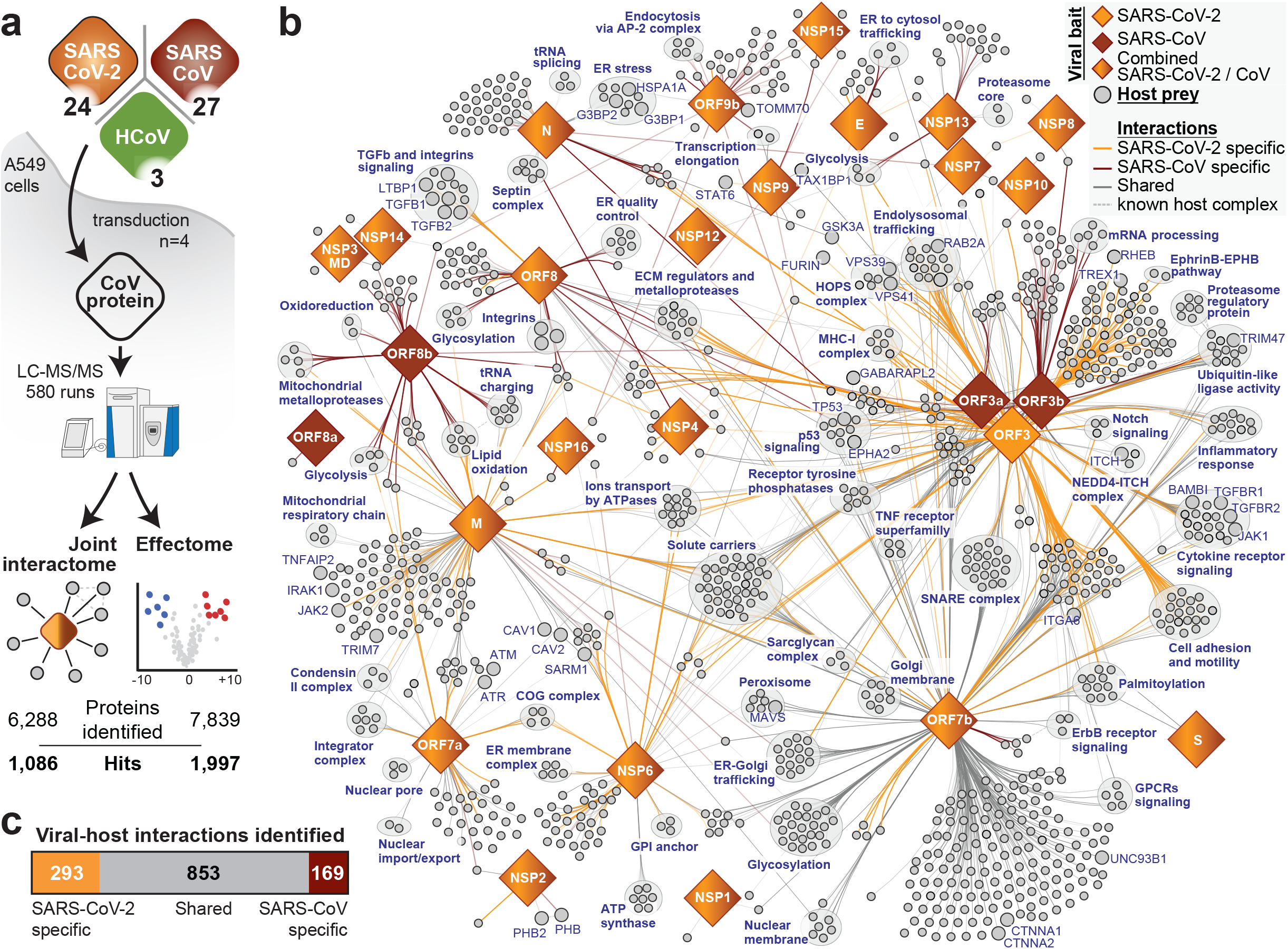
Joint analysis of SARS-CoV-2 and SARS-CoV protein-protein virus-host interactomes. **(a)** Systematic comparison of interactomes and host proteome changes (“effectomes”) of the homologous SARS-CoV-2 and SARS-CoV viral proteins, with ORF3 homologs of HCoV-NL63 and HCoV-229E as reference for pan-coronavirus specificity. **(b)** Combined virus-host protein interaction network of SARS-CoV-2 and SARS-CoV measured by AP-MS. Homologous viral proteins are displayed as a single node. Shared and virus-specific interactions are denoted by the edge color. The edge color gradient reflects the p-value of the interaction. **(c)** The numbers of unique and shared host interactions between the homologous proteins of SARS-CoV-2 and SARS-CoV. AP-MS: affinity-purification coupled to mass spectrometry; MD: Macro domain; NSP: Non-structural protein.

To map the virus-host interactions to the functions of viral proteins, we have conducted an unprecedented study of total proteomes of A549 cells expressing 54 individual viral proteins, the “effectome” (Figure 1a; Supplementary Table 3). This dataset provides clear links between protein expression changes and virus-host interactions, as exemplified by ORF9b, which leads to a dysregulation of mitochondrial functions and binds to TOMM70, a known regulator of mitophagy^2,17^ (Figure 1b; Supplementary Tables 2, 3). Global pathway enrichment analysis of the effectome dataset confirmed such mitochondrial dysregulation by ORF9b of both viruses^2,18^ (Extended data Fig. 2c; Supplementary Table 3) and further highlighted virus-specific effects, as exemplified by the exclusive upregulation of proteins involved in cholesterol metabolism (CYP51A1, DHCR7, IDI1, SQLE) by SARS-CoV-2 NSP6. Intriguingly, cholesterol metabolism was recently shown to be implicated in SARS-CoV-2 replication and suggested as a promising target for drug development^19–21^. Beside perturbations at the pathway level, viral proteins specifically modulated single host proteins, possibly explaining more distinct molecular mechanisms involved in viral protein function. Focusing on the 180 most affected host proteins, we identified RCOR3, a putative transcriptional corepressor, as strongly upregulated by NSP4 of both viruses (Extended data Fig. 2d, 3a). Remarkably, the apolipoprotein B (APOB) was substantially regulated by ORF3 and NSP1 of SARS-CoV-2, suggesting its importance for SARS-CoV-2 biology (Extended data Fig. 3b).

### Multi-omics profiling of SARS-CoV-2 and SARS-CoV infection

While interactome and effectome provide in-depth information on the activity of individual viral proteins, we wished to directly study their concerted activities in the context of viral infection. To this end, we infected ACE2-expressing A549 cells (Extended data Fig. 4a, b) with SARS-CoV-2 and SARS-CoV, and profiled the impact of viral infection on Mrna expression, protein abundance, ubiquitination and phosphorylation in a time-resolved manner (Figure 2 a-b).

**Figure 2.**
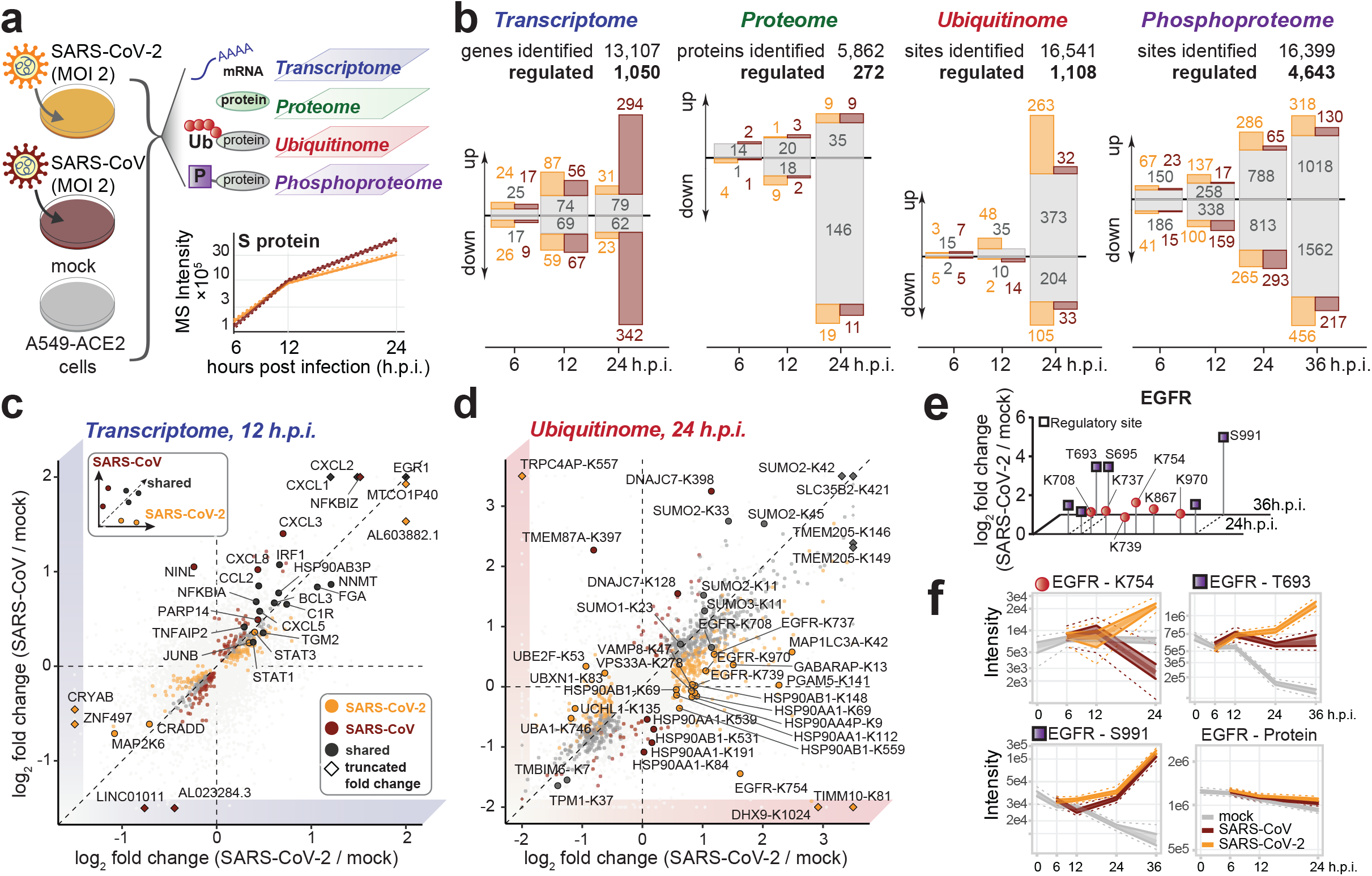
Multi-level profiling of SARS-CoV-2 and SARS-CoV infection. **(a)** Time-resolved profiling of parallel SARS-CoV-2 and SARS-CoV infection by multiple omics methods. The plot shows the MS intensity estimates for spike proteins of SARS-CoV-2 and SARS-CoV over time (n=4 independent experiments). **(b)** The numbers of distinct transcripts, proteins, ubiquitination and phosphorylation sites, significantly up- or downregulated at given time points after the infection (in comparison to the mock samples at the same time point). Color denotes transcripts/proteins/sites that are regulated similarly by SARS-CoV-2 and SARS-CoV infection (grey), or specifically by SARS-CoV-2 (orange) or SARS-CoV (brown). **(c-d)** Scatter plots comparing the host transcriptome and ubiquitinome respectively of SARS-CoV-2 (*x*-axis) and SARS-CoV (*y*-axis) infection at the indicated time after infection (log_2_ fold change in comparison to the mock infection samples at the same time point). Significantly regulated transcripts/sites (moderated t-test FDR-corrected two-sided p-value ≥ 0.05 (c), Bayesian linear model-based unadjusted two-sided p-value ≥ 10^−3^, |log_2_ fold change|≥ 0.5 (d), n=3 independent experiments), are colored according to their specificity in both infections. Diamonds indicate that the actual log_2_ fold change was truncated to fit into the plot. **(e)** Phosphorylation (purple square) and ubiquitination (red circle) sites on epidermal growth factor receptor (EGFR) regulated upon SARS-CoV-2 infection. The plot shows median log_2_ fold changes of site intensities compared to mock at 24 and 36 h.p.i. Regulatory sites are indicated with a thick black border. **(f)** Profile plots of time-resolved EGFR K754 ubiquitination, T693 and S991 phosphorylation, and total protein levels in SARS-CoV-2 or SARS-CoV-infected A549-ACE2 cells, with indicated median, 50% and 95% confidence intervals. n=3 (ubiquitination) or 4 (phosphorylation, total protein level) independent experiments. h.p.i.: hours post-infection.

In line with previous reports^9,22^, both SARS-CoV-2 and SARS-CoV share the ability to down-regulate type-I interferon response and activate a pro-inflammatory signature at transcriptome and proteome levels (Figure 2a-c, Extended data Fig. 4c-f, i, Supplementary Table 4, 8, Supplementary discussion 1). However, SARS-CoV elicited a more pronounced activation of the NFkB pathway, correlating with its higher replication rate and potentially explaining the reduced severity of pulmonary disease in case of SARS-CoV-2^23^ (Supplementary Tables 4, 5). In contrast, SARS-CoV-2 infection led to higher expression of FN1 and SERPINE1, which may be linked to the specific recruitment of TGFB factors (Figure 1b) and supporting regulation of TGF-β signaling by SARS-CoV-2.

To better understand the mechanisms underlying perturbation of cellular signaling, we performed comparative ubiquitination and phosphorylation profiling of SARS-CoV-2 and SARS-CoV infection. This analysis identified 1□108 of 16□541 detected ubiquitination sites to be differentially regulated by SARS-CoV-2 or SARS-CoV infection (Figure 2a, b, d, Extended data Fig. 5a; Supplementary Table 6). More than half of the significant sites were regulated in a similar manner by both viruses. These included sites on SLC35 and SUMO family proteins, indicating possible regulation of sialic acid transport and the process of SUMO-regulation itself. SARS-CoV-2 specifically increased ubiquitination on autophagy-related factors (MAP1LC3A, GABARAP, VPS33A, VAMP8) as well as particular sites on EGFR (*e*.*g*. K739, K754, K970). Sometimes the two viruses targeted distinct sites on the same cellular protein, as exemplified by HSP90 family members (HSP90AA1-K84, -K191 and -K539) (Figure 2d). Notably, a number of proteins (*e*.*g*. ALCAM, ALDH3B1, CTNNA1, EDF1 and SLC12A2) exhibited concomitant ubiquitination and a decrease at the protein level after infection, pointing to ubiquitination-mediated protein degradation (Figure 2d; Extended data Fig. 4f, 5a; Supplementary Tables 5, 6). Among these downregulated proteins, EDF1 has a pivotal role in the maintenance of endothelial integrity and may be a link to endothelial dysfunctions described for COVID-19^24,25^. Profound regulation of cellular signaling pathways was also observed at the phosphoproteomic level: among 16□399 total quantified phosphorylation sites, 4□□643 showed significant changes after SARS-CoV-2 or SARS-CoV infection (Extended data Fig. 5b, c; Supplementary Table 7). Highly regulated sites were identified for the proteins of the MAPK pathways (*e*.*g*. MAPKAPK2, MAP2K1, JUN, SRC) together with proteins involved in autophagy signaling (*e*.*g*. DEPTOR, RICTOR, OPTN, SQSTM1, LAMTOR1) and viral entry (*e*.*g*. ACE2, RAB7A) (Extended data Fig. 5b, d). Notably, RAB7A was recently shown to be an important host factor for SARS-CoV-2 infection that assists endosomal trafficking of ACE2 to the plasma membrane^26^. Simultaneously, we observed significantly higher phosphorylation at S72 of RAB7A in SARS-CoV-2 infection compared to SARS-CoV or mock, a site implicated in its intracellular localization and molecular association^27^. The regulation of known phosphosites suggests an involvement of central kinases (CDKs, AKT, MAPKs, ATM, and CHEK1) linked to cell survival, cell cycle progression, cell growth and motility, stress responses and the DNA damage response, which was also supported by the analysis of enriched motifs (Extended data Fig. 5e, f; Supplementary Tables 7 - 8). Notably, only SARS-CoV-2 but not SARS-CoV led to phosphorylation of the antiviral kinase EIF2AK2/PKR at the critical regulatory residue S33^28^. This differential activation of EIF2AK2/PKR could contribute to the difference in growth kinetics of the two SARS viruses (Supplementary Table 4, 5).

Our data clearly point to an interplay of phosphorylation and ubiquitination patterns on individual host proteins. EGFR, for instance, showed increased ubiquitination on six lysine residues at 24 hours post-infection (h.p.i.) accompanied by increased phosphorylation of T693, S695 and S991 after 24 and 36 hours (Figure 2e, f). Ubiquitination of all six lysine residues on EGFR was more pronounced upon SARS-CoV-2 infection. Moreover, vimentin, a central co-factor for coronavirus entry^29^ and pathogenicity^30,31^, displayed distinct phosphorylation and ubiquitination patterns on several sites early (*e*.*g*. S420) or late (*e*.*g*. S56, S72, K334) in infection (Extended data Fig. 6a, b). These discoveries underscore the value of testing different post-translational modifications simultaneously and suggest a concerted engagement of regulatory machineries to modify target protein’s functions and abundance.

### Phosphorylation and ubiquitination of functional domains of viral proteins

The majority of viral proteins were also post-translationally modified. Of the 27 detected SARS coronavirus proteins, 21 were ubiquitinated, among which N, S, NSP2, and NSP3 were the most frequently modified proteins in both viruses (Extended data Fig. 6c, Supplementary Table 6). Many of these ubiquitination sites were shared between the two viruses. Around half of the sites specifically regulated in either of the two viruses were conserved but differentially ubiquitinated, while the other half was encoded by either of the two pathogens, indicating that such acquired adaptations are also post-translationally modified and could recruit cellular proteins with appropriate functions (Figure 3a). Our interactome data identified several host E3 ligases (*e*.*g*. SARS-CoV-2 ORF3 with TRIM47, WWP1/2, STUB1; M and TRIM7; NSP13 and RING1) and deubiquitinating enzymes (*e*.*g*. SARS-CoV-2 ORF3 with USP8; ORF7a with USP34; SARS-CoV N with USP9X) and likely indicate a crosstalk between ubiquitination and viral protein functions (Figure 1b, Extended data Fig. 6d, Supplementary Table 2). Of particular interest are extensive ubiquitination events on the spike protein S of both viruses (K97, K528, K825, K835, K921 and K947) distributed on functional domains (N-terminal domain, C-terminal domain, fusion peptide and Heptad repeat 1 domain) potentially indicating critical regulatory functions that are conserved among the two viruses (Extended data Fig. 6e). Mapping of the phosphorylation events identified 5 SARS-CoV-2 (M, N, S, NSP3, ORF9b) and 8 SARS-CoV (M, N, S, NSP1, NSP2, NSP3, ORF3 and ORF9b) proteins to be phosphorylated (Extended data Fig. 6f, Supplementary Table 7), which corresponds to known recognition motifs. In particular, CAMK4 and MAPKAPK2 potentially phosphorylate sites on S and N, respectively. Inferred from phosphorylation of cellular proteins, the activities of these kinases were enriched in SARS-CoV-2 and SARS-CoV infected cells (Extended data Fig. 5e, f, 6e, g). Moreover, N proteins of both SARS coronaviruses recruit GSK3, which could potentially be linked to phosphorylation events on these viral proteins (Figure 1b, Extended data Fig. 6g, Supplementary Table 7). Particularly interesting are newly identified post-translationally modified sites located at functional domains of viral proteins. We identified SARS-CoV-2 N K338 ubiquitination and SARS-CoV-2/SARS-CoV N S310/311 phosphorylation (Extended data Fig. 6g). Mapping those sites to the atomic structure of the C-terminal domain (CTD)^32,33^ highlights critical positions for the functionality of the protein (Figure 3c, Extended data Fig. 6h, Supplementary discussion 2). Collectively, while the identification of differentially regulated sites may indicate pathogen-specific functions, insights gleaned from conserved post-translational modifications provide useful knowledge for the development of targeted pan-antiviral therapies.

**Figure 3.**
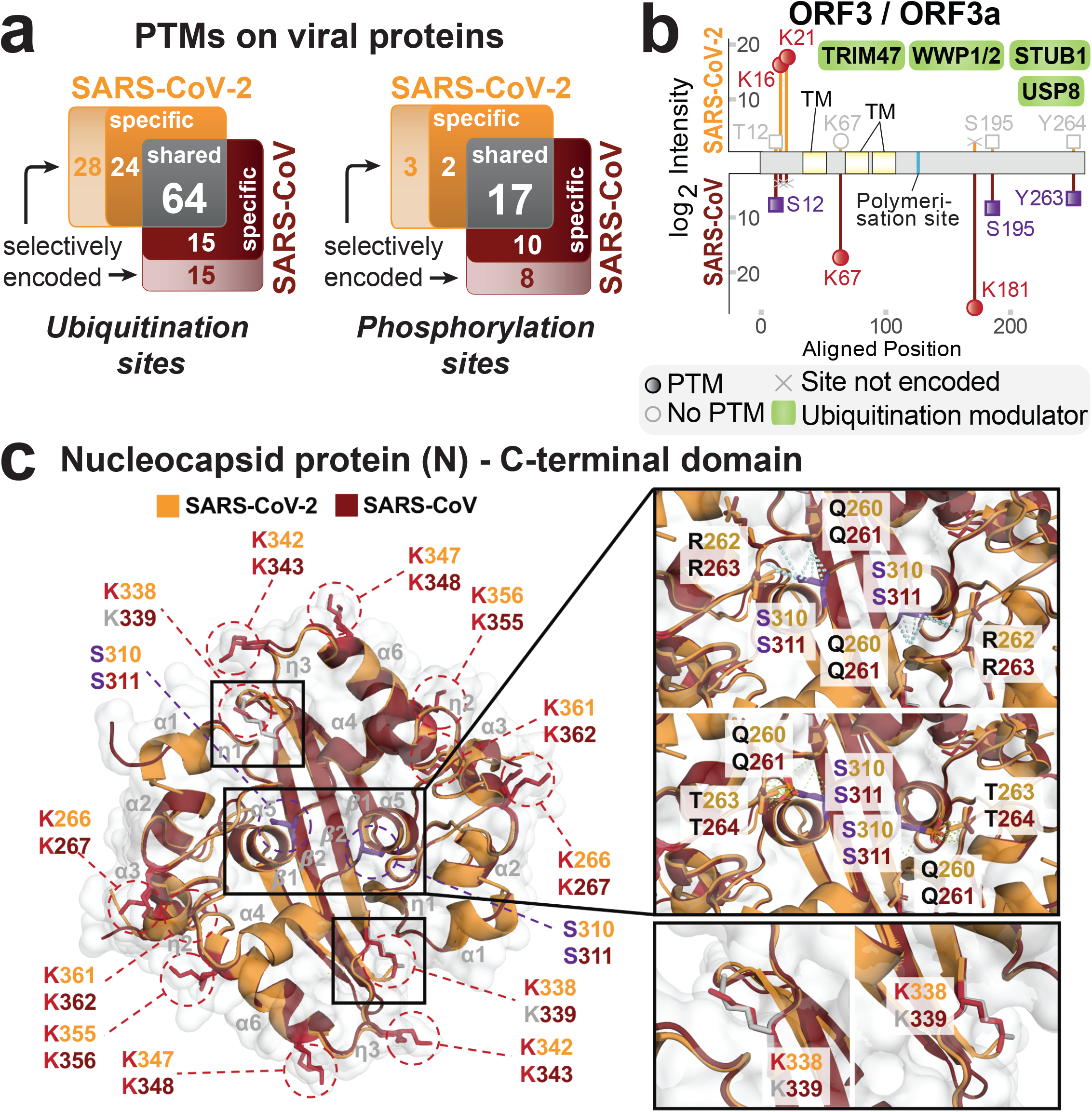
Integration of data from SARS-CoV-2 and SARS-CoV infection identifies coordinated regulation between omics layers. **(a)** Venn diagram presenting the distribution of all identified shared, differentially regulated and selectively encoded (sequence-specific) ubiquitination and phosphorylation sites on SARS-CoV-2 and SARS-CoV homologous proteins as measured after infection of A549-ACE2 cells. **(b)** Mapping of the ubiquitination (red circle) and phosphorylation (purple square) sites of SARS-CoV-2 ORF3 / SARS-CoV ORF3a proteins on their aligned sequence with median log_2_ intensities in A549-ACE2 cells infected with the respective virus at 24 h.p.i. Functional (blue) and topological (yellow) domains are mapped on each sequence. Binding of ubiquitin modifying enzymes to ORF3/ORF3a as identified in our AP-MS experiments (Extended data Fig. 1b) are indicated (green).. **(c)** Surface and ribbon representation of superimposed SARS-CoV (PDB: 2CJR, brown) and SARS-CoV-2 (PDB: 6YUN, orange) N CTD dimers (r.m.s.d. values of 0.492 Å for matching 108 Cα atoms). Side chains are colored in red, purple or grey as they belong to ubiquinated, phosphorylated or unmodified sites respectively. K338 ubiquitination site unique to SARS-CoV-2 is shown as close-up for both monomers (lower). Close-ups of inter-chain residue interactions established by non-phosphorylated (upper) and phosphorylated (center) SARS-CoV-2 S310/SARS-CoV S311. CTD: C-terminal domain; hACE2: binding site of human ACE2; FP: fusion peptide; HR1/2: Heptad region 1/2; CP: cytoplasmic region. CoV2 Cleav.: SARS-CoV-2 cleavage sites; r.m.s.d.: root-mean-square deviation.

### Integrative analysis highlights the perturbation of key cellular pathways

Our unified experimental design in a syngeneic system permitted direct time-resolved comparison of SARS-CoV-2 and SARS-CoV infection across different levels. Integrative pathway enrichment analysis demonstrated that both viruses largely perturb the same cellular processes at multiple levels albeit with varying temporal patterns (Extended data Fig. 7a). Transcriptional downregulation of proteins involved in tau-protein kinase activity and iron ions sequestration at 6 h.p.i., for instance, was followed by a decrease in protein abundance after 12 h.p.i. (Supplementary Table 8). RHO GTPase activation, mRNA processing and role of ABL in ROBO-SLIT signaling appeared to be regulated mostly through phosphorylation (Extended data Fig. 7a). In contrast, processes connected to cellular integrity such as the formation of senescence-associated heterochromatin foci, apoptosis-induced DNA fragmentation and amino acid transport across the plasma membrane were modulated through concomitant phosphorylation and ubiquitination events, providing insights into the molecular relationships of these post-translational modifications. Ion transporters, especially the SLC12 family (cation-coupled chloride cotransporters), previously identified as cellular factors in pulmonary inflammation^34^, were also regulated at multiple levels, evidenced by reduced protein abundance as well as differential post-translational modifications (Extended data Fig. 7a).

The pathway enrichment analysis provided a global and comprehensive picture of how SARS-CoV-2 and SARS-CoV affect the host. We next applied an automated approach to systematically explore the underlying molecular mechanisms contained in the viral interactome and effectome data. We mapped the measured interactions and effects of each viral protein onto the global network of cellular interactions^35^ and applied a network diffusion approach^36^ (Figure 4a). Such analysis utilizes known cellular protein-protein interactions, signaling and regulation events to identify connection points between the interactors of the viral protein and the proteins affected by its expression (Extended data Fig. 1b, 2d, Supplementary Tables 2, 3). The connections inferred from the real data were significantly shorter than for randomized data, confirming both the relevance of the approach and the data quality (Extended data Fig. 8a, b). Amongst many other findings, this approach pointed towards the potential mechanisms of autophagy regulation by ORF3 and NSP6; the modulation of innate immunity by M, ORF3 and ORF7b; and the Integrin-TGF-β-EGFR-RTK signaling perturbation by ORF8 of SARS-CoV-2 (Figure 4b, Extended data Fig. 8c, d). Enriching these subnetworks with SARS-CoV-2 infection-dependent mRNA abundance, protein abundance, phosphorylation and ubiquitination (Figure 4a) provided novel insights into the regulatory mechanisms employed by SARS-CoV-2. For instance, this analysis confirmed a role of NSP6 in autophagy^37^ and revealed the inhibition of autophagic flux by ORF3 protein, unique to SARS-CoV-2, leading to the accumulation of autophagy receptors (SQSTM1, GABARAP(L2), NBR1, CALCOCO2, MAP1LC3A/B, TAX1BP1), also observed in virus-infected cells (MAP1LC3B) (Figure 4c, Extended data Fig. 8e, f). This inhibition may be due to the interaction of the ORF3 protein with the HOPS complex (VPS11, -16, -18, -39, -41), which is essential for autophagosome-lysosome fusion, as well as by the differential phosphorylation of regulatory sites (*e*.*g*. on TSC2, mTORC1 complex, ULK1, RPS6, SQSTM1) and ubiquitination of key components (MAP1LC3A, GABARAP(L2), VPS33A, VAMP8) (Figure 4c, Extended data Fig. 8g). This inhibition of autophagosome function may have direct consequences for protein degradation. The abundance of APOB, a protein degraded *via* autophagy^38^, was selectively increased after SARS-CoV-2 infection or expression of the SARS-CoV-2 ORF3 (Extended data Fig. 3b, 8h). Accumulating APOB levels could exacerbate the risk of arterial thrombosis^39^, one of the main complications contributing to lung, heart and kidney failure in COVID-19 patients^40^. The inhibition of the IFN-α/β response observed at transcriptional and proteome levels was similarly explained by the network diffusion analysis (Extended data Fig. 8i), which implicated multiple proteins of SARS-CoV-2 in the disruption of antiviral immunity. Additional experiments functionally corroborated the inhibition of IFN-α/β induction or signaling by ORF3, ORF6, ORF7a, ORF7b, ORF9b (Extended data Fig. 8j). Upon virus infection, we observed the regulation of TGF-β and EGFR pathways modulating cell survival, motility and innate immune responses (Extended data Fig. 9a-d). Specifically, our network diffusion analysis revealed a connection between the binding of the ORF8 and ORF3 proteins to TGF-β-associated factors (TGFB1, TGFB2, LTBP1, TGFBR2, FURIN, BAMBI), the differential expression of ECM regulators (FERMT2, CDH1) and the virus-induced upregulation of fibrinogens (FGA, FGB), fibronectin (FN1) and SERPINE1 (Extended data Fig. 9a, b)^41^. The increased phosphorylation of proteins involved in MAPK (*e*.*g*. SHC1-S139, SOS1-S1134/1229, JUN-S63/S73, MAPKAPK2-T334, p38-T180/Y182) and receptor tyrosine kinase signaling (*e*.*g*. phosphorylation of PI3K complex members, PDPK1 (S241) and RPS6KA1 (S380)) as well as a higher expression of JUN, FOS and EGR1 are further indicative of TGF-β and EGFR pathways regulation (Extended data Fig. 9a, c, d). In turn, TGF-β and EGFR signaling are known to be potentiated by integrin signaling and activation of YAP-dependent transcription^42^, which we observed to be regulated in a time-dependent manner upon SARS-CoV-2 infection (Extended data Fig. 9a). Besides promoting virus replication, activation of these pathways has been implicated in fibrosis^13–15^, one of the hallmarks of COVID-19^16^.

**Figure 4.**
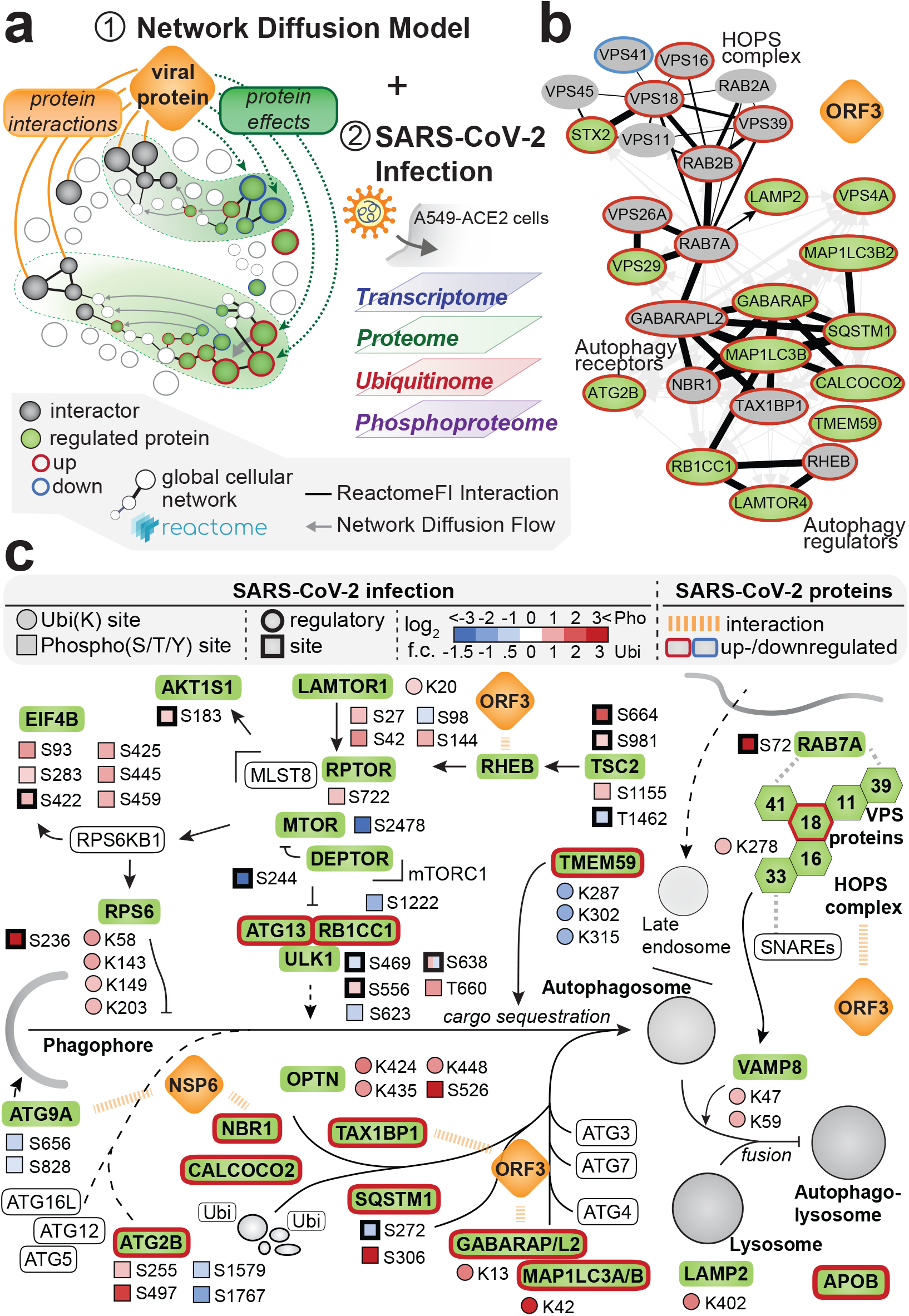
Network diffusion approach identifies molecular pathways linking protein-protein interactions with downstream changes in the host proteome. **(a)** Network diffusion approach to identify functional connections between the host targets of a viral protein and downstream proteome changes. The results of network diffusion are integrated with omics datasets of SARS coronavirus infection to streamline the identification of affected host pathways. **(b)** Subnetworks of the network diffusion predictions linking host targets of SARS-CoV-2 ORF3 to the factors involved in autophagy. The thickness of directed edges is proportional to the random walk transition probability. Black edges denote the connections present in ReactomeFI. **(c)** Overview of perturbations to host-cell autophagy induced by SARS-CoV-2. The pathway regulation is derived from the network diffusion model of SARS-CoV-2 ORF3 and NSP6 and overlaid with the changes in protein levels, ubiquitination and phosphorylation induced by SARS-CoV-2 infection.

### Data-guided drug testing reveals hotspots for antiviral therapies

Taken together, the viral-host protein-protein interactions and pathway regulations observed at multiple levels identify potential vulnerability points of SARS-CoV and SARS-CoV-2 that we decided to target by well-characterized selective drugs for antiviral therapies. To test antiviral efficacy, we established time-lapse fluorescent microscopy of SARS-CoV-2 GFP-reporter virus infection^43^. Inhibition of virus replication by IFN-α/β treatment corroborated previous conclusions that efficient SARS-CoV-2 replication involves an inactivation of this pathway at an early step and confirmed the reliability of this screening approach (Extended data Fig. 10a)^9,44^. We tested a panel of 48 drugs modulating the pathways perturbed by the virus for their effects on SARS-CoV-2 replication (Figure 5a, Supplementary Table 9). Notably, B-RAF (Sorafenib, Regorafenib, Dabrafenib), JAK1/2 (Baricitinib) and MAPK (SB 239063) inhibitors, which are commonly used to treat cancer and autoimmune diseases^45–47^ led to a significant increase of virus growth in our *in vitro* infection setting (Figure 5a, Extended data Fig. 10b, Supplementary Table 9). In contrast, inducers of DNA damage (Tirapazamine, Rabusertib) or a mTOR inhibitor (Rapamycin) led to suppression of virus growth. The highest antiviral activity was observed for Gilteritinib (a designated FLT3/AXL inhibitor), Ipatasertib (AKT inhibitor), Prinomastat and Marimastat (matrix metalloproteases (MMPs) inhibitors) (Figure 5a, b, Extended data Fig. 10c, Supplementary Table 9). These compounds profoundly inhibited replication of SARS-CoV-2 while having no or minor effects on cell growth (Extended data Fig. 10b, Supplementary Table 9). Quantitative PCR analysis indicated antiviral activities for Gilteritinib and Tirapazamine against SARS-CoV-2 and SARS-CoV (Figure 5c, Extended data Fig. 10d, e). Notably, Prinomastat and Marimastat, specific inhibitors of MMP-2 and MMP-9, showed selective activity against SARS-CoV-2 but not against SARS-CoV (Figure 5c, Extended data Fig. 10f, g). MMPs activities have been linked to TGF-β activation and pleural effusions, alveolar damage and neuroinflammation (*e*.*g*. Kawasaki disease), all of which are characteristics of COVID-19^23,48–51^.

**Figure 5.**
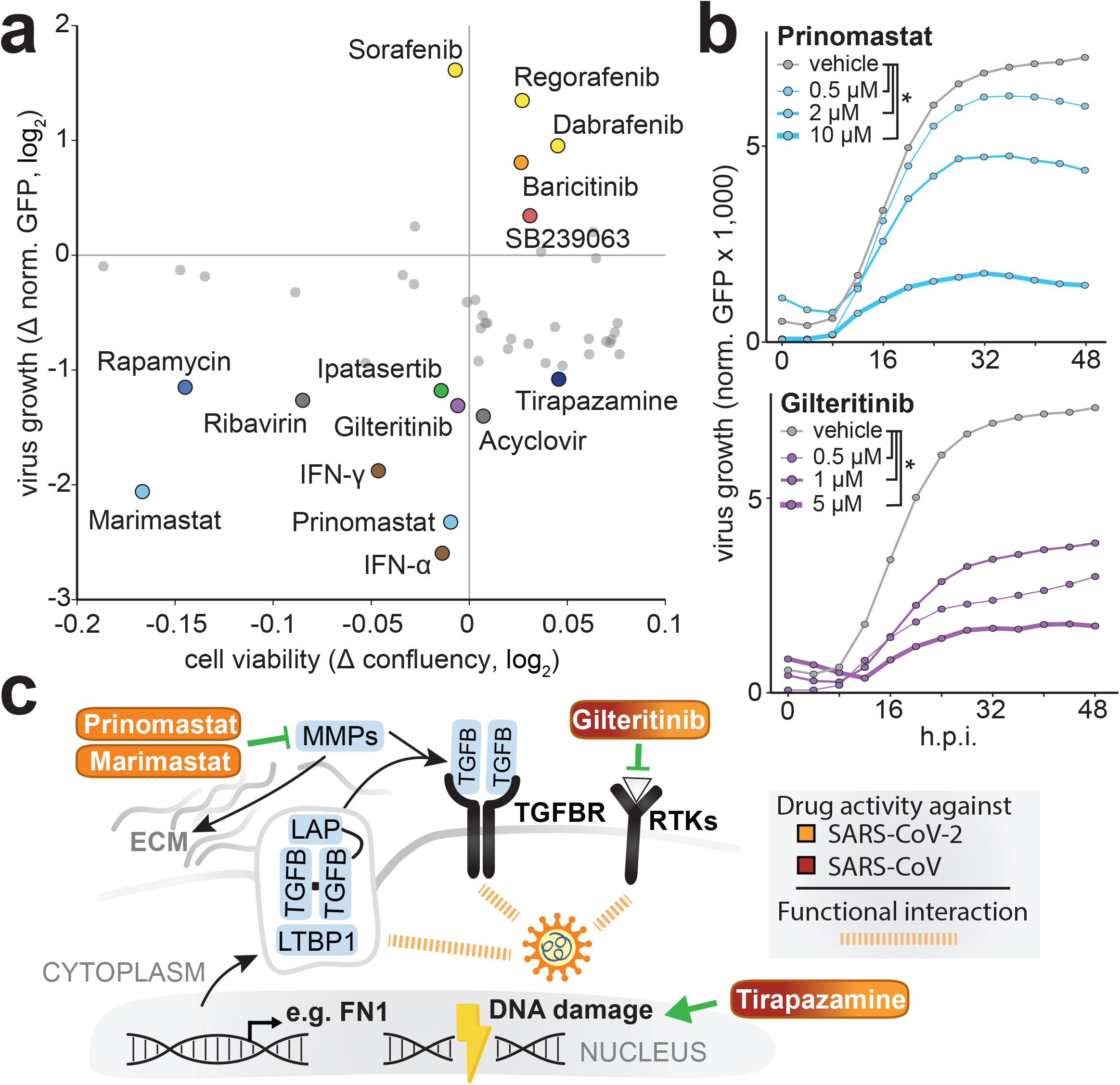
SARS-CoV-2-targeted pathways, as revealed by a multi-omics profiling approach, allow systematic testing of candidate antiviral therapies. **(a)** A549-ACE2 cells were treated with the indicated drugs 6 hours prior to infection with SARS-CoV-2-GFP (MOI 3). Scatter plot shows cell viability changes (*x*-axis, confluence log_2_ fold change in uninfected cells) and virus growth changes (*y*-axis, normalized GFP area log_2_ fold change in SARS-CoV-2-GFP-infected cells) of drug-treated in comparison to non-treated A549-ACE2 cells at 48 h.p.i. A confluence cutoff of −0.2 log_2_ fold change was applied to remove cytotoxic compounds. **(b)** shows time-courses of virus replication after Prinomastat or Gilteritinib pre-treatment. Asterisk indicates the significance in comparison to the control treatment (n=4 independent experiments, Wilcoxon test; unadjusted two-sided p-value ≥ 0.01). **(c)** Drugs potentially targeting pathways identified in our study. Color indicates antiviral activity against SARS-CoV-2/SARS-CoV (brown-orange gradient) or SARS-CoV-2 specifically (orange) as inferred from *in vitro* experiments. MOI: multiplicity of infection; h.p.i.: hours post-infection.

This drug screen demonstrates the value of our combined dataset that profiles SARS-CoV-2 infection at multiple levels. We hope that further exploration of these rich data by the scientific community and additional studies of the interplay between different omics levels will substantially advance our molecular understanding of coronaviruses biology, including the pathogenicity associated with specific human coronaviruses, such as SARS-CoV-2 and SARS-CoV. Moreover, this resource, together with complementary approaches by the community^26,52–54^, will streamline the search for antiviral compounds and serve as a base for rational design of combination therapies that target the virus from multiple synergistic angles, thus potentiating the effect of individual drugs while minimizing potential side-effects on healthy tissues.

## Supporting information

Supplementary_Table_1_List and sequences of coronavirus proteins

Supplementary_Table_2_Virus-host protein-protein interaction network of SARS-CoV-2 and SARS-CoV in A549 cells

Supplementary_Table_3_Total proteome of A549 cells expressing individual coronavirus proteins

Supplementary_Table_4_Transcriptome of A549-ACE2 cells infected with SARS-CoV-2 and SARS-CoV

Supplementary_Table_5_Total proteome of A549-ACE2 cells infected with SARS-CoV-2 and SARS-CoV

Supplementary_Table_6_Ubiquitinome of A549-ACE2 cells infected with SARS-CoV-2 and SARS-CoV

Supplementary_Table_7_Phosphoproteome of A549-ACE2 cells infected with SARS-CoV-2 and SARS-CoV

Supplementary_Table_8_Biological functions and pathways enriched in A549-ACE2 cells infected with SARS-CoV-2 and SARS-CoV

Supplementary_Table_9_Viral inhibitor Assay

Supplementary_Table_10_Intersection with other SARS-CoV-2 studies to date

Supplementary_Data_1_Diffusion networks of SARS-CoV-2 and SARS-CoV proteins

Supplementary Discussion

## Material and Methods

### Cell lines and reagents

HEK293T, A549, Vero E6 and HEK293-R1 cells and their respective culturing conditions were described previously^55^. All cell lines were tested to be mycoplasma-free. Expression constructs for C-terminal HA tagged viral ORFs were synthesized (Twist Bioscience and BioCat) and cloned into pWPI vector as described previously^56^ with the following modifications: starting ATG codon was added, internal canonical splicing sites were replaced with synonymous mutations and C-terminal HA-tag, followed by amber stop codon, was added to individual viral open reading frames. C-terminally hemagglutinin(HA)-tagged ACE2 sequence was amplified from an ACE2 expression vector (kindly provided by Stefan Pöhlmann)^57^ into the lentiviral vector pWPI-puro. A549 cells were transduced twice, and ACE2-expressing A549 (A549-ACE2) cells were selected with puromycin. Lentiviruses production, transduction of cells and antibiotic selection were performed as described previously^52^. RNA-isolation (Macherey-Nagel NucleoSpin RNA plus), reverse transcription (TaKaRa Bio PrimeScript RT with gDNA eraser) and RT-qPCR (Thermo-Fisher Scientific PowerUp SYBR green) were performed as described previously^54^. RNA-isolation for NGS applications was performed according to manufacturer’s protocol (Qiagen RNeasy mini kit, RNase free DNase set). For detection of protein abundance by western blotting, HA-HRP (Sigma-Aldrich; H6533; 1:2500 dilution), ACTB-HRP (Santa Cruz; sc-47778; 1:5000 dilution), MAP1LC3B (Cell Signaling; 3868; 1:1000 dilution), MAVS (Cell Signaling; 3993; 1:1000 dilution), HSPA1A (Cell Signaling; 4873; 1:1000 dilution), TGFβ (Cell Signaling; 3711; 1:1000 dilution), phospho-p38 (T180/Y182) (Cell Signaling; 4511; 1:1000 dilution), p38 (Cell Signaling; 8690; 1:1000 dilution) and SARS-CoV-2/SARS-CoV N protein (Sino Biological; 40143-MM05; 1:1000 dilution) antibodies were used Secondary antibodies detecting mouse (Cell Signaling; 7076; 1:5000 dilution/Jackson ImmunoResearch; 115-035-003; 1:5000 dilution), rat (Invitrogen; 31470; 1:5000 dilution), and rabbit IgG (Cell Signaling; 7074; 1:5000 dilution) were horseradish peroxidase (HRP)-coupled. For AP-MS and AP-WB applications, HA-beads (Sigma-Aldrich and Thermo Fisher Scientific) and Streptactin II beads (IBA Lifesciences) were used. WB imaging was performed as described previously^58^. For the stimulation of cells in the reporter assay, recombinant human IFN-α was a kind gift from Peter Stäheli, recombinant human IFN-γ were purchased from PeproTech and IVT4 was produced as described before^59^. All compounds tested during the viral inhibitor assay are listed in Supplementary Table 9.

### Virus strains, stock preparation, plaque assay and *in vitro* infection

SARS-CoV-Frankfurt-1, SARS-CoV-2-MUC-IMB-1 and SARS-CoV-2-GFP strains^43^ were produced by infecting Vero E6 cells cultured in DMEM medium (10% FCS, 100 ug/ml Streptomycin, 100 IU/ml Penicillin) for 2 days (MOI 0.01). Viral stock was harvested and spun twice (1000g/10min) before storage at −80°C. Titer of viral stock was determined by plaque assay. Confluent monolayers of VeroE6 cells were infected with serial five-fold dilutions of virus supernatants for 1 hour at 37□°C. The inoculum was removed and replaced with serum-free MEM (Gibco, Life Technologies) containing 0.5% carboxymethylcellulose (Sigma-Aldrich). Two days post-infection, cells were fixed for 20 minutes at room temperature with formaldehyde directly added to the medium to a final concentration of 5%. Fixed cells were washed extensively with PBS before staining with H2O containing 1% crystal violet and 10% ethanol for 20 minutes. After rinsing with PBS, the number of plaques was counted and the virus titer was calculated. A549-ACE2 cells were infected with either SARS-CoV-Frankfurt-1 or SARS-CoV-2-MUC-IMB-1 strains (MOI 2) for the subsequent experiments. At each time point, the samples were washed once with 1x TBS buffer and harvested in SDC lysis buffer (100 mM Tris HCl pH 8.5; 4% SDC) or 1x SSB lysis buffer (62.5 mM Tris HCl pH 6.8; 2% SDS; 10% glycerol; 50 mM DTT; 0.01% bromophenol blue) or RLT (Qiagen) for proteome-phosphoproteome-ubiquitinome, western blot, and transcriptome analyses, respectively. The samples were heat-inactivated and frozen at −80°C until further processing, as described in the following sections.

### Affinity purification and mass spectrometric analyses of SARS-CoV-2, SARS-CoV and HCoV-229E/NL63 proteins expressed in A549 cells

To determine the interactomes of SARS-CoV-2 and SARS-CoV and the interactomes of an accessory protein (encoded by ORF4/ORF4a of HCoV-229E or ORF3 of HCoV-NL63) that presumably represents a homolog of the ORF3 and ORF3a proteins of SARS-CoV-2 and SARS-CoV, respectively, four replicate affinity purifications were performed for each HA-tagged viral protein. A549 cells (6×10^6^ cells per 15-cm dish) were transduced with lentiviral vectors encoding HA-tagged SARS-CoV-2, SARS-CoV or HCoV-229E/NL63 proteins and protein lysates were prepared from cells harvested three days post-transduction. Cell pellets of two 15-cm dishes were lysed in lysis buffer (50 mM Tris-HCl pH 7.5, 100 mM NaCl, 1.5 mM MgCl_2_, 0.2% (v/v) NP-40, 5% (v/v) glycerol, cOmplete protease inhibitor cocktail (Roche), 0.5% (v/v) 750 U/µl Sm DNAse) and sonicated (5 min, 4°C, 30 sec on, 30 sec off, low settings; Bioruptor, Diagenode SA). Following normalization of protein concentrations of cleared lysates, virus protein-bound host proteins were enriched by adding 50 µl anti-HA-agarose slurry (Sigma-Aldrich, A2095) with constant agitation for 3 hours at 4°C. Non-specifically bound proteins were removed by four subsequent washes with lysis buffer followed by three detergent-removal steps with washing buffer (50 mM Tris-HCl pH 7.5, 100 mM NaCl, 1.5 mM MgCl_2_, 5% (v/v) glycerol). Enriched proteins were denatured, reduced, alkylated and digested by addition of 200 µl digestion buffer (0.6 M guanidinium chloride, 1 mM TCEP, 4 mM CAA, 100 mM Tris-HCl pH 8, 0.5 µg LysC (WAKO Chemicals), 0.5 µg trypsin (Promega) at 30°C overnight. Peptide purification on StageTips with three layers of C18 Empore filter discs (3M) and subsequent mass spectrometry analysis was performed as described previously^55,56^. Briefly, purified peptides were loaded onto a 20□cm reverse-phase analytical column (75□µm diameter; ReproSil-Pur C18-AQ 1.9□µm resin; Dr. Maisch) and separated using an EASY-nLC 1200 system (Thermo Fisher Scientific). A binary buffer system consisting of buffer A (0.1% formic acid in H_2_O) and buffer B (80% acetonitrile, 0.1% formic acid in H_2_O) with a 90 min gradient (5-30% buffer B (65 min), 30-95% buffer B (10 min), wash out at 95% buffer B (5 min), decreased to 5% buffer B (5 min), and 5% buffer B (5 min)) was used at a flow rate of 300 nl□per min. Eluting peptides were directly analysed on a Q-Exactive HF mass spectrometer (Thermo Fisher Scientific). Data-dependent acquisition included repeating cycles of one MS1 full scan (300–1□650□m/z, R□=□60□000 at 200□m/z) at an ion target of 3×10^6^, followed by 15 MS2 scans of the highest abundant isolated and higher-energy collisional dissociation (HCD) fragmented peptide precursors (R = 15□000 at 200□m/z). For MS2 scans, collection of isolated peptide precursors was limited by an ion target of 1×10^5^ and a maximum injection time of 25□ms. Isolation and fragmentation of the same peptide precursor was eliminated by dynamic exclusion for 20□s. The isolation window of the quadrupole was set to 1.4□m/z and HCD was set to a normalized collision energy of 27%.

### Proteome analyses of cells expressing SARS-CoV-2, SARS-CoV and HCoV-229E/NL63 proteins

For the determination of proteome changes in A549 cells expressing SARS-CoV-2, SARS-CoV or HCoV-229E/NL63 proteins, a fraction of 1×10^6^ lentivirus-transduced cells from the affinity purification samples were lysed in guanidinium chloride buffer (6 M GdmCl, 10 mM TCEP, 40 mM CAA, 100 mM Tris-HCl pH 8), boiled at 95°C for 8 min and sonicated (10 min, 4°C, 30 sec on, 30 sec off, high settings). Protein concentrations of cleared lysates were normalized to 50 µg and proteins were pre-digested with 1 µg LysC at 37°C for 1 hour followed by a 1:10 dilution (100 mM Tris-HCl pH 8) and overnight digestion with 1 µg trypsin at 30°C. Peptide purification on StageTips with three layers of C18 Empore filter discs (3M) and subsequent mass spectrometry analysis was performed as described previously^55,56^. Briefly, 300 ng of purified peptides were loaded onto a 50 cm reversed phase column (75 μm inner diameter, packed in house with ReproSil-Pur C18-AQ 1.9 μm resin [Dr. Maisch GmbH]). The column temperature was maintained at 60°C using a homemade column oven. A binary buffer system, consisting of buffer A (0.1% formic acid (FA)) and buffer B (80% ACN, 0.1% FA), was used for peptide separation, at a flow rate of 300 nl/min. An EASY-nLC 1200 system (Thermo Fisher Scientific), directly coupled online with the mass spectrometer (Q Exactive HF-X, Thermo Fisher Scientific) *via* a nano-electrospray source, was employed for nano-flow liquid chromatography. Peptides were eluted by a linear 80 min gradient from 5% to 30% buffer B (0.1% v/v formic acid, 80% v/v acetonitrile), followed by a 4 min increase to 60% B, a further 4 min increase to 95% B, a 4 min plateau phase at 95% B, a 4 min decrease to 5% B and a 4 min wash phase of 5% B. To acquire MS data, the data-independent acquisition (DIA) scan mode operated by the XCalibur software (Thermo Fisher) was used. DIA was performed with one full MS event followed by 33 MS/MS windows in one cycle resulting in a cycle time of 2.7 seconds. The full MS settings included an ion target value of 3×10^6^ charges in the 300 – 1□650 m/z range with a maximum injection time of 60 ms and a resolution of 120□000 at m/z 200. DIA precursor windows ranged from 300.5 m/z (lower boundary of first window) to 1□649.5 m/z (upper boundary of 33rd window). MS/MS settings included an ion target value of 3×10^6^ charges for the precursor window with an Xcalibur-automated maximum injection time and a resolution of 30□000 at m/z 200.

To generate the proteome library for DIA measurements purified peptides from the first and the fourth replicates of all samples were pooled separately and 25 µg of peptides from each pool were fractionated into 24 fractions by high pH reversed-phase chromatography as described earlier^60^. During each separation, fractions were concatenated automatically by shifting the collection tube every 120 seconds. In total 48 fractions were dried in a vacuum centrifuge, resuspended in buffer A* (0.2% TFA, 2% ACN) and subsequently analyzed by a top12 data-dependent acquisition (DDA) scan mode using the same LC gradient and settings. The mass spectrometer was operated by the XCalibur software (Thermo Fisher). DDA scan settings on full MS level included an ion target value of 3×10^6^ charges in the 300 – 1□650 m/z range with a maximum injection time of 20 ms and a resolution of 60□000 at m/z 200. At the MS/MS level the target value was 10^5^ charges with a maximum injection time of 60 ms and a resolution of 15□000 at m/z 200. For MS/MS events only, precursor ions with 2-5 charges that were not on the 20 s dynamic exclusion list were isolated in a 1.4 m/z window. Fragmentation was performed by higher-energy C-trap dissociation (HCD) with a normalized collision energy of 27eV.

### Infected time-course proteome-phosphoproteome-diGly proteome sample preparation

Frozen lysates of infected A549-ACE2 cells harvested at 6, 12 and 24 hours (also 36 hours only in phosphoproteomics study) post-infection were thawed on ice, boiled for 5 min at 95°C and sonicated for 15 min (Branson Sonifierer). Protein concentrations were estimated by tryptophan assay^61^. To reduce and alkylate proteins, samples were incubated for 5 min at 45°C with TCEP (10 mM) and CAA (40 mM). Samples were digested overnight at 37°C using trypsin (1:100 w/w, enzyme/protein, Sigma-Aldrich) and LysC (1:100 w/w, enzyme/protein, Wako). For proteome analysis, 10 µg of peptide material were desalted using SDB-RPS StageTips (Empore)^61^. Briefly, samples were diluted with 1% TFA in isopropanol to a final volume of 200 µl and loaded onto StageTips, subsequently washed with 200 µl of 1% TFA in isopropanol and 200 µl 0.2% TFA/ 2% ACN. Peptides were eluted with 75 µl of 1.25% Ammonium hydroxide (NH4OH) in 80% ACN and dried using a SpeedVac centrifuge (Eppendorf, Concentrator plus). They were resuspended in buffer A* (0.2% TFA/ 2% ACN) prior to LC-MS/MS analysis. Peptide concentrations were measured optically at 280 nm (Nanodrop 2000, Thermo Scientific) and subsequently equalized using buffer A*. 1µg peptide was subjected to LC-MS/MS analysis.

The rest of the samples was four-fold diluted with 1% TFA in isopropanol and loaded onto SDB-RPS cartridges (Strata™-X-C, 30 mg/ 3 ml, Phenomenex Inc), pre-equilibrated with 4 ml 30% MeOH/1% TFA and washed with 4 ml 0.2% TFA. Samples were washed twice with 4 ml 1% TFA in isopropanol, once with 0.2% TFA/ 2% ACN and eluted twice with 2 ml 1.25% NH4OH/ 80% ACN. Eluted peptides were diluted with ddH_2_O to a final ACN concentration of 35%, snap frozen and lyophilized.

For phosphopeptide enrichment, lyophilized peptides were resuspended in 105 µl of equilibration buffer (1% TFA/ 80% ACN) and the peptide concentration was measured optically at 280nm (Nanodrop 2000, Thermo Scientific) and subsequently equalized using equilibration buffer. The AssayMAP Bravo robot (Agilent) performed the enrichment for phosphopeptides (150µg) by priming AssayMAP cartridges (packed with 5 µl Fe(III)-NTA) with 0.1% TFA in 99% ACN followed by equilibration in equilibration buffer and loading of peptides. Enriched phosphopeptides were eluted with 1 % Ammonium hydroxide, which was evaporated by Speedvac’ing samples for 20 minutes. Dried peptides were resuspended in 6 µl buffer A* and 5 µl was subjected to LC-MS/MS analysis.

For diGly peptide enrichment, lyophilized peptides were reconstituted in IAP buffer (50 mM MOPS, pH 7.2, 10 mM Na_2_HPO_4_, 50 mM NaCl) and the peptide concentration was estimated by tryptophan assay. K-□-GG remnant containing peptides were enriched using the PTMScan® Ubiquitin Remnant Motif (K-□-GG) Kit (Cell Signaling Technology). Crosslinking of antibodies to beads and subsequent immunopurification was performed with slight modifications as previously described^62^. Briefly, two vials of crosslinked beads were combined and equally split into 16 tubes (∼31 µg of antibody per tube). Equal peptide amounts (600 µg) were added to crosslinked beads and the volume was adjusted with IAP buffer to 1 ml. After 1 hour of incubation at 4°C and gentle agitation, beads were washed twice with cold IAP and 5 times with cold ddH_2_O. Thereafter, peptides were eluted twice with 50 µl 0.15% TFA. Eluted peptides were desalted and dried as described for proteome analysis with the difference that 0.2% TFA instead of 1%TFA in isopropanol was used for the first wash. Eluted peptides were resuspended in 9 µl buffer A* and 4 µl was subjected to LC-MS/MS analysis.

### DIA Measurements

Samples were loaded onto a 50 cm reversed phase column (75 μm inner diameter, packed in house with ReproSil-Pur C18-AQ 1.9 μm resin [Dr. Maisch GmbH]). The column temperature was maintained at 60°C using a homemade column oven. A binary buffer system, consisting of buffer A (0.1% formic acid (FA)) and buffer B (80% ACN plus 0.1% FA) was used for peptide separation, at a flow rate of 300 nl/min. An EASY-nLC 1□200 system (Thermo Fisher Scientific), directly coupled online with the mass spectrometer (Orbitrap Exploris 480, Thermo Fisher Scientific) *via* a nano-electrospray source, was employed for nano-flow liquid chromatography. The FAIMS device was placed between the nanoelectrospray source and the mass spectrometer and was used for measurements of the proteome and the PTM-library samples. Spray voltage was set to 2□650 V, RF level to 40 and heated capillary temperature to 275°C. For proteome measurements we used a 100 min gradient starting at 5% buffer B followed by a stepwise increase to 30% in 80 min, 60% in 4 min and 95% in 4 min. The buffer B concentration stayed at 95% for 4 min, decreased to 5% in 4 min and stayed there for 4 min. The mass spectrometer was operated in data-independent mode (DIA) with a full scan range of 350-1□650 m/z at 120□000 resolution at 200 m/z, normalized automatic gain control (AGC) target of 300% and a maximum fill time of 28 ms. One full scan was followed by 22 windows with a resolution of 15□000, normalized automatic gain control (AGC) target of 1□000% and a maximum fill time of 25 ms in profile mode using positive polarity. Precursor ions were fragmented by higher-energy collisional dissociation (HCD) (NCE 30%). Each of the selected CVs (−40, −55 and −70) was applied to sequential survey scans and MS/MS scans; the MS/MS CV was always paired with the appropriate CV from the corresponding survey scan.

For phosphopeptide samples, 5 µl were loaded and eluted with a 70 min gradient starting at 3% buffer B followed by a stepwise increase to 19% in 40 min, 41% in 20 min, 90% in 5 min and 95% in 5 min. The mass spectrometer was operated in data-independent mode (DIA) with a full scan range of 300-1□400 m/z at 120□000 resolution at 200 m/z and a maximum fill time of 60 ms. One full scan was followed by 32 windows with a resolution of 30□000. Normalized automatic gain control (AGC) target and maximum fill time were set to 1□000% and 54 ms, respectively, in profile mode using positive polarity. Precursor ions were fragmented by higher-energy collisional dissociation (HCD) (NCE stepped 25-27.5-30%). For the library generation, we enriched A549 cell lysates for phosphopeptides and measured them with 7 different CV settings (−30, −40, −50, −60, −70, −80 or −90 V) using the same DIA method. The noted CVs were applied to the FAIMS electrodes throughout the analysis. For the analysis of K-□-GG peptide samples, half of the samples were loaded. We used a 120 min gradient starting at 3% buffer B followed by a stepwise increase to 7% in 6 min, 20% in 49 min, 36% in 39 min, 45% in 10 min and 95% in 4 min. The buffer B concentration stayed at 95% for 4 min, decreased to 5% in 4 min and stayed there for 4 min. The mass spectrometer was operated in data-independent mode (DIA) with a full scan range of 300-1□350 m/z at 120□000 resolution at m/z 200, normalized automatic gain control (AGC) target of 300% and a maximum fill time of 20 ms. One full scan was followed by 46 windows with a resolution of 30□000. Normalized automatic gain control (AGC) target and maximum fill time were set to 1□000% and 54 ms, respectively, in profile mode using positive polarity. Precursor ions were fragmented by higher-energy collisional dissociation (HCD) (NCE 28%). For K-□-GG peptide library, we mixed the first replicate of each sample and measured them with eight different CV setting (−35, −40, −45, −50, −55, −60, −70 or −80 V) using the same DIA method.

### Processing of raw MS data

#### AP-MS data

Raw MS data files of AP-MS experiments conducted in DDA mode were processed with MaxQuant (version 1.6.14) using the standard settings and label-free quantification enabled (LFQ min ratio count 1, normalization type none, stabilize large LFQ ratios disabled). Spectra were searched against forward and reverse sequences of the reviewed human proteome including isoforms (UniprotKB, release 2019.10) and C-terminally HA-tagged SARS-CoV-2, SARS-CoV and HCoV proteins by the built-in Andromeda search engine^63^.

In-house Julia scripts^64^ were used to define alternative protein groups: only the peptides identified in AP-MS samples were considered for being protein group-specific, protein groups that differed by the single specific peptide or had less than 25% different specific peptides were merged to extend the set of peptides used for protein group quantitation and reduce the number of protein isoform-specific interactions.

#### Viral protein overexpression DIA MS data

Spectronaut version 13 (Biognosys) with the default settings was used to generate the proteome libraries from DDA runs by combining files of respective fractionations using the human fasta file (Uniprot, 2019.10, 42□431 entries) and viral bait sequences. Proteome DIA files were analyzed using the proteome library with the default settings and disabled cross run normalization.

#### SARS-CoV-2/SARS-CoV-infected proteome/PTM DIA MS data

Spectronaut version 14 (Biognosys)^65^ was used to generate the libraries and analyze all DIA files using the human fasta file (UniprotKB, release 2019.10) and sequences of SARS-CoV-2/SARS-CoV proteins (UniProt, release 2020.08). Orf1a polyprotein sequences were split into separate protein chains according to the cleavage positions specified in the UniProt. For the generation of the PTM-specific libraries, the DIA single CV runs were combined with the actual DIA runs and either phosphorylation at Serine/Threonine/Tyrosine or GlyGly at Lysine was added as variable modification to default settings. Maximum number of fragment ions per peptide was increased to 25. The proteome DIA files were analyzed using direct DIA approach with default settings and disabled cross run normalization. All PTM DIA files were analyzed using their respective hybrid library and either phosphorylation at Serine/Threonine/Tyrosine or GlyGly at Lysine was added as an additional variable modification to default settings with LOESS normalization and disabled PTM localization filter.

A collection of in-house Julia scripts^64^ were used to process the elution group (EG)-level Spectronaut reports, identify PTMs and assign EG-level measurements to PTMs. The PTM was considered if at least once it was detected with ≥ 0.75 localization probability in EG with q-value ≤ 10^−3^. For further analysis of given PTM, only the measurements with ≥ 0.5 localization probability and EG q-value ≤ 10^−2^ were used.

### Bioinformatic analysis

Unless otherwise specified, the bioinformatic analysis was done in R (version 3.6), Julia (version 1.5) and Python (version 3.8) using a collection of in-house scripts^64,66^.

#### Statistical analysis of MS data

MaxQuant and Spectronaut output files were imported into R using in-house maxquantUtils R package^67^. For all MS datasets, the Bayesian linear random effects models were used to define how the abundances of proteins change between the conditions. To specify and fit the models we employed msglm R package^68^, which utilizes rstan package (version 2.19)^69^ for inferring the posterior distribution of the model parameters. In all the models, the effects corresponding to the experimental conditions have regularized horseshoe+ priors^70^, while the batch effects have normally distributed priors. Laplacian distribution was used to model the instrumental error of MS intensities. For each MS instrument used, the heteroscedastic intensities noise model was calibrated with the technical replicate MS data of the instrument. These data were also used to calibrate the logit-based model of missing MS data (the probability that the MS instrument will fail to identify the protein given its expected abundance in the sample). The model was fit using unnormalized MS intensities data. Instead of transforming the data by normalization, the inferred protein abundances were scaled by the normalization multiplier of each individual MS sample to match the expected MS intensity of that sample. This allows taking the signal-to-noise variation between the samples into account when fitting the model. Due to high computational intensity, the model was applied to each protein group separately. For all the models, 4□000 iterations (2□000 warmup + 2□000 sampling) of the No-U-Turn Markov Chain Monte Carlo were performed in 7 or 8 independent chains, every 4th sample was collected for posterior distribution of the model parameters. For estimating the statistical significance of protein abundance changes between the two experimental conditions, the p-value was defined as the probability that a random sample from the posterior distribution of the first condition would be smaller (or larger) than a random sample drawn from the second condition. No multiple hypothesis testing corrections were applied, since this is handled by the choice of the model priors.

#### Statistical analysis of AP-MS data and filtering for specific interactions

The statistical model was applied directly to the MS1 intensities of protein group-specific LC peaks (*evidence*.*txt* table of MaxQuant output). In R GLM formula language, the model could be specified as

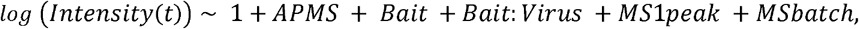

where *APMS* effect models the average shift of intensities in AP-MS data in comparison to full proteome samples, *Bait* is the average enrichment of a protein in AP-MS experiments of homologous proteins of both SARS-CoV and SARS-CoV-2, and *Bait:Virus* corresponds to the virus-specific changes in protein enrichment. *MS1peak* is the log-ratio between the intensity of a given peak and the total protein abundance (the peak is defined by its peptide sequence, PTMs and the charge; it is assumed that the peak ratios do not depend on experimental conditions^71^), and *MSbatch* accounts for batch-specific variations of protein intensity. *APMS, Bait* and *Bait:Virus* effects were used to reconstruct the batch effect-free abundance of the protein in AP-MS samples.

The modeling provided the enrichment estimates for each protein in each AP experiment. Specific AP-MS interactions had to pass the two tests. In the first test, the enrichment of the candidate protein in a given bait AP was compared against the background, which was dynamically defined for each interaction to contain the data from all other baits, where the abundance of the candidate was within 50%-90% percentile range (excluding top 10% baits from the background allowed the protein to be shared by a few baits in the resulting AP-MS network). The non-targeting control and Gaussia luciferase baits were always preserved in the background. Similarly, to filter out any potential side-effects of very high bait protein expression, the ORF3 homologs were always present in the background of M interactors and vice versa. To rule out the influence of the batch effects, the second test was applied. It was defined similarly to the first one, but the background was constrained to the baits of the same batch, and 40%-80% percentile range was used. In both tests, the protein has to be 4 times enriched against the background (16 times for highly expressed baits: ORF3, M, NSP13, NSP5, NSP6, ORF3a, ORF7b, ORF8b, HCoV-229E ORF4a) with the p-value ≤ 10^−3^.

Additionally, we excluded the proteins that, in the viral protein expression data, have shown upregulation, and their enrichment in AP-MS data was less than 16 times stronger than observed upregulation effects. Finally, to exclude the carryover of material between the samples sequentially analyzed by MS, we removed the putative interactors, which were also enriched at higher levels in the samples of the preceding bait, or the one before it. For the analysis of interaction specificity between the homologous viral proteins, we estimated the significance of interaction enrichment difference (corrected by the average difference between the enrichment of the shared interactors to adjust for the bait expression variation). Specific interactions have to be 4 times enriched in comparison to the homolog with p-value ≤ 10^−3^.

#### Statistical analysis of DIA proteome effects upon viral protein overexpression

The statistical model of the viral protein overexpression data set was similar to AP-MS data, except that protein-level intensities provided by Spectronaut were used. The PCA analysis of the protein intensities has identified that the 2nd principal component is associated with the batch-dependent variations between the samples. To exclude their influence, this principal component was added to the experimental design matrix as an additional batch effect.

As with AP-MS data, the two statistical tests were used to identify the significantly regulated proteins (column “is_change” in Supplementary Table 3). First, the absolute value of median log_2_-fold change of the protein abundance upon overexpression of a given viral protein in comparison to the background had to be above 1.0 with p-value ≤ 10^−3^. The background was individually defined for each analyzed protein. It was composed of experiments, where the abundance of given protein was within the 20%-80% percentile range of all measured samples. Second, the protein had to be significantly regulated (same median log_2_-fold change and p-value thresholds applied) against the batch-specific background (defined similarly to the global background, but using only the samples of the same batch).

An additional stringent criterion was applied to select the most significant changes (column “is_top_change” in Supplementary Table 3; Extended data Fig. 1i).

For each protein we classified bait-induced changes as:

- “high” when |median log_2_ fold-change|≥ 1 and p-value ≤ 10^−10^ both in background and batch comparisons
- “medium” if 10^−10^< p-value ≤ 10^−4^ with same fold-change requirement and
- “low” if 10^−4^ < p-value ≤ 10^−2^ with same fold-change requirement,

all other changes were considered non-significant.

We then required that “shared” top-regulated proteins should have exactly one pair of SARS-CoV-2/SARS-CoV “high”- or “medium”-significant homologous baits among the baits with either up- or downregulated changes and no other baits with significant changes of the same-type.

We further defined “SARS-CoV-2-specific” or “SARS-CoV-specific” top-regulated proteins to be the ones with exactly one “high”-significant change, and no other significant changes of the same sign. For “specific” hits we additionally required that in comparison of “high”-significant bait to its homolog |median log_2_ fold-change|≥ 1 and p-value ≤ 10^−3^. When the homologous bait was missing (SARS-CoV-2 NSP1, SARS-CoV ORF8a and SARS-CoV ORF8b), we instead required that in the comparison of the “high”-significant change to the background |median log_2_ fold-change|≥ 1.5.

The resulting network of most affected proteins was imported and prepared for publication in Cytoscape v.3.8.1^72^.

#### Statistical analysis of DIA proteomic data of SARS-CoV-2 and SARS-CoV-infected A549-ACE2 cells

Similarly to the AP-MS DDA data, the linear Bayesian model was applied to the elution group (EG) level intensities. To model the protein intensity, the following linear model (in R notation) was used:

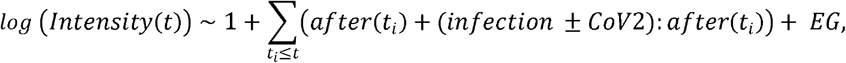

where

- *after(t*_*i*_) effect corresponds to the protein abundance changes in mock-infected samples that happened between *t*_*i-1*_ and *t*_*i*_ h.p.i. and it is applied to the modeled intensity at all time points starting from *t*_*i*_;
- *infection:after(t*_*i*_) (*t*_*i*_=6, 12, 24) is the common effect of SARS-CoV-2 & SARS-CoV infections occurred between *t*_*i-1*_ and *t*_*i*_;
- *CoV2:after(t*_*i*_) is the virus-specific effect within *t*_*i-1*_ and *t*_*i*_ h.p.i. that is added to the log intensity for SARS-CoV-2-infected samples and subtracted from the intensity for SARS-CoV ones;
- *EG* is the elution group-specific shift in the measured log-intensities.

The absolute value of median log_2_ fold change between the conditions above 0.25 and the corresponding unadjusted p-value ≤ 10^−3^ were used to define the significant changes at a given time point in comparison to mock infection. We also required that the protein group is quantified in at least two replicates of at least one of the compared conditions. Additionally, if for one of the viruses (*e*.*g*. SARS-CoV-2) only the less stringent condition (|median log_2_ fold-change|≥ 0.125, p*-*value ≤ 10^−2^) was fulfilled, but the change was significant in the infection of the other virus (SARS-CoV), and the difference between the viruses was not significant, the observed changes were considered significant for both viruses.

#### Statistical analysis of DIA phosphoproteome and ubiquitinome data of SARS-CoV-2 and SARS-CoV infections

The data from single-double- and triple-modified peptides were analyzed separately and, for a given PTM, the most significant result was reported.

The data was analyzed with the same Bayesian linear model as proteome SARS-CoV/-2 infection data. In addition to the intensities normalization, for each replicate sample the scale of the effects in the experimental design matrix was adjusted, so that on average the correlation between log fold-changes of the replicates was 1:1. The same logic as for the proteome analysis, was applied to identify significant changes, but the median log_2_ fold change had to be larger than 0.5, or 0.25 for the less stringent test. We additionally required that the PTM peptides are quantified in at least two replicates of at least one of the compared conditions. To ignore the changes in PTM site intensities that are due to proteome-level regulation, we excluded PTM sites on significantly regulated proteins if the direction of protein and PTM site changes was the same and the difference between their median log_2_ fold changes was less than 2. Phosphoproteomics data were further analyzed with Ingenuity Pathway Analysis software (QIAGEN Inc., https://www.qiagenbioinformatics.com/products/ingenuity-pathway-analysis)

#### Transcriptomic analysis of SARS-CoV-2 and SARS-CoV infected A549-ACE2 cells

As for the analysis of the transcriptome data, Gencode gene annotations v28 and the human reference genome GRCh38 were derived from the Gencode homepage (EMBL-EBI). Viral genomes were derived from GenBank (SARS-CoV-2 - LR824570.1, and SARS-CoV - AY291315.1). Dropseq tool v1.12 was used for mapping raw sequencing data to the reference genome. The resulting UMI filtered count matrix was imported into R v3.4.4. CPM (counts per million) values were calculated for the raw data and genes having a mean cpm value less than 1 were removed from the dataset. A dummy variable combining the covariates infection status (mock, SARS-CoV, SARS-CoV-2) and time point was used for modeling the data within Limma (v3.46.0)^73^.

Data was transformed with the Voom method^73^ followed by quantile normalization. Differential testing was performed between infection states at individual timepoints by calculating moderated t-statistics and p-values for each host gene. A gene was considered to be significantly regulated if the FDR adjusted p-value was below 0.05. The data for this study have been deposited in the European Nucleotide Archive (ENA) at EMBL-EBI under accession number PRJEB38744.

#### Gene Set Enrichment Analysis

We have used Gene Ontology, Reactome and other EnrichmentMap gene sets of human proteins (version 2020.10)^74^ as well as protein complexes annotations from IntAct Complex Portal (version 2019.11)^75^ and CORUM (version 2019)^76^. PhosphoSitePlus (version 2020.08) was used for known kinase-substrate and regulatory sites annotations, Perseus (version 1.6.14.0)^77^ was used for annotation of known kinase motifs. For transcription factor enrichment analysis (Extended data Fig. 2e) the significantly regulated transcripts were submitted to ChEA3 web-based application^78^ and ENCODE data on transcription factor– target gene associations were used^79^.

To find the nonredundant collection of annotations describing the unique and shared features of multiple experiments in a dataset (Figure 1d, Extended data Fig. 2l, m), we have used in-house Julia package OptEnrichedSetCover.jl^80^, which employs evolutionary multi-objective optimization technique to find a collection of annotation terms that have both significant enrichments in the individual experiments and minimal pairwise overlaps.

The resulting set of terms was further filtered by requiring that the annotation term has to be significant with the specified unadjusted Fisher’s Exact Test p-value cutoff at least in one of the experiments or comparisons (the specific cutoff value is indicated in the figure legend of the corresponding enrichment analysis).

The generation of diagonally-split heatmaps was done with VegaLite.jl package (https://github.com/queryverse/VegaLite.jl).

#### Viral PTMs alignment

For matching the PTMs of SARS-CoV-2 and SARS-CoV the protein sequences were aligned using the BioAlignments.jl Julia package (v.2.0, https://github.com/BioJulia/BioAlignments.jl). Needleman-Wunsch algorithm with BLOSUM80 substitution matrix, −5 and −3 penalties for the gap and extension, respectively. As for the cellular proteins, we required that the viral phosphorylation or ubiquitination site is observed with q-value ≤ 10^−3^ and localization probability ≥ 0.75. For the PTMs with lower confidence (q-value ≤ 10^−2^ and localization probability ≥ 0.5) we required that the same site is observed with high confidence at the matching position of the orthologous protein of the other virus.

#### Network diffusion analysis

To systematically detect functional interactions, which may connect the cellular targets of each viral protein (interactome dataset) with the downstream changes it induces on proteome level (effectome dataset), we have used the network diffusion-based HierarchicalHotNet method^36^ as implemented in Julia package HierarchicalHotNet.jl^81^. Specifically, for network diffusion with restart, we used the ReactomeFI network (version 2019)^35^ of cellular functional interactions, reversing the direction of functional interaction (*e*.*g*. replacing kinase→substrate interaction with substrate→kinase). The proteins with significant abundance changes upon bait overexpression (|median(log_2_ fold change)|≥ 0.25, p-value ≥ 10^−2^ both in the comparison against the controls and against the baits of the same batch) were used as the sources of signal diffusion with weights set to 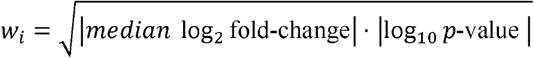,otherwise the node weight was set to zero. The weight of the edge *g*_*i*_ *⍰ g*_*j*_ was set to *w* _*i,j*_ =1 +*w*_*j*_ The restart probability was set to 0.4, as suggested in the original publication, so that the probability of the random walk to stay in the direct neighborhood of the node is the same as the probability to visit more distant nodes. To find the optimal cutting threshold of the resulting hierarchical tree of strongly connected components (SCCs) of the weighted graph corresponding to the stationary distribution of signal diffusion and to confirm the relevance of predicted functional connections, the same procedure was applied to 1□000 random permutations of vertex weights as described in Reyna *et al*.^36^ (vertex weights are randomly shuffled between the vertices with similar in- and out-degrees). Since cutting the tree of SCCs at any threshold *t* (keeping only the edges with weights above *t*) and collapsing each resulting SCC into a single node produces the directed acyclic graph of connections between SCCs, it allowed efficient enumeration of the paths from the “source” nodes (proteins strongly perturbed by viral protein expression with vertex weight *w, w* ≥ *1*.*5*) to the “sink” nodes (interactors of the viral protein). At each threshold *t*, the average inverse of the path length from source to sink nodes was calculated as:

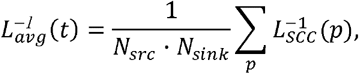

where *N*_*src*_ is the number of “sources”, *N*_*sink*_ is the number of “sinks”, *L*_*SCC*_*(p)* is the number of SCCs that the given path *p* from source to sink goes through, and the sum is for all paths from sources to sinks. The metric changes from 1 (all sources and sinks in the same SCC) to 0 (no or infinitely long paths between sources and sinks). For the generation of the diffusion networks we were using the *t*_*opt*_ threshold that maximized the difference between 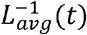 for the real data and the third quartile of 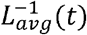 for randomly shuffled data.

In the generated SCC networks, the direction of the edges was reverted back, and the results were exported as GraphML files using in-house Julia scripts^64^. The catalogue of the networks for each viral bait is available as Supplementary Data 1.

To assess the significance of edges in the resulting network, we calculated the p-value of the edge *g*_*i*_*⍰g*_*j*_ as the probability that the permuted data-based transition probability between the given pair of genes is higher than the real data-based one:

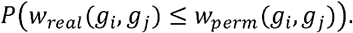

This p-value was stored as the “prob_perm_walkweight_greater” edge attribute of GraphML output. The specific subnetworks predicted by the network diffusion (Figure 4b-d) were filtered for edges with p-value ≤ 0.05.

When the *g*_*i*_*⍰ g*_*j*_ connection was not present in the ReactomeFI network, to recover the potential short pathways connecting *g*_*i*_ and *g*_*j*_, ReactomeFI was searched for intermediate *g*_*k*_ nodes, such that the edges *g*_*i*_*⍰g*_*k*_ and *g*_*i*_*⍰g*_*j*_ are present in ReactomeFI. The list of these short pathways is provided as “flowpaths” edge attribute in GraphML output.

The GraphML output of network diffusion was prepared for publication using yEd (v.3.20, www.yworks.com).

#### Intersection with other SARS coronavirus datasets

The intersection between the data generated by this study and other publicly available datasets was done using the information from respective supplementary tables. When multiple viruses were used in a study, only the comparisons with SARS-CoV and SARS-CoV-2 were included. For time-resolved data, all time points up to 24 h.p.i. were considered. The dataset coverage was defined as the number of reported distinct protein groups for proteomic studies and genes for transcriptomic studies. Confident interactions/significant regulations were filtered according to the criteria specified in the original study. A hit was considered as “confirmed” when it was significant both in this and external data and showed the same trend.

### qRT-PCR analysis

RNA isolation from SARS-CoV and SARS-CoV-2 infected A549-ACE2 cells was performed as described above (Qiagen). 500 ng total RNA was used for reverse transcription with PrimeScript RT with gDNA eraser (Takara). For relative transcript quantification PowerUp SYBR Green (Applied Biosystems) was used. Primer sequences can be provided upon request.

### Co-immunoprecipitation and western blot analysis

HEK293T cells were transfected with pWPI plasmid encoding single HA-tagged viral proteins, alone or together with pTO-SII-HA expressing host factor of interest. 48 hours after transfection, cells were washed in PBS, flash frozen in liquid nitrogen and kept at −80°C until further processing. Co-immunoprecipitation experiments were performed as described previously^55,56^. Briefly, cells were lysed in lysis buffer (50 mM Tris-HCl pH 7.5, 100 mM NaCl, 1.5 mM MgCl_2_, 0.2% (v/v) NP-40, 5% (v/v) glycerol, cOmplete protease inhibitor cocktail (Roche), 0.5% (v/v) 750 U/µl Sm DNAse) and sonicated (5 min, 4°C, 30 sec on, 30 sec off, low settings; Bioruptor, Diagenode SA). HA or Streptactin beads were added to cleared lysates and samples were incubated for 3 hours at 4°C under constant rotation. Beads were washed six times in the lysis buffer and resuspended in 1x SDS sample buffer 62,5 mM Tris-HCl pH 6.8, 2% SDS, 10% glycerol, 50 mM DTT, 0.01% bromophenol blue). After boiling for 5 minutes at 95°C, a fraction of the input lysate and elution were loaded on NuPAGE™ Novex™ 4-12% Bis-Tris (Invitrogen), and further submitted to western blotting using Amersham Protran nitrocellulose membranes. Imaging was performed by HRP luminescence (ECL, Perkin Elmer).

SARS-CoV-2 infected A549-ACE2 cell lysates were sonicated (10 min, 4°C, 30 sec on, 30 sec off, low settings; Bioruptor, Diagenode SA). Protein concentration was adjusted based on Pierce660 assay supplemented with ionic detergent compatibility reagent. After boiling for 5 min at 95°C and brief max g centrifugation, the samples were loaded on NuPAGE™ Novex™ 4-12% Bis-Tris (Invitrogen), and blotted onto 0,22 µm Amersham^™^ Protran^®^ nitrocellulose membranes (Merck). Primary and secondary antibody stainings were performed according to the manufacturer’s recommendations. Imaging was performed by HRP luminescence using Femto kit (ThermoFischer Scientific) or Western Lightning PlusECL kit (Perkin Elmer).

### Mapping of identified post-translational modification sites on the C-terminal domain structure of the Nucleocapsid protein

N CTD dimers of SARS-CoV-2 (PDB: 6YUN) and SARS-CoV (PDB: 2CJR) were superimposed by aligning the α-carbons backbone over 111 residues (from position 253/254 to position 364/365 following SARS-CoV-2/SARS-CoV numbering) by using the tool MatchMaker^82^ as implemented in the Chimera software^83^. Ubiquitination sites were visually inspected and mapped by using the PyMOL software (https://pymol.org). Phosphorylation on Ser310/311 was simulated in silico by using the PyTMs plugin as implemented in PyMOL^84^. Inter-chain residue contacts, dimer interface area, free energy and complex stability were comparatively analyzed between non-phosphorylated and phosphorylated SARS-CoV-2 and SARS-CoV N CTD by using the PDBePISA server^85^. Poisson– Boltzmann electrostatic surface potential of native and post-translationally modified N CTD was calculated by using the PBEQ Solver tool on the CHARMM-GUI server by preserving existing hydrogen bonds^86^. Molecular graphics depictions were produced with the PyMOL software.

### Reporter Assay and IFN Bioassay

The following reporter constructs were used in this study: pISRE-luc was purchased from Stratagene, EF1-α-ren from Engin Gürlevik, pCAGGS-Flag-RIG-I from Chris Basler, pIRF1-GAS-ff-luc, pWPI-SMN1-flag and pWPI-NS5 (ZIKV)-HA was described previously^56,87^.

For the reporter assay, HEK293-R1 cells were plated in 24-well plates 24 hours prior to transfection. Firefly reporter and Renilla transfection control were transfected together with plasmids expressing viral proteins using polyethylenimine (PEI, Polysciences) for untreated and treated conditions. In 18 hours cells were stimulated for 8 hours with a corresponding inducer and harvested in the passive lysis buffer (Promega). Luminescence of Firefly and Renilla luciferases was measured using dual-luciferase-reporter assay (Promega) according to the manufacturer’s instructions in a microplate reader (Tecan).

Total amounts of IFN-α/β in cell supernatants were measured by using 293T cells stably expressing the firefly luciferase gene under the control of the mouse Mx1 promoter (Mx1-luc reporter cells)^88^. Briefly, HEK293-R1 cells were seeded, transfected with pCAGGS-flag-RIG-I plus viral protein constructs and stimulated as described above. Cell supernatants were harvested in 8 hours. Mx1-luc reporter cells were seeded into 96-well plates in triplicates and were treated 24 hours later with supernatants. At 16 hours post-incubation, cells were lysed in the passive lysis buffer (Promega), and luminescence was measured with a microplate reader (Tecan). The assay sensitivity was determined by a standard curve.

### Viral inhibitor assay

A549-ACE2 cells were seeded into 96-well plates in DMEM medium (10% FCS, 100 ug/ml Streptomycin, 100 IU/ml Penicillin) one day before infection. Six hours before infection, or at the time of infection, the medium was replaced with 100μl of DMEM medium containing either the compounds of interest or DMSO as a control. Infection was performed by adding 10μl of SARS-CoV-2-GFP (MOI 3) per well and plates were placed in the IncuCyte S3 Live-Cell Analysis System where whole well real-time images of mock (Phase channel) and infected (GFP and Phase channel) cells were captured every 4 hours for 48 hours. Cell viability (mock) and virus growth (mock and infected) were assessed as the cell confluence per well (Phase area) and GFP area normalized on cell confluence per well (GFP area/Phase area) respectively using IncuCyte S3 Software (Essen Bioscience; version 2019B Rev2).

For comparative analysis of antiviral treatment activity against SARS-CoV and SARS-CoV-2, A549-ACE2 cells were seeded in 24-well plates, as previously described. Treatment was performed for 6 hours with 0.5ml of DMEM medium containing either the compounds of interest or DMSO as a control, and infected with SARS-CoV-Frankfurt-1 or SARS-CoV-2-MUC-IMB-1 (MOI 1) for 24 hours. Total cellular RNA was harvested and analyzed by qRT-PCR, as previously described.

## Main text statements

## Acknowledgements

We thank Stefan Pöhlmann for sharing ACE2 plasmids and Robert Baier for technical assistance, Monika Trojan for advice on drugs, Juan Pancorbo and Johannes Albert-von der Gönna from the Leibniz Supercomputing Centre (www.lrz.de) for technical assistance. We further thank the Karl Max von Bauernfeind - Verein for support for the screening microscope. Work in the authors’ laboratories was supported by an ERC consolidator grant (ERC-CoG ProDAP, 817798), the German Research Foundation (PI 1084/3, PI 1084/4, PI 1084/5, TRR179/TP11, TRR237/A07) and the German Federal Ministry of Education and Research (COVINET) to A.P. This work was also supported by the German Federal Ministry of Education and Research (CLINSPECT-M) to BK. AW was supported by the China Scholarship Council (CSC). The work of JZ was supported by the German Research Foundation (SFB1021, A01 and B01; KFO309, P3), the State of Hessen through the LOEWE Program (DRUID, B02) and the German Ministry for Education and Research (COVINET, RAPID). We specially thank Roberto Polakiewicz, Florian Gnad and Cell Signaling Technology (CST) for gifting of PTMScan® Ubiquitin Remnant Motif (K-□-GG) Kits. The work of P.S. at the Heinrich Pette Institute is supported by the Free and Hanseatic City of Hamburg and the Federal Ministry of Health.

## Author contribution statements

Conceptualization: A.S., V.Gi., V.Gr., O.K., V.B., C.U., D.A.H., Y.H., J.Z., P.S., M.M.,A.Pic.

Investigation: V.Gi., V.Gr., O.K., V.B., C.U., D.A.H., Y.H., L.O., A.W., A.Pir., F.M.H., M.C.R., I.P., T.M.L., R.E., J.J., P.S.

Data analysis: A.S., V.Gi., V.Gr., V.B., O.K., C.U., D.A.H., Y.H., S.M.H., F.M.H., M.C.R., L.Z., T.E., M.R.

Funding acquisition: R.W., B.K., U.P., R.R., J.Z., V.T., M.M., A.Pic. Supervision: M.M., R.R., A.Pic.

Writing: A.S., V.Gi., V.Gr., O.K., V.B., C.U., D.A.H., Y.H., L.Z., M.M., A.Pic.

## Additional Information statement

### Supplementary Information

is available for this paper. Correspondence and requests for materials should be addressed to Andreas Pichlmair. Reprints and permissions information is available at www.nature.com/reprints.

## Data availability statement

The raw sequencing data for this study have been deposited in the European Nucleotide Archive (ENA) at EMBL-EBI under accession number PRJEB38744 (https://www.ebi.ac.uk/ena/data/view/PRJEB38744). The mass spectrometry proteomics data have been deposited to the ProteomeXchange Consortium *via* the PRIDE^89^ partner repository with the dataset identifier PXD022282, PXD020461 and PXD020222.

The protein interactions from this publication have been submitted to the IMEx (http://www.imexconsor-tium.org) consortium through IntAct^90^ with the identifier IM-28109.

The data and analysis results are accessible online *via* the interactive web interface at https://covinet.innatelab.org.

The following public data sets were used in the study:

‐ Gene Ontology and Reactome annotations (http://download.baderlab.org/EM_Genesets/April_01_2019/Human/UniProt/Human_GO_AllPathways_with_GO_iea_April_01_2019_UniProt.gmt)
‐ IntAct Protein Interactions (https://www.ebi.ac.uk/intact/, v2019.12)
‐ IntAct Protein Complexes (https://www.ebi.ac.uk/complexportal/home, v2019.12)
‐ CORUM Protein Complexes (http://mips.helmholtz-muenchen.de/corum/download/allComplexes.xml.zip, v2018.3)
‐ Reactome Functional Interactions (https://reactome.org/download/tools/ReatomeFIs/FIsInGene_020720_with_annotations.txt.zip)
‐ Human (v2019.10), Human-CoV, SARS-CoV-2 and SARS-CoV (v2020.08) protein sequences: https://uniprot.org.

## Code availability statement

In-house R and Julia packages and scripts used for the bioinformatics analysis of the data have been deposited to public GitHub repositories: https://doi.org/10.5281/zenodo.4536605, https://doi.org/10.5281/zenodo.4536603, https://doi.org/10.5281/zenodo.4536590, https://doi.org/10.5281/zenodo.4536596, https://doi.org/10.5281/zenodo.4541090, https://doi.org/10.5281/zenodo.4541082.

## Extended data legends

**Extended data Figure 1.**
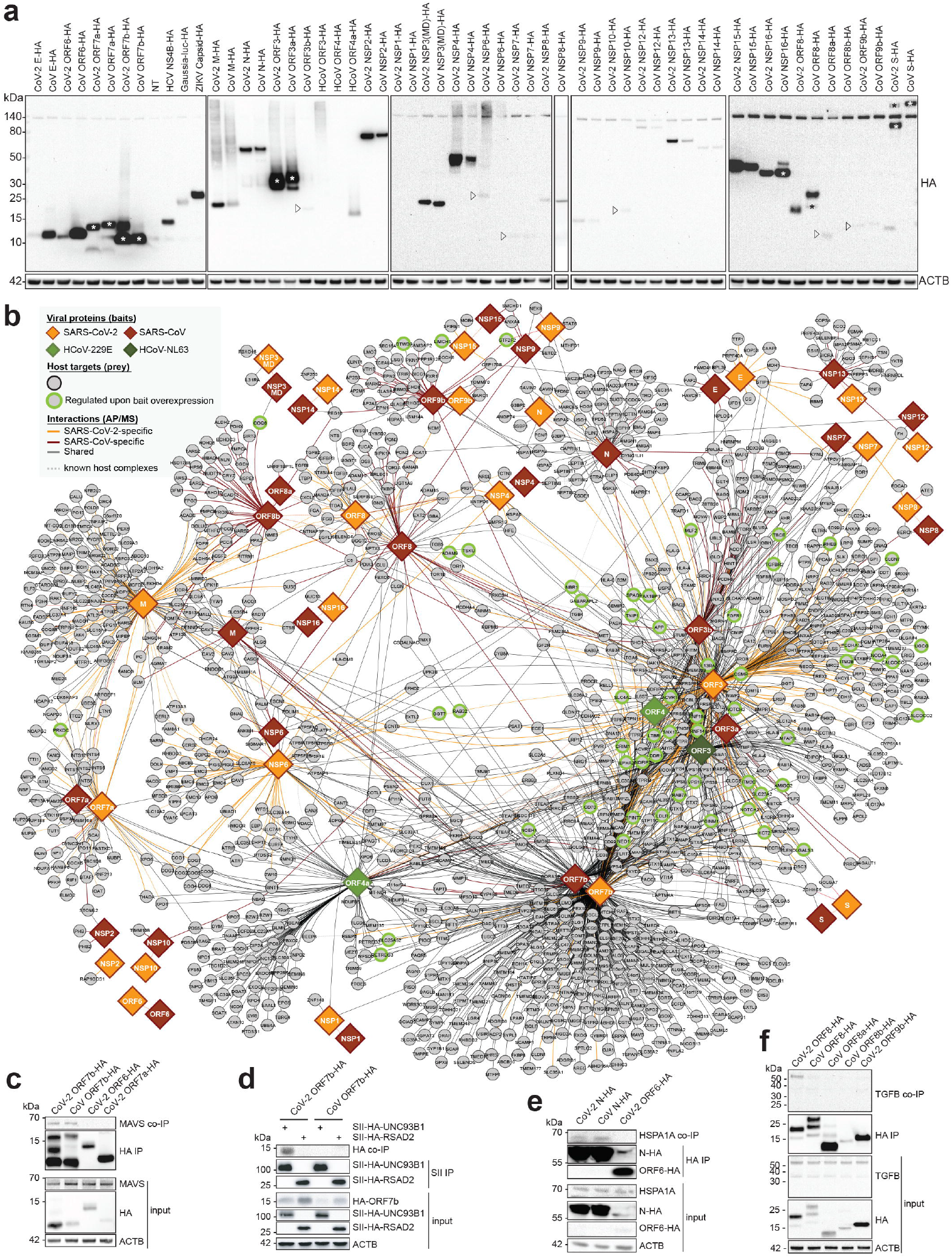
SARS-CoV-2 and SARS-CoV proteins expressed in A549 cells target host proteins. **(a)** Expression of HA-tagged viral proteins, in stably transduced A549 cells, used in AP-MS and proteome expression measurements. When several bands are present in a single lane, * marks the band with expected molecular weight (n = 4 independent experiments). For gel source data, see Supplementary Figure 1. **(b)** Extended version of the virus-host protein-protein interaction network with 24 SARS-CoV-2 and 27 SARS-CoV proteins, as well as ORF3 of HCoV-NL63 and ORF4 and ORF4a of HCoV-229E, used as baits. Host targets regulated upon viral protein overexpression are highlighted (see the in-plot legend). **(c-f)** Co-precipitation experiments in HEK293T cells showing a specific enrichment of **(c)** endogenous MAVS co-precipitated with C-terminal HA-tagged ORF7b of SARS-CoV-2 and SARS-CoV (negative controls: SARS-CoV-2 ORF6-HA, ORF7a-HA), **(d)** ORF7b-HA of SARS-CoV-2 and SARS-CoV co-precipitated with SII-HA-UNC93B1 (control precipitation: SII-HA-RSAD2), **(e)** endogenous HSPA1A co-precipitated with N-HA of SARS-CoV-2 and SARS-CoV (control: SARS-CoV-2 ORF6-HA) and **(f)** endogenous TGF-β with ORF8-HA of SARS-CoV-2 vs ORF8-HA, ORF8a-HA, ORF8b-HA of SARS-CoV or ORF9b-HA of SARS-CoV-2. (n=2 independent experiments). For gel source data, see Supplementary Figure 1. AP-MS: affinity-purification coupled to mass spectrometry; MD: Macro domain; NSP: Non-structural protein.

**Extended data Figure 2.**
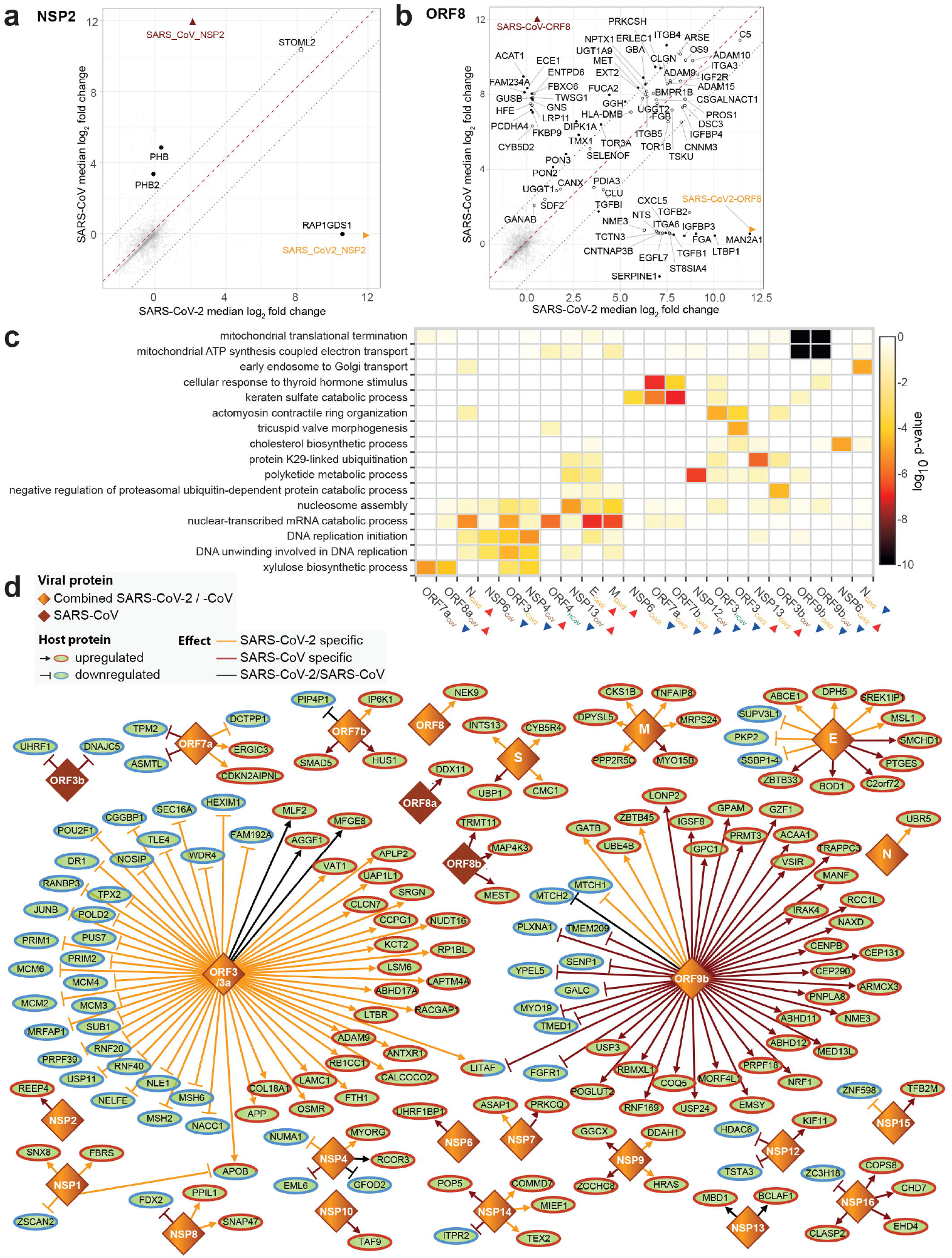
SARS-CoV-2 and SARS-CoV proteins trigger shared and specific interactions with host factors, and induce changes to the host proteome. **(a-b)** Differential enrichment of proteins in **(a)** NSP2 and **(b)** ORF8 of SARS-CoV-2 (*x*-axis) vs SARS-CoV (*y*-axis) AP-MS experiments (n=4 independent experiments). **(c)** Gene Ontology Biological Processes enriched among the cellular proteins that are up-(red arrow) or down-(blue arrow) regulated upon overexpression of individual viral proteins. **(d)** The most affected proteins from the effectome data of protein changes upon viral bait overexpression in A549 cells (see materials and methods for the exact protein selection criteria). Homologous viral proteins are displayed as a single node. Shared and virus-specific effects are denoted by the edge color. NSP: Non-structural protein.

**Extended data Figure 3.**
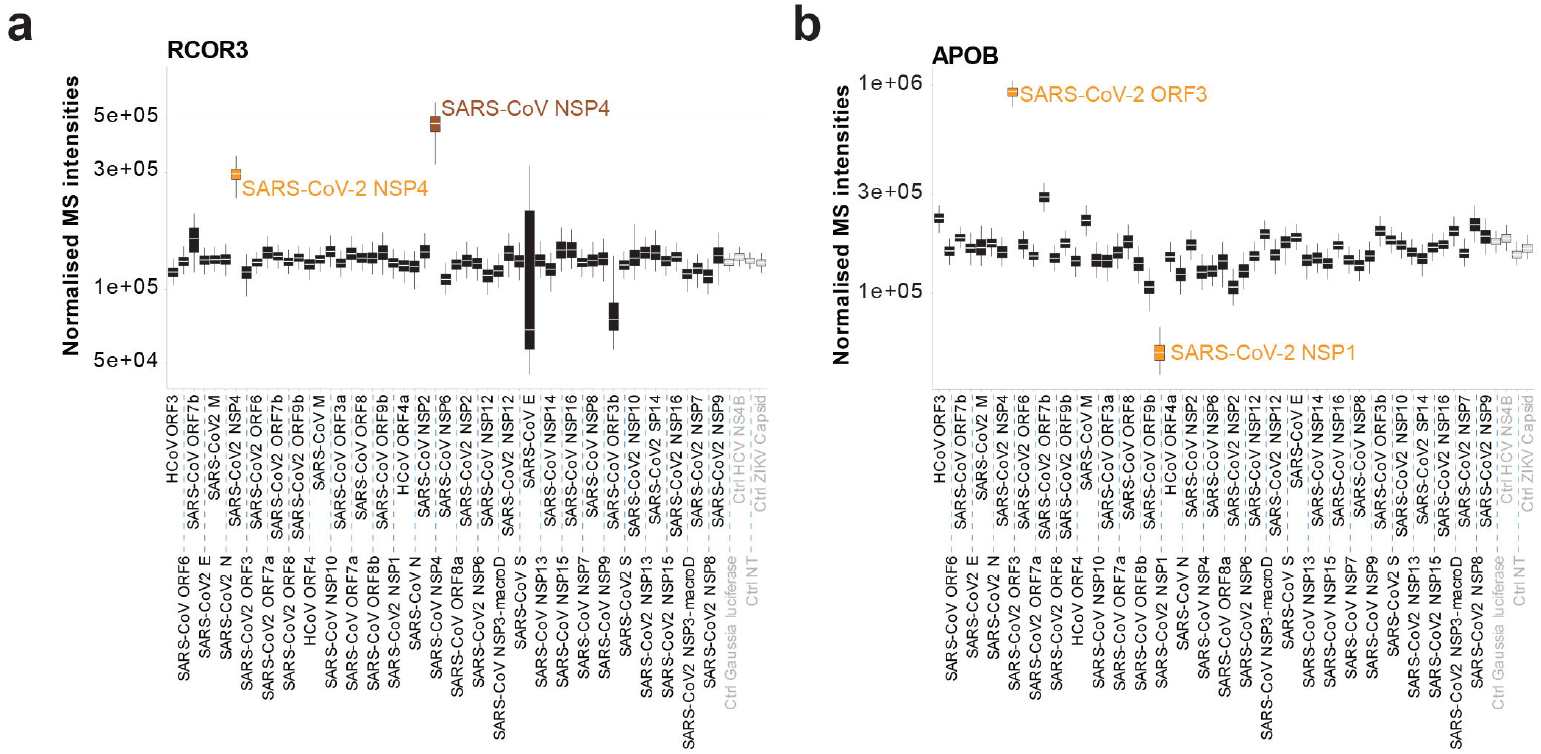
RCOR3 and APOB regulation upon SARS-CoV-2 and SARS-CoV protein over-expression. **(a-b)** Normalized intensities of selected candidates specifically perturbed by individual viral proteins: **(a)** RCOR3 was upregulated both by SARS-CoV-2 and SARS-CoV NSP4 proteins, **(b)** APOB was upregulated by ORF3 and downregulated by NSP1 specifically to SARS-CoV-2. The box and the whiskers represent 50% and 95% confidence intervals, and the white line corresponds to the median of the log_2_ fold-change upon viral protein overexpression (n=4 independent experiments).

**Extended data Figure 4.**
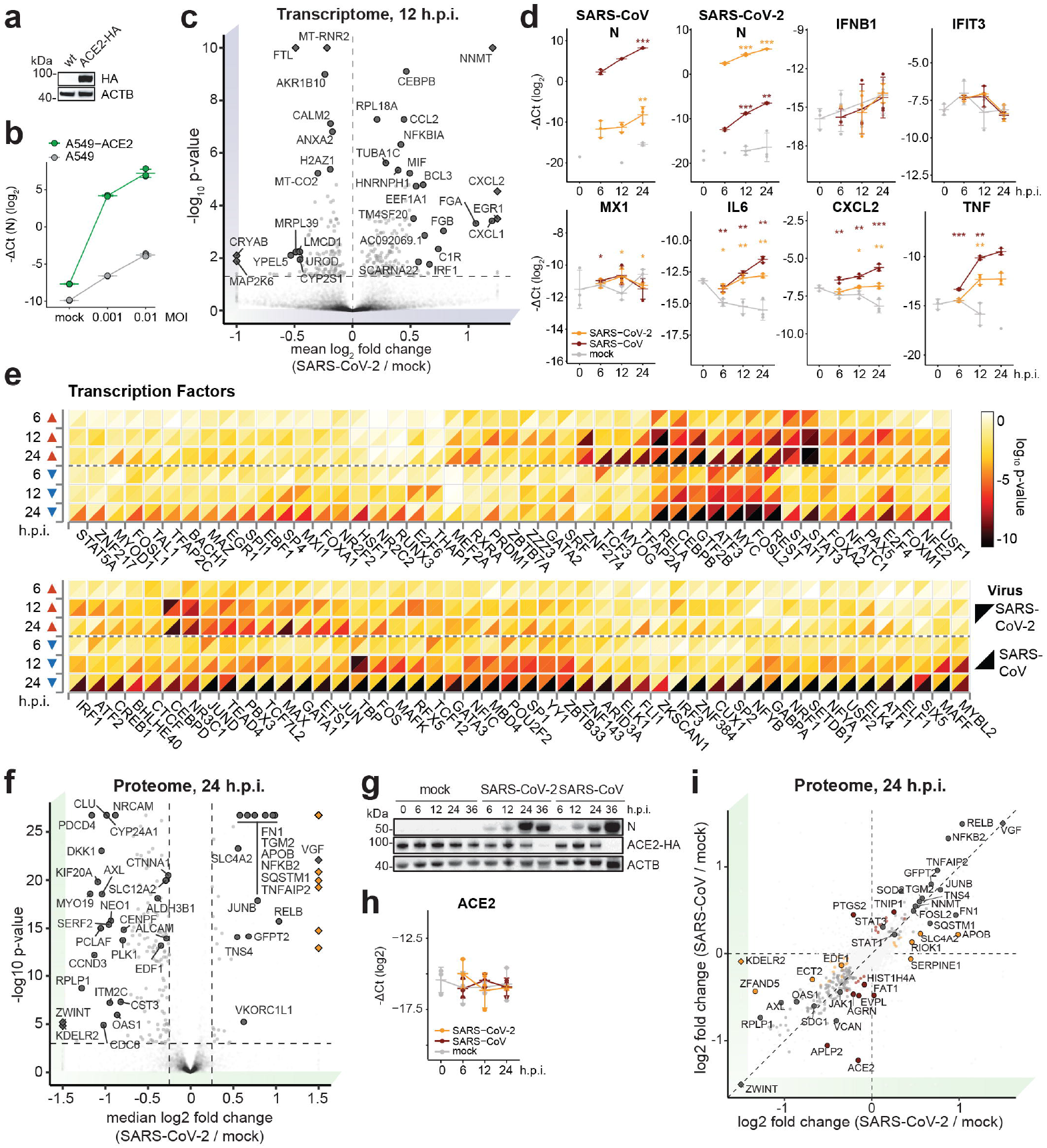
Tracking of virus-specific changes in infected A549-ACE2 cells by transcriptomics and proteomics. **(a)** Western blot showing ACE2-HA expression levels in A549 cells untransduced (wild-type) or transduced with ACE2-HA-encoding lentivirus (n = 2 independent experiments). For gel source data, see Supplementary Figure 1.**(b)** mRNA expression levels of SARS-CoV-2 *N* relative to *RPLP0* as measured by qRT-PCR upon infection of wild-type A549 and A549-ACE2 cells at the indicated MOIs. Error bars represent mean and standard deviation (n=3 independent experiments). **(c)** Volcano plot of mRNA expression changes of A549-ACE2 cells, infected with SARS-CoV-2 at an MOI of 2 in comparison to mock infection at 12 h.p.i. Significant hits are highlighted in gray (moderated t-test FDR-corrected two-sided p-value, n=3 independent experiments). Diamonds indicate that the actual log_2_ fold change or p-value were truncated to fit into the plot. **(d)** Expression levels, as measured by qRT-PCR, of SARS-CoV-2/SARS-CoV *N* and host transcripts relative to *RPLP0* in infected (MOI 2) A549-ACE2 cells with SARS-CoV-2 (orange) and SARS-CoV (brown) at indicated time points. Error bars correspond to mean and standard deviation (Two-sided student t-test, unadjusted p-value, n=3 independent experiments). *: p-value ≥ 0.05; **: p-value ≥ 0.01; ***: p-value ≥ 10^−3^. **(e)** Analysis of transcription factors, whose targets are significantly enriched among up-(red arrow) and down-(blue arrow) regulated genes of A549-ACE2 cells infected with SARS-CoV-2 (upper triangle) and SARS-CoV (lower triangle) for indicated time points (Fisher’s exact test unadjusted one-sided p-value ≤ 10^−4^). **(f)** Volcano plot of SARS-CoV-2-induced protein abundance changes at 24 h.p.i. in comparison to mock. Viral proteins are highlighted in orange, selected significant hits are marked in black (Bayesian linear model-based unadjusted two-sided p-value ≤ 10^−3^, |median log_2_ fold change|≥ 0.25, n=4 independent experiments). Diamonds indicate that the actual log_2_ fold change was truncated to fit into the plot. **(g)** Western blot showing the total levels of ACE2-HA protein at 6, 12, 24 and 36 h.p.i. (mock, SARS-CoV-2 and SARS-CoV infections); N viral protein as infection and ACTB as loading controls (n = 3 independent experiments). For gel source data, see Supplementary Figure 1. **(h)** Stable expression of *ACE2* mRNA transcript relative to *RPLP0*, as measured by qRT-PCR, after SARS-CoV-2 and SARS-CoV infections (MOI 2) of A549-ACE2 cells at indicated h.p.i. (error bars show mean and standard deviation, n=3 independent experiments). **(i)** Scatter plots comparing the host proteome of SARS-CoV-2 (*x*-axis) and SARS-CoV (*y*-axis) infection at 24 h.p.i. (log_2_ fold change in comparison to the mock infection samples at the same time point). Significantly regulated proteins (Bayesian linear model-based unadjusted two-sided p-value ≤ 10^−3^, |log_2_ fold change|≥ 0.25, n=4 independent experiments), are colored according to their specificity in both infections. Diamonds indicate that the actual log_2_ fold change was truncated to fit into the plot. h.p.i.: hours post-infection; MOI: multiplicity of infection.

**Extended data Figure 5.**
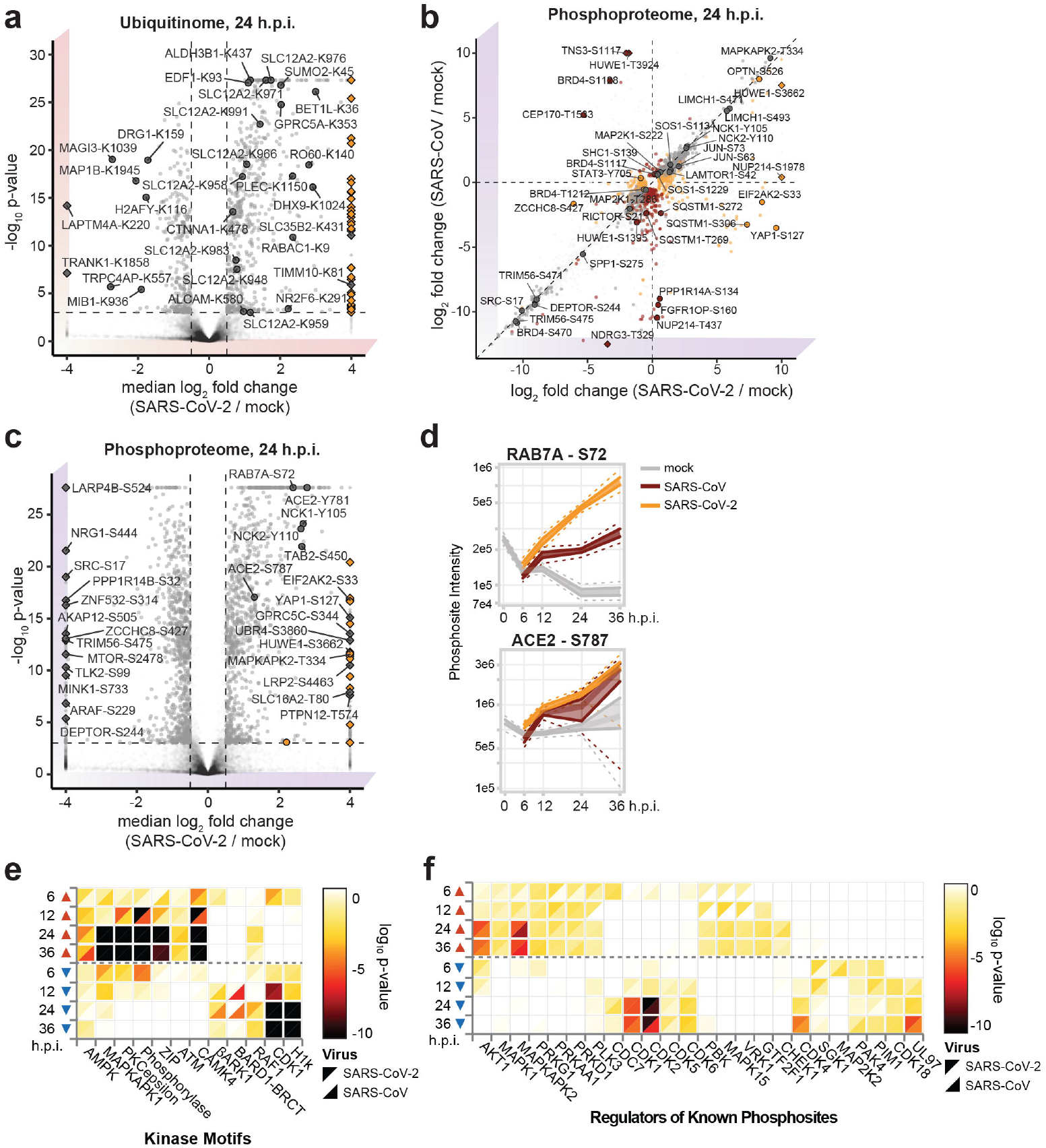
Post-translational modifications modulated during SARS-CoV-2 or SARS-CoV infection. **(a)** Volcano plots of SARS-CoV-2-induced ubiquitination changes at 24 h.p.i. in comparison to mock. The viral PTM sites are highlighted in orange and selected significant hits in black. **(b)** Scatter plots comparing the host phosphoproteome of SARS-CoV-2 (*x*-axis) and SARS-CoV (*y*-axis) infection at 24 h.p.i. (log_2_ fold change in comparison to the mock infection samples at the same time point). Significantly regulated sites are colored according to their specificity in both infections. **(c)** Volcano plots of SARS-CoV-2-induced phosphorylation changes at 24 h.p.i. in comparison to mock. The viral PTM sites are highlighted in orange and selected significant hits in black. For **(a-c)**, a change is defined significant if its Bayesian linear model-based unadjusted two-sided p-value ≤ 10^−3^ and |log_2_ fold change|≥ 0.5, n=3 independent experiments for ubiquitination and n=4 independent experiments for phosphorylation data. Diamonds in (a-c) indicate that the actual log_2_ fold change was truncated to fit into the plot. **(d)** Profile plots showing the time-resolved phosphorylation of ACE2 (S787) and RAB7A (S72) with indicated median, 50% and 95% confidence intervals. n = 4 independent experiments **(e)** The enrichment of host kinase motifs among the significantly regulated phosphorylation sites of SARS-CoV-2 (upper triangle) and SARS-CoV-infected (lower triangle) A549-ACE2 cells (MOI 2) at the indicated time points (Fisher’s exact test, unadjusted one-sided p-value ≤ 10^−3^). **(f)** The enrichment of specific kinases among the ones known to phosphorylate significantly regulated sites at the indicated time points and annotated in PhosphoSitePlus database (Fisher’s exact test, unadjusted one-sided p-value ≤ 10^−2^). h.p.i.: hours post-infection.

**Extended Data Figure 6.**
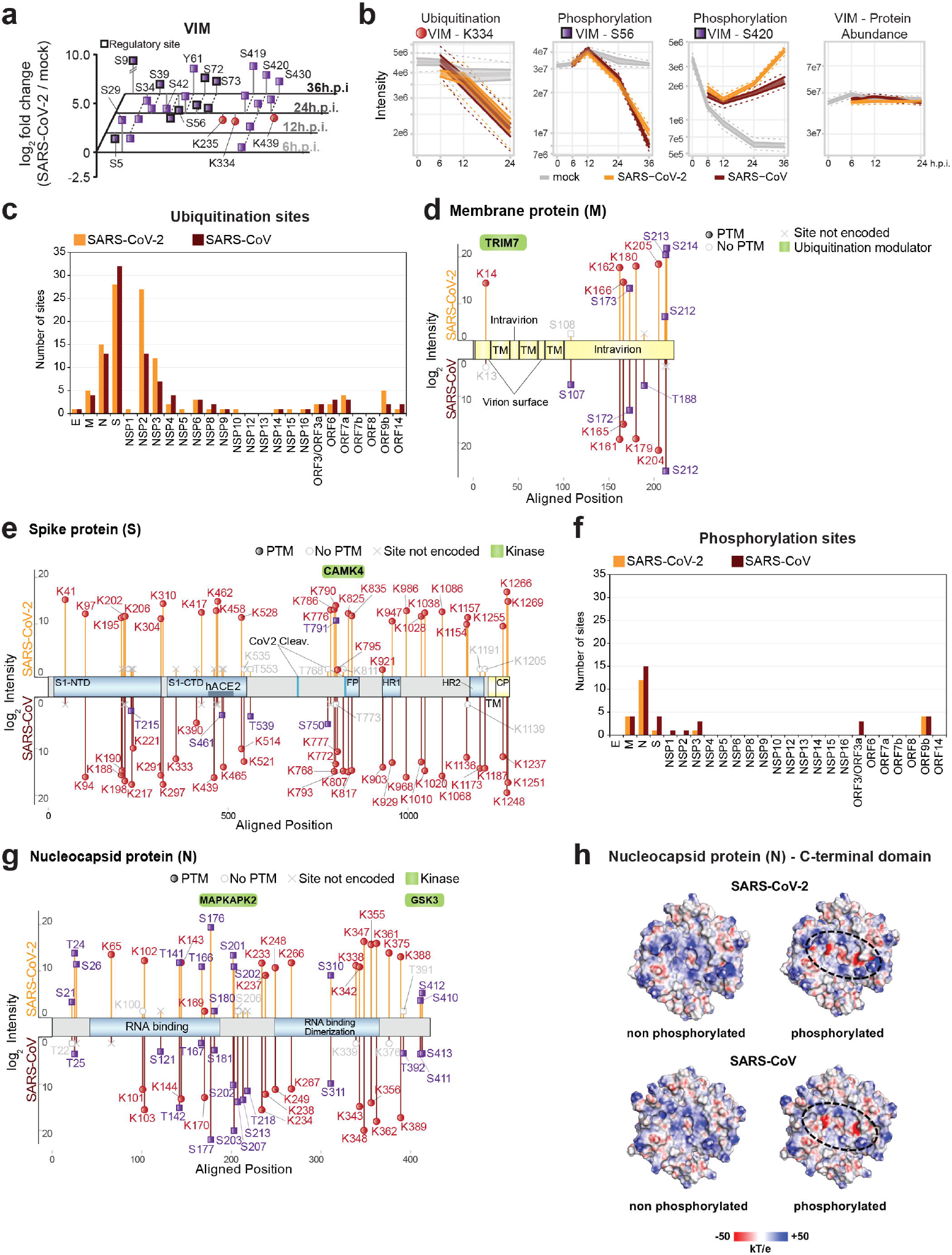
Integration of multi-omics data from SARS-CoV-2 and SARS-CoV infection identified co-regulation of host and viral factors. **(a)** Phosphorylation (purple square) and ubiquitination (red circles) sites on vimentin (VIM) regulated upon SARS-CoV-2 infection. The plot shows the medians of log_2_ fold changes compared to mock at 6, 12, 24, and 36 h.p.i., regulatory sites are indicated with a thick black border. **(b)** Profile plots of VIM K334 ubiquitination, S56 and S72 phosphorylation, and total protein levels in SARS-CoV-2 or SARS-CoV infected A549-ACE2 cells at indicated times after infection, with indicated median, 50% and 95% confidence intervals. n=3 (ubiquitination) or 4 (total protein levels, phosphorylation) independent experiments **(c)** Number of ubiquitination sites identified on each SARS-CoV-2 or SARS-CoV proteins in infected A549-ACE2 cells. **(d-e)** Mapping the ubiquitination and phosphorylation sites of SARS-CoV-2/SARS-CoV M and S proteins on their aligned sequence with median log_2_ intensities in infected A549-ACE2 cells at 24 h.p.i. (n=4 independent experiments for phosphorylation and n = 3 independent experiments for ubiquitination data). Functional (blue) and topological (yellow) domains are mapped on each sequence. Binding of ubiquitin modifying enzymes to both M proteins and the host kinases that potentially recognise motifs associated with the reported sites and overrepresented among cellular motifs enriched upon infection (Extended data Fig. 5e, f) or interacting with given viral protein (Extended data Fig. 1b) are indicated (green). **(f)** Number of phosphorylation sites identified on each SARS-CoV-2 or SARS-CoV proteins in infected A549-ACE2 cells. **(g)** Mapping the ubiquitination (red circle) and phosphorylation (purple square) sites of SARS-CoV-2/SARS-CoV N protein on their aligned sequence with median log_2_ intensities in A549-ACE2 cells infected with the respective virus at 24 h.p.i. (n=4 independent experiments). Functional (blue) domains are mapped on each sequence. The host kinases that potentially recognise motifs associated with the reported sites and overrepresented among cellular motifs enriched upon infection (Extended data Fig. 5e, f) or interacting with given viral protein (Extended data Fig. 1b) (green). **(h)** Electrostatic surface potential analysis of non-phosphorylated and phosphorylated SARS-CoV and SARS-CoV-2 N CTD dimers is shown on the right panels; red, white and blue regions represent areas with negative, neutral and positive electrostatic potential, respectively (scale from −50 to +50 kT e^−1^). h.p.i.: hours post-infection; TM: transmembrane domain; CTD: C-terminal domain.

**Extended Data Figure 7.**
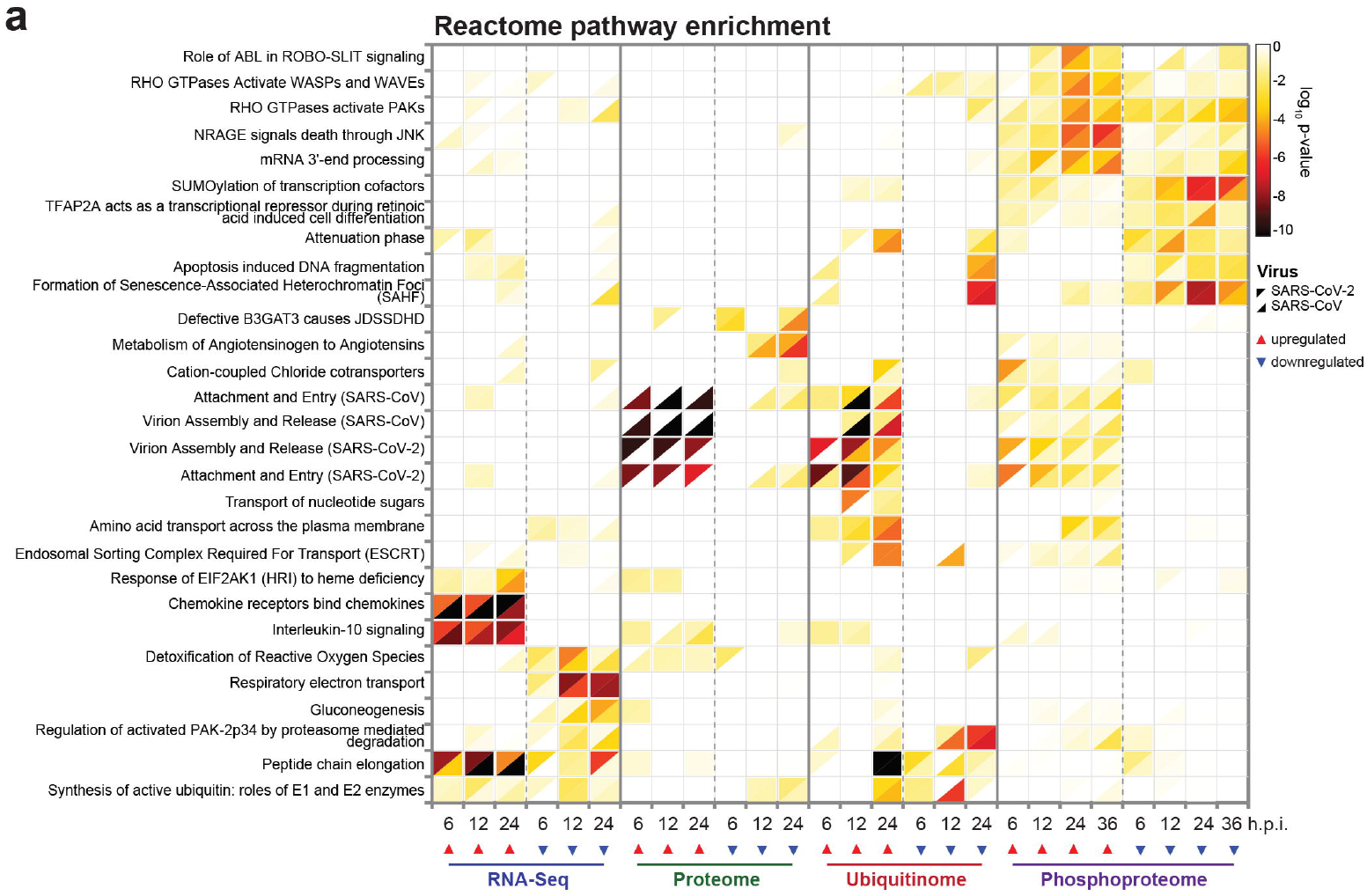
Reactome pathways enrichment in multi-omics data of SARS-CoV-2 and SARS-CoV infection. **(a)** Reactome pathways enriched in up-(red arrow) or downregulated (blue arrow) transcripts, proteins, ubiquitination and phosphorylation sites (Fisher’s exact test unadjusted p-value ≤ 10^−4^) in SARS-CoV-2 or SARS-CoV-infected A549-ACE2 cells at indicated times after infection. h.p.i.: hours post-infection.

**Extended Data Figure 8.**
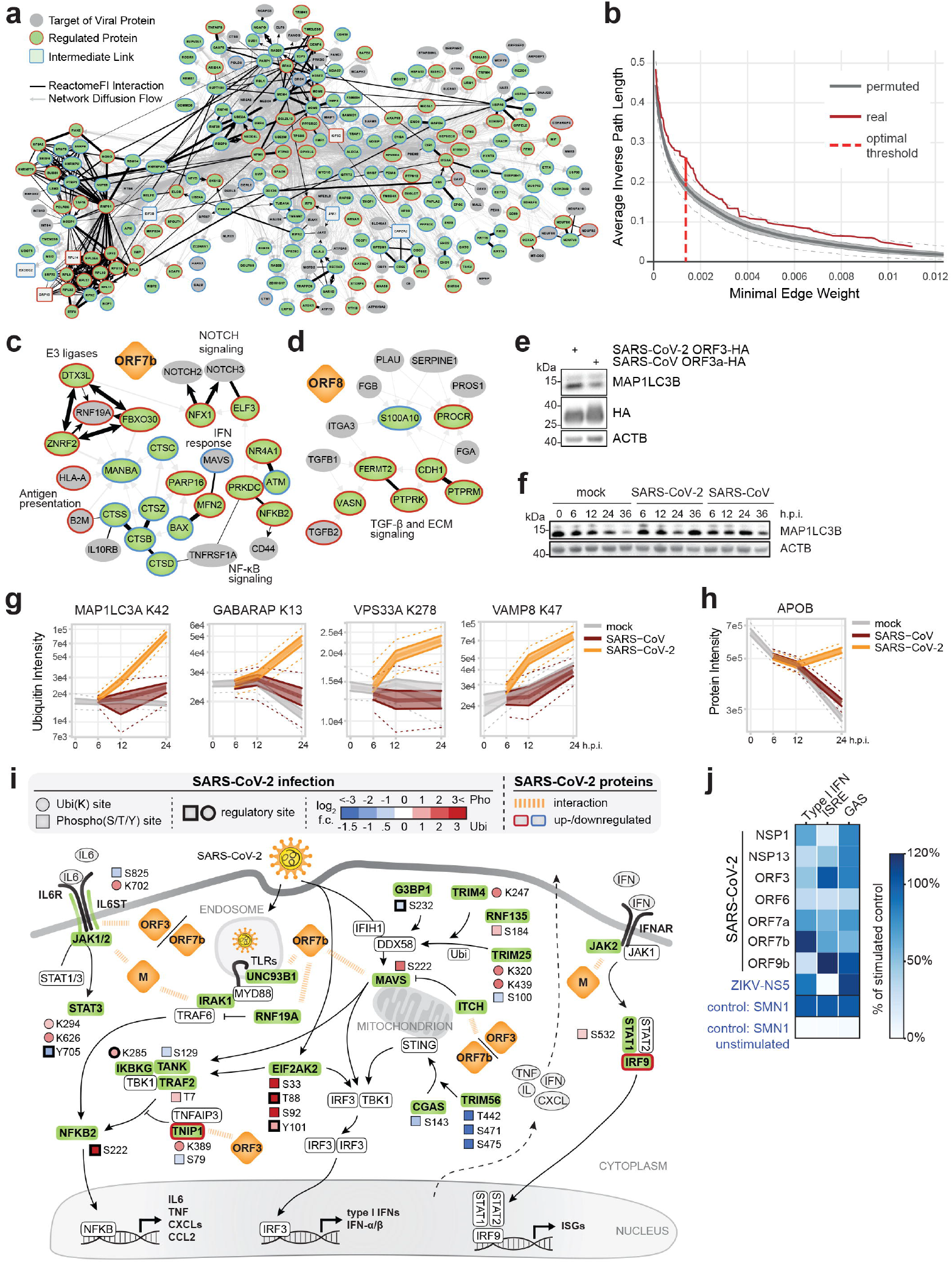
SARS-CoV-2 uses a multi-pronged approach to perturb host-pathways at several levels. **(a)** The host subnetwork perturbed by SARS-CoV-2 M predicted by the network diffusion approach. Edge thickness reflects the transition probability in random walk with restart, directed edges represent the walk direction, and ReactomeFI connections are highlighted in black. **(b)** Selection of the optimal threshold for the network diffusion model of SARS-CoV-2 M-induced proteome changes. The plot shows the relationship between the minimal allowed edge weight of the random walk graph (*x*-axis) and the mean inverse length of the path from the regulated proteins to the host targets of the viral protein along the edges of the resulting filtered subnetwork (*y*-axis). The red curve represents the metric for the network diffusion analysis of the actual data. The grey band shows 50% confidence interval, and dashed lines correspond to 95% confidence interval for the average inverse path length distribution for 1000 randomised datasets. Optimal edge weight threshold that maximizes the difference between the metric based on real data and its 3rd quartile based on randomized data is highlighted by the red vertical line. **(c-d)** Subnetworks of the network diffusion predictions linking host targets of SARS-CoV-2 **(c)** ORF7b to the factors involved in innate immunity and **(d)** ORF8 to the factors involved in TGF-β signaling. **(e-f)** Western blot showing the accumulation of the autophagy-associated factor MAP1LC3B upon **(e)** SARS-CoV-2 ORF3 expression in HEK293-R1 cells (n=3 independent experiments) and **(f)** SARS-CoV-2/SARS-CoV infection of A549-ACE2 cells (n=3 independent experiments). For gel source data, see Supplementary Figure 1. **(g-h)** Profile plots showing the time-resolved **(g)** ubiquitination of the autophagy regulators MAP1LC3A, GABARAP, VPS33A and VAMP8 (n=3 independent experiments), as well as **(h)** an increase in total protein abundance of APOB with indicated median, 50% and 95% confidence intervals (n=4 independent experiments). **(i)** Overview of perturbations to host-cell innate immunity-related pathways, induced by distinct proteins of SARS-CoV-2, derived from the network diffusion model and overlaid with transcriptional, ubiquitination and phosphorylation changes upon SARS-CoV-2 infection. **(j)** Heatmap showing the effects of the indicated SARS-CoV-2 proteins on type-I IFN expression levels, ISRE and GAS promoter activation in HEK293-R1. Accumulation of type-I IFN in the supernatant was evaluated by testing supernatants of PPP-RNA (IVT4) stimulated cells on MX1-luciferase reporter cells, ISRE promoter activation – by luciferase assay after IFN-α stimulation, and GAS promoter activation – by luciferase assay after IFN-γ stimulation in cells expressing SARS-CoV-2 proteins as compared to the controls (ZIKV NS5 and SMN1) (n=3 independent experiments).

**Extended Data Figure 9.**
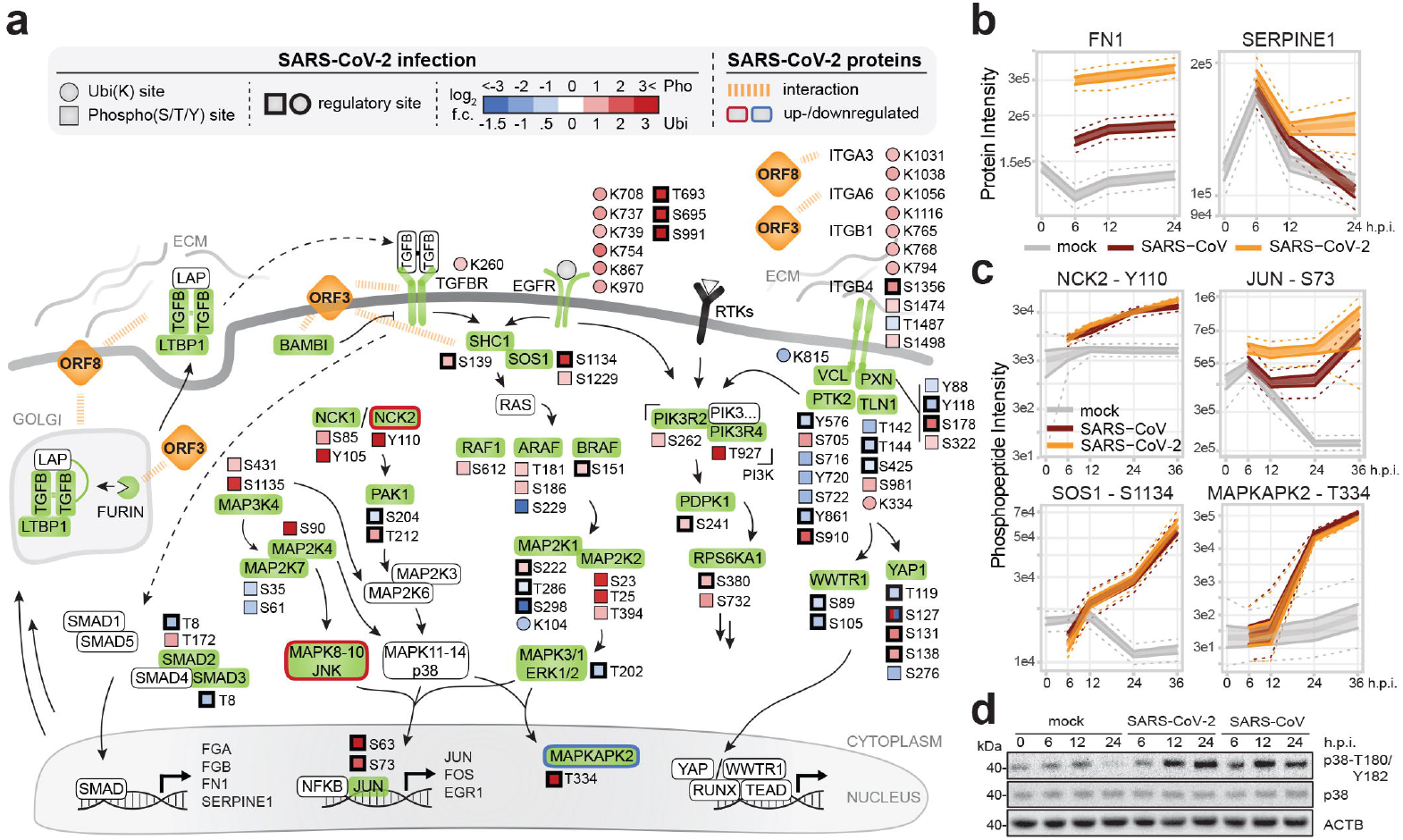
Perturbation of host integrin-TGF-β-EGFR-RTK signaling by SARS-CoV-2. **(a)** Overview of perturbations to host-cell Integrin-TGF-β-EGFR-RTK signaling, induced by distinct proteins of SARS-CoV-2, derived from the network diffusion model and overlaid with transcriptional, ubiquitination and phosphorylation changes upon SARS-CoV-2 infection. **(b)** Profile plots of total protein levels of SERPINE1 and FN1 in SARS-CoV-2 or SARS-CoV-infected A549-ACE2 cells at 6, 12, and 24 h.p.i., with indicated median, 50% and 95% confidence intervals. (n = 4 independent experiments) **(c)** Profile plots showing intensities of indicated phosphosites on NCK2, JUN, SOS1 and MAPKAPK2 in SARS-CoV-2 or SARS-CoV-infected A549-ACE2 cells at 6, 12, 24 and 36 h.p.i., with indicated median, 50% and 95% confidence intervals. (n = 4 independent experiments) **(d)** Western blot showing phosphorylated (T180/Y182) and total protein levels of p38 in SARS-CoV-2 or SARS-CoV infected A549-ACE2 cells. (n = 3 independent experiments) For gel source data, see Supplementary Figure 1. h.p.i.: hours post-infection.

**Extended data Figure 10.**
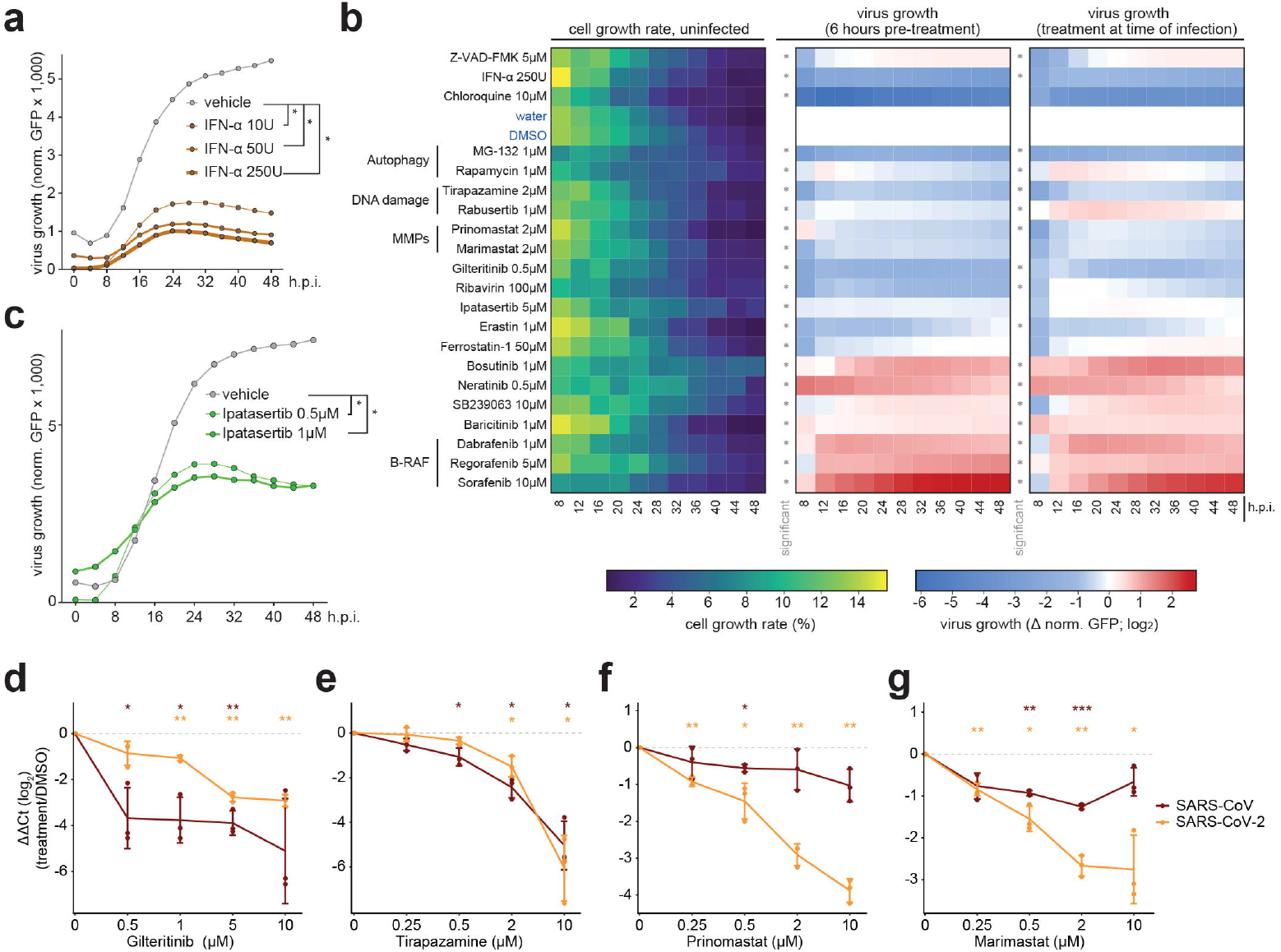
Drug repurposing screen, focusing on pathways perturbed by SARS-CoV-2, reveals potential candidates for use in antiviral therapy. **(a)** A549-ACE2 cells exposed for 6 hours to the specified concentrations of IFN-α and infected with SARS-CoV-2-GFP reporter virus (MOI 3). GFP signal and cell confluency were analyzed by live-cell imaging for 48 h.p.i. Time-courses show virus growth over time as the mean of GFP-positive area normalized to the total cell area (n=4 independent experiments). **(b)** A549-ACE2 cells were pre-treated for 6 hours or treated at the time of infection with SARS-CoV-2-GFP reporter virus (MOI 3). GFP signal and cell growth were tracked for 48 h.p.i. by live-cell imaging using an Incucyte S3 platform. Left heatmap: the cell growth rate (defined as the ratio of cell confluence change between the confluence at t and t-1) over time in drug-treated uninfected conditions. Middle (6 hours of pre-treatment) and right (treatment at the time of infection) heatmaps: treatment-induced changes in virus growth over time (GFP signal normalized to total cell confluence log_2_ fold change between the treated and control (water, DMSO) conditions). Only non-cytotoxic treatments with significant effects on SARS-CoV-2-GFP are shown. Asterisks indicate significance of the difference to the control treatment (Wilcoxon test; unadjusted two-sided p-value ≤ 0.05, n=4 independent experiments). **(c)** A549-ACE2 cells exposed for 6 hours to the specified concentrations of Ipatasertib and infected with SARS-CoV-2-GFP reporter virus (MOI 3). GFP signal and cell confluency were analyzed by live-cell imaging for 48 h.p.i. Time-courses show virus growth over time as the mean of GFP-positive area normalized to the total cell area (n=4 independent experiments). **(d-g)** mRNA expression levels at 24 h.p.i. of SARS-CoV-2 (orange) and SARS-CoV (brown) *N* relative to *RPLP0*, compared to DMSO-treated cells, as measured by qRT-PCR in infected A549-ACE2 cells (MOI 1) pre-treated for 6 hours with **(d)** Gilteritinib, **(e)** Tirapazamine, **(f)** Prinomastat or **(g)** Marimastat. Error bars represent mean and standard deviation (Student t-test, two-sided, unadjusted p-value, n=3 independent experiments). *: p-value ≥ 0.05; **: p-value ≥ 0.01; ***: p-value ≥ 10^−3^. h.p.i.: hours post-infection, MOI: multiplicity of infection.

**Extended data Table 1.**
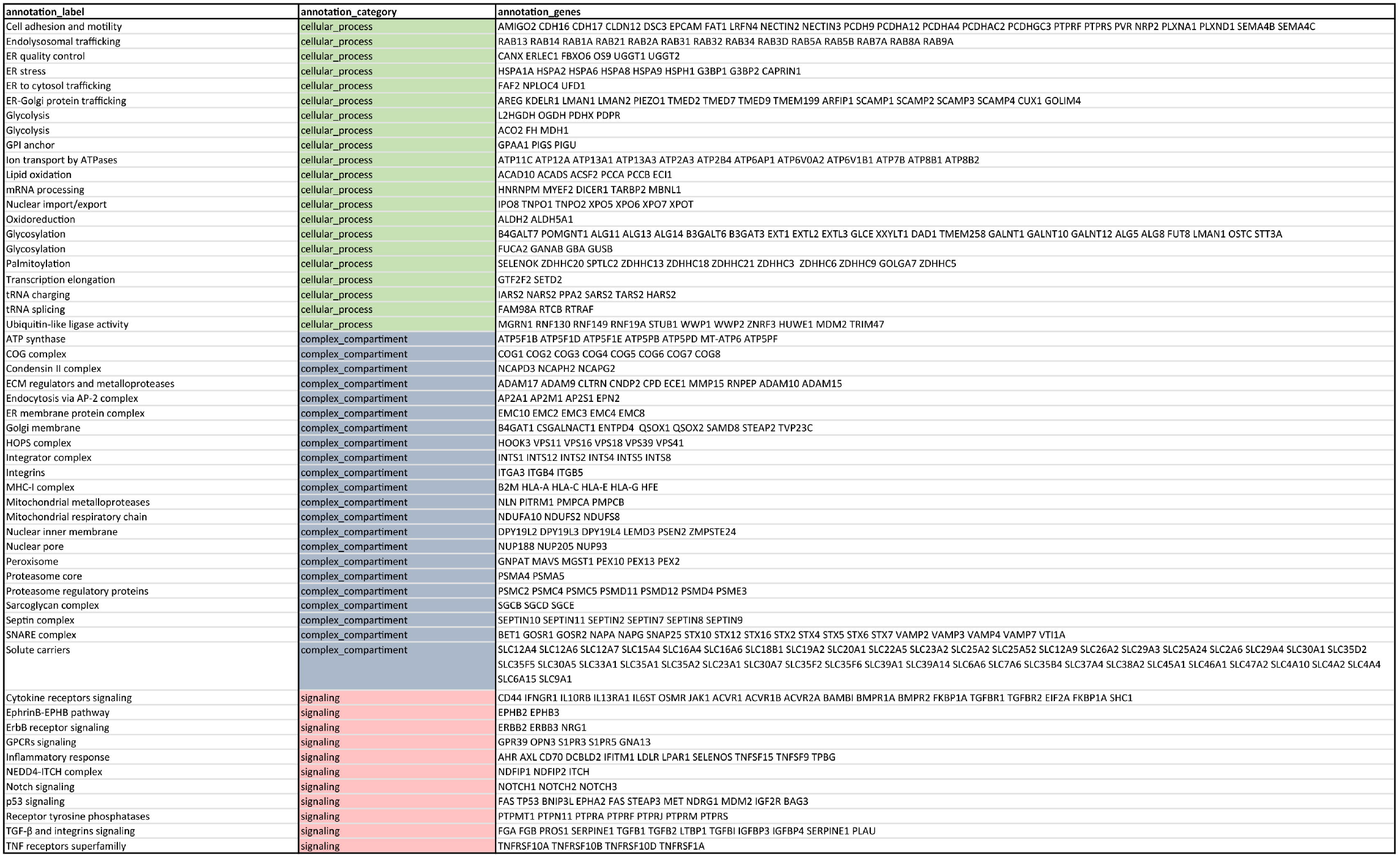
Functional annotations of the protein-protein interaction network of SARS-CoV-2 and SARS-CoV (AP-MS). Proteins identified as SARS-CoV-2 and/or SARS-CoV host binders *via* AP-MS (Figure 1b) grouped based on functional enrichment analysis of GOBP, GPCC, GPMF and Reactome terms (Supplementary table 2).

## References (main)

1. Gordon, D. E. et al. A SARS-CoV-2 protein interaction map reveals targets for drug repurposing. Nature 583, 459–468 (2020).

2. Gordon, D. E. et al. Comparative host-coronavirus protein interaction networks reveal pan-viral disease mechanisms. Science (2020) doi:10.1126/science.abe9403.

3. Bojkova, D. et al. Proteomics of SARS-CoV-2-infected host cells reveals therapy targets. Nature 583, 469–472 (2020).

4. Messner, C. B. et al. Ultra-high-throughput clinical proteomics reveals classifiers of COVID-19 infection. Cell Systems (2020) doi:10.1016/j.cels.2020.05.012.

5. Samavarchi-Tehrani, P. et al. A SARS-CoV-2 – host proximity interactome. bioRxiv 2020.09.03.282103 (2020) doi:10.1101/2020.09.03.282103.

6. Laurent, E. M. N. et al. Global BioID-based SARS-CoV-2 proteins proximal interactome unveils novel ties between viral polypeptides and host factors involved in multiple COVID19-associated mechanisms. bioRxiv 2020.08.28.272955 (2020) doi:10.1101/2020.08.28.272955.

7. Klann, K. et al. Growth Factor Receptor Signaling Inhibition Prevents SARS-CoV-2 Replication. Molecular Cell 80, 164–174.e4 (2020).

8. Huang, J. et al. SARS-CoV-2 Infection of Pluripotent Stem Cell-Derived Human Lung Alveolar Type 2 Cells Elicits a Rapid Epithelial-Intrinsic Inflammatory Response. Cell Stem Cell (2020) doi:10.1016/j.stem.2020.09.013.

9. Blanco-Melo, D. et al. Imbalanced Host Response to SARS-CoV-2 Drives Development of COVID-19. Cell 181, 1036–1045.e9 (2020).

10. Li, J. et al. Virus-Host Interactome and Proteomic Survey Reveal Potential Virulence Factors Influencing SARS-CoV-2 Pathogenesis. Med (2020) doi:10.1016/j.medj.2020.07.002.

## Additional references

11. von Brunn, A. et al. Analysis of Intraviral Protein-Protein Interactions of the SARS Coronavirus ORFeome. PLoS One 2, (2007).

12. Cornillez-Ty, C. T., Liao, L., Yates, J. R., Kuhn, P. & Buchmeier, M. J. Severe Acute Respiratory Syndrome Coronavirus Nonstructural Protein 2 Interacts with a Host Protein Complex Involved in Mitochondrial Biogenesis and Intracellular Signaling. Journal of Virology 83, 10314–10318 (2009).

13. Andrianifahanana, M. et al. ERBB receptor activation is required for profibrotic responses to transforming growth factor beta. Cancer Res. 70, 7421–7430 (2010).

14. Pittet, J.-F. et al. TGF-β is a critical mediator of acute lung injury. J Clin Invest 107, 1537–1544 (2001).

15. George, P. M., Wells, A. U. & Jenkins, R. G. Pulmonary fibrosis and COVID-19: the potential role for antifibrotic therapy. Lancet Respir Med 8, 807–815 (2020).

16. Mo, X. et al. Abnormal pulmonary function in COVID-19 patients at time of hospital discharge. European Respiratory Journal (2020) doi:10.1183/13993003.01217-2020.

17. Heo, J.-M. et al. Integrated proteogenetic analysis reveals the landscape of a mitochondrial-autophagosome synapse during PARK2-dependent mitophagy. Science Advances 5, eaay4624 (2019).

18. Shi, C.-S. et al. SARS-Coronavirus Open Reading Frame-9b Suppresses Innate Immunity by Targeting Mitochondria and the MAVS/TRAF3/TRAF6 Signalosome. The Journal of Immunology 193, 3080–3089 (2014).

19. Hoagland, D. A. et al. Modulating the transcriptional landscape of SARS-CoV-2 as an effective method for developing antiviral compounds. bioRxiv 2020.07.12.199687 (2020) doi:10.1101/2020.07.12.199687.

20. Castiglione, V., Chiriacò, M., Emdin, M., Taddei, S. & Vergaro, G. Statin therapy in COVID-19 infection. Eur Heart J Cardiovasc Pharmacother (2020) doi:10.1093/ehjcvp/pvaa042.

21. Radenkovic, D., Chawla, S., Pirro, M., Sahebkar, A. & Banach, M. Cholesterol in Relation to COVID-19: Should We Care about It? J Clin Med 9, (2020).

22. Chu, H. et al. Comparative replication and immune activation profiles of SARS-CoV-2 and SARS-CoV in human lungs: an ex vivo study with implications for the pathogenesis of COVID-19. Clin Infect Dis doi:10.1093/cid/ciaa410.

23. Zhu, Z. et al. From SARS and MERS to COVID-19: a brief summary and comparison of severe acute respiratory infections caused by three highly pathogenic human coronaviruses. Respiratory Research 21, 224 (2020).

24. Cazzaniga, A., Locatelli, L., Castiglioni, S. & Maier, J. The Contribution of EDF1 to PPARγ Transcriptional Activation in VEGF-Treated Human Endothelial Cells. Int J Mol Sci 19, (2018).

25. Gavriilaki, E. et al. Endothelial Dysfunction in COVID-19: Lessons Learned from Coronaviruses. Curr Hypertens Rep 22, 63 (2020).

26. Daniloski, Z. et al. Identification of Required Host Factors for SARS-CoV-2 Infection in Human Cells. Cell (2020) doi:10.1016/j.cell.2020.10.030.

27. Shinde, S. R. & Maddika, S. PTEN modulates EGFR late endocytic trafficking and degradation by dephosphorylating Rab7. Nat Commun 7, 10689 (2016).

28. Wang, D. et al. Auto-phosphorylation Represses Protein Kinase R Activity. Scientific Reports 7, 44340 (2017).

29. Yu, Y. T.-C. et al. Surface vimentin is critical for the cell entry of SARS-CoV. Journal of Biomedical Science 23, 14 (2016).

30. dos Santos, G. et al. Vimentin regulates activation of the NLRP3 inflammasome. Nature Communications 6, 6574 (2015).

31. Ramos, I., Stamatakis, K., Oeste, C. L. & Pérez-Sala, D. Vimentin as a Multifaceted Player and Potential Therapeutic Target in Viral Infections. Int J Mol Sci 21, (2020).

32. Zinzula, L. et al. High-resolution structure and biophysical characterization of the nucleocapsid phosphoprotein dimerization domain from the Covid-19 severe acute respiratory syndrome coronavirus 2. Biochem Biophys Res Commun (2020) doi:10.1016/j.bbrc.2020.09.131.

33. Chen, C.-Y. et al. Structure of the SARS Coronavirus Nucleocapsid Protein RNA-binding Dimerization Domain Suggests a Mechanism for Helical Packaging of Viral RNA. Journal of Molecular Biology 368, 1075–1086 (2007).

34. Perry, J. S. A. et al. Interpreting an apoptotic corpse as anti-inflammatory involves a chloride sensing pathway. Nature Cell Biology 21, 1532–1543 (2019).

35. Wu, G., Dawson, E., Duong, A., Haw, R. & Stein, L. ReactomeFIViz: a Cytoscape app for pathway and network-based data analysis. F1000Res 3, 146 (2014).

36. Reyna, M. A., Leiserson, M. D. M. & Raphael, B. J. Hierarchical HotNet: identifying hierarchies of altered subnetworks. Bioinformatics 34, i972–i980 (2018).

37. Cottam, E. M., Whelband, M. C. & Wileman, T. Coronavirus NSP6 restricts autophagosome expansion. Autophagy 10, 1426–1441 (2014).

38. Ohsaki, Y., Cheng, J., Fujita, A., Tokumoto, T. & Fujimoto, T. Cytoplasmic Lipid Droplets Are Sites of Convergence of Proteasomal and Autophagic Degradation of Apolipoprotein B. Mol Biol Cell 17, 2674–2683 (2006).

39. Khalil Maged F., Wagner William D., & Goldberg Ira J. Molecular Interactions Leading to Lipoprotein Retention and the Initiation of Atherosclerosis. Arteriosclerosis, Thrombosis, and Vascular Biology 24, 2211–2218 (2004).

40. Nicolai Leo et al. Immunothrombotic Dysregulation in COVID-19 Pneumonia Is Associated With Respiratory Failure and Coagulopathy. Circulation 142, 1176–1189 (2020).

41. Zavadil, J. et al. Genetic programs of epithelial cell plasticity directed by transforming growth factor-β. Proc Natl Acad Sci U S A 98, 6686–6691 (2001).

42. Qin, Z., Xia, W., Fisher, G. J., Voorhees, J. J. & Quan, T. YAP/TAZ regulates TGF-β/Smad3 signaling by induction of Smad7 via AP-1 in human skin dermal fibroblasts. Cell Commun. Signal 16, 18 (2018).

43. Thao, T. T. N. et al. Rapid reconstruction of SARS-CoV-2 using a synthetic genomics platform. Nature 1–8 (2020) doi:10.1038/s41586-020-2294-9.

44. Mantlo, E., Bukreyeva, N., Maruyama, J., Paessler, S. & Huang, C. Antiviral activities of type I interferons to SARS-CoV-2 infection. Antiviral Res 179, 104811 (2020).

45. Seebacher, N. A., Stacy, A. E., Porter, G. M. & Merlot, A. M. Clinical development of targeted and immune based anti-cancer therapies. Journal of Experimental & Clinical Cancer Research 38, 156 (2019).

46. O’Shea, J. J., Kontzias, A., Yamaoka, K., Tanaka, Y. & Laurence, A. Janus kinase Inhibitors in autoimmune diseases. Ann Rheum Dis 72, ii111–ii115 (2013).

47. Yong, H.-Y., Koh, M.-S. & Moon, A. The p38 MAPK inhibitors for the treatment of inflammatory diseases and cancer. Expert Opinion on Investigational Drugs 18, 1893–1905 (2009).

48. Hsieh, W.-Y. et al. ACE/ACE2 ratio and MMP-9 activity as potential biomarkers in tuberculous pleural effusions. Int J Biol Sci 8, 1197–1205 (2012).

49. Ueland, T. et al. Distinct and early increase in circulating MMP-9 in COVID-19 patients with respiratory failure. J Infect 81, e41–e43 (2020).

50. Villalta, P. C., Rocic, P. & Townsley, M. I. Role of MMP2 and MMP9 in TRPV4-induced lung injury. Am J Physiol Lung Cell Mol Physiol 307, L652–L659 (2014).

51. Marten, N. W. & Zhou, J. The Role of Metalloproteinases in Corona Virus Infection. Experimental Models of Multiple Sclerosis 839–848 (2005) doi:10.1007/0-387-25518-4_48.

52. Zhang, J.-Y. et al. Single-cell landscape of immunological responses in patients with COVID-19. Nature Immunology 21, 1107–1118 (2020).

53. Genomewide Association Study of Severe Covid-19 with Respiratory Failure. New England Journal of Medicine 383, 1522–1534 (2020).

54. Ali, A. & Vijayan, R. Dynamics of the ACE2–SARS-CoV-2/SARS-CoV spike protein interface reveal unique mechanisms. Scientific Reports 10, 14214 (2020).

55. Hubel, P. et al. A protein-interaction network of interferon-stimulated genes extends the innate immune system landscape. Nat. Immunol. 20, 493–502 (2019).

56. Scaturro, P. et al. An orthogonal proteomic survey uncovers novel Zika virus host factors. Nature 561, 253–257 (2018).

57. Hoffmann, M. et al. SARS-CoV-2 Cell Entry Depends on ACE2 and TMPRSS2 and Is Blocked by a Clinically Proven Protease Inhibitor. Cell 181, 271–280.e8 (2020).

58. Gebhardt, A. et al. The alternative cap-binding complex is required for antiviral defense in vivo. PLoS Pathog. 15, e1008155 (2019).

59. Goldeck, M., Schlee, M., Hartmann, G. & Hornung, V. Enzymatic synthesis and purification of a defined RIG-I ligand. Methods Mol. Biol. 1169, 15–25 (2014).

60. Kulak, N. A., Geyer, P. E. & Mann, M. Loss-less Nano-fractionator for High Sensitivity, High Coverage Proteomics. Mol. Cell Proteomics 16, 694–705 (2017).

61. Kulak, N. A., Pichler, G., Paron, I., Nagaraj, N. & Mann, M. Minimal, encapsulated proteomic-sample processing applied to copy-number estimation in eukaryotic cells. Nat. Methods 11, 319–324 (2014).

62. Udeshi, N. D., Mertins, P., Svinkina, T. & Carr, S. A. Large-scale identification of ubiquitination sites by mass spectrometry. Nat Protoc 8, 1950–1960 (2013).

63. Tyanova, S., Temu, T. & Cox, J. The MaxQuant computational platform for mass spectrometry-based shotgun proteomics. Nat Protoc 11, 2301–2319 (2016).

64. Alexey Stukalov. innatelab/cov2_utils_jl v1.0.0. (Zenodo, 2021). doi:10.5281/zenodo.4541090.

65. Bober, M. & Miladinovic, S. General guidelines for validation of decoy models for HRM/DIA/SWATH as exemplified using spectronaut. F1000posters.

66. Alexey Stukalov. innatelab/cov2 v1.0.0. (Zenodo, 2021). doi:10.5281/zenodo.4541082.

67. Alexey Stukalov. innatelab/maxquantUtils v0.1.0. (Zenodo, 2021). doi:10.5281/zenodo.4536603.

68. Alexey Stukalov. innatelab/msglm v0.5.0. (Zenodo, 2021). doi:10.5281/zenodo.4536605.

69. Carpenter, B. et al. Stan: A Probabilistic Programming Language. Journal of Statistical Software 76, 1–32 (2017).

70. Bhadra, A., Datta, J., Polson, N. G. & Willard, B. The Horseshoe+ Estimator of Ultra-Sparse Signals. Bayesian Anal. 12, 1105–1131 (2017).

71. Goeminne, L. J. E., Gevaert, K. & Clement, L. Peptide-level Robust Ridge Regression Improves Estimation, Sensitivity, and Specificity in Data-dependent Quantitative Label-free Shotgun Proteomics. Mol. Cell Proteomics 15, 657–668 (2016).

72. Shannon, P. et al. Cytoscape: A Software Environment for Integrated Models of Biomolecular Interaction Networks. Genome Res 13, 2498–2504 (2003).

73. Ritchie, M. E. et al. limma powers differential expression analyses for RNA-sequencing and microarray studies. Nucleic Acids Res 43, e47 (2015).

74. Merico, D., Isserlin, R., Stueker, O., Emili, A. & Bader, G. D. Enrichment map: a network-based method for gene-set enrichment visualization and interpretation. PLoS ONE 5, e13984 (2010).

75. Meldal, B. H. M. et al. Complex Portal 2018: extended content and enhanced visualization tools for macromolecular complexes. Nucleic Acids Res. 47, D550–D558 (2019).

76. Giurgiu, M. et al. CORUM: the comprehensive resource of mammalian protein complexes-2019. Nucleic Acids Res. 47, D559–D563 (2019).

77. Tyanova, S. et al. The Perseus computational platform for comprehensive analysis of (prote)omics data. Nat. Methods 13, 731–740 (2016).

78. Keenan, A. B. et al. ChEA3: transcription factor enrichment analysis by orthogonal omics integration. Nucleic Acids Res. 47, W212–W224 (2019).

79. Landt, S. G. et al. ChIP-seq guidelines and practices of the ENCODE and modENCODE consortia. Genome Res. 22, 1813–1831 (2012).

80. Alexey Stukalov. alyst/OptEnrichedSetCover.jl v0.5.0. (Zenodo, 2021). doi:10.5281/zenodo.4536596.

81. Alexey Stukalov. alyst/HierarchicalHotNet.jl v1.0.0. (Zenodo, 2021). doi:10.5281/zenodo.4536590.

82. Meng, E. C., Pettersen, E. F., Couch, G. S., Huang, C. C. & Ferrin, T. E. Tools for integrated sequence-structure analysis with UCSF Chimera. BMC Bioinformatics 7, 339 (2006).

83. Pettersen, E. F. et al. UCSF Chimera--a visualization system for exploratory research and analysis. J Comput Chem 25, 1605–1612 (2004).

84. Warnecke, A., Sandalova, T., Achour, A. & Harris, R. A. PyTMs: a useful PyMOL plugin for modeling common post-translational modifications. BMC Bioinformatics 15, (2014).

85. Paxman, J. J. & Heras, B. Bioinformatics Tools and Resources for Analyzing Protein Structures. Methods Mol Biol 1549, 209–220 (2017).

86. Jo, S., Vargyas, M., Vasko-Szedlar, J., Roux, B. & Im, W. PBEQ-Solver for online visualization of electrostatic potential of biomolecules. Nucleic Acids Res 36, W270–275 (2008).

87. Vogt, C. et al. The interferon antagonist ML protein of thogoto virus targets general transcription factor IIB. J. Virol. 82, 11446–11453 (2008).

88. Jorns, C. et al. Rapid and simple detection of IFN-neutralizing antibodies in chronic hepatitis C non-responsive to IFN-alpha. J. Med. Virol. 78, 74–82 (2006).

89. Perez-Riverol, Y. et al. The PRIDE database and related tools and resources in 2019: improving support for quantification data. Nucleic Acids Res 47, D442–D450 (2019).

90. Orchard, S. et al. The MIntAct project—IntAct as a common curation platform for 11 molecular interaction databases. Nucleic Acids Res 42, D358–D363 (2014).

