## Supplementary_Data_1_Diffusion networks of SARS-CoV-2 and SARS-CoV proteins for "Multilevel proteomics reveals host perturbations by SARS-CoV-2 and SARS-CoV": readme.txt

The archive contains the output of the network diffusion analysis of AP-MS
and viral protein overexpression data generated by HierarchicalHotNet.jl package.

For each viral protein XXX it contains:
1. cov2_XXX_flow_20200609.graphml: the subnetwork of the full ReactomeFI-based network diffusion network,
   enriched with paths from protein perturbed by viral protein expression to its host targets.,,
   The network is in GraphML format, best viewed with yEd v3.20 (https://www.yworks.com/products/yed)
2. cov2_XXX_flow_distance_20200609.pdf: the comparison of the average path length in the real data to 1000 random permutations of the data.
   See Materials and Methods and Extended Figure 3b for the description of the plot and the randomization process.
3. cov2_XXX_maxcomponent_size_20200609.pdf: the comparison of the size of maximal strongly connected component for the real data and for the 1000 random permutations.
   See the original HierarchicalHotNet paper (https://academic.oup.com/bioinformatics/article/34/17/i972/5093236) for the description of the plot.
   
If the network diffusion analysis didn't identify any enriched subnetwork for a given viral protein, the corresponding files would be empty.
