## Supplementary figures and images for "Multilevel proteomics reveals host perturbations by SARS-CoV-2 and SARS-CoV"

### cov2_SARS_CoV_ORF8_flow_distance_20200609.pdf

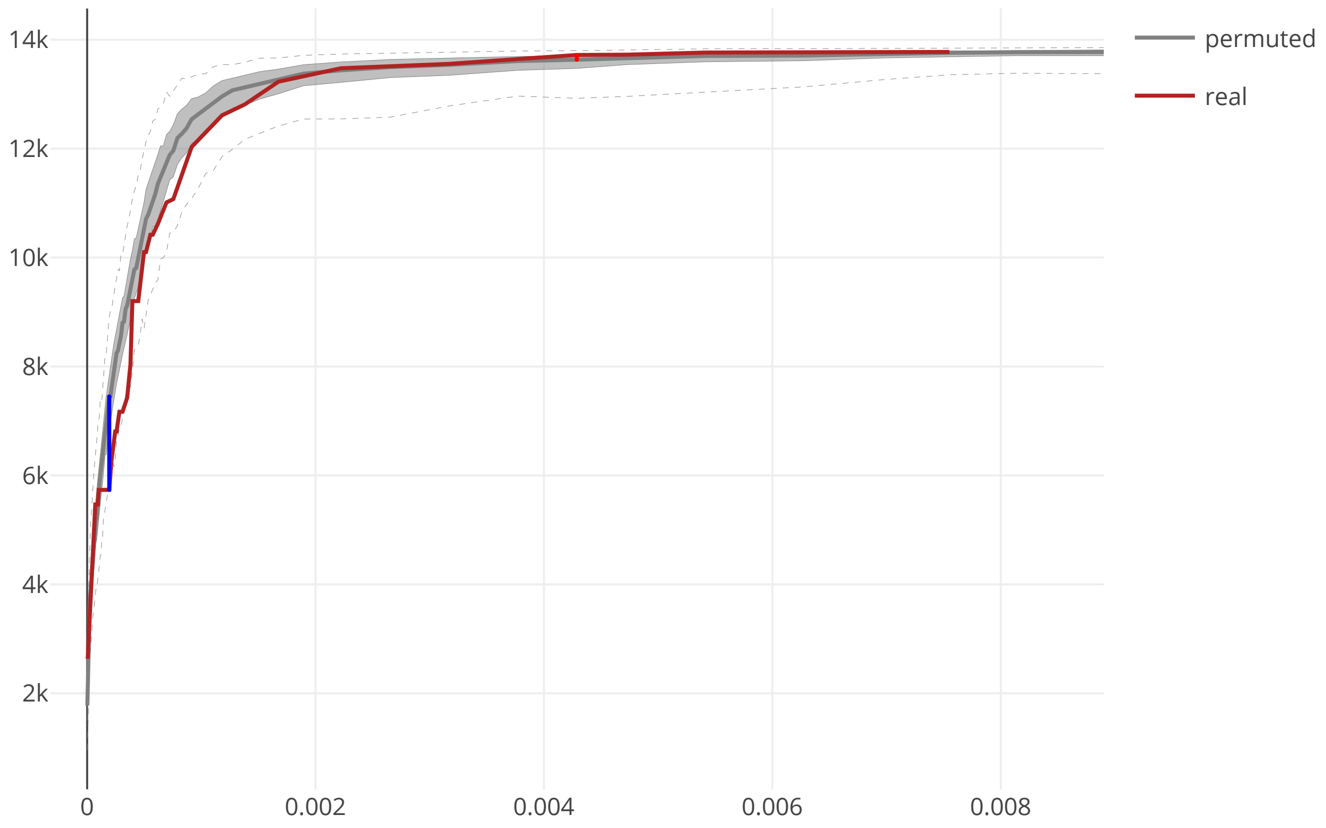

### cov2_SARS_CoV_ORF8_maxcomponent_size_20200609.pdf

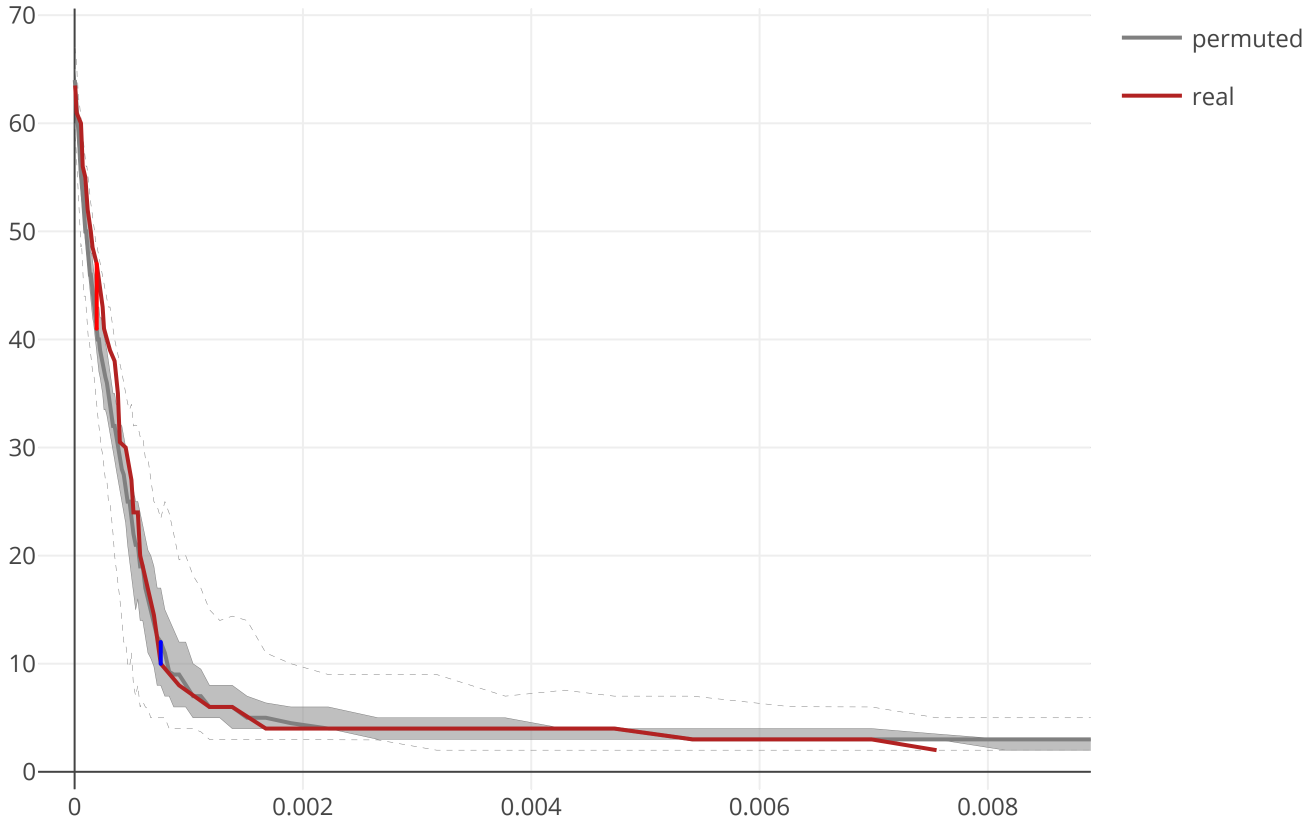

### cov2_SARS_CoV_ORF9b_flow_distance_20200609.pdf

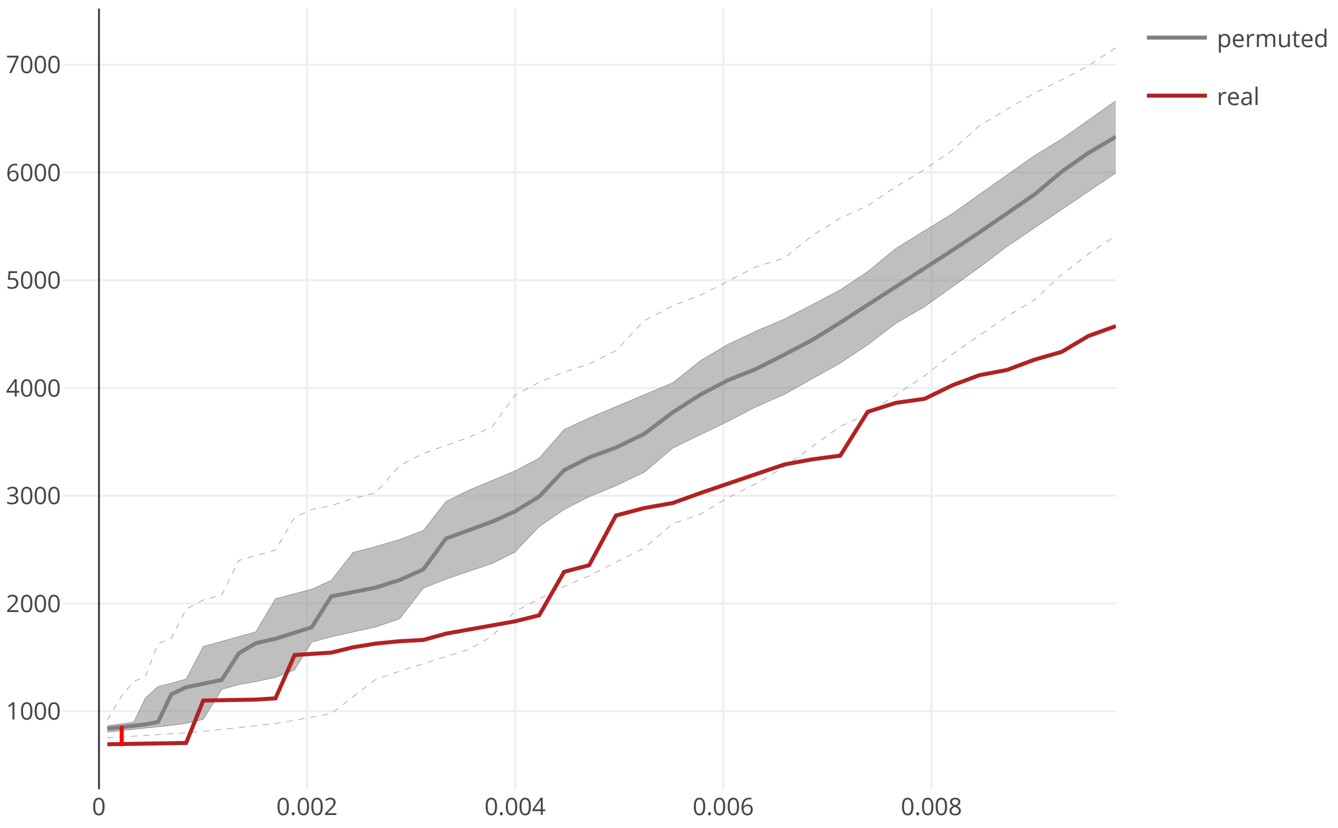

### cov2_SARS_CoV_ORF9b_maxcomponent_size_20200609.pdf

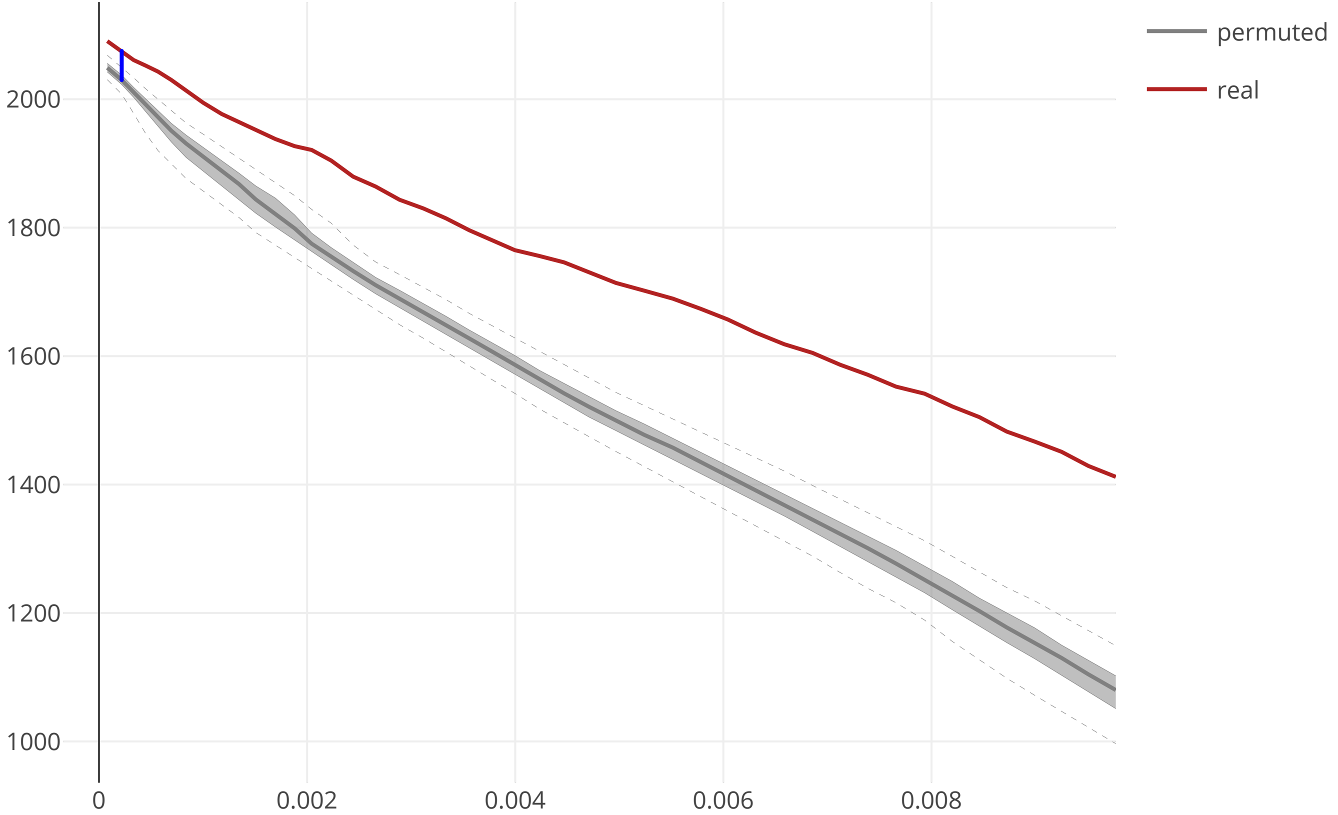

### cov2_SARS_CoV_S_flow_distance_20200609.pdf

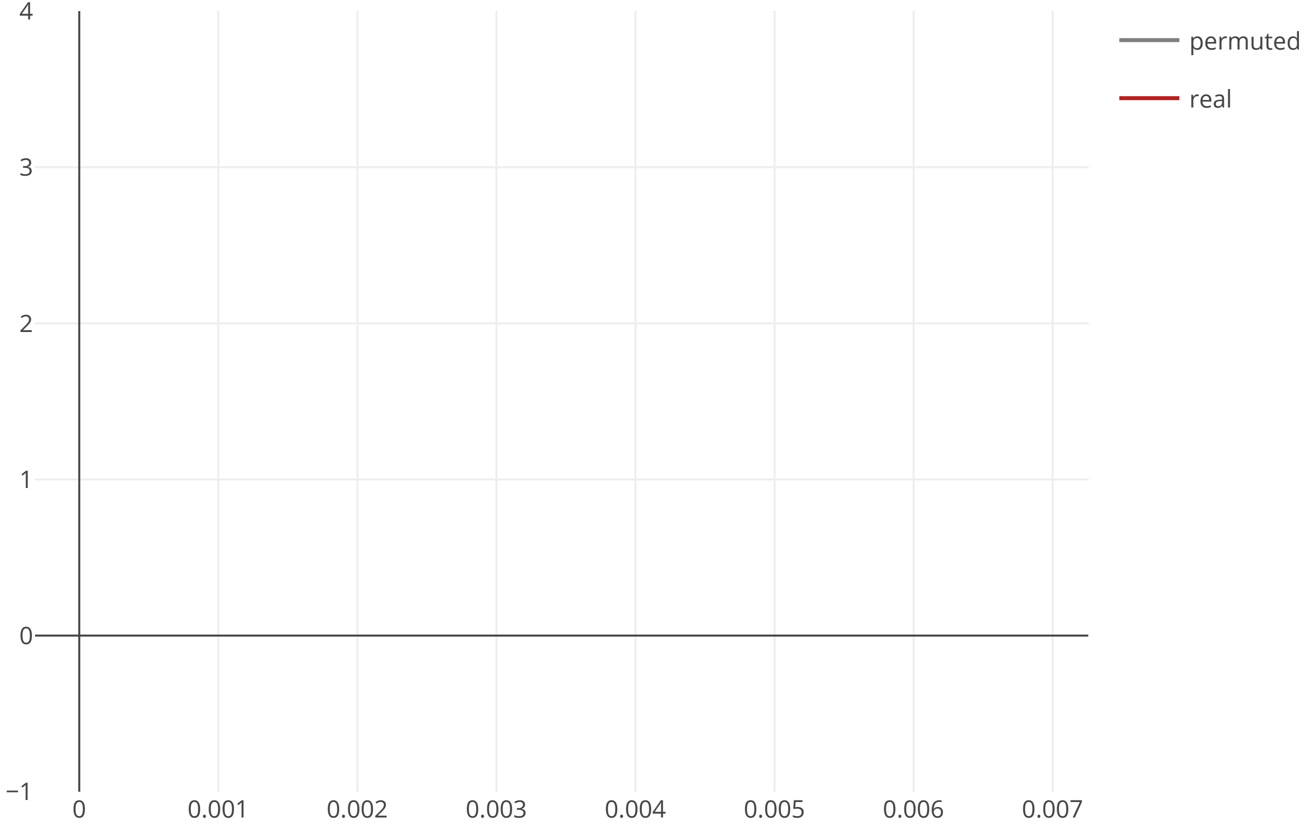

### cov2_SARS_CoV_S_maxcomponent_size_20200609.pdf

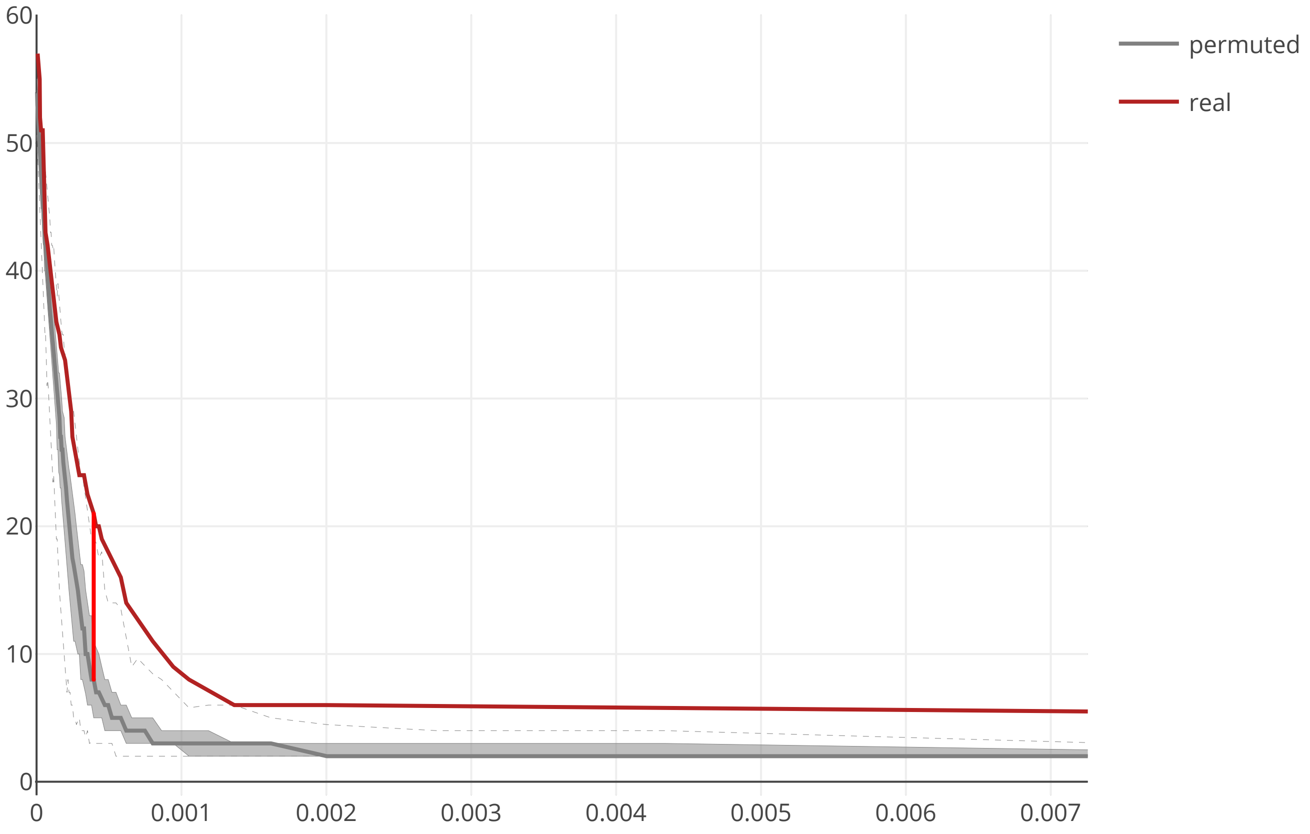
