## Supplementary figures and images for "Multilevel proteomics reveals host perturbations by SARS-CoV-2 and SARS-CoV"

### cov2_HCoV_ORF3_flow_distance_20200609.pdf

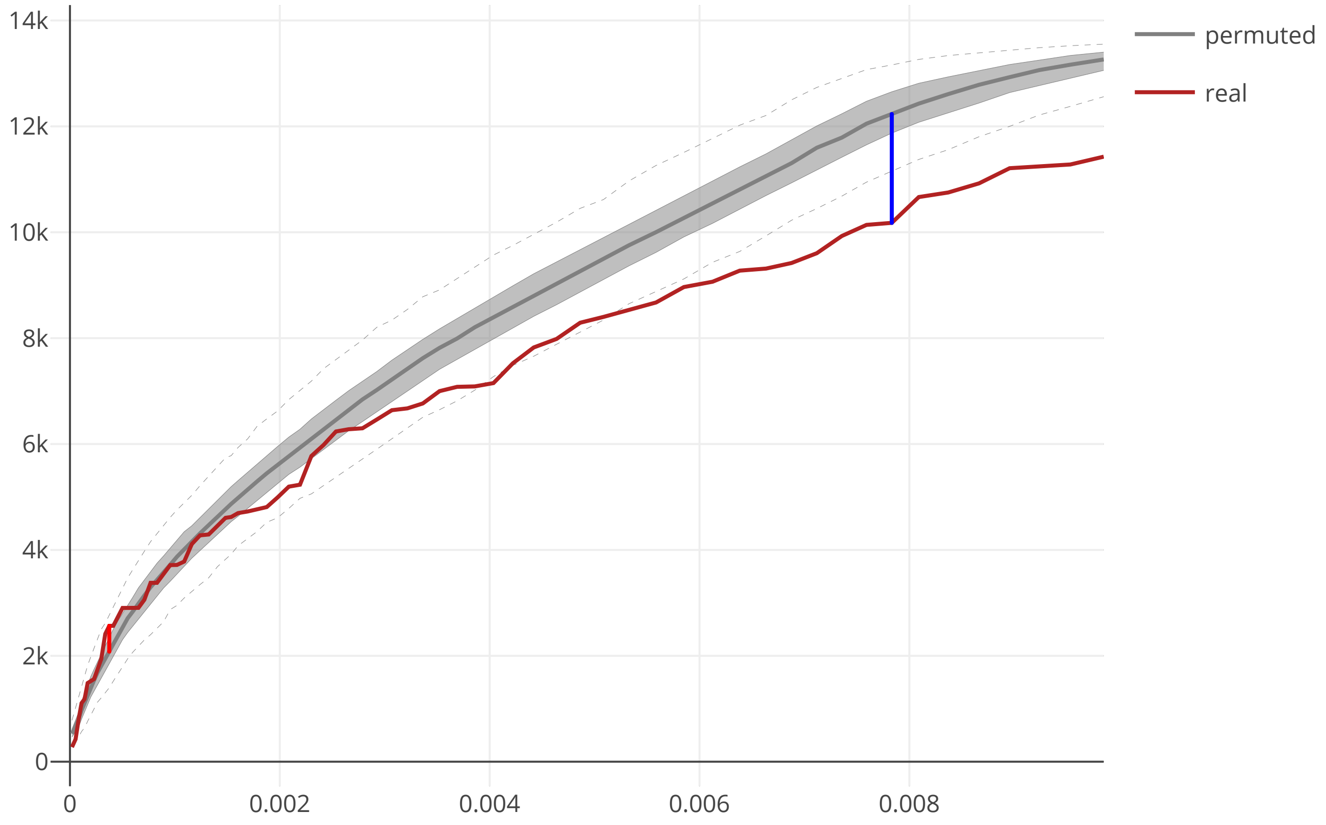

### cov2_HCoV_ORF3_maxcomponent_size_20200609.pdf

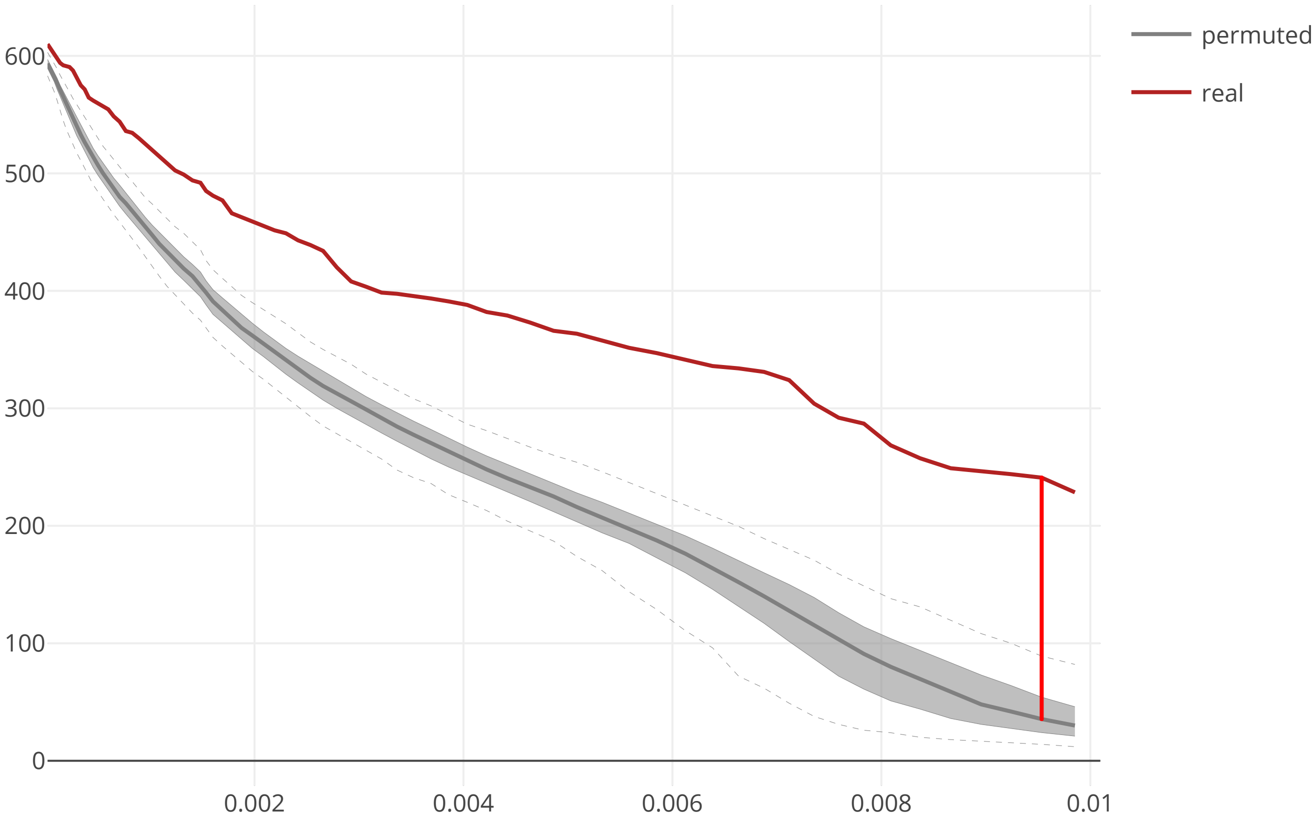

### cov2_HCoV_ORF4_flow_distance_20200609.pdf

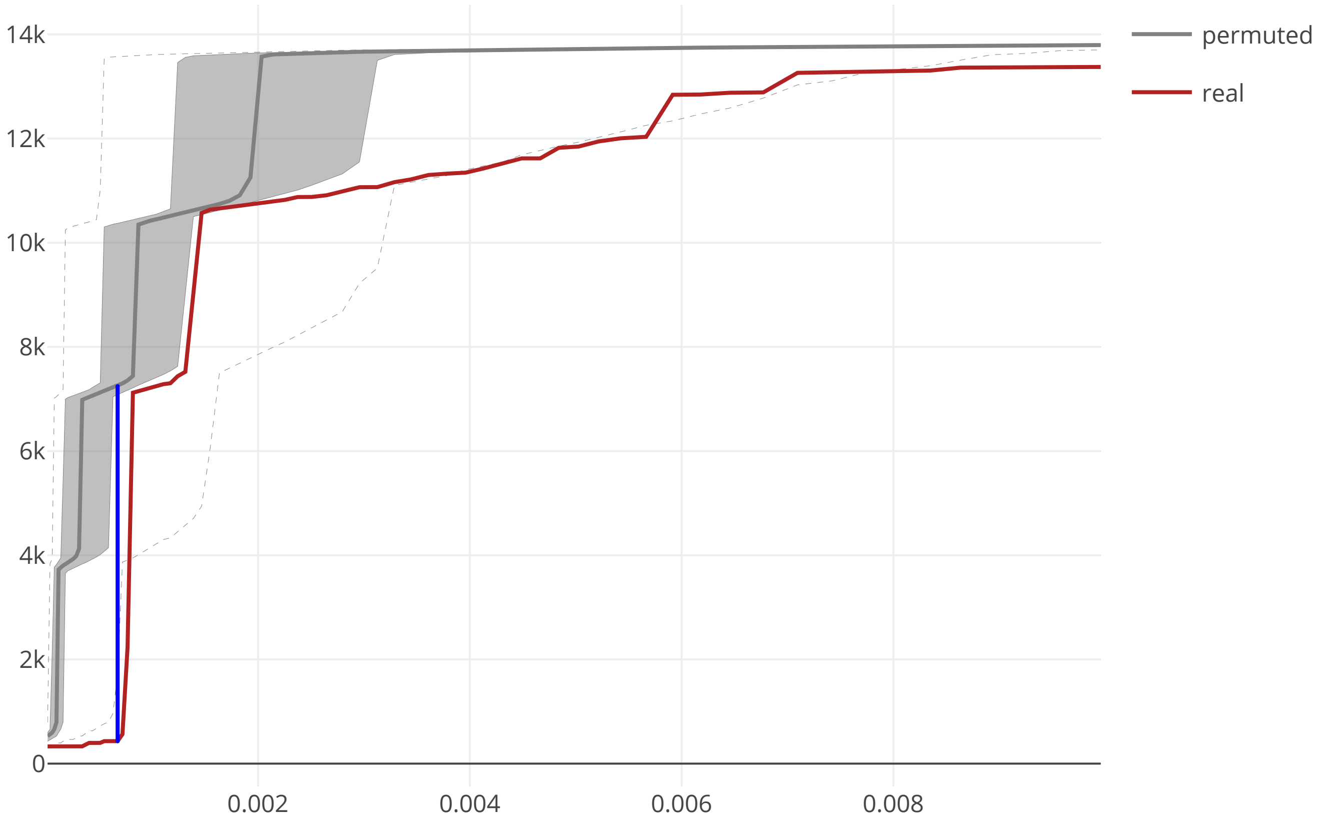

### cov2_HCoV_ORF4_maxcomponent_size_20200609.pdf

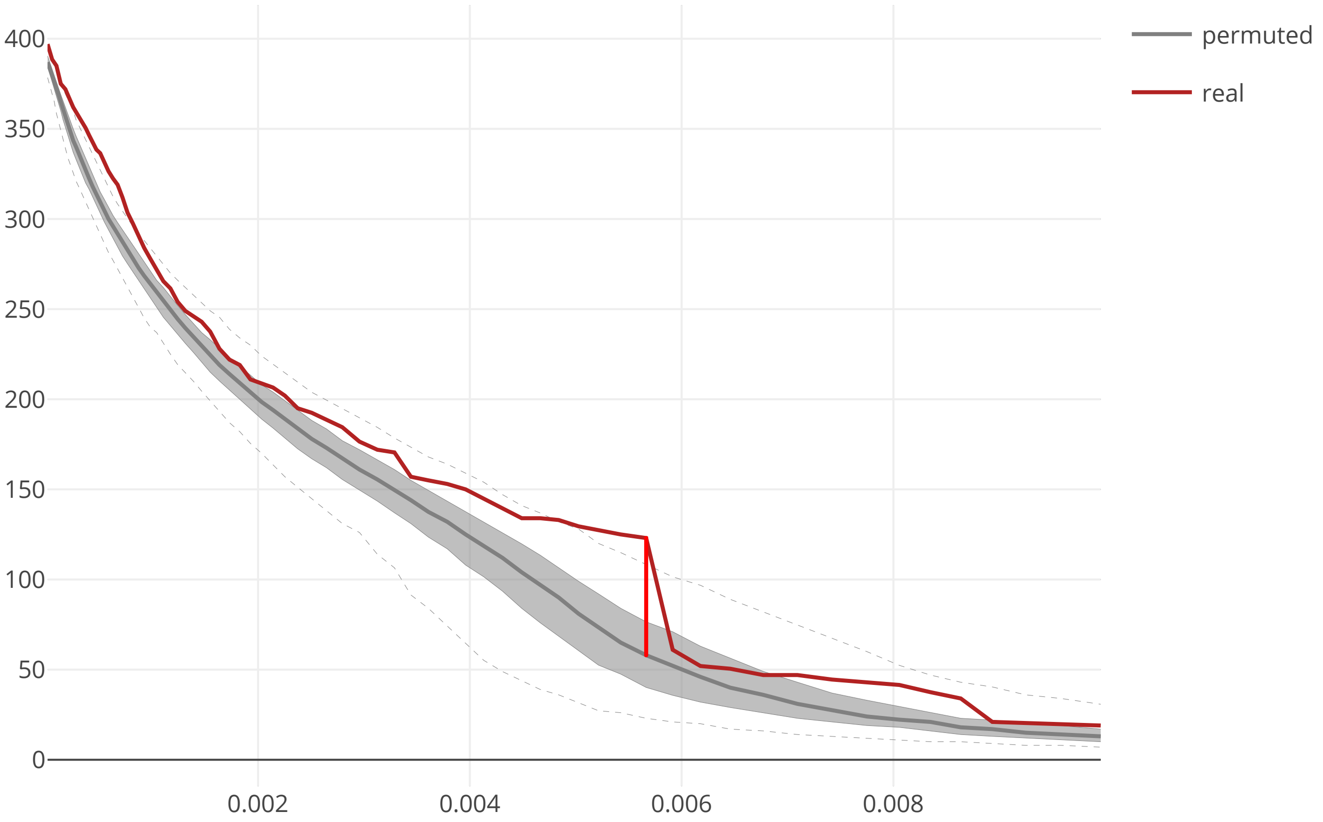

### cov2_HCoV_ORF4a_flow_distance_20200609.pdf

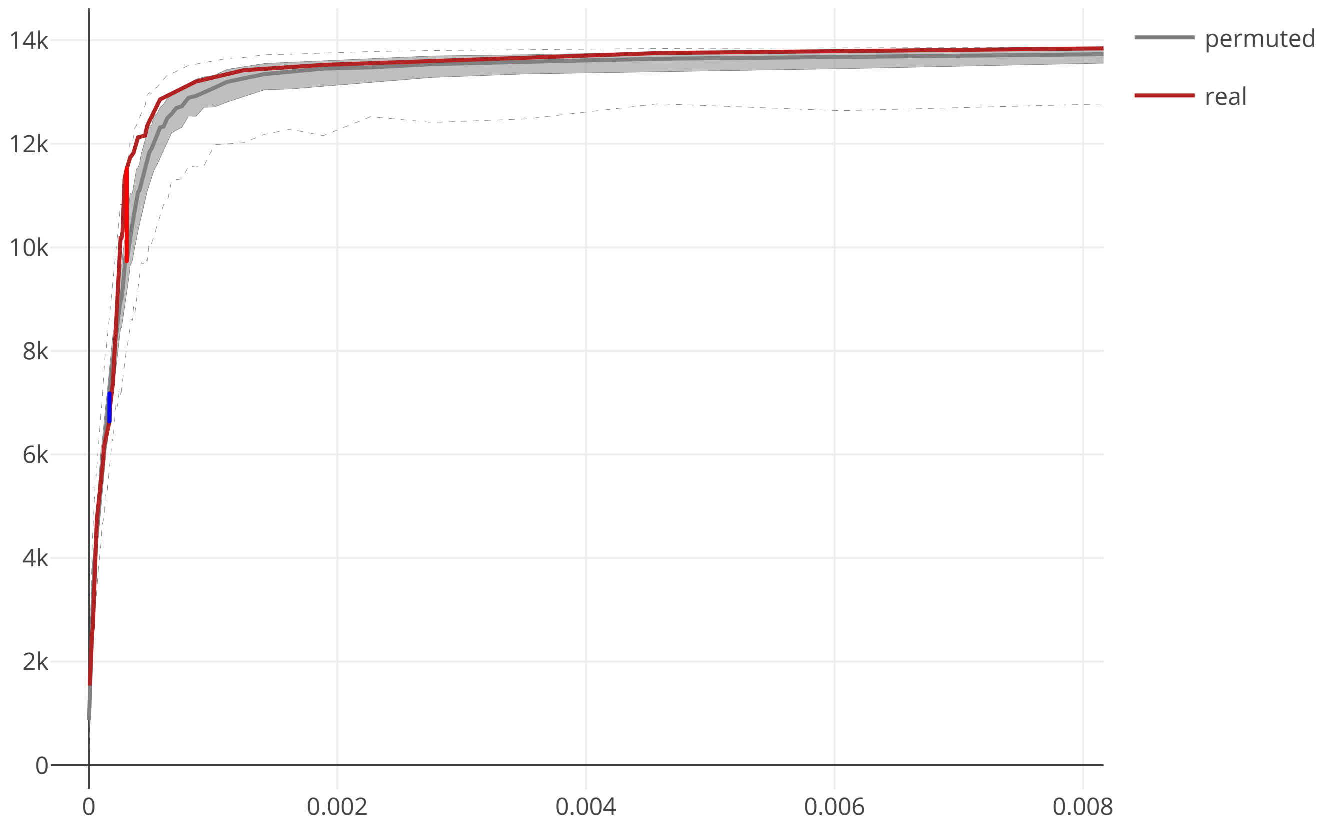

### cov2_HCoV_ORF4a_maxcomponent_size_20200609.pdf

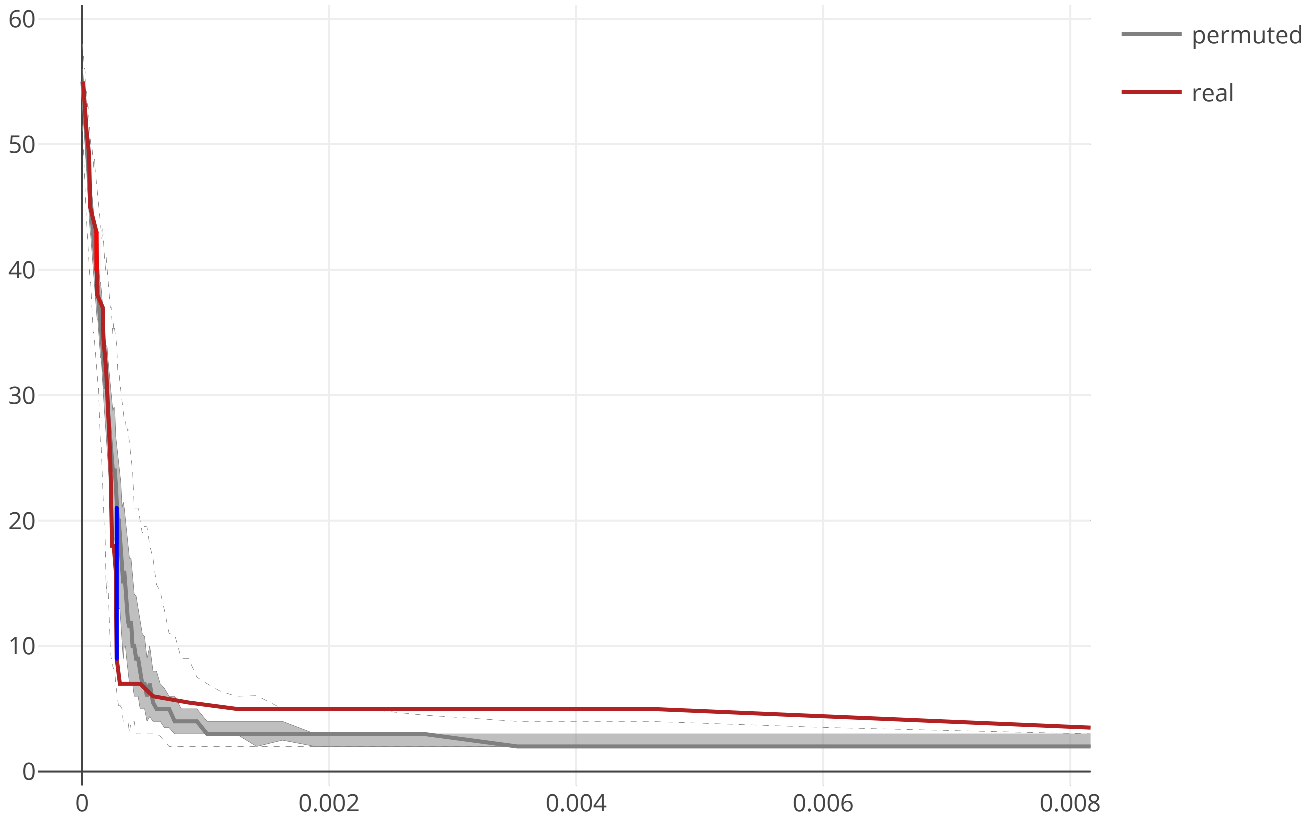

### cov2_SARS_CoV2_E_flow_distance_20200609.pdf

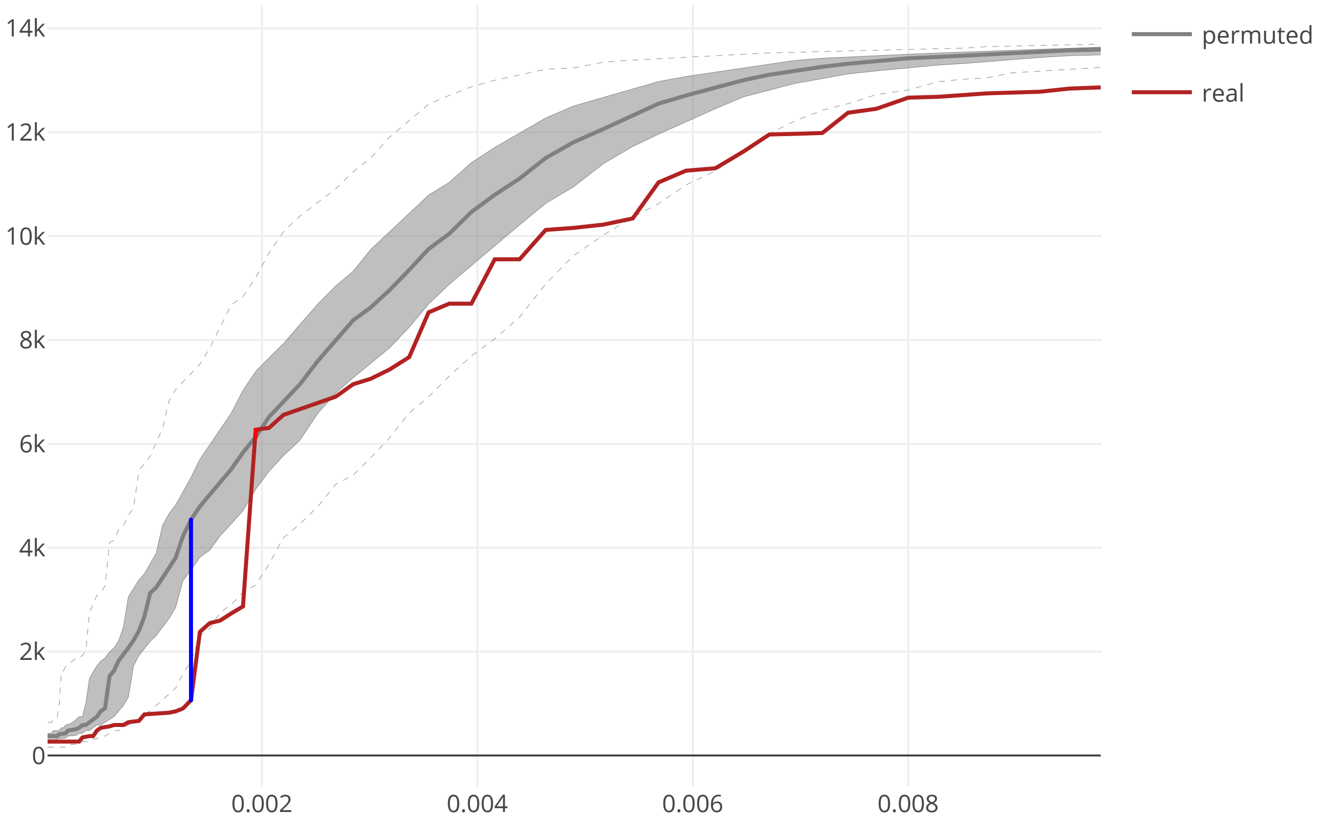

### cov2_SARS_CoV2_E_maxcomponent_size_20200609.pdf

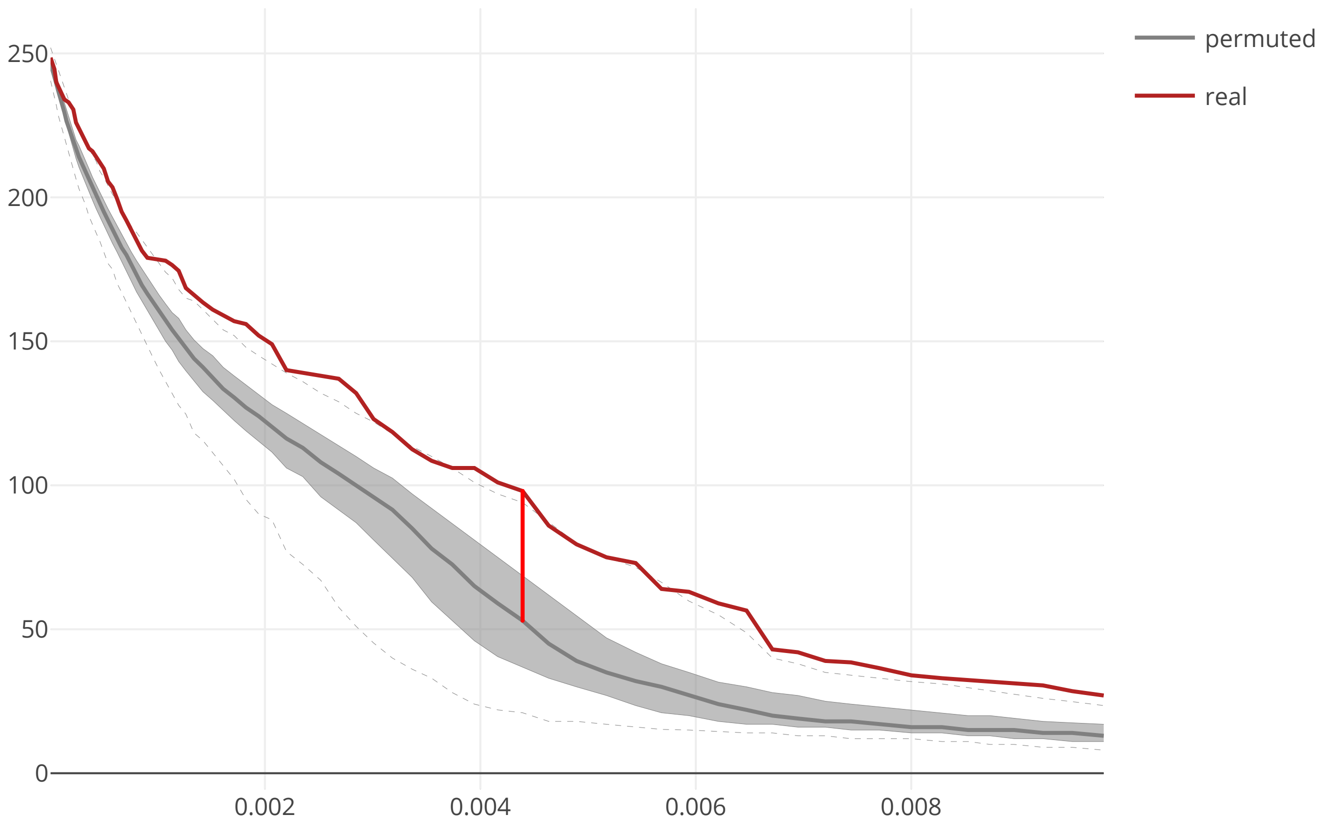

### cov2_SARS_CoV2_M_flow_distance_20200609.pdf

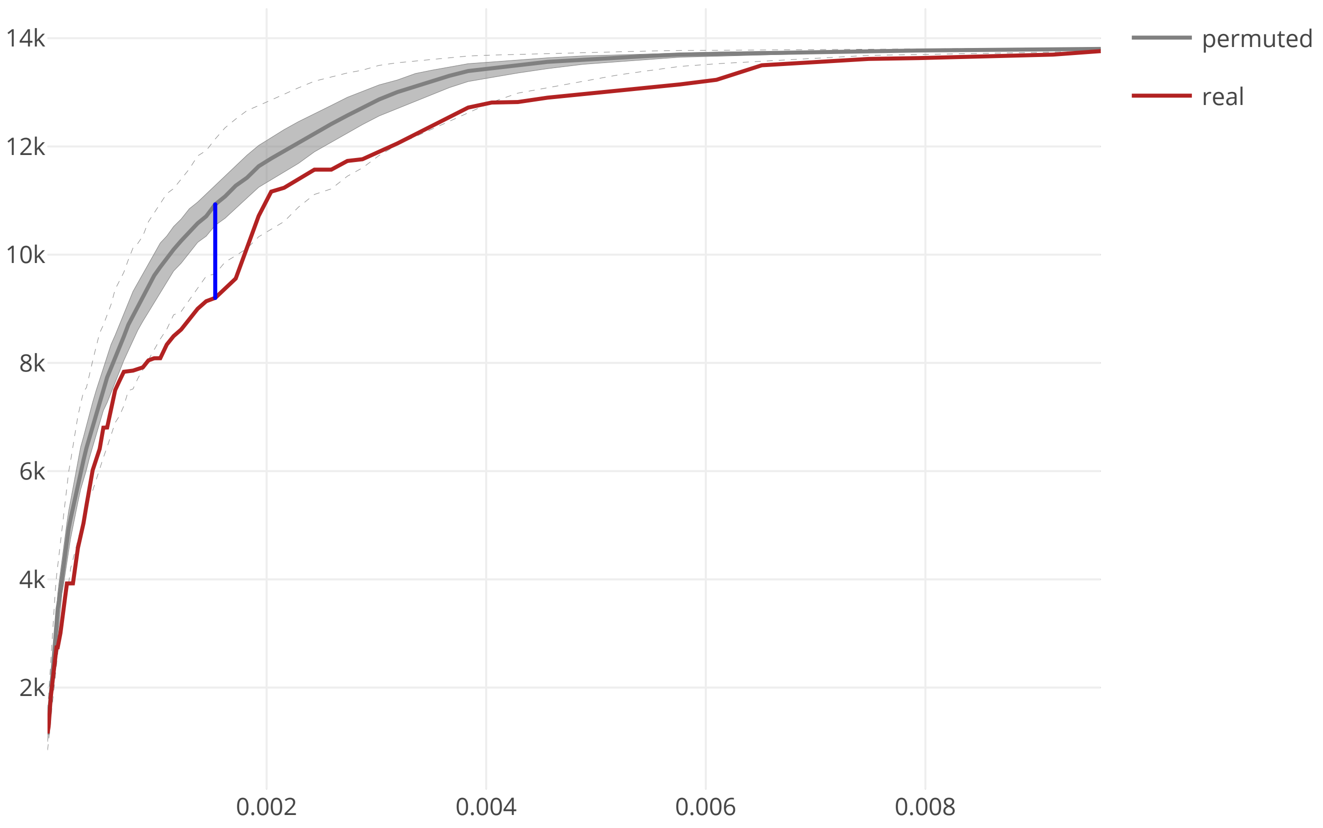

### cov2_SARS_CoV2_M_maxcomponent_size_20200609.pdf

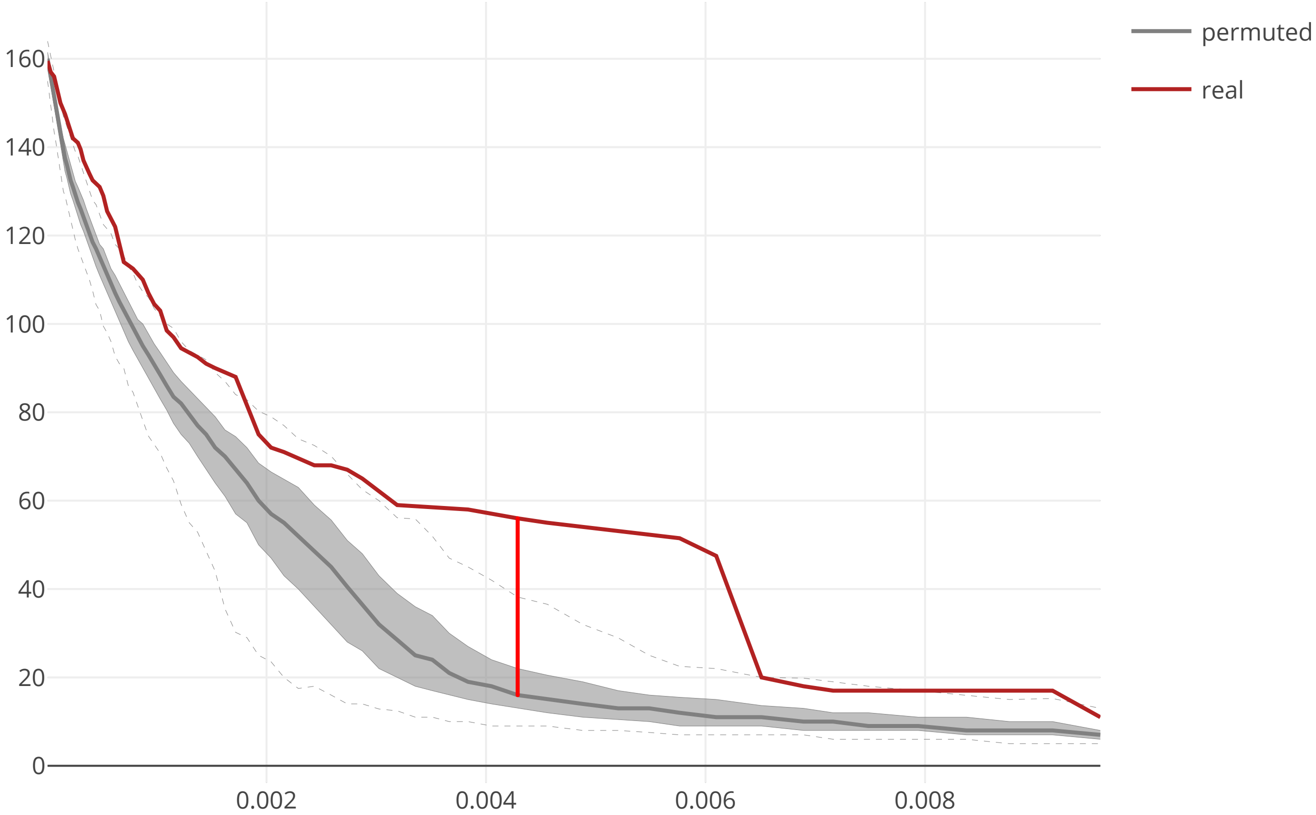

### cov2_SARS_CoV2_N_flow_distance_20200609.pdf

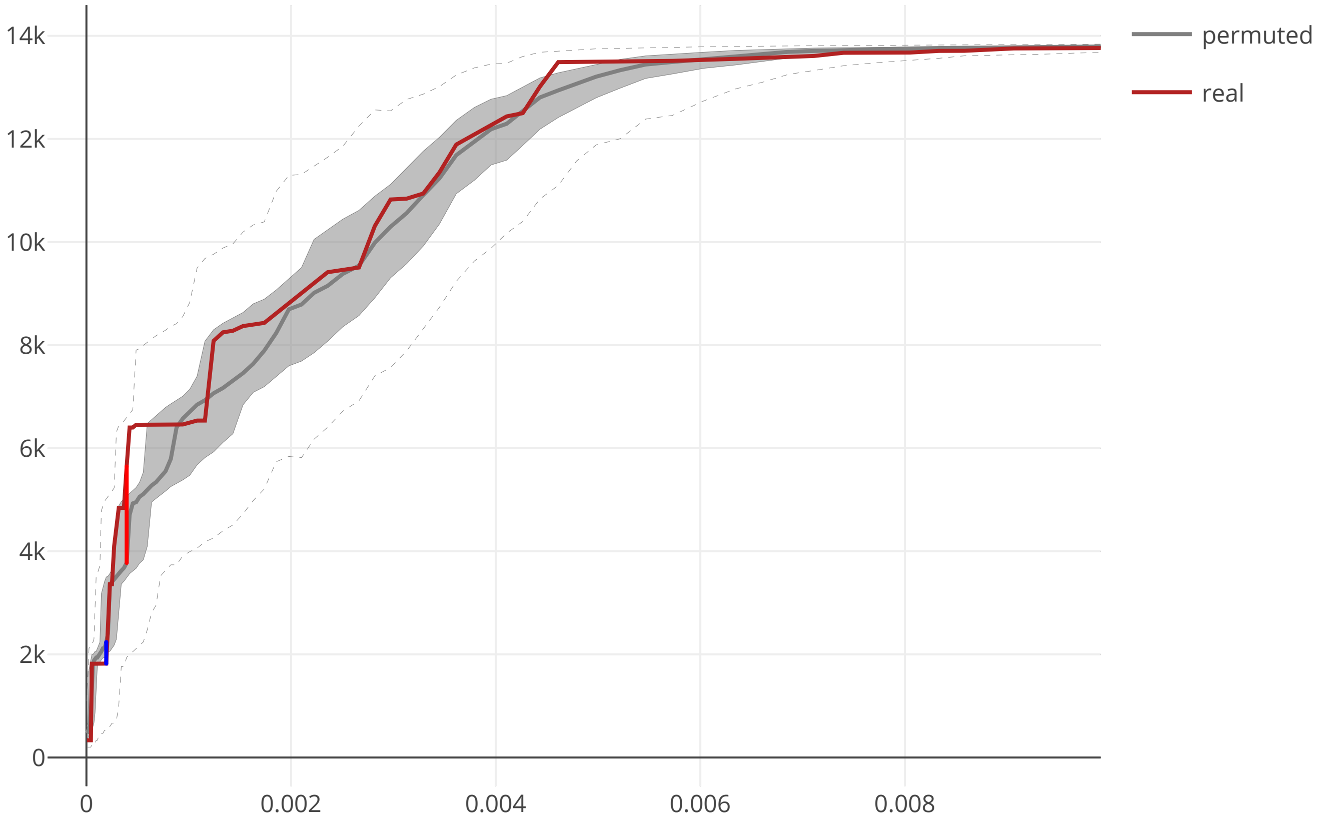

### cov2_SARS_CoV2_N_maxcomponent_size_20200609.pdf

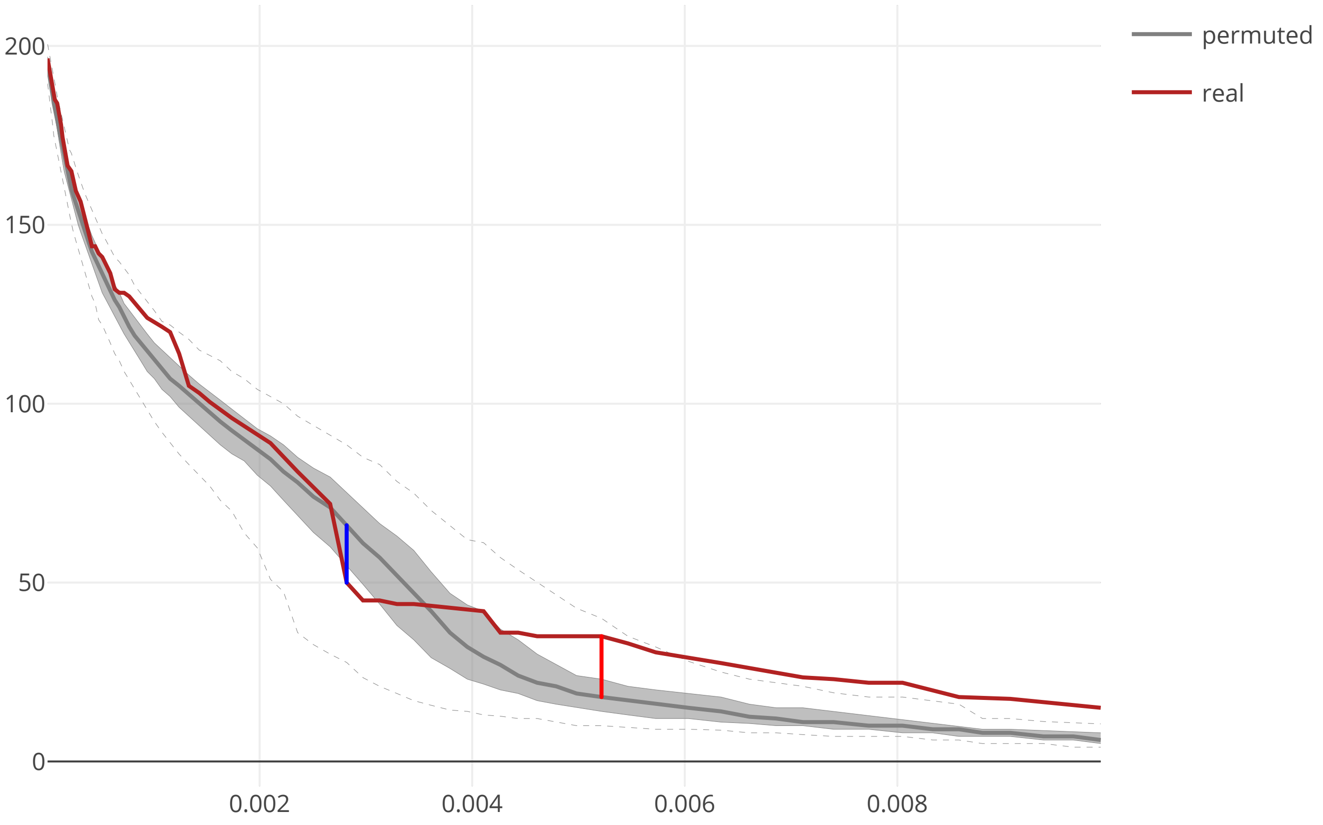

### cov2_SARS_CoV2_NSP1_flow_distance_20200609.pdf

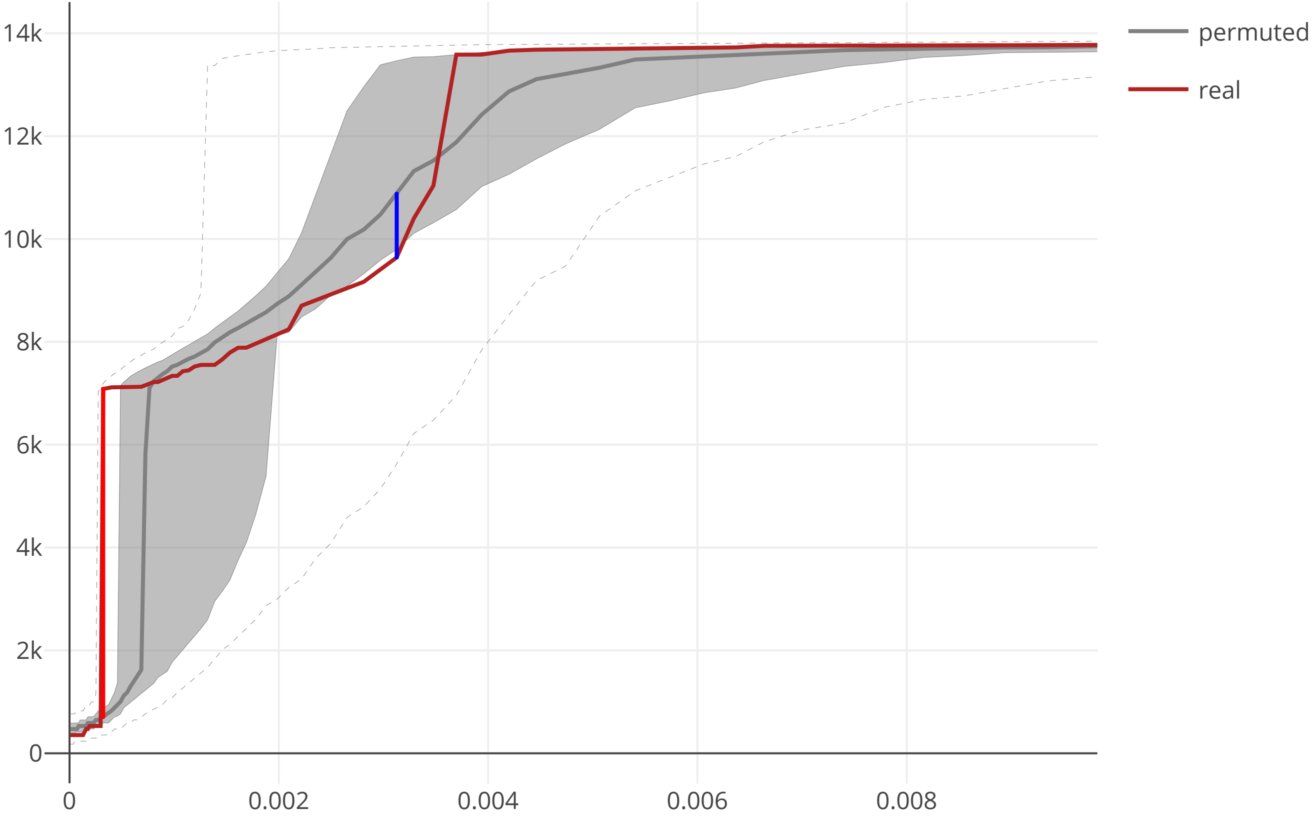

### cov2_SARS_CoV2_NSP1_maxcomponent_size_20200609.pdf

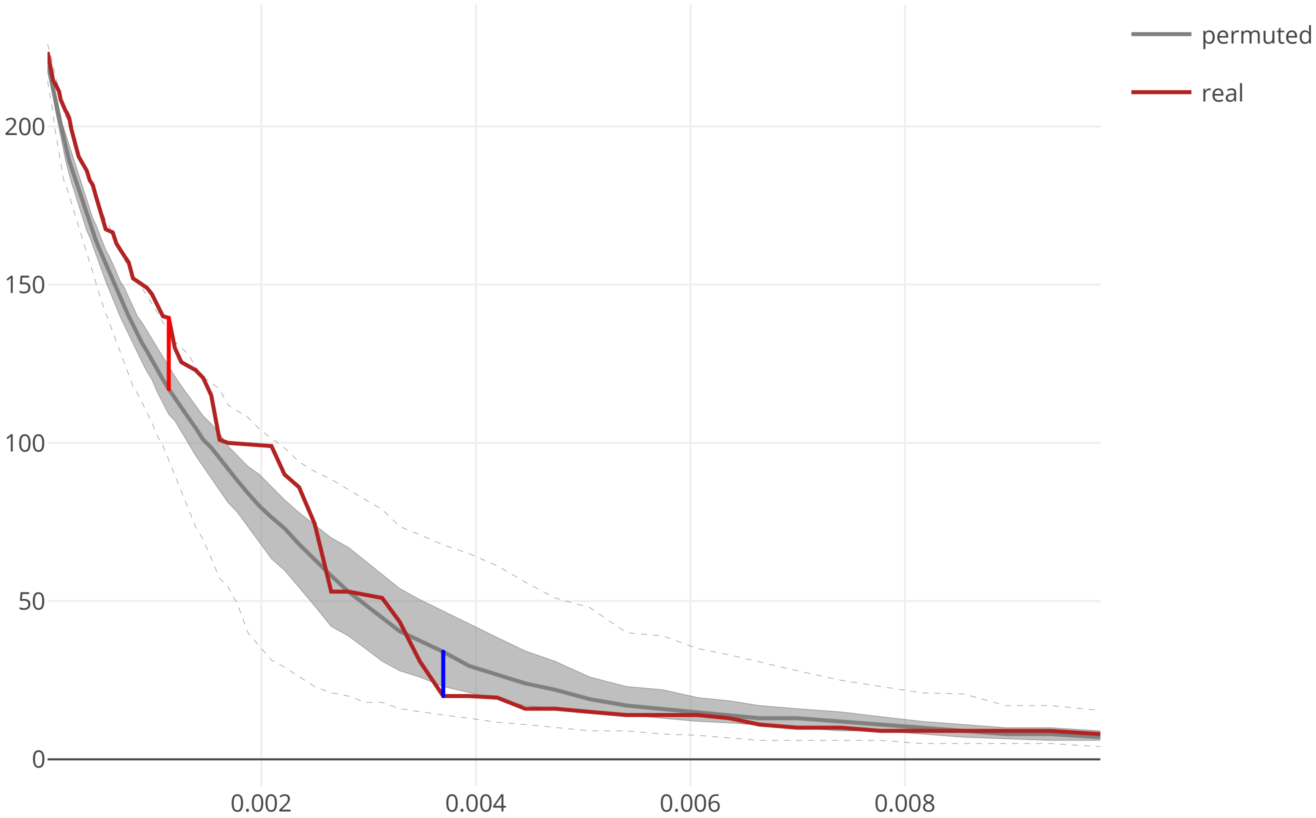

### cov2_SARS_CoV2_NSP2_flow_distance_20200609.pdf

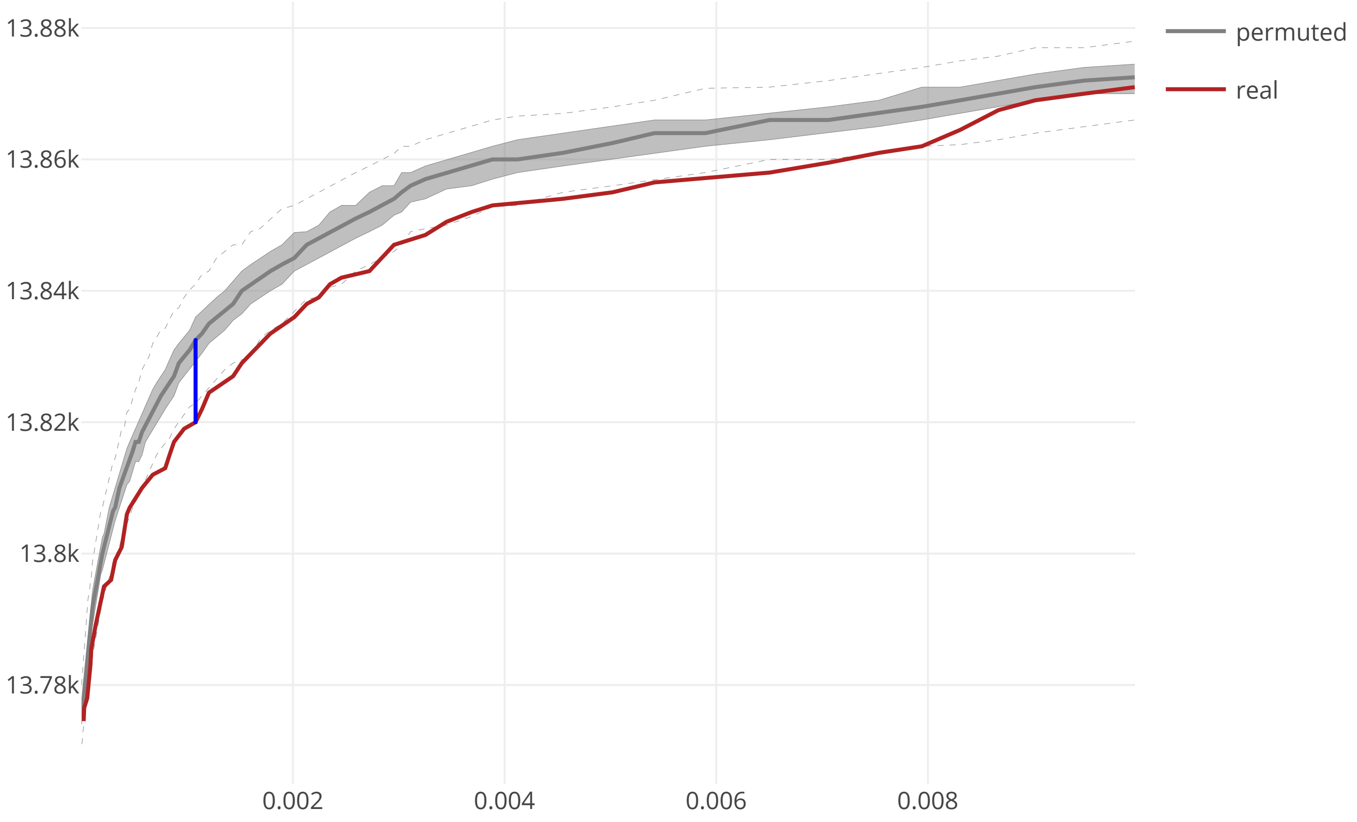

### cov2_SARS_CoV2_NSP2_maxcomponent_size_20200609.pdf

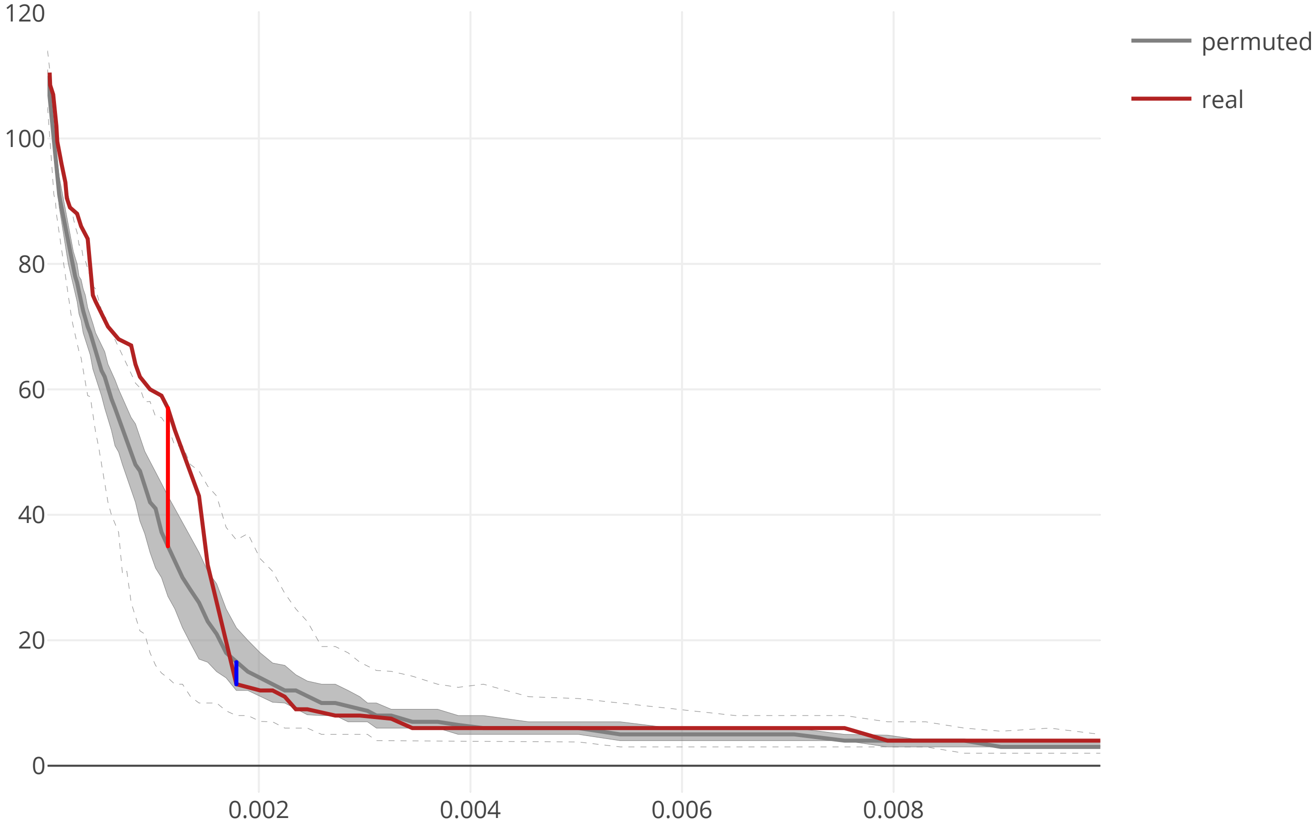

### cov2_SARS_CoV2_NSP3_macroD_flow_distance_20200609.pdf

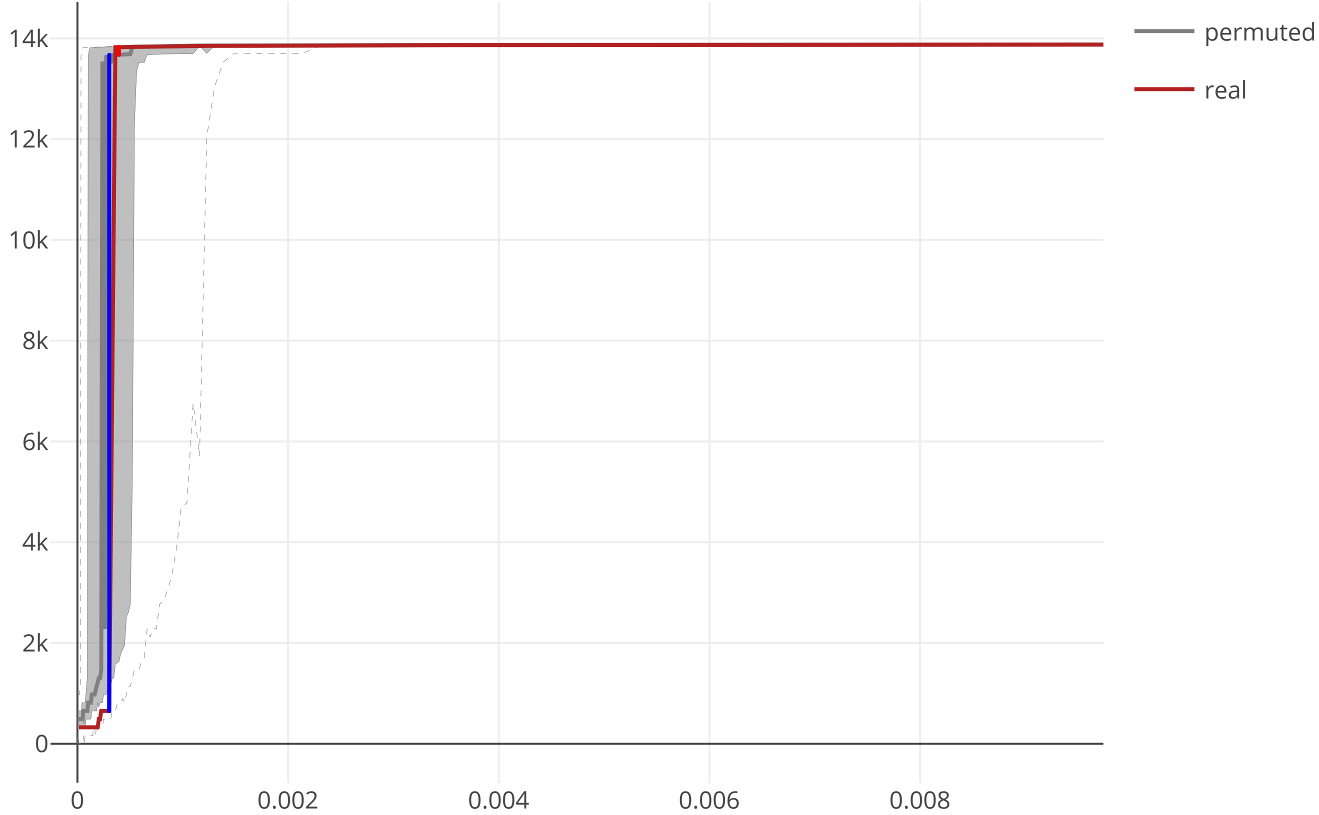

### cov2_SARS_CoV2_NSP3_macroD_maxcomponent_size_20200609.pdf

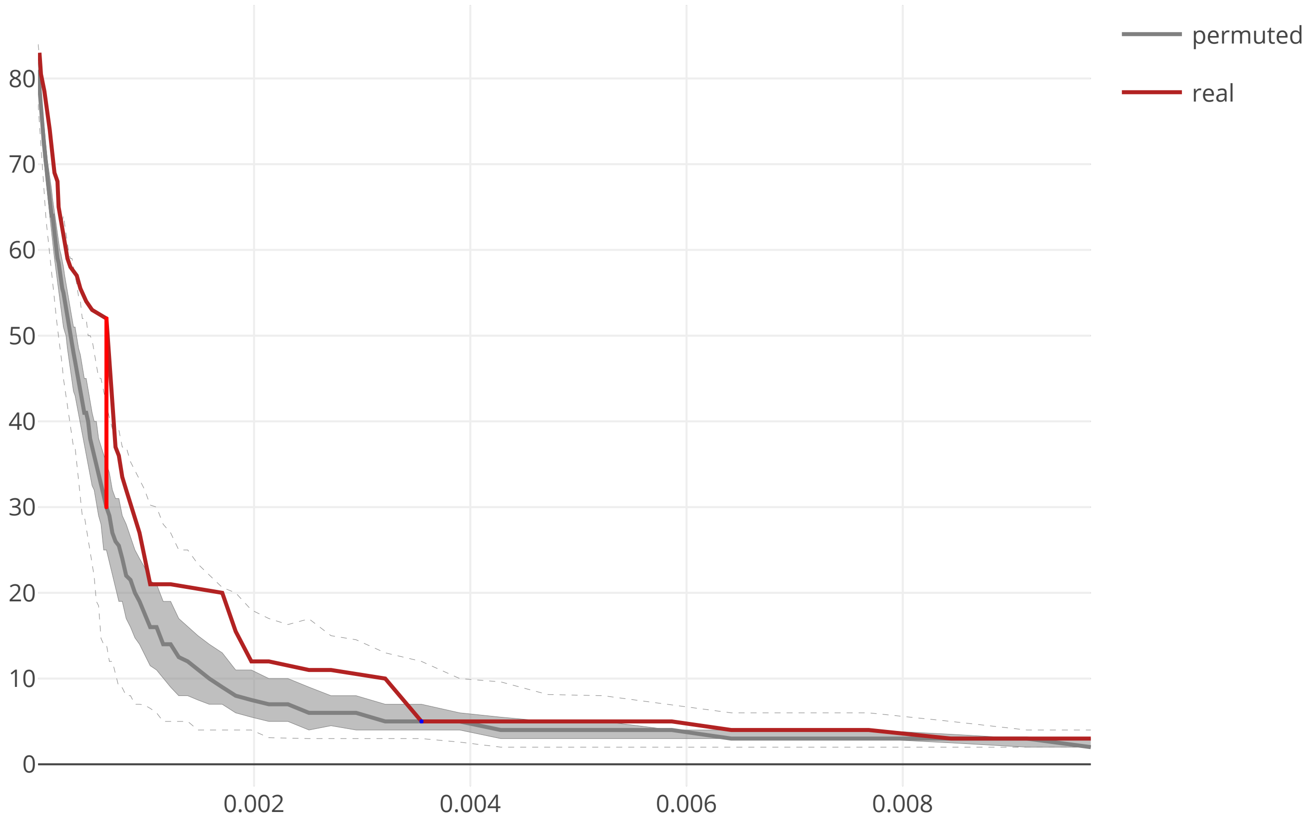

### cov2_SARS_CoV2_NSP4_flow_distance_20200609.pdf

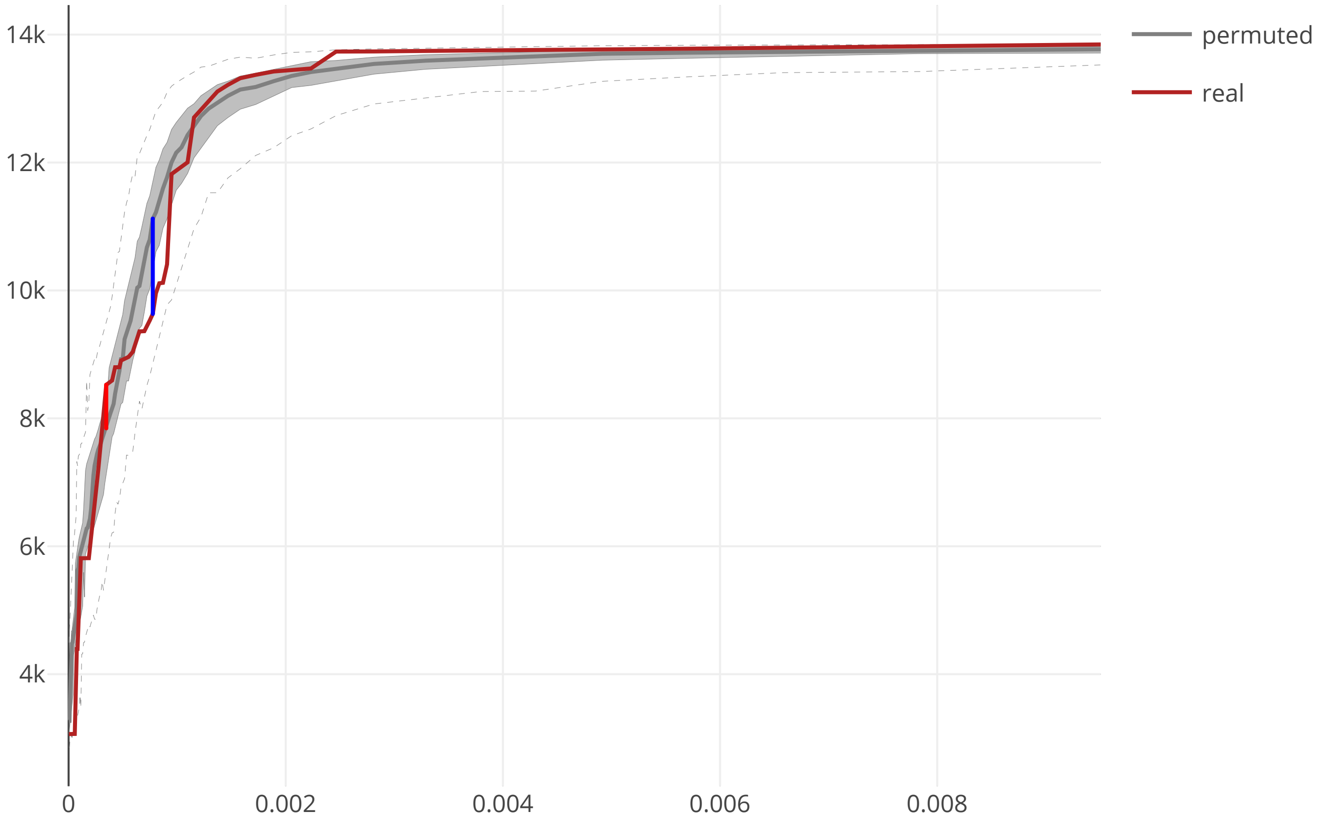

### cov2_SARS_CoV2_NSP4_maxcomponent_size_20200609.pdf

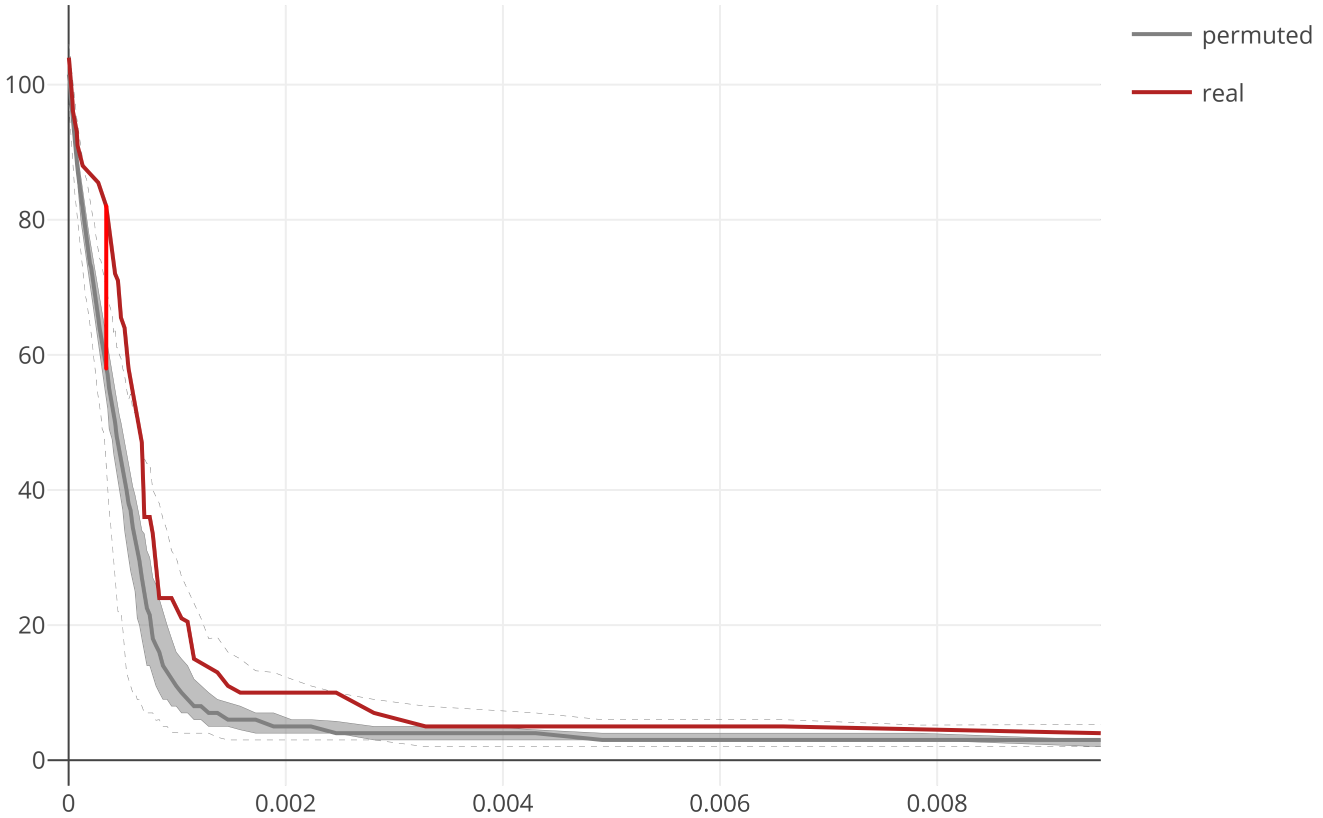

### cov2_SARS_CoV2_NSP6_flow_distance_20200609.pdf

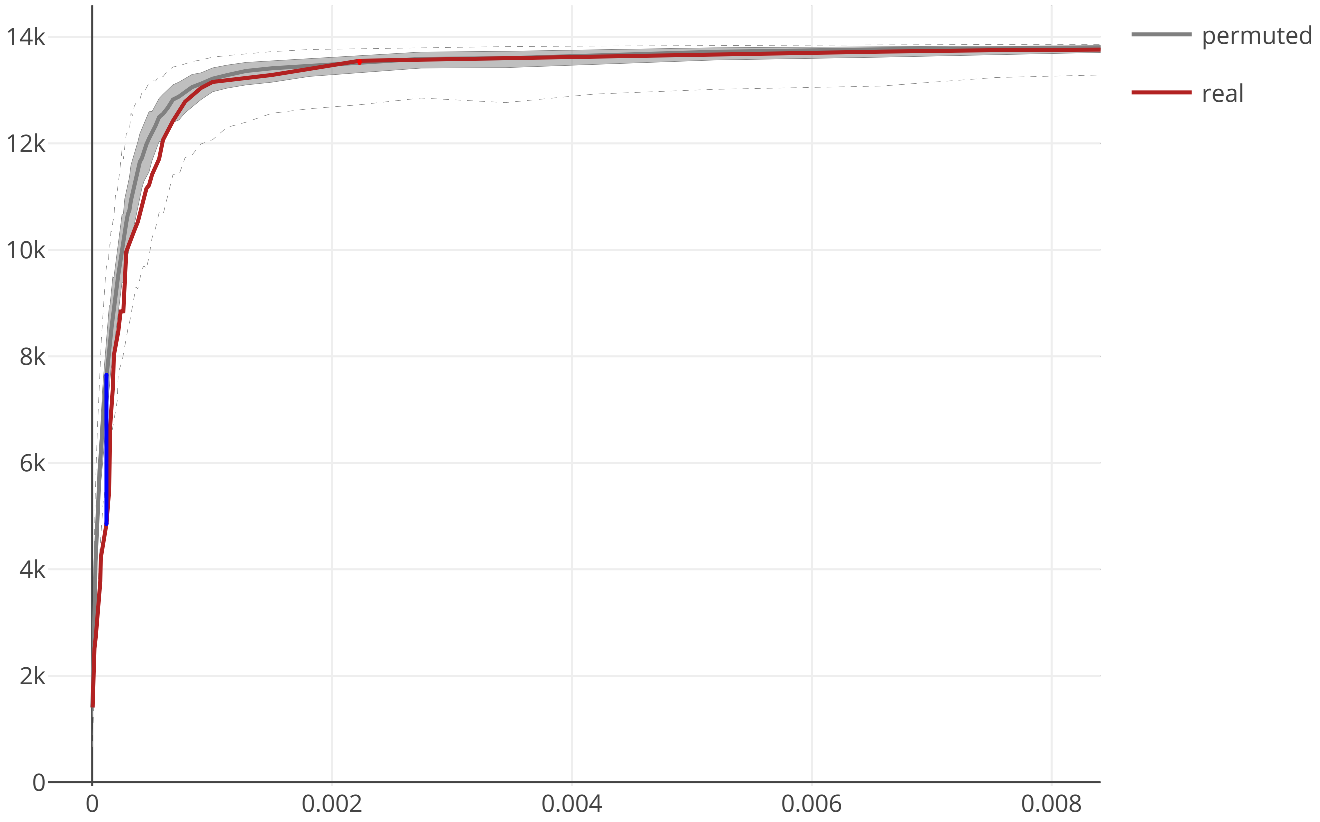

### cov2_SARS_CoV2_NSP6_maxcomponent_size_20200609.pdf

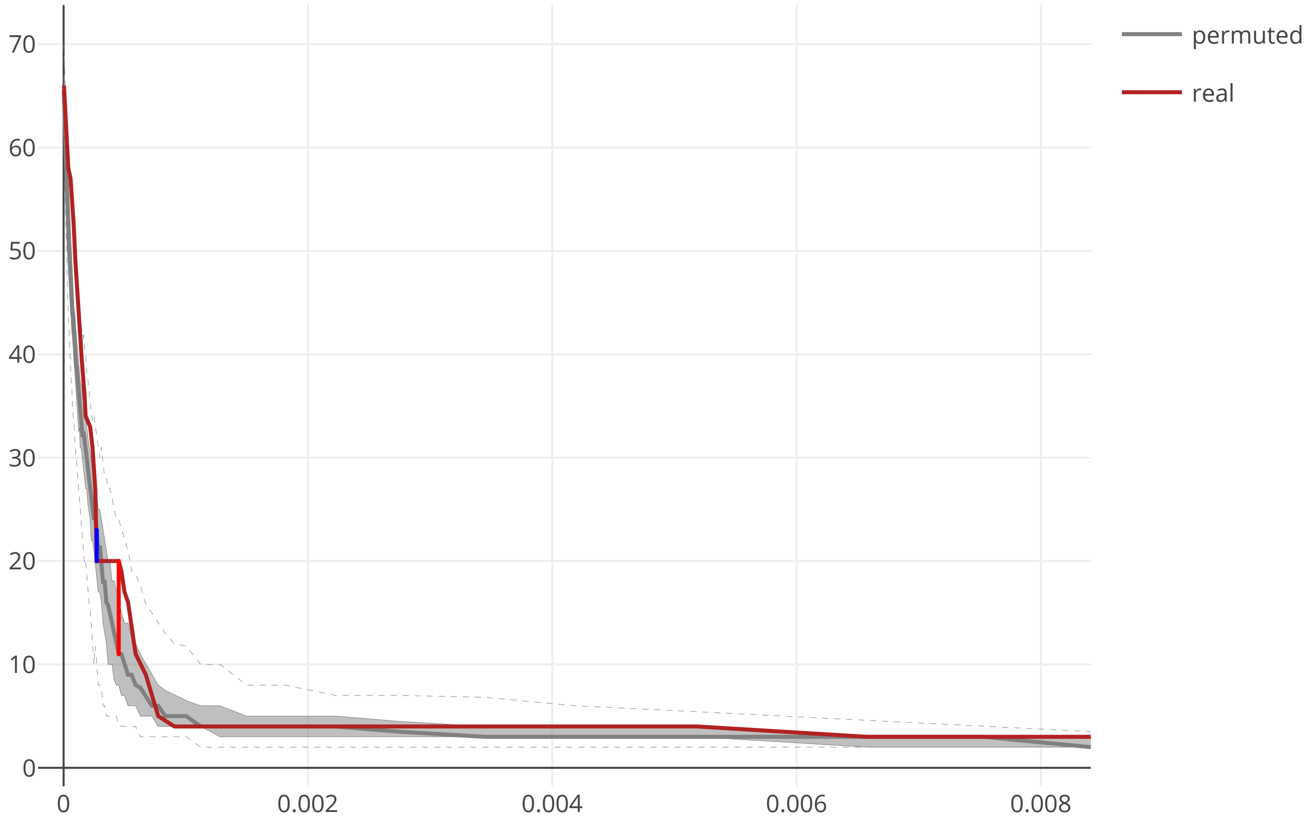

### cov2_SARS_CoV2_NSP7_flow_distance_20200609.pdf

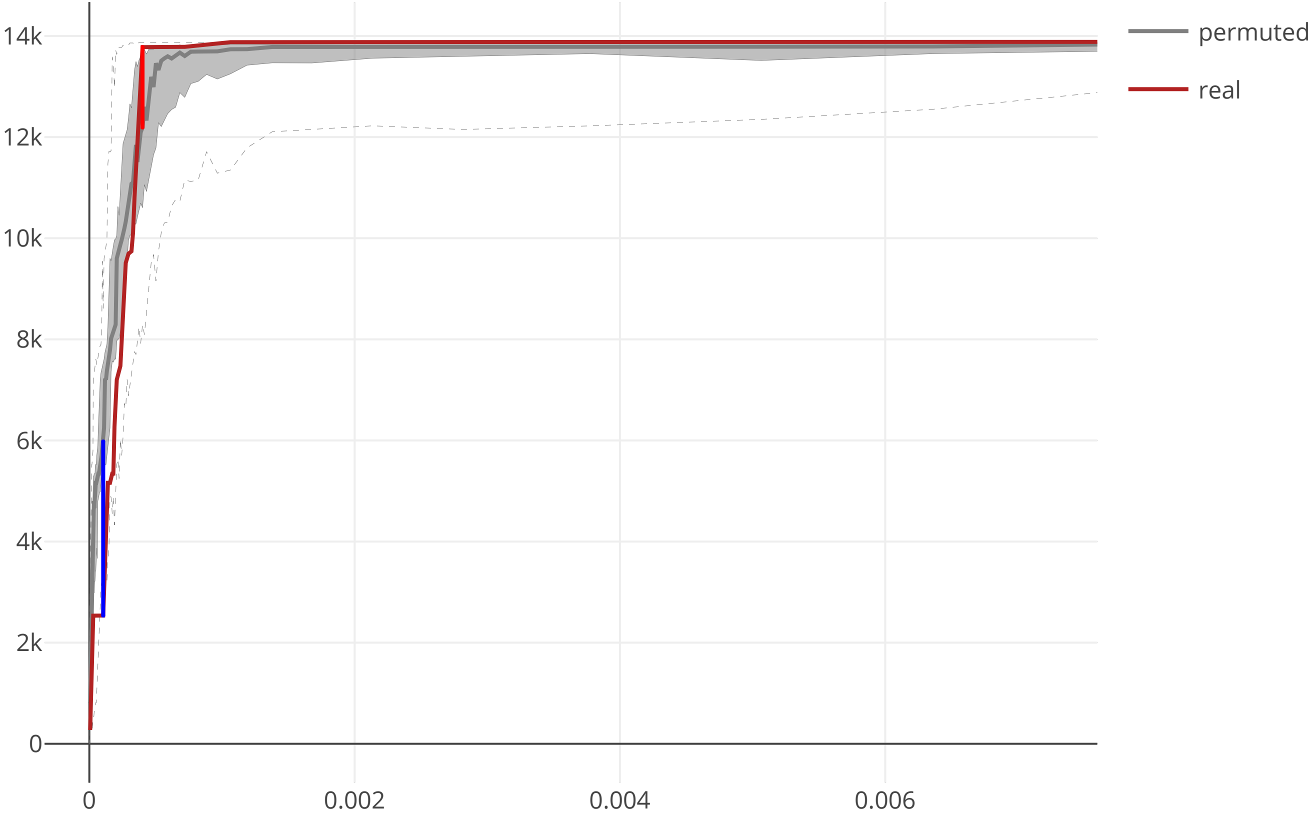

### cov2_SARS_CoV2_NSP7_maxcomponent_size_20200609.pdf

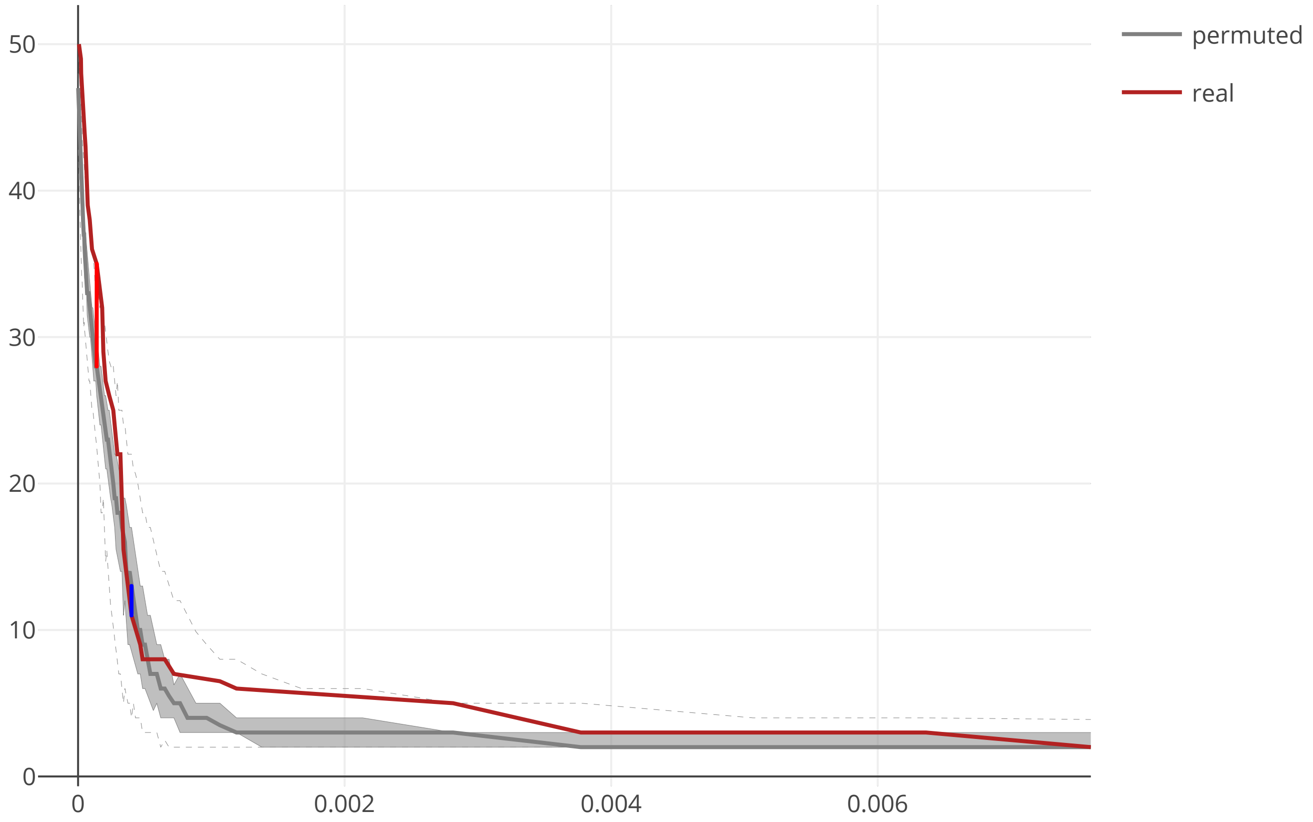

### cov2_SARS_CoV2_NSP8_flow_distance_20200609.pdf

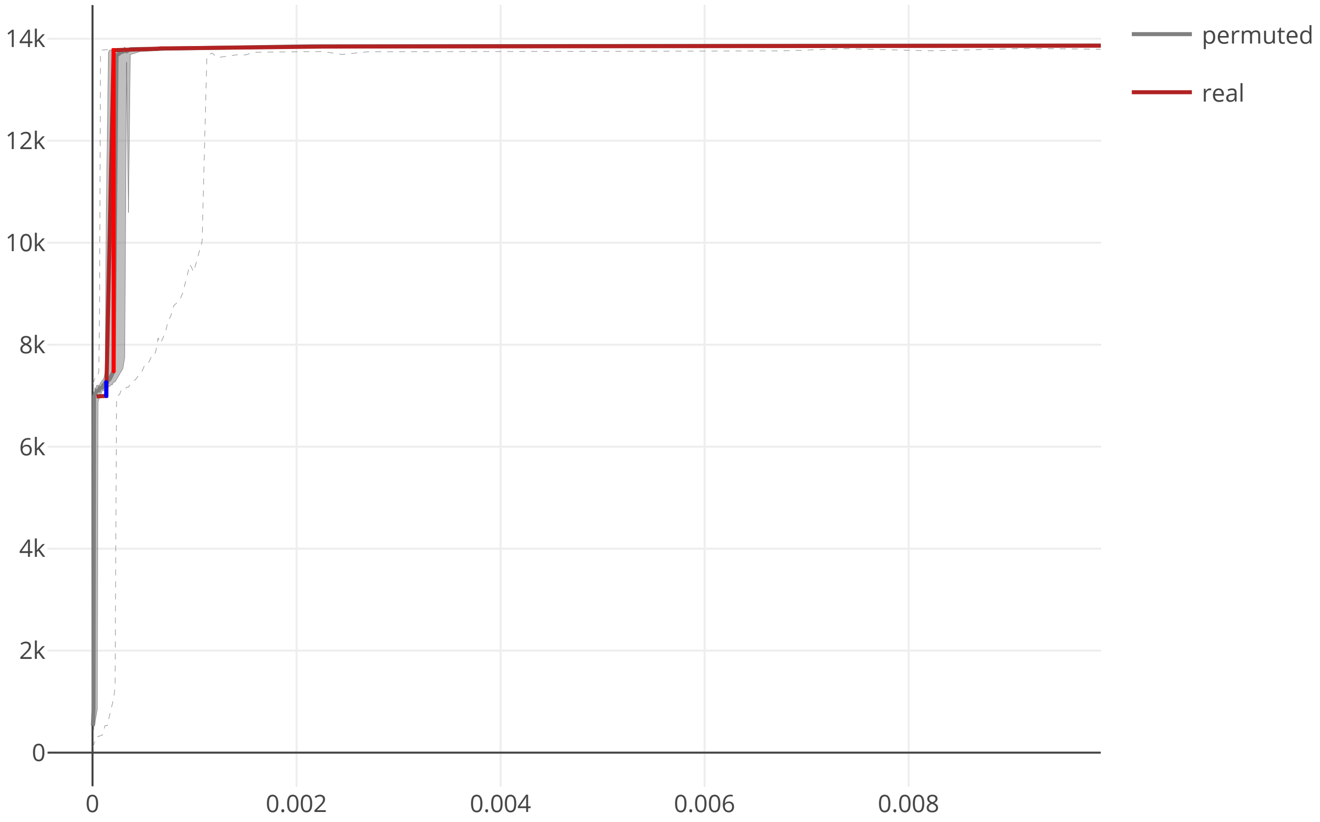

### cov2_SARS_CoV2_NSP8_maxcomponent_size_20200609.pdf

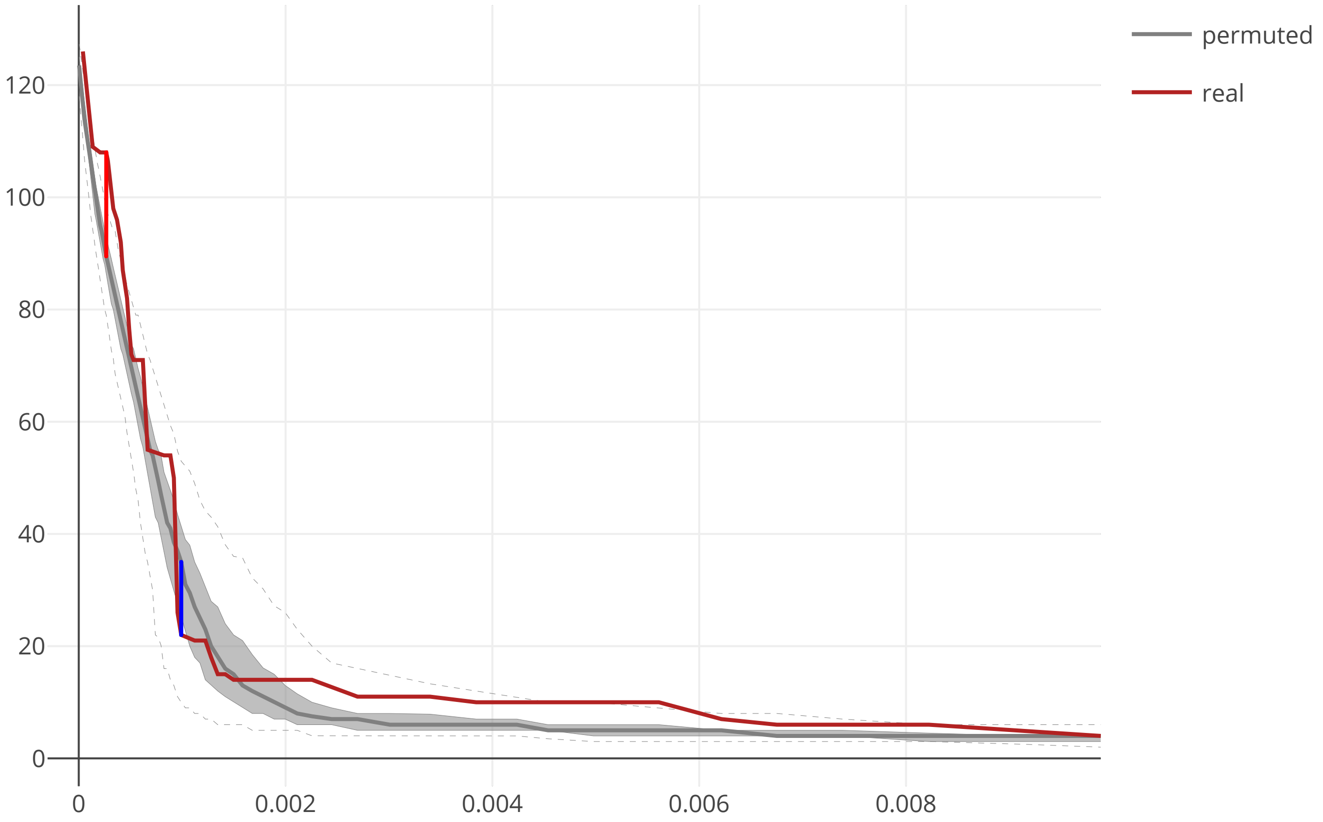

### cov2_SARS_CoV2_NSP9_flow_distance_20200609.pdf

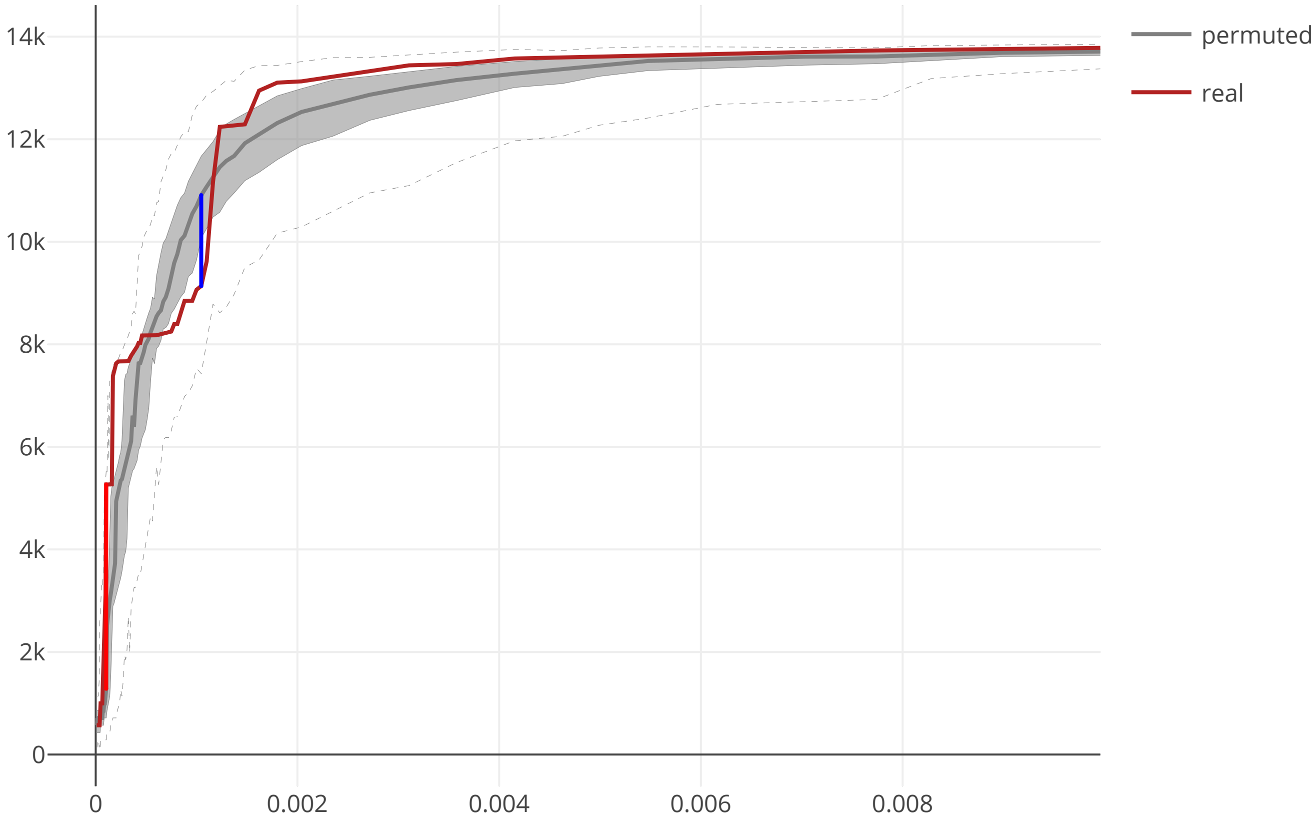

### cov2_SARS_CoV2_NSP9_maxcomponent_size_20200609.pdf

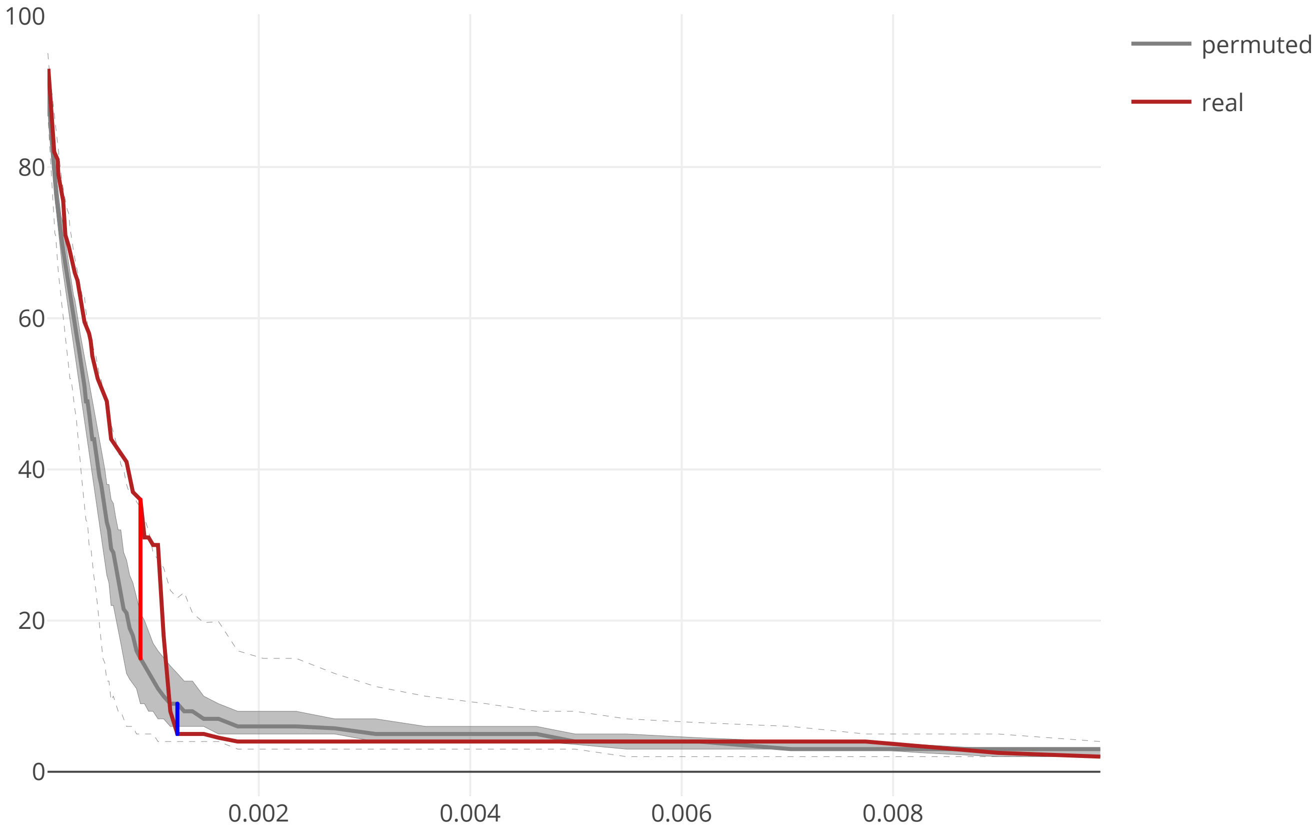

### cov2_SARS_CoV2_NSP10_flow_distance_20200609.pdf

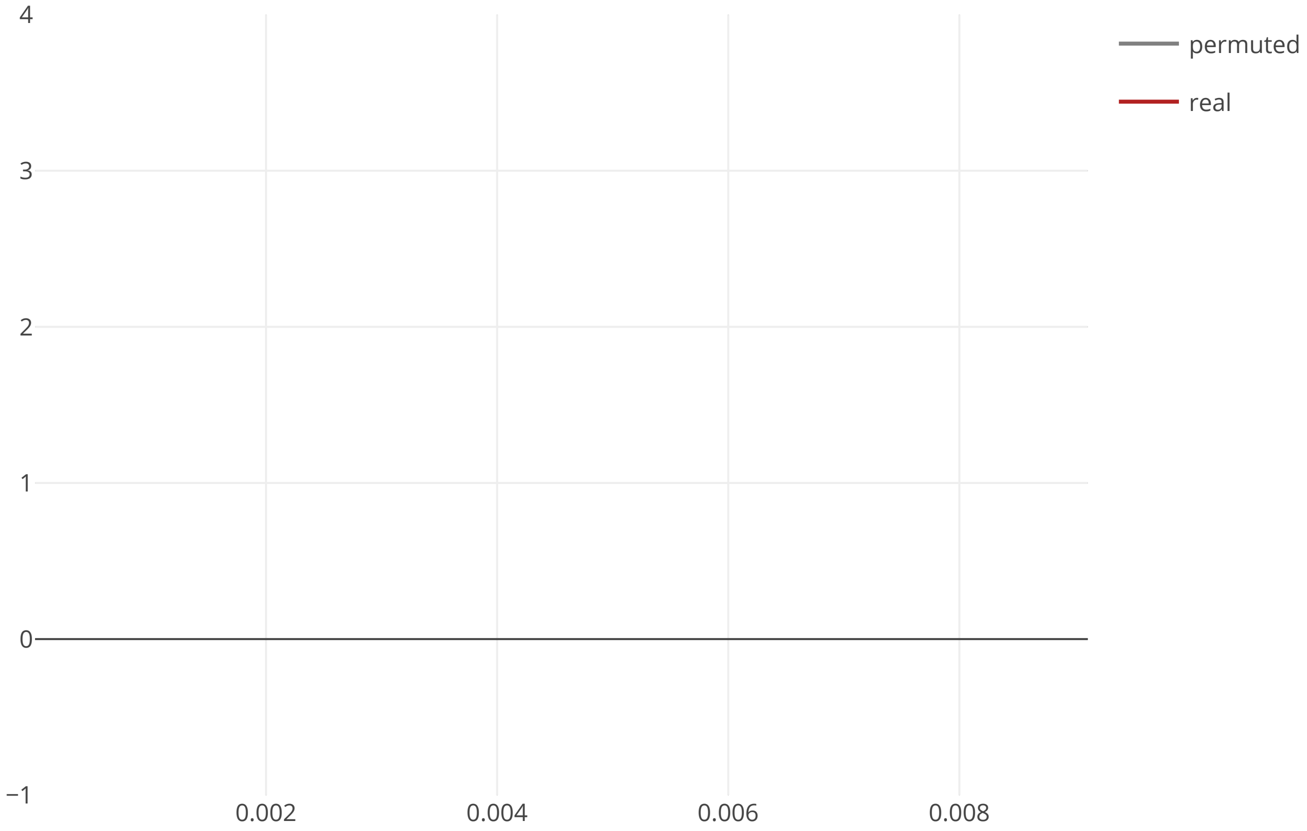

### cov2_SARS_CoV2_NSP10_maxcomponent_size_20200609.pdf

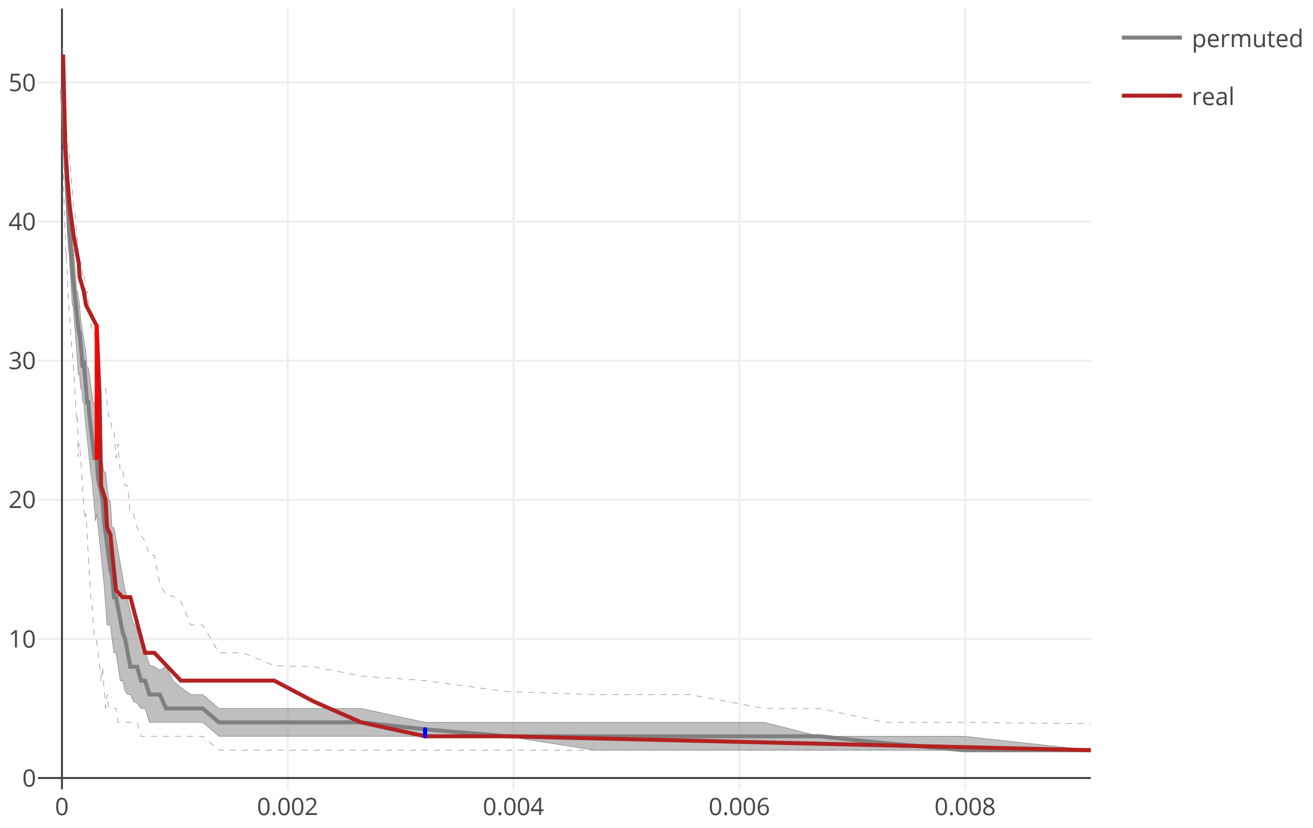

### cov2_SARS_CoV2_NSP12_flow_distance_20200609.pdf

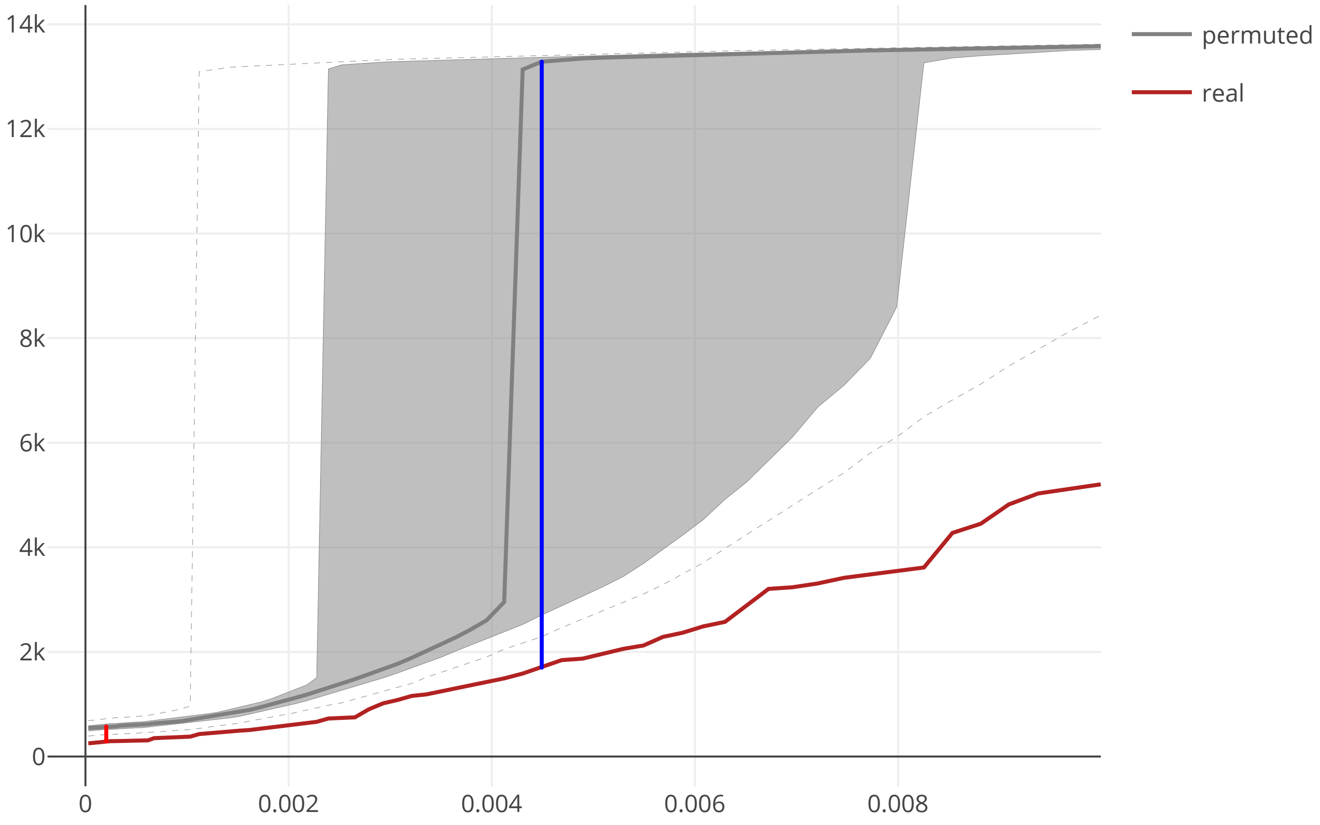

### cov2_SARS_CoV2_NSP12_maxcomponent_size_20200609.pdf

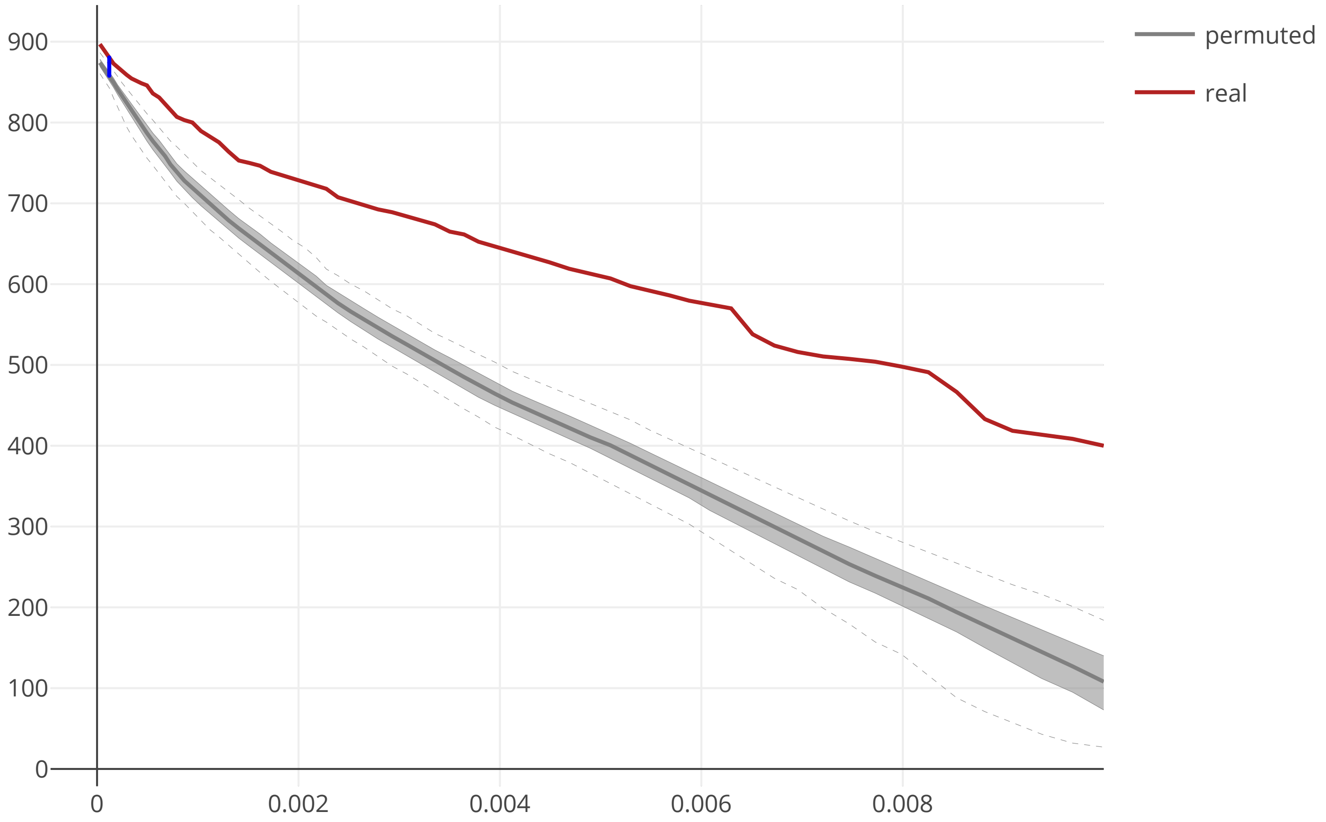

### cov2_SARS_CoV2_NSP13_flow_distance_20200609.pdf

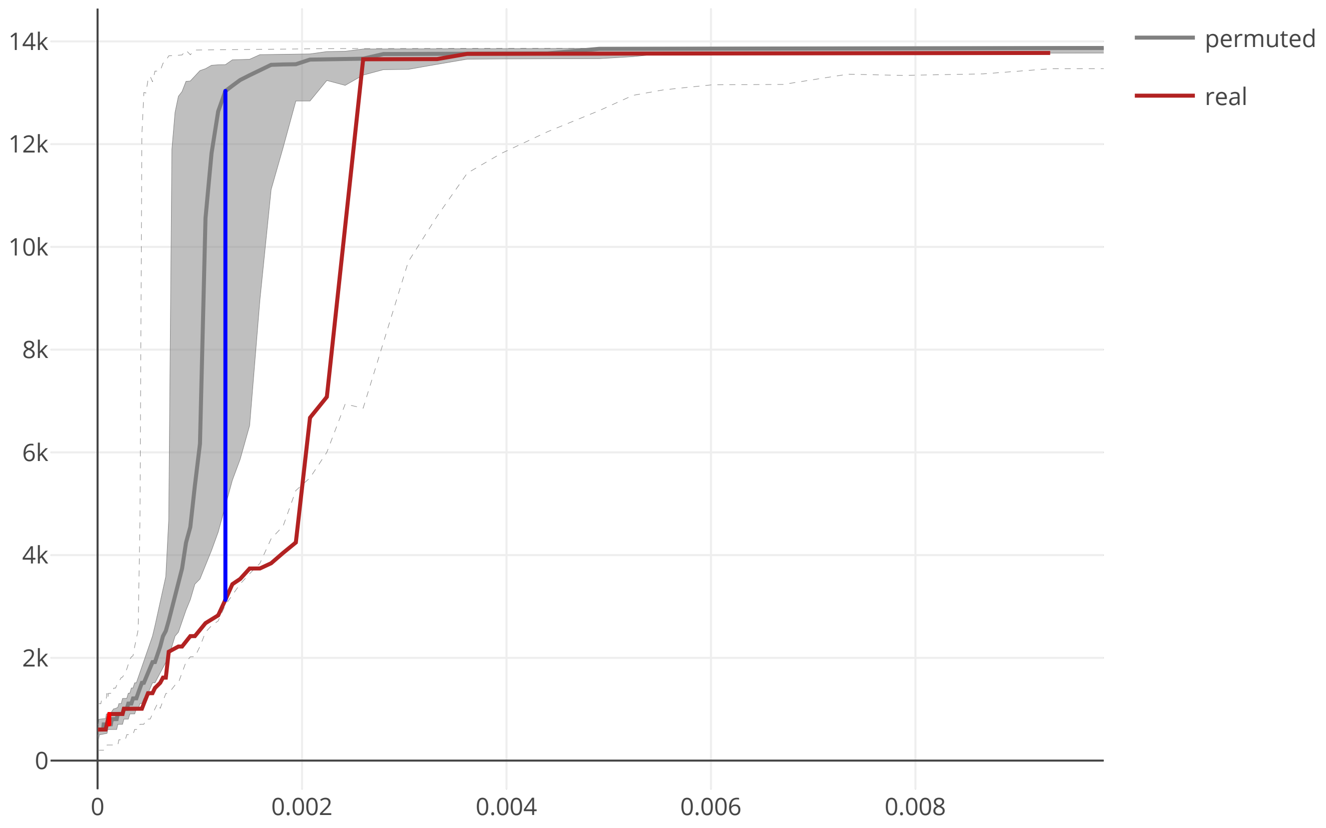

### cov2_SARS_CoV2_NSP13_maxcomponent_size_20200609.pdf

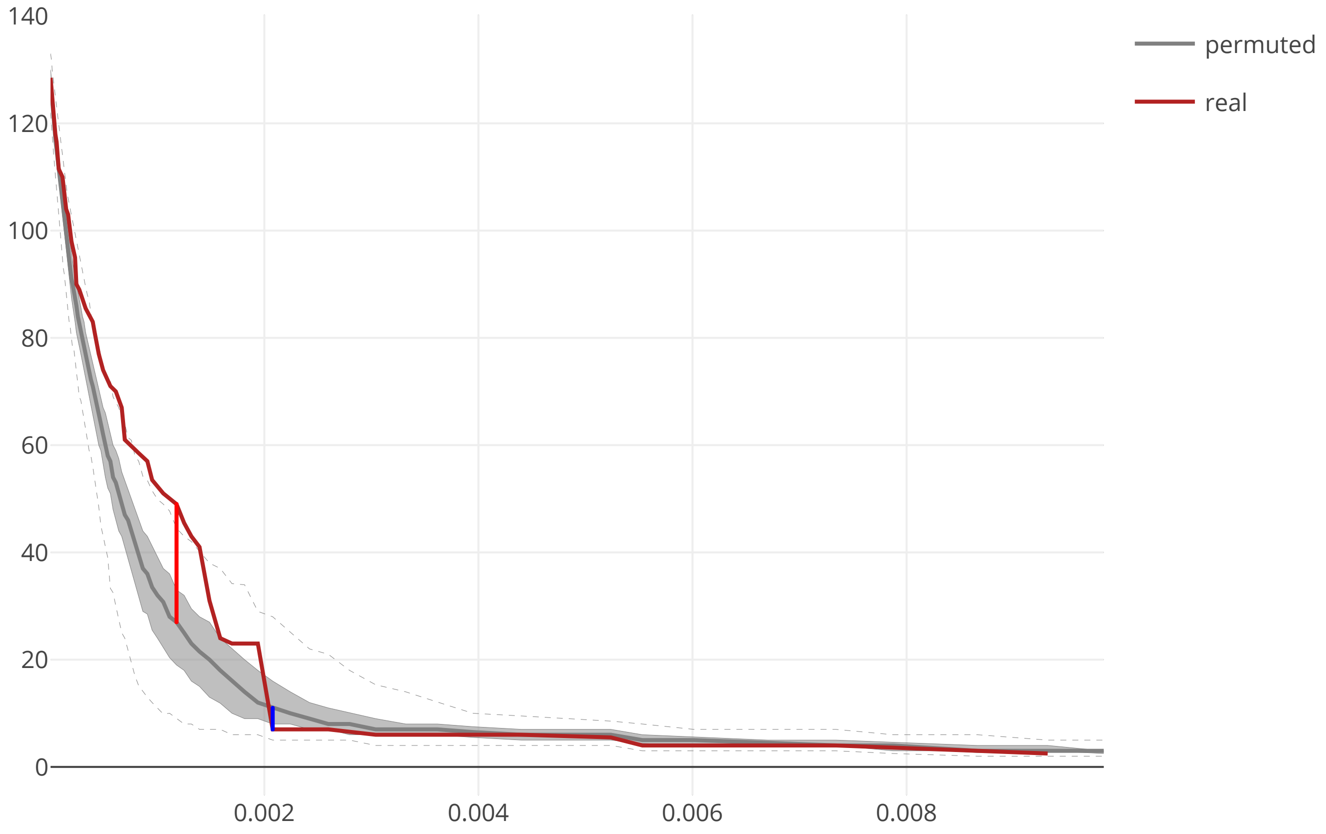

### cov2_SARS_CoV2_NSP14_flow_distance_20200609.pdf

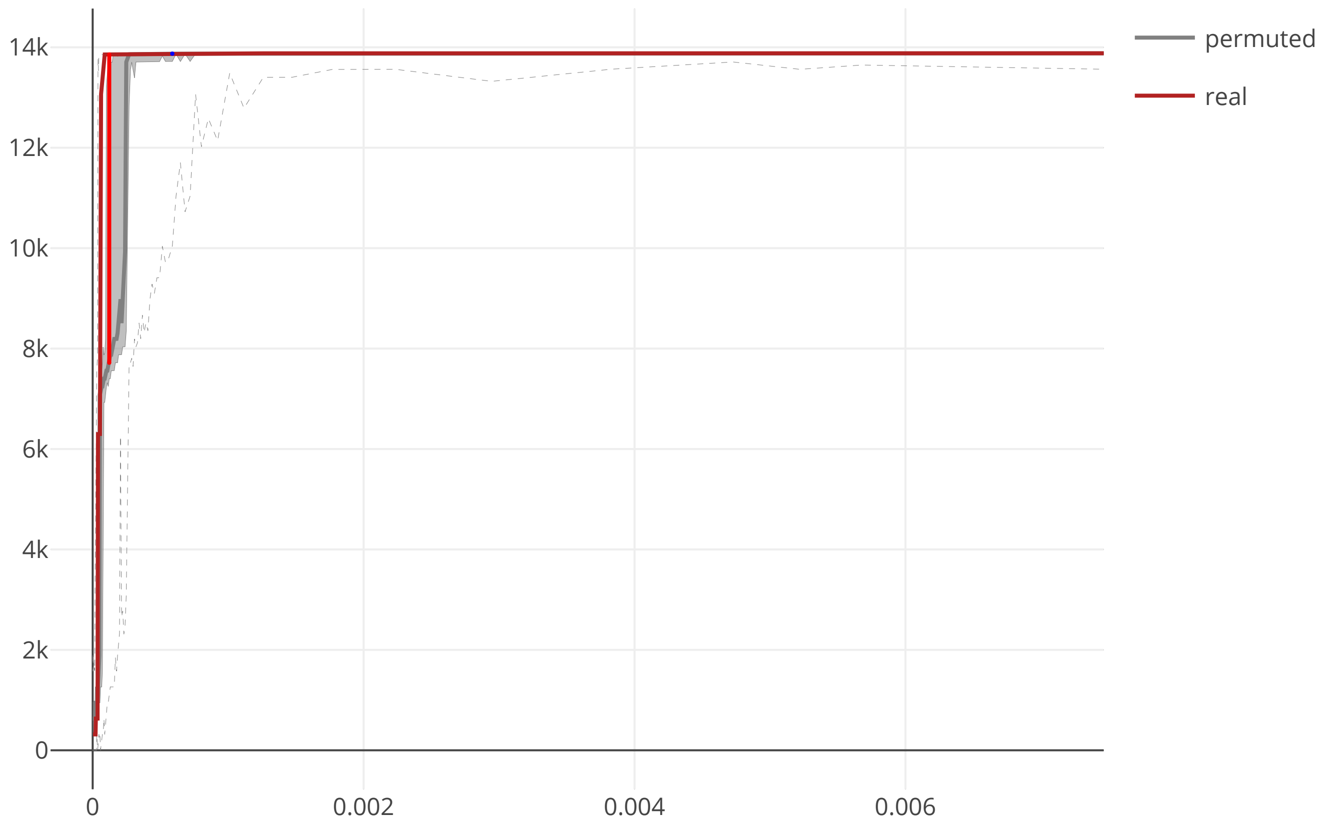

### cov2_SARS_CoV2_NSP14_maxcomponent_size_20200609.pdf

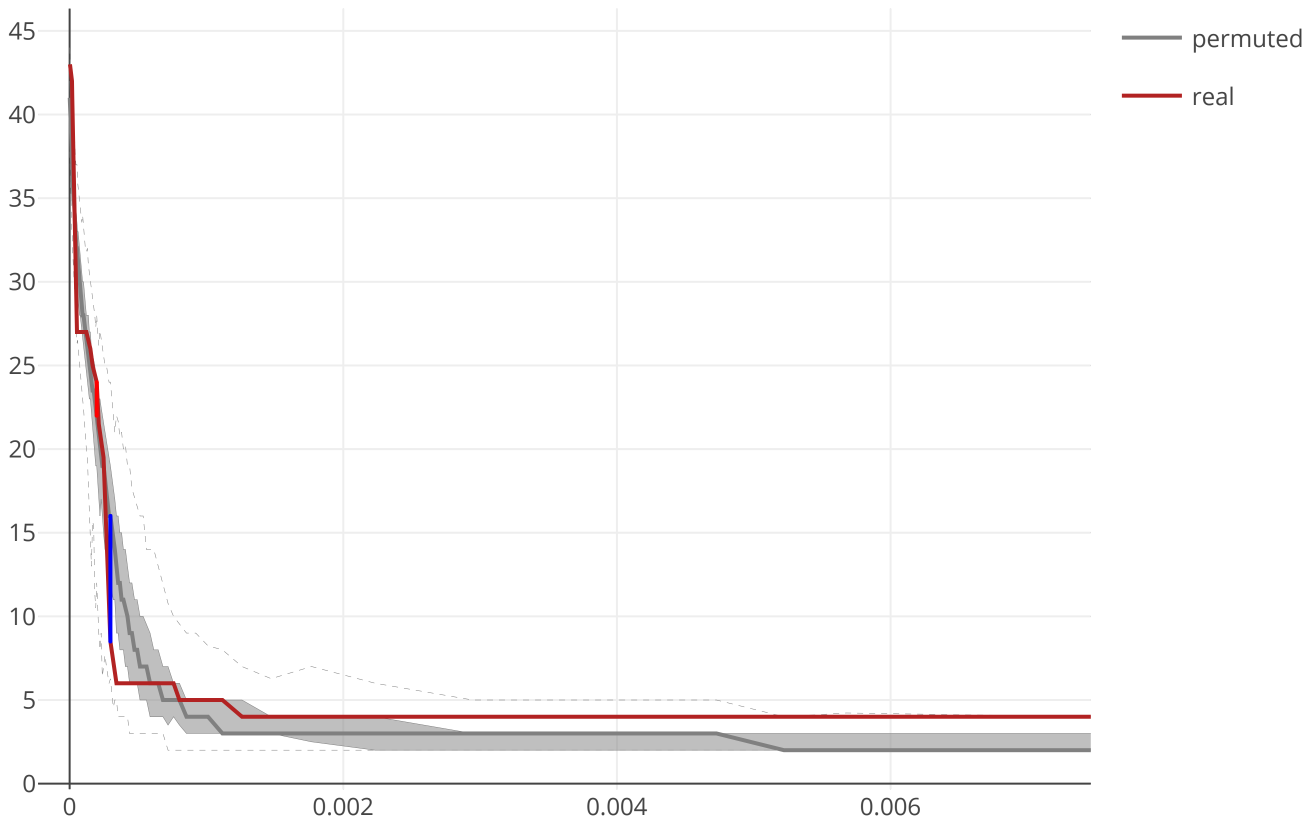

### cov2_SARS_CoV2_NSP15_flow_distance_20200609.pdf

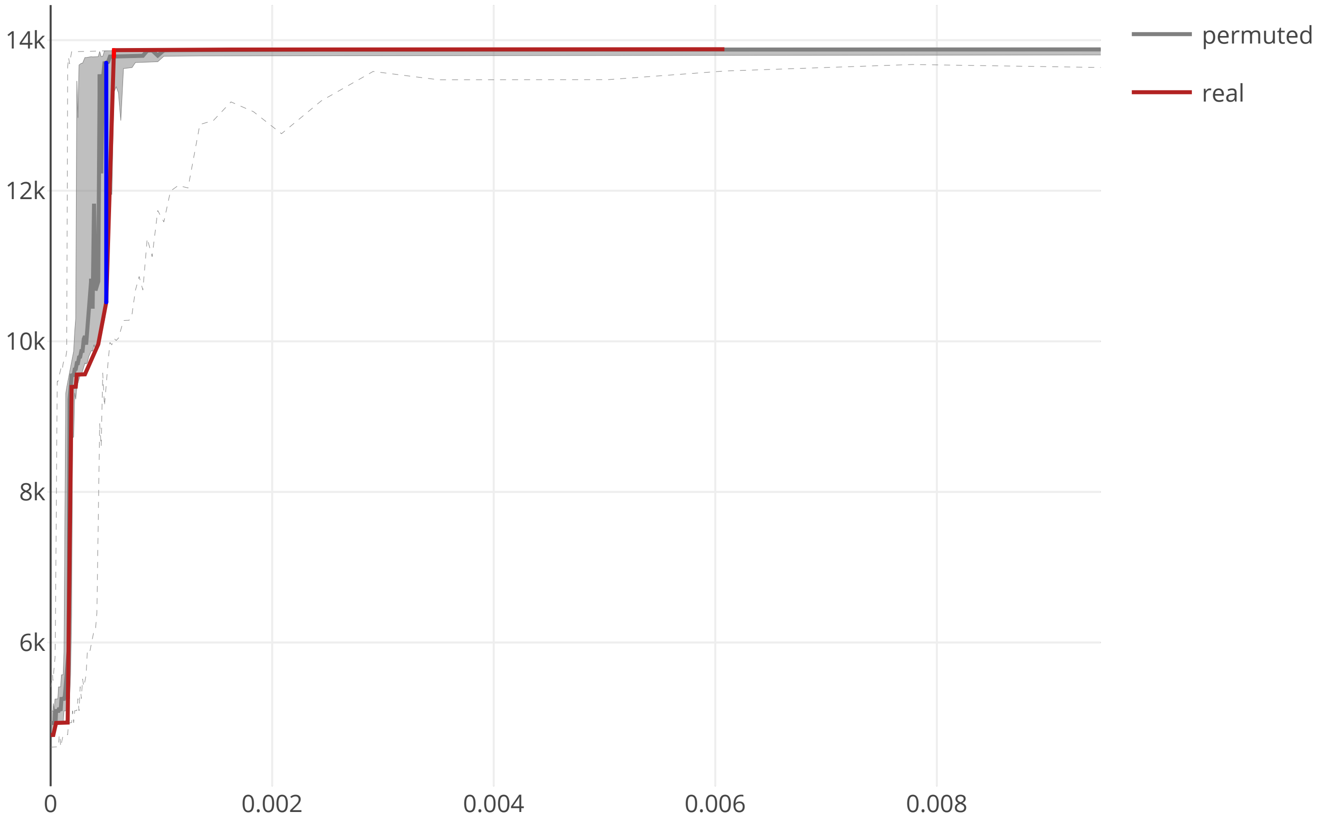

### cov2_SARS_CoV2_NSP15_maxcomponent_size_20200609.pdf

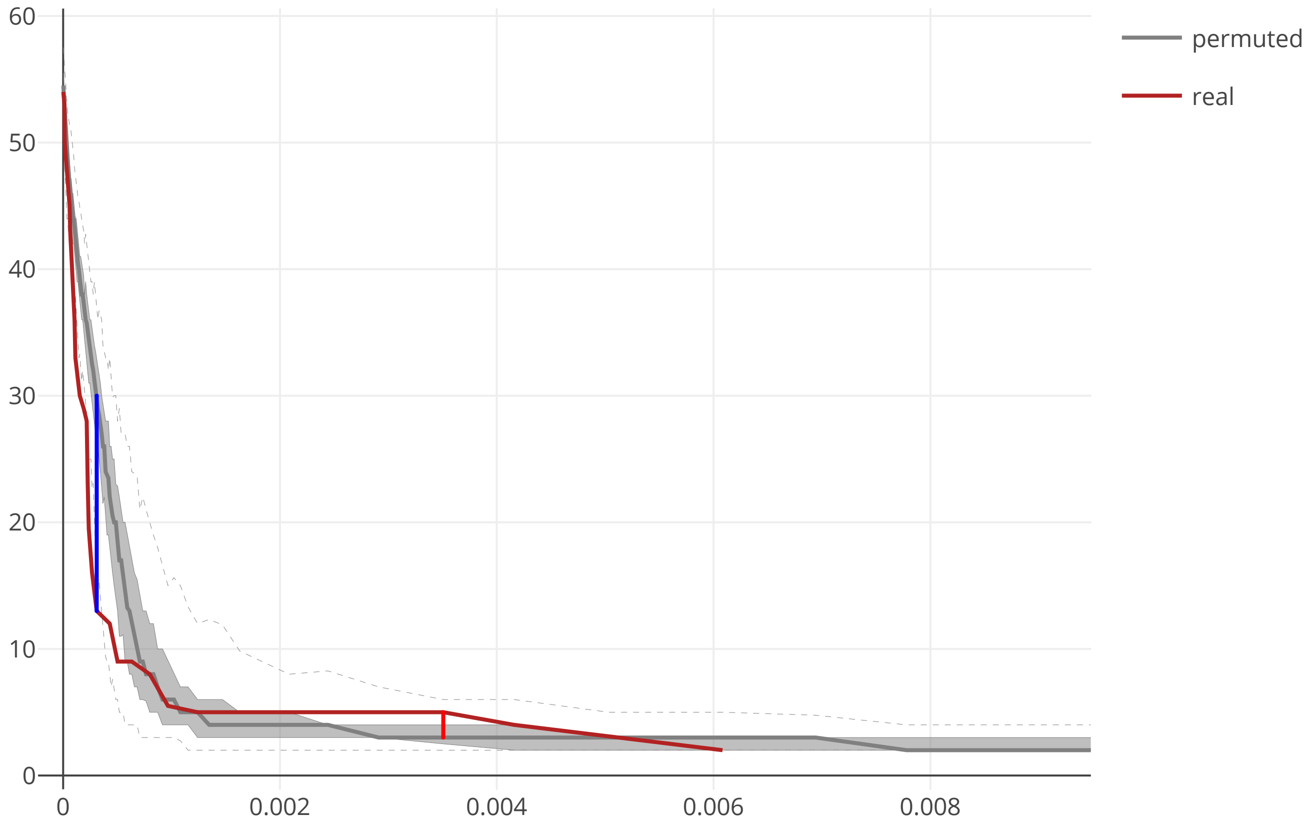

### cov2_SARS_CoV2_NSP16_flow_distance_20200609.pdf

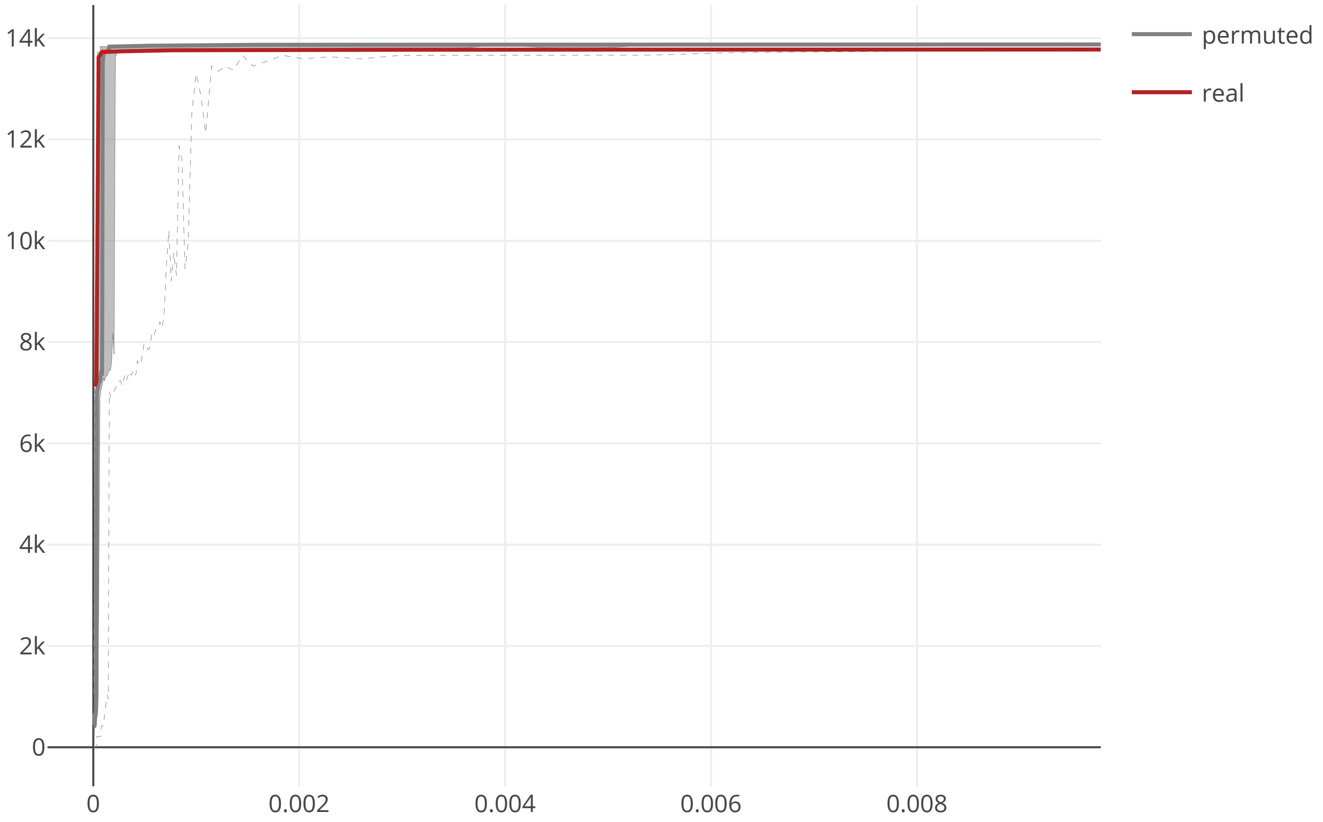

### cov2_SARS_CoV2_NSP16_maxcomponent_size_20200609.pdf

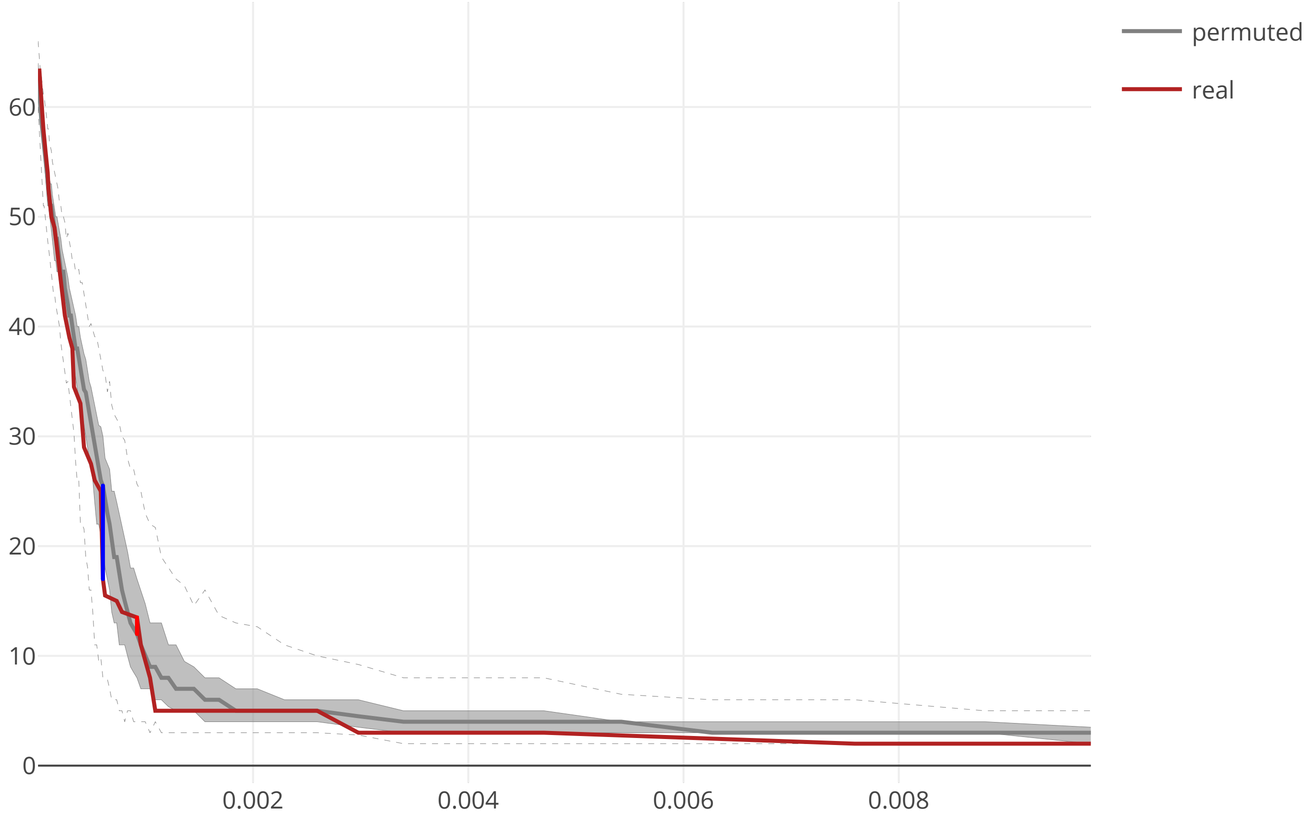

### cov2_SARS_CoV2_ORF3_flow_distance_20200609.pdf

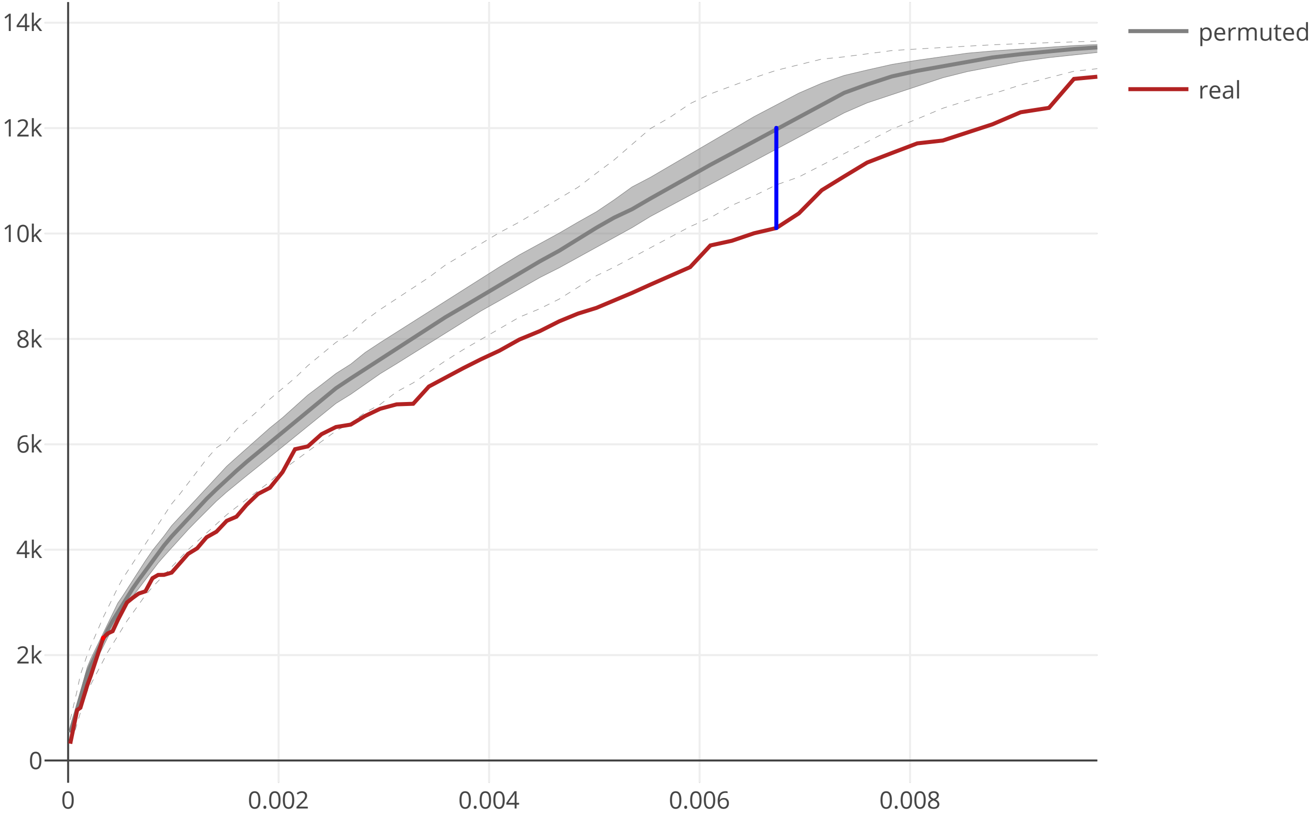

### cov2_SARS_CoV2_ORF3_maxcomponent_size_20200609.pdf

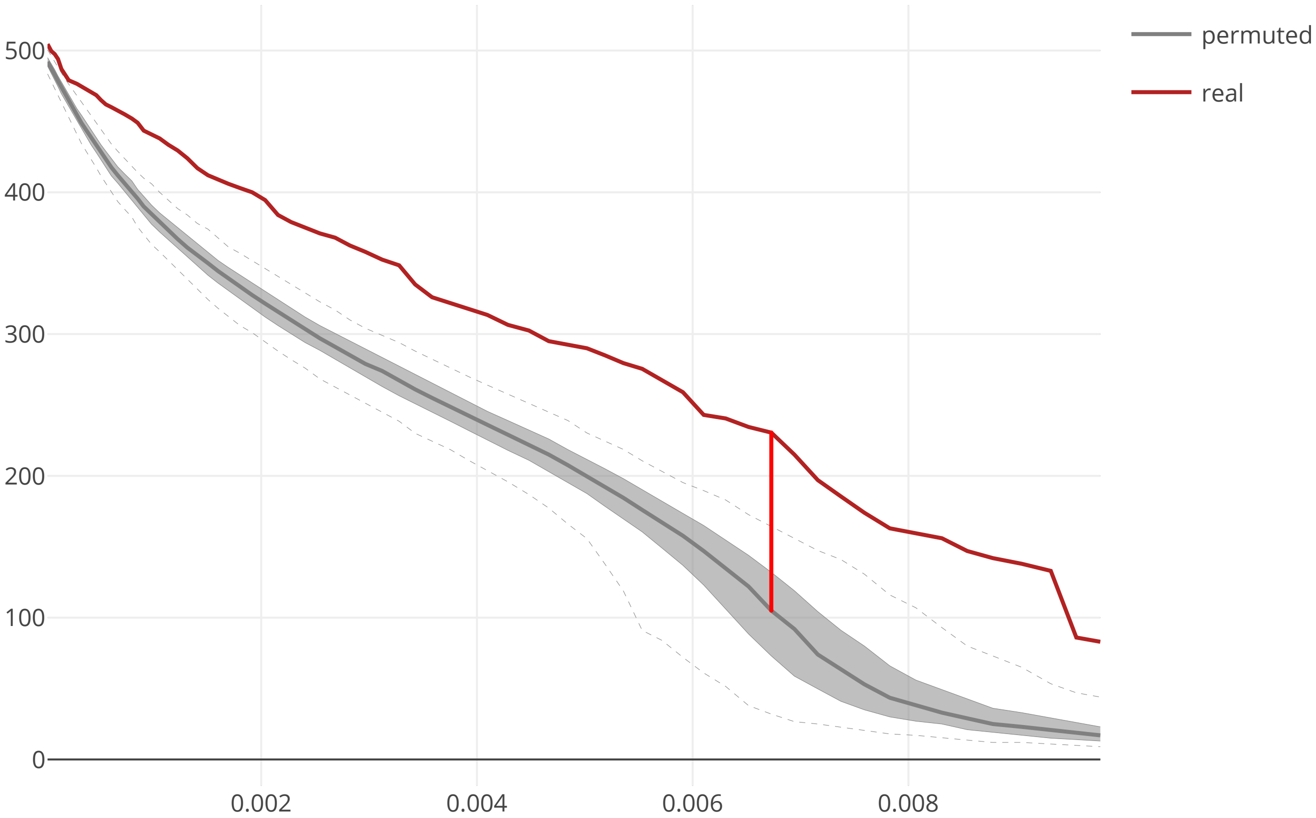

### cov2_SARS_CoV2_ORF6_flow_distance_20200609.pdf

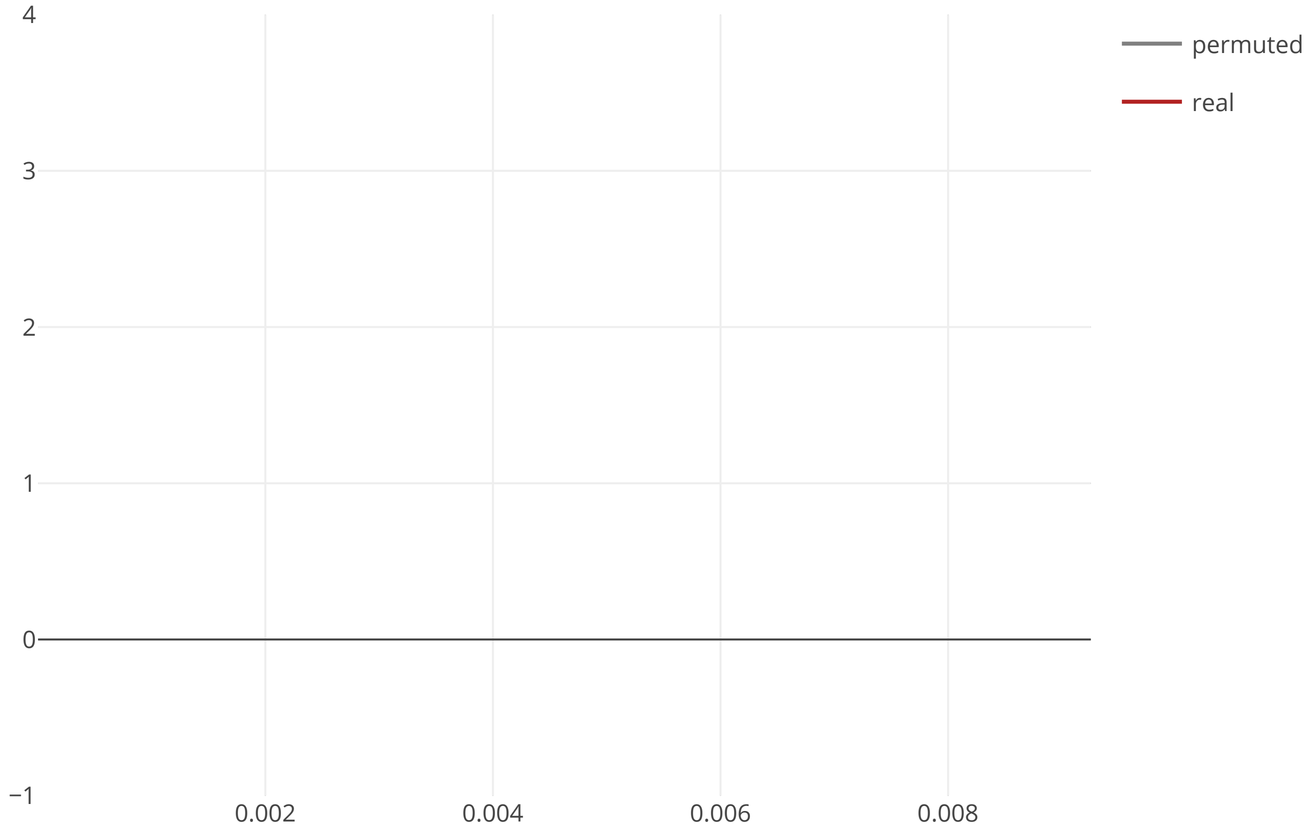

### cov2_SARS_CoV2_ORF6_maxcomponent_size_20200609.pdf

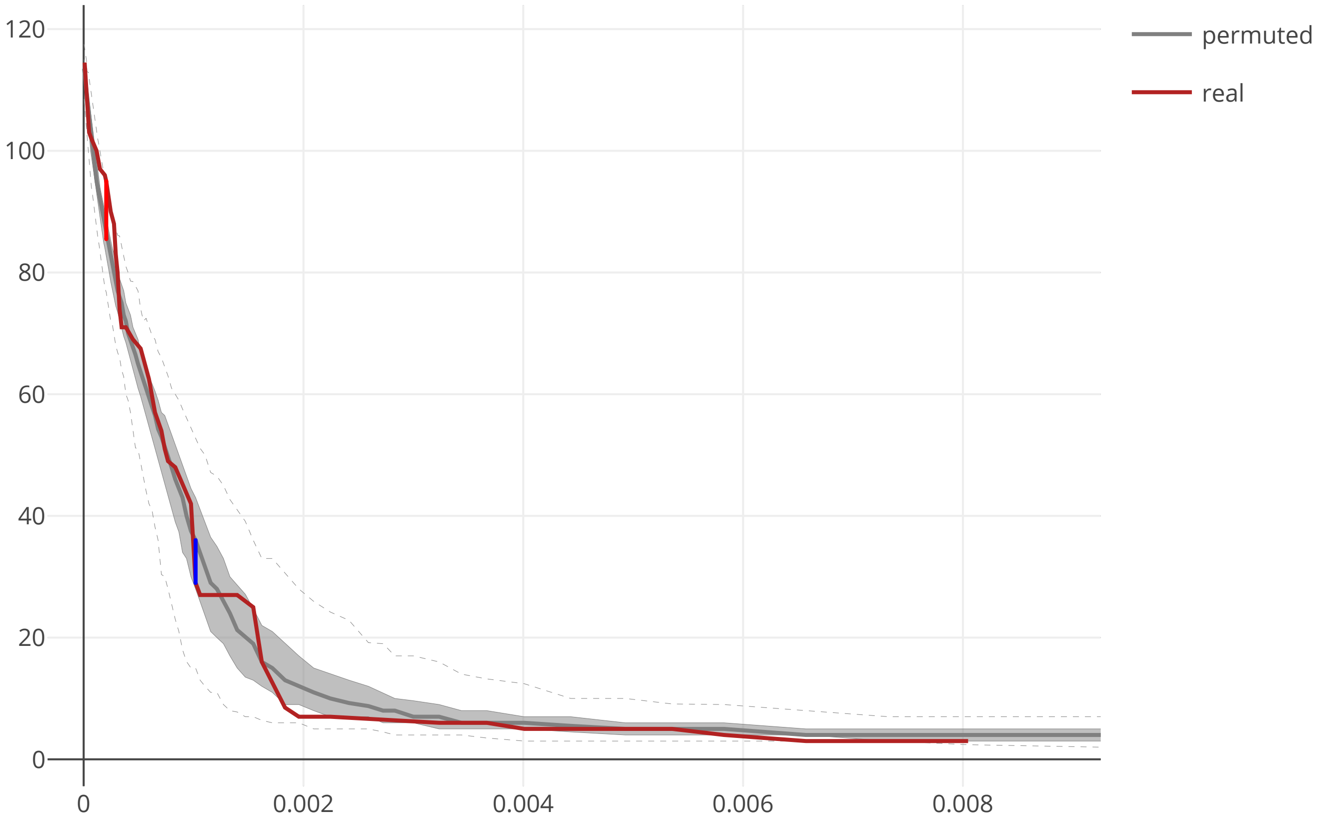

### cov2_SARS_CoV2_ORF7a_flow_distance_20200609.pdf

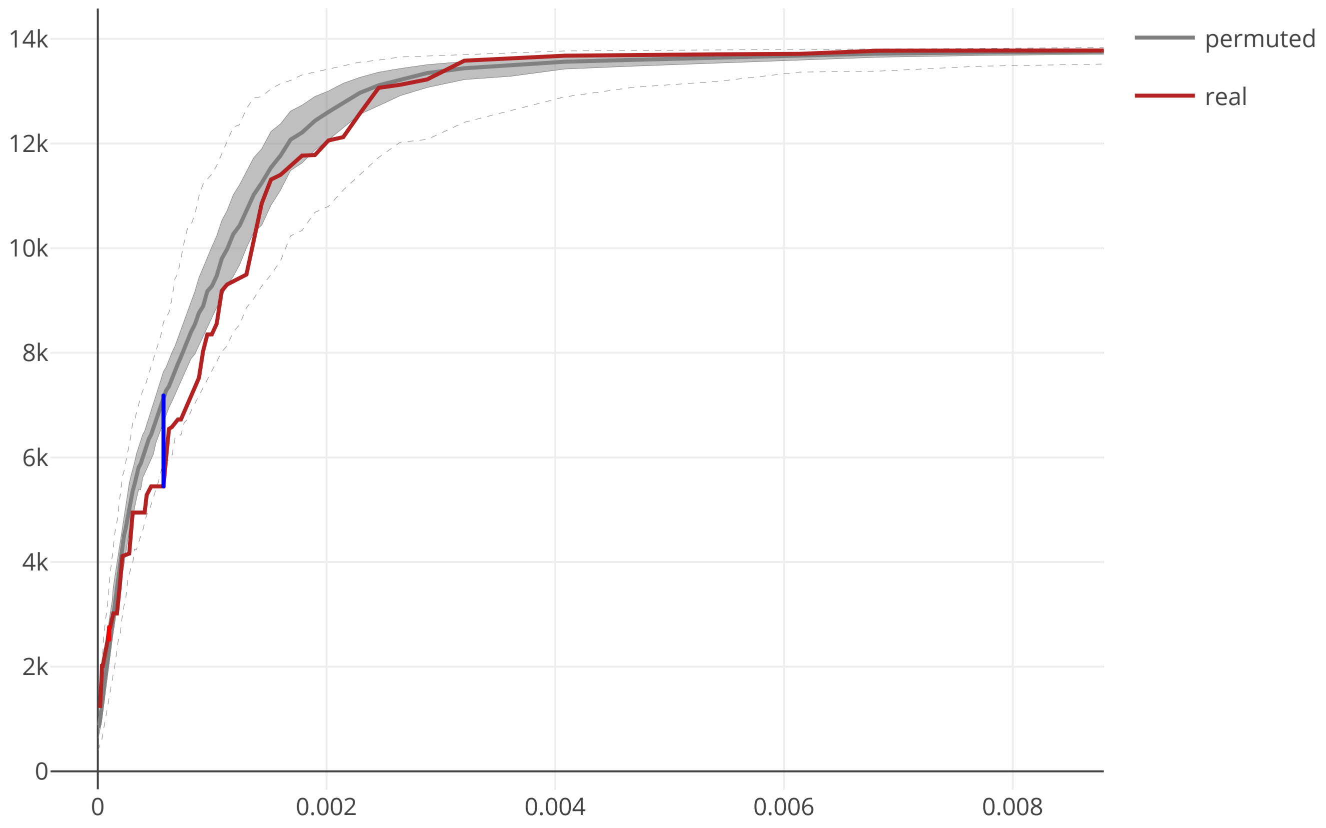

### cov2_SARS_CoV2_ORF7a_maxcomponent_size_20200609.pdf

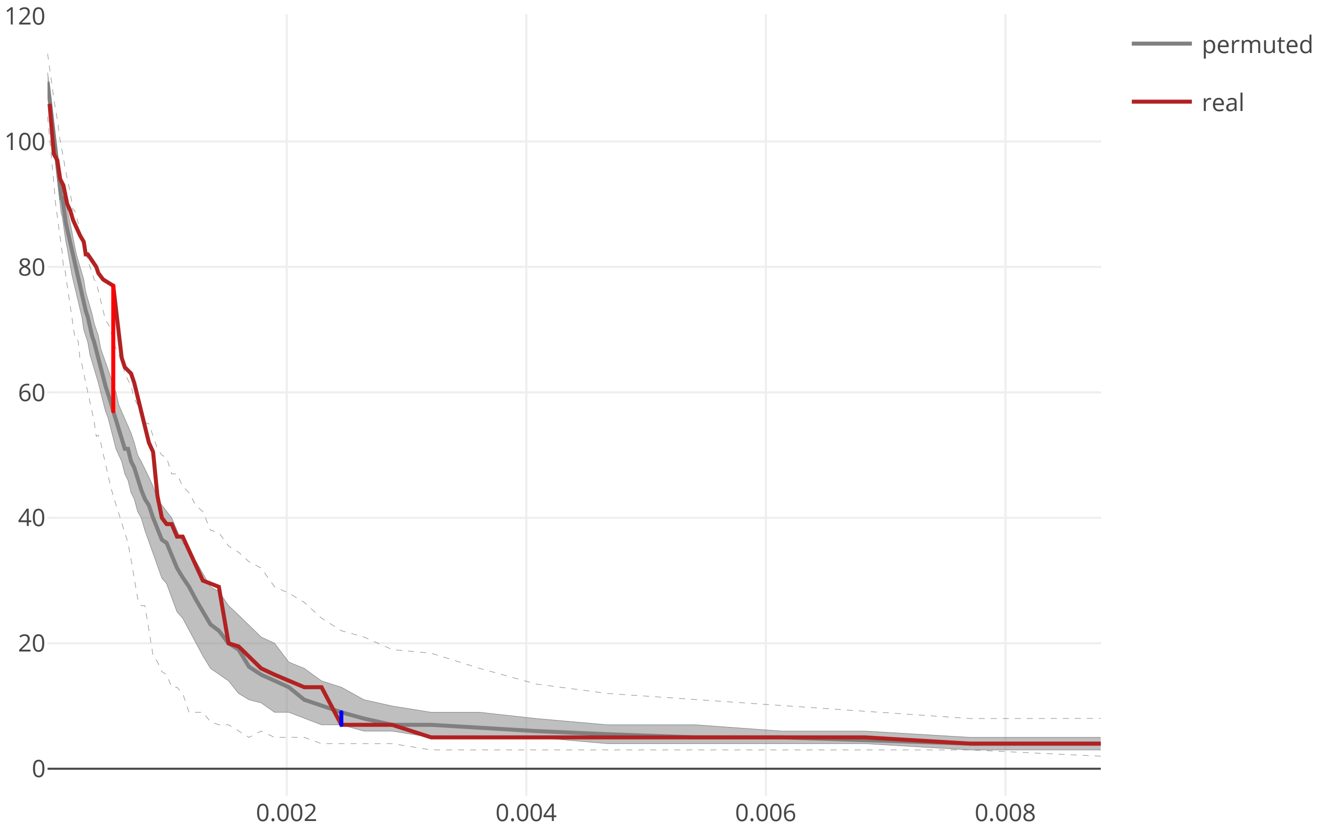

### cov2_SARS_CoV2_ORF7b_flow_distance_20200609.pdf

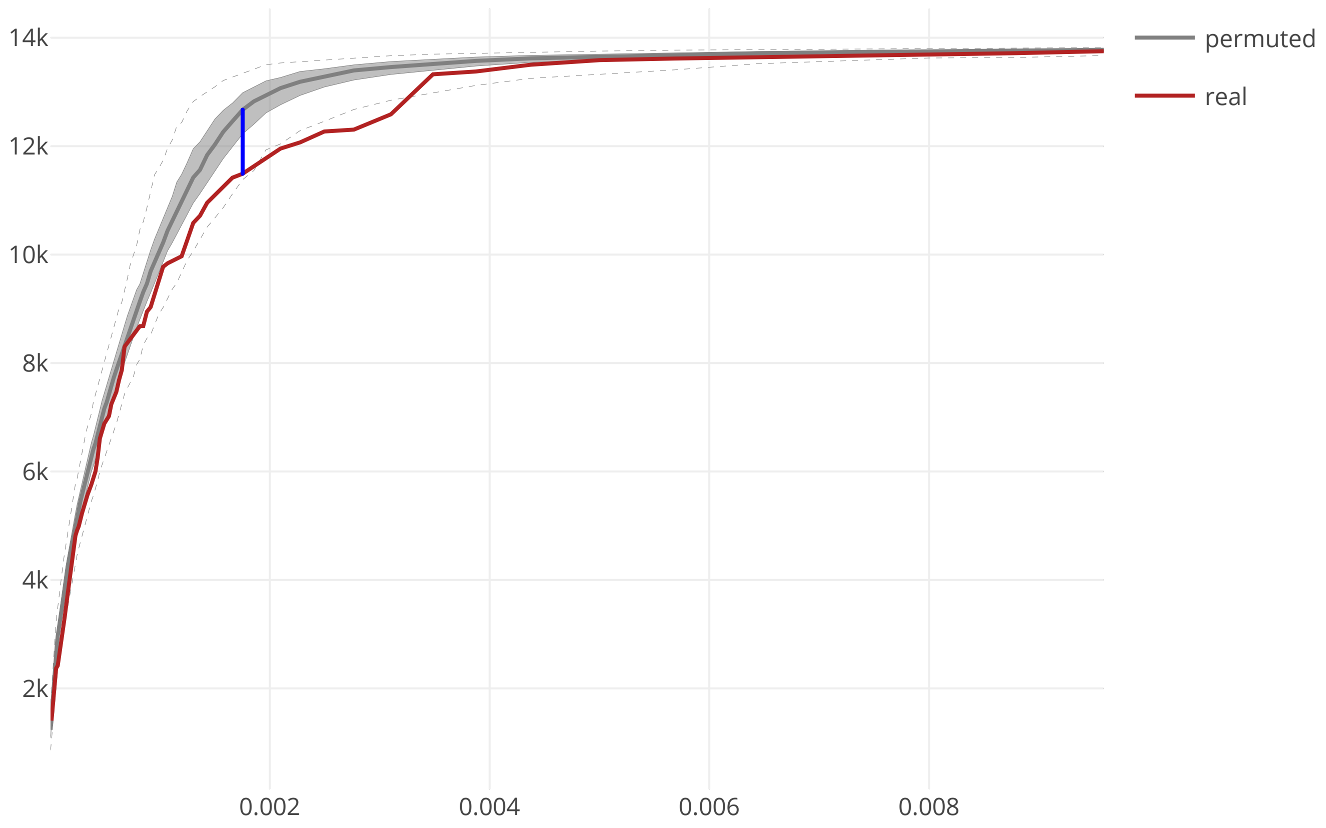

### cov2_SARS_CoV2_ORF7b_maxcomponent_size_20200609.pdf

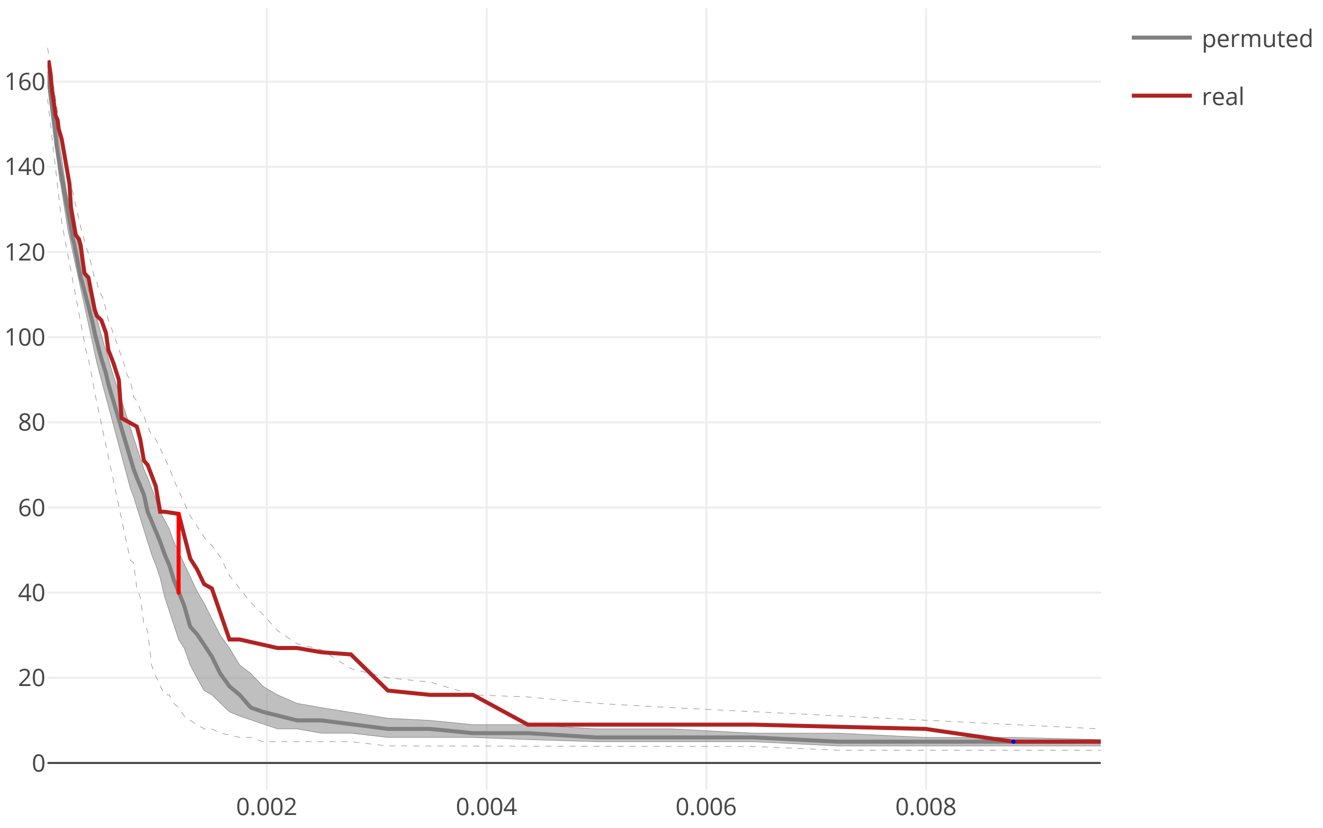

### cov2_SARS_CoV2_ORF8_flow_distance_20200609.pdf

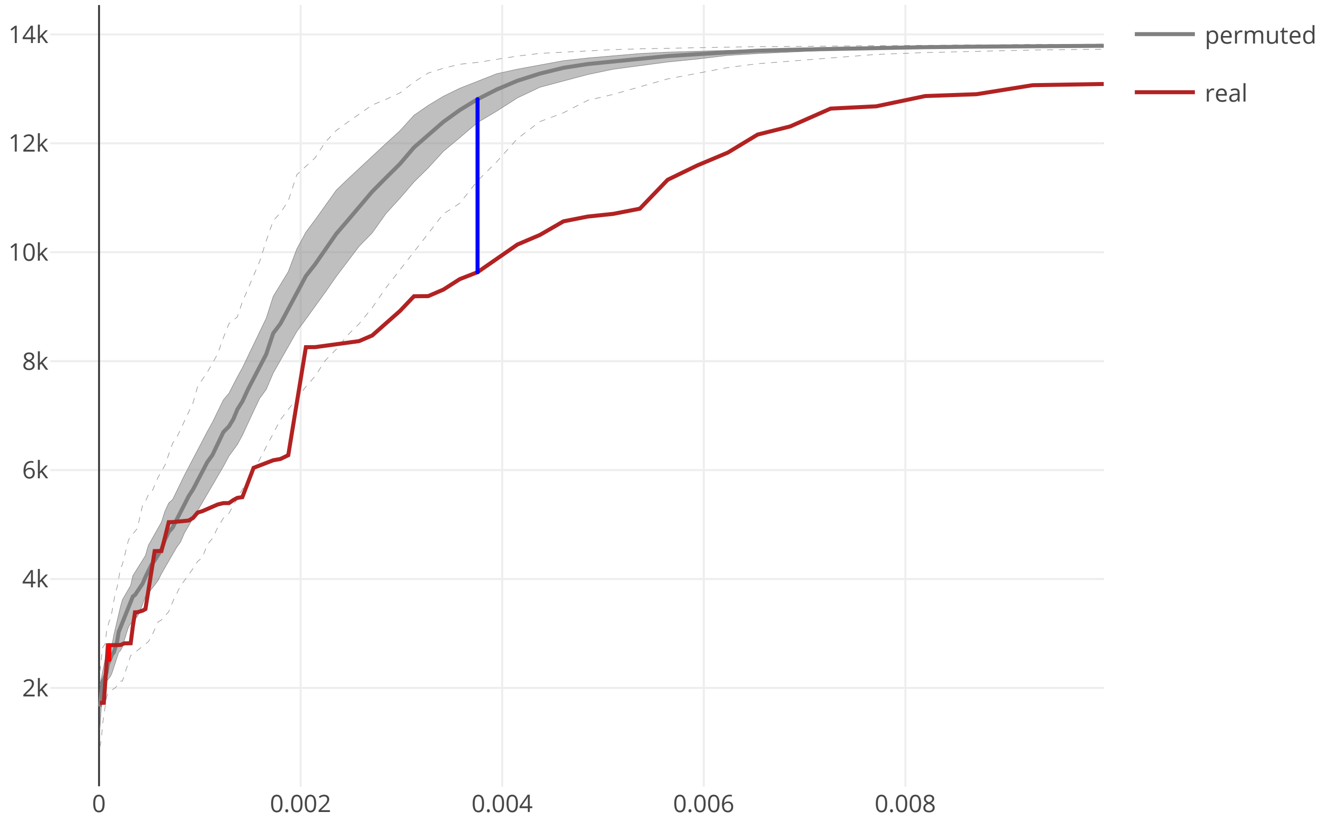

### cov2_SARS_CoV2_ORF8_maxcomponent_size_20200609.pdf

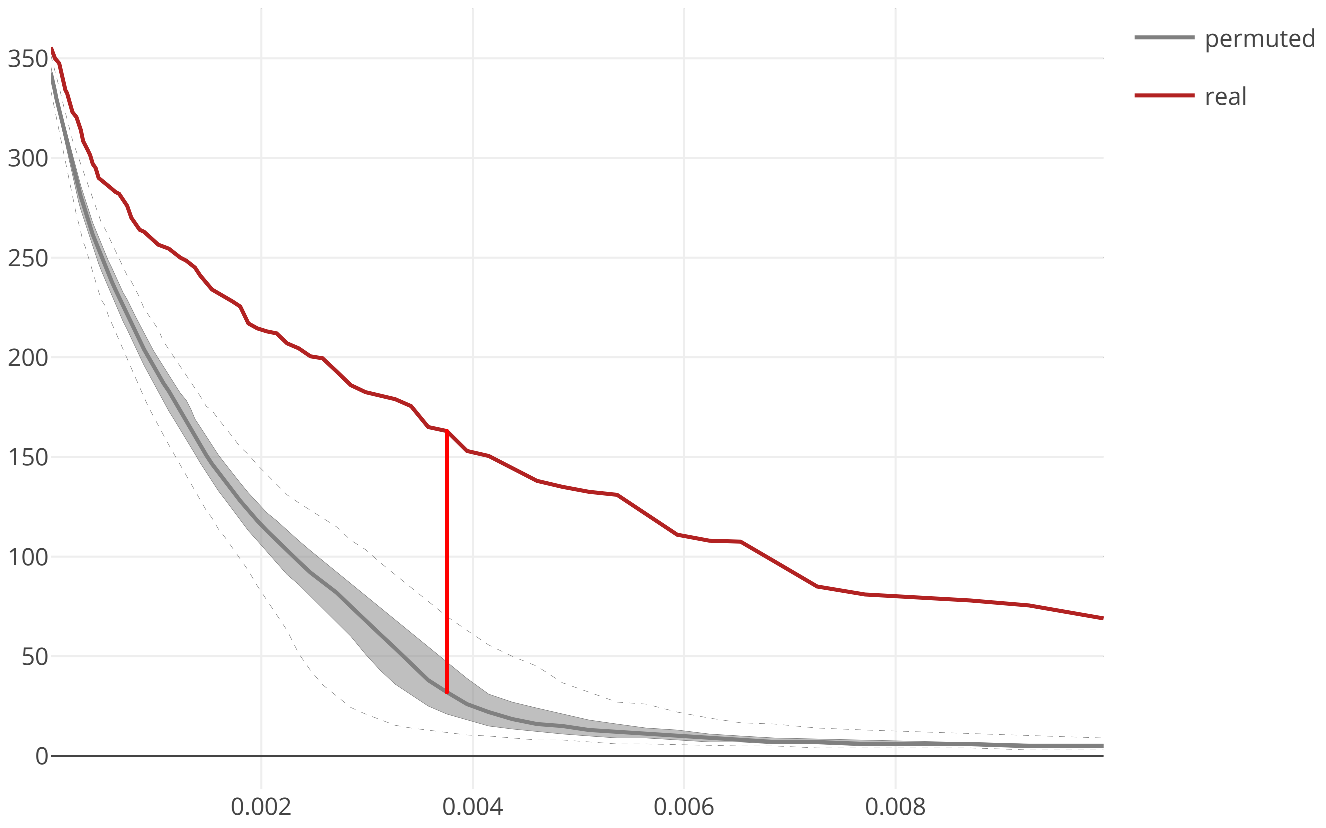

### cov2_SARS_CoV2_ORF9b_flow_distance_20200609.pdf

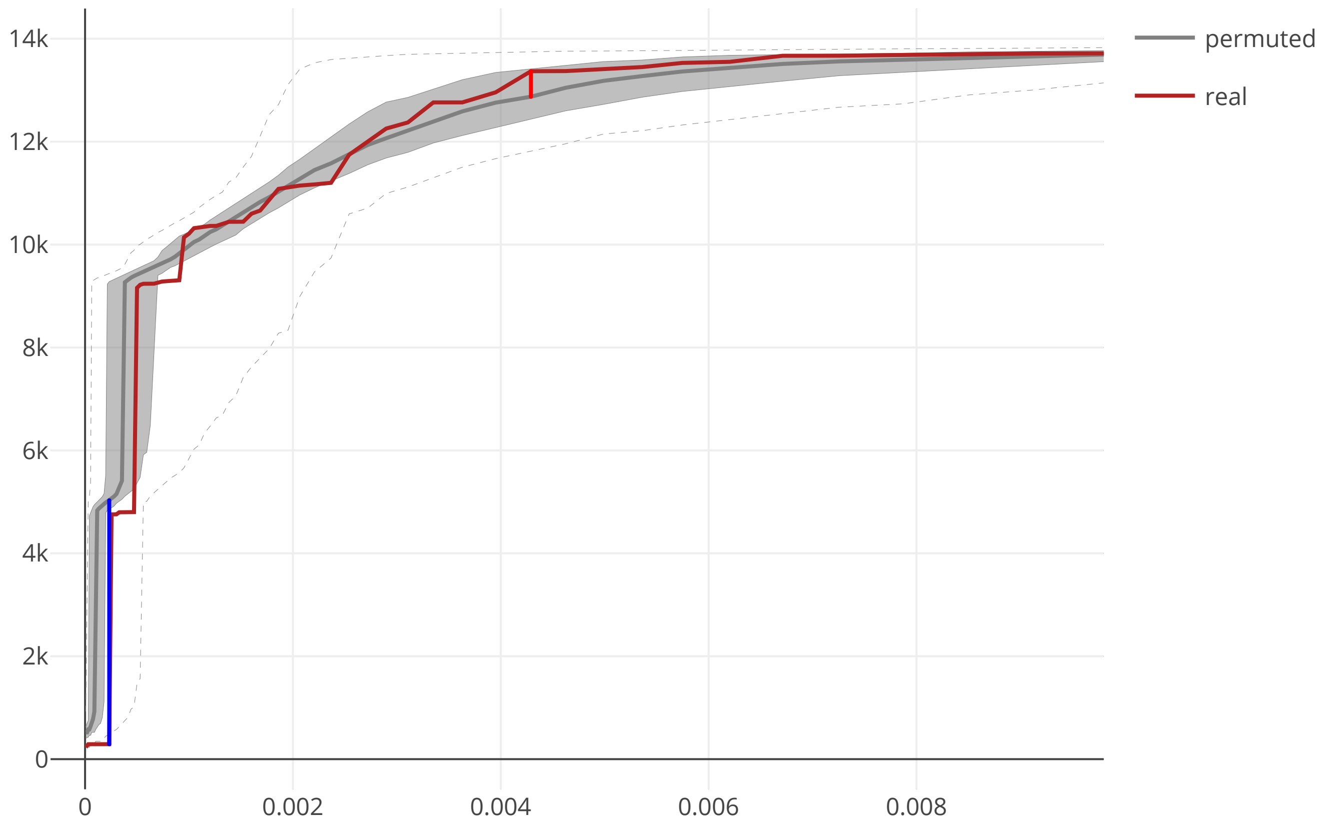

### cov2_SARS_CoV2_ORF9b_maxcomponent_size_20200609.pdf

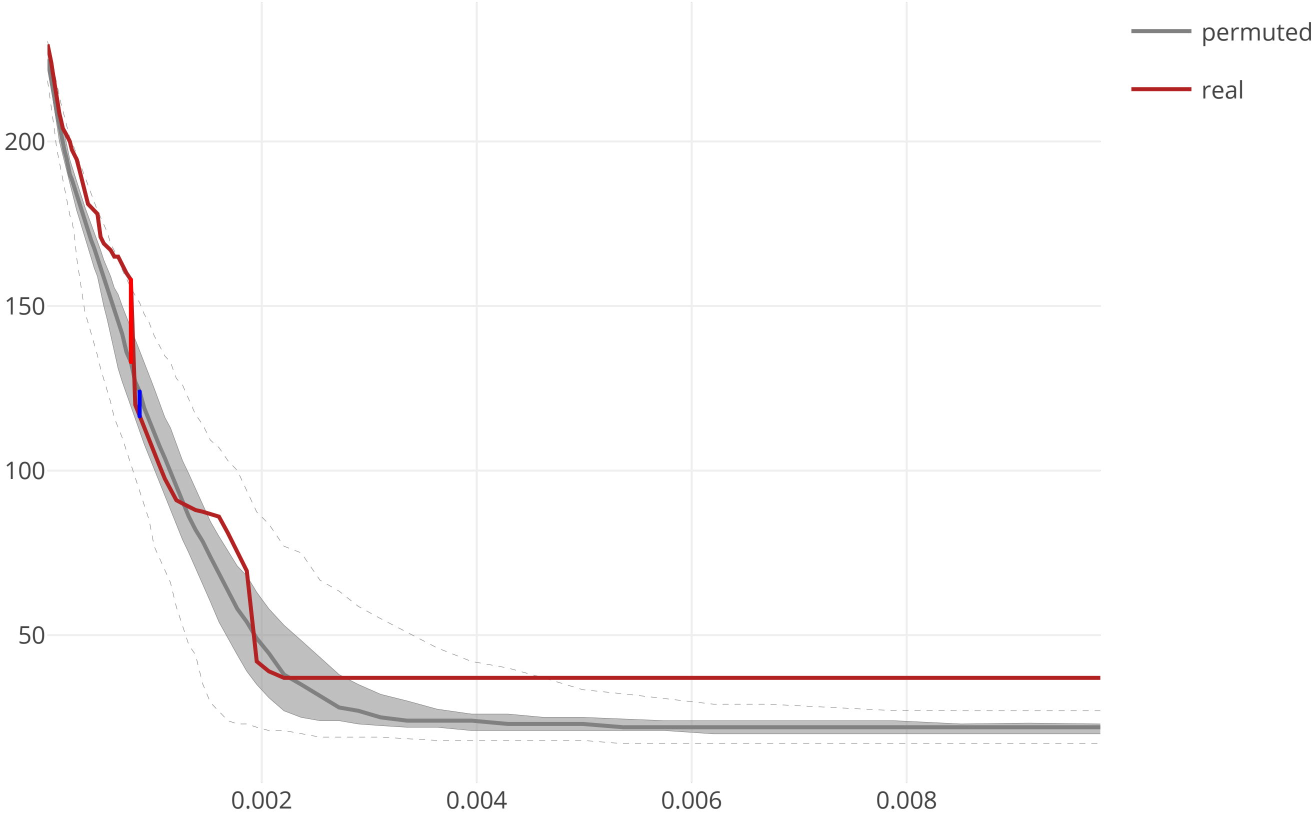

### cov2_SARS_CoV2_S_flow_distance_20200609.pdf

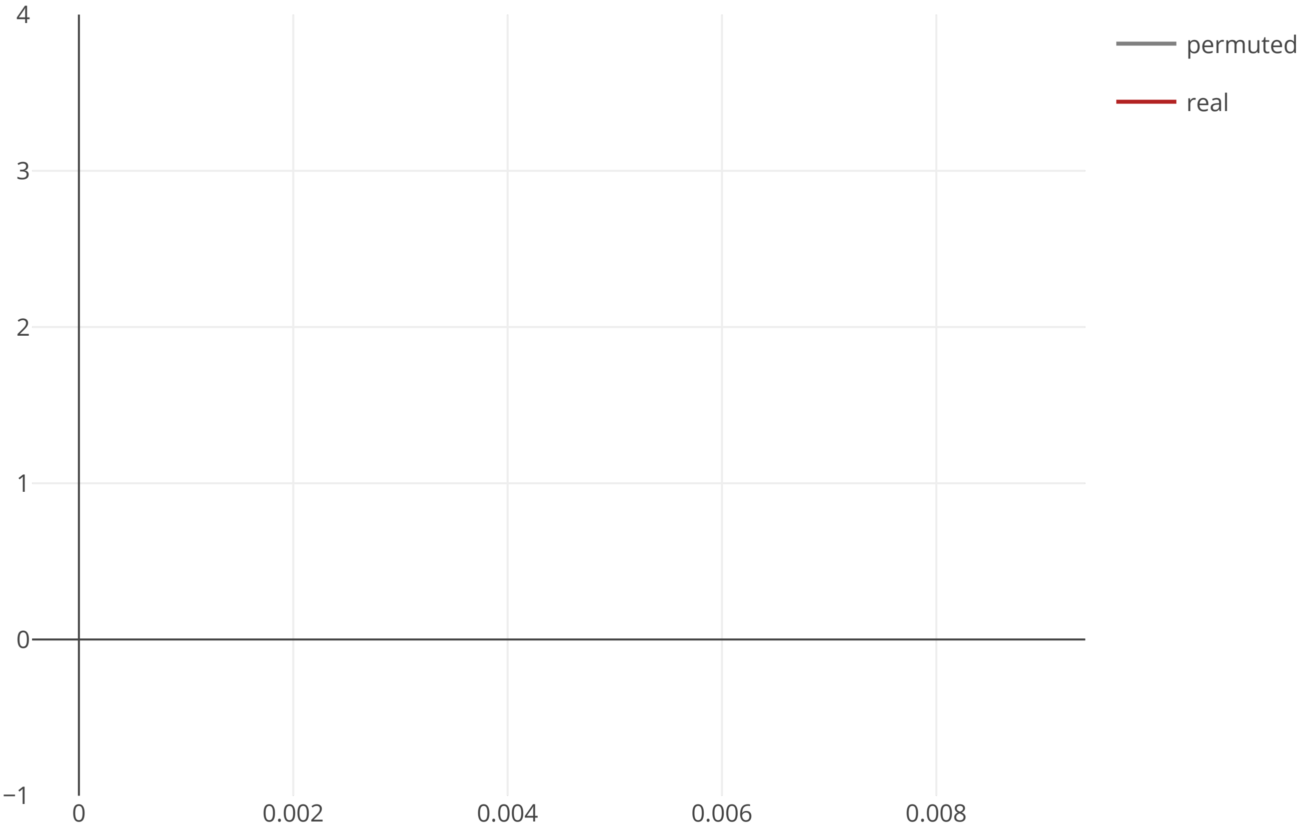

### cov2_SARS_CoV2_S_maxcomponent_size_20200609.pdf

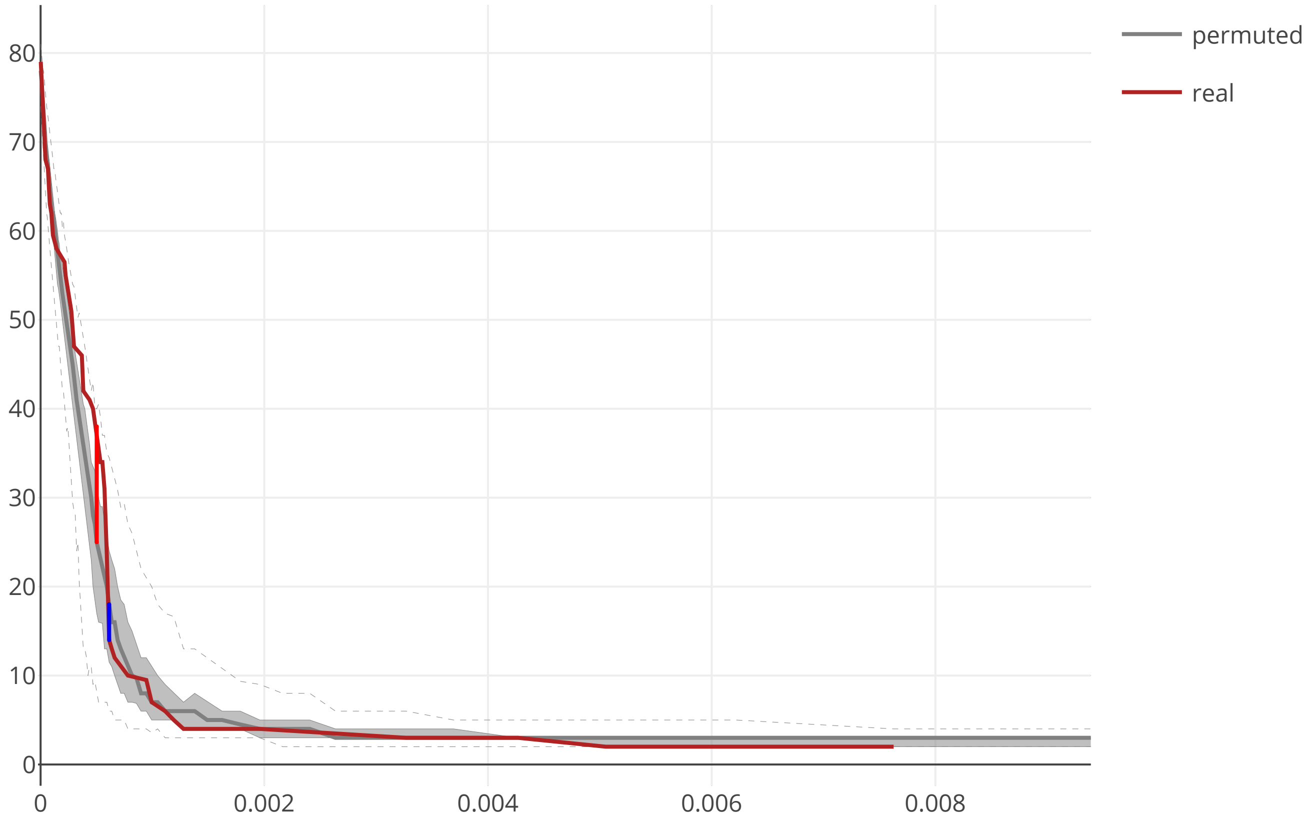

### cov2_SARS_CoV_E_flow_distance_20200609.pdf

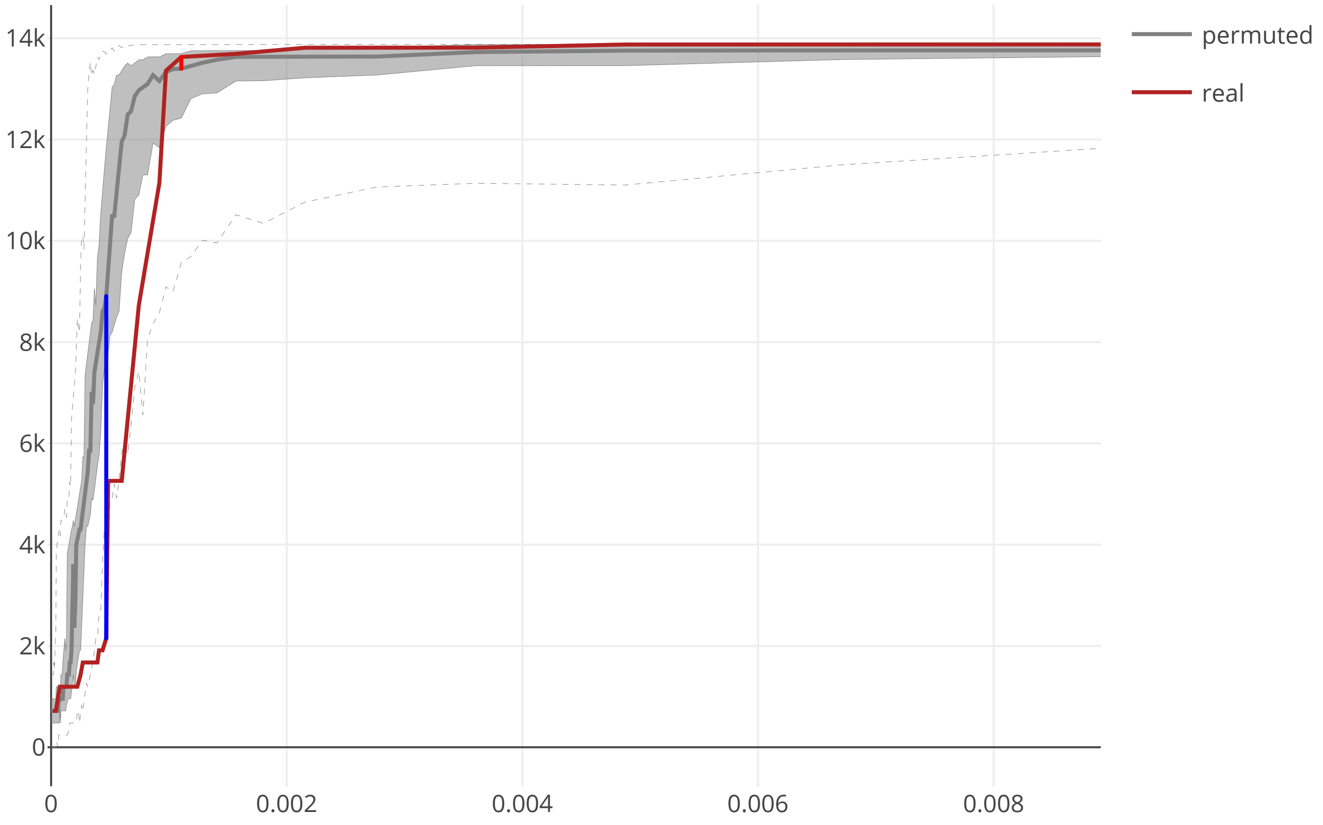

### cov2_SARS_CoV_E_maxcomponent_size_20200609.pdf

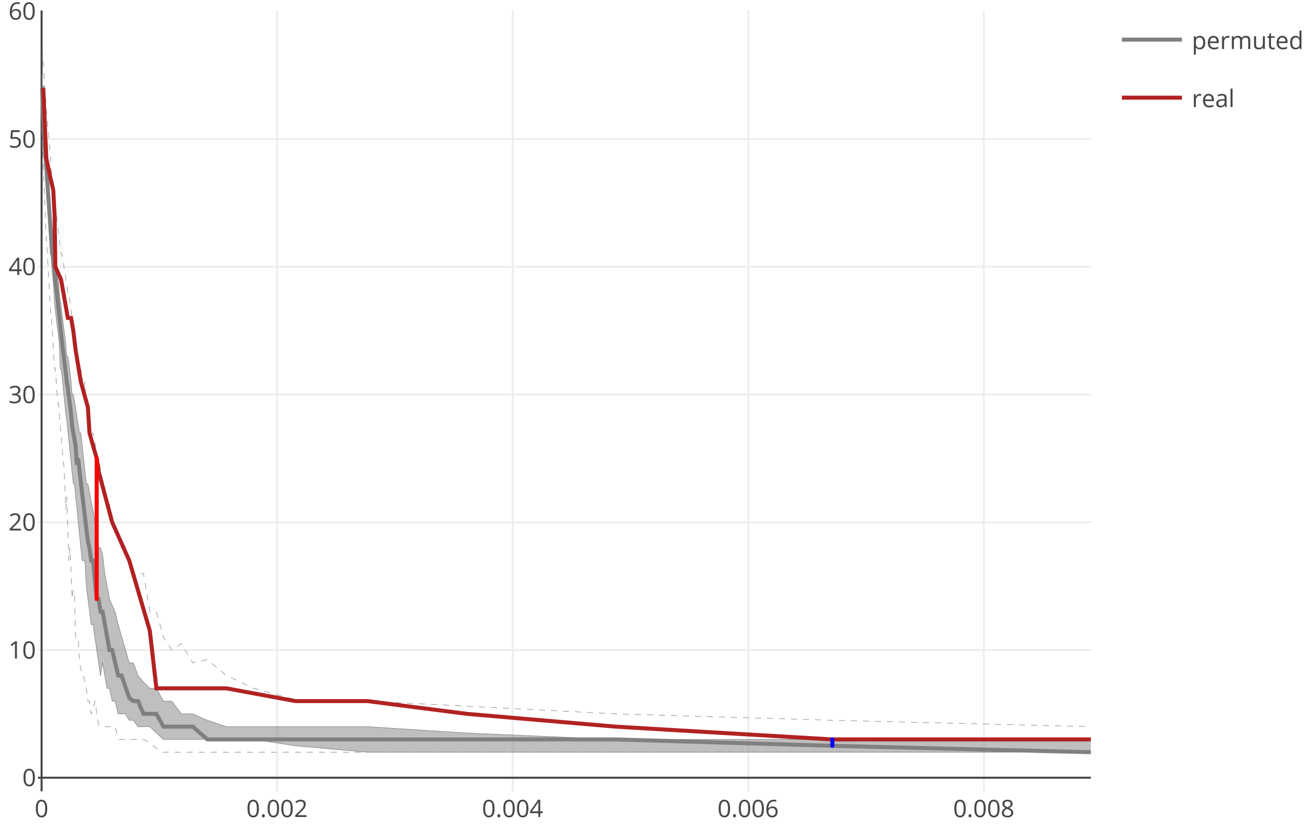

### cov2_SARS_CoV_M_flow_distance_20200609.pdf

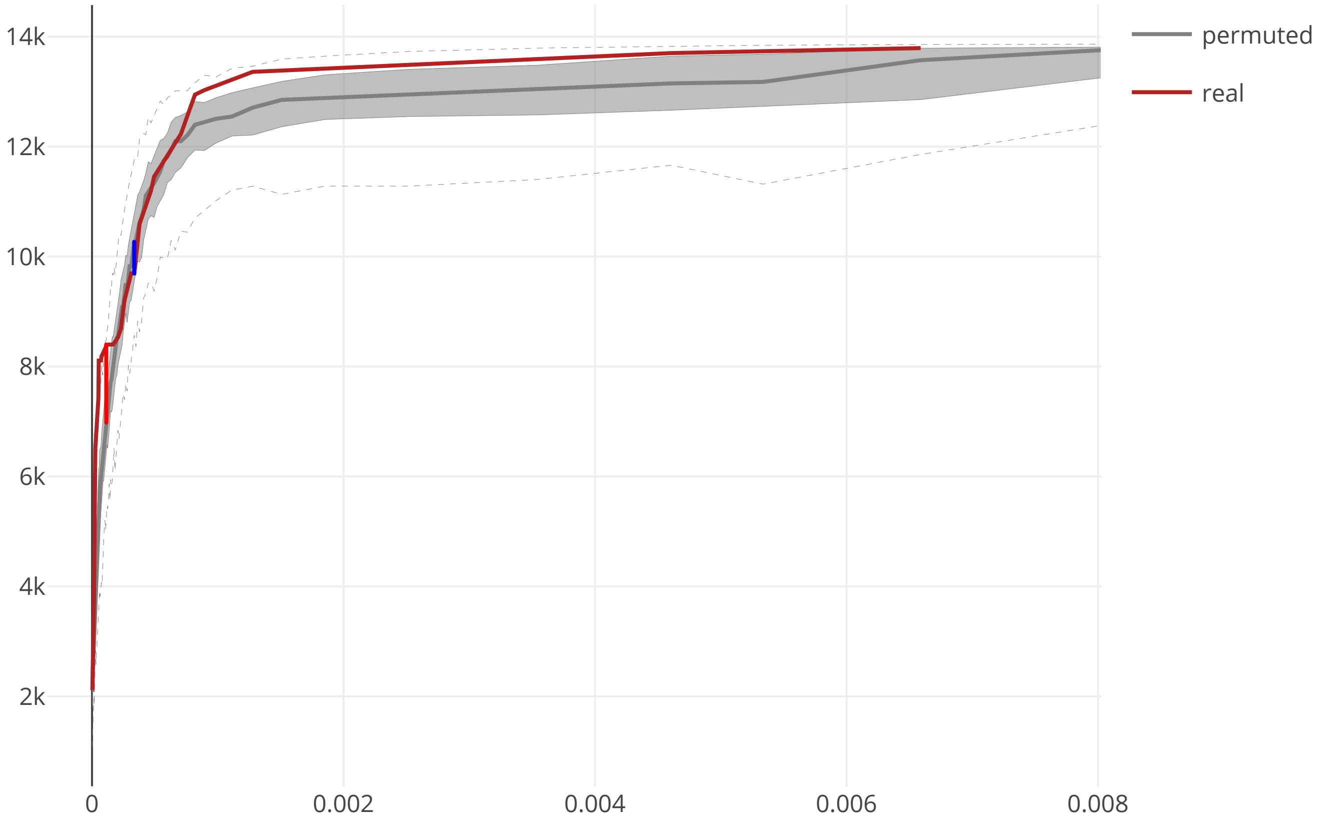

### cov2_SARS_CoV_M_maxcomponent_size_20200609.pdf

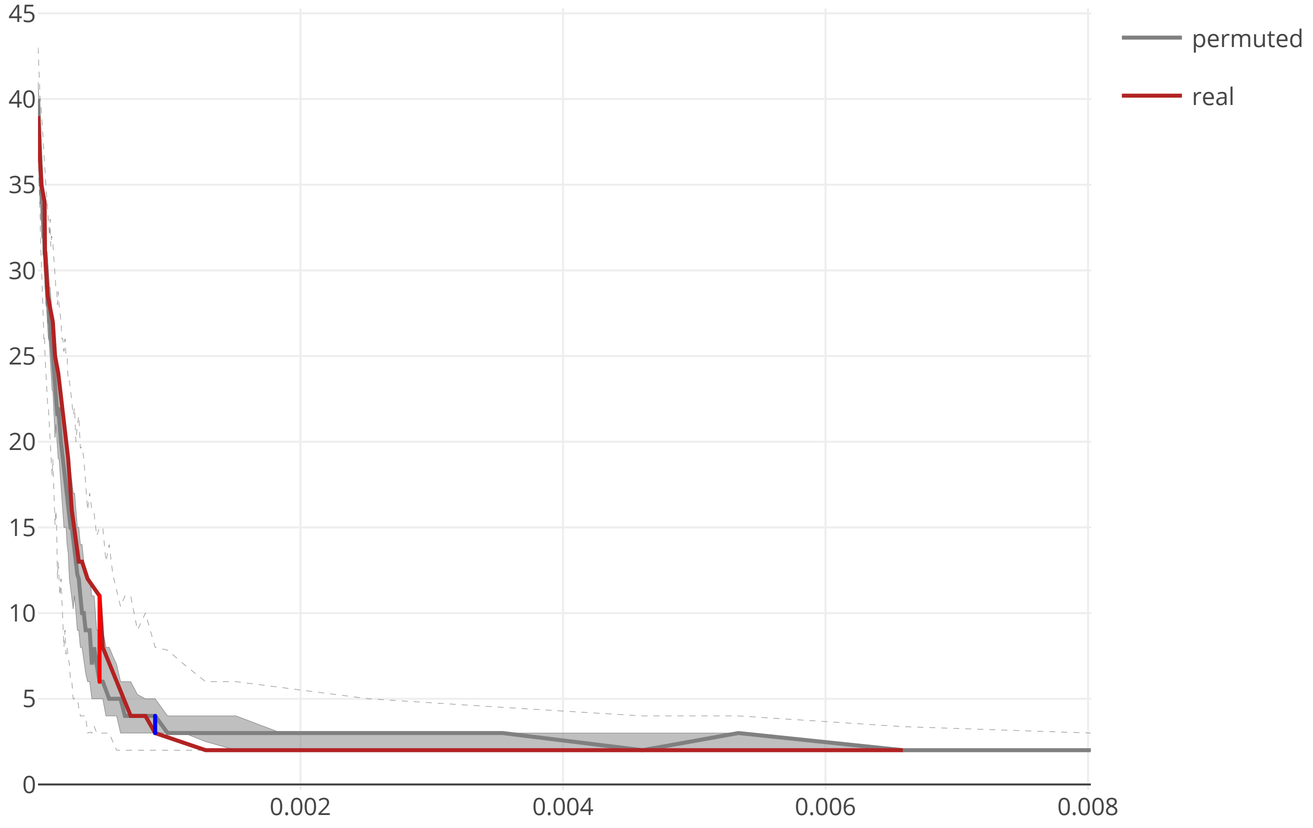

### cov2_SARS_CoV_N_flow_distance_20200609.pdf

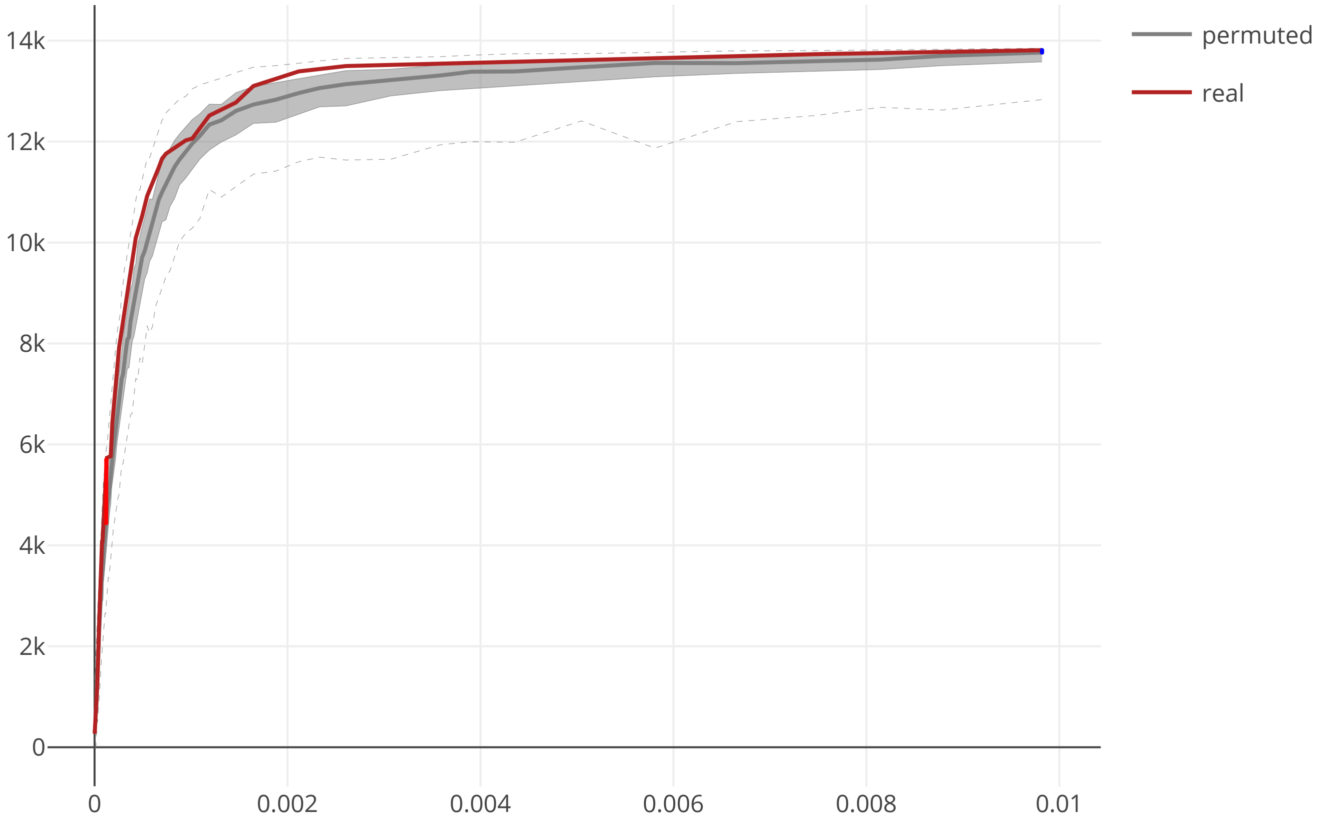

### cov2_SARS_CoV_N_maxcomponent_size_20200609.pdf

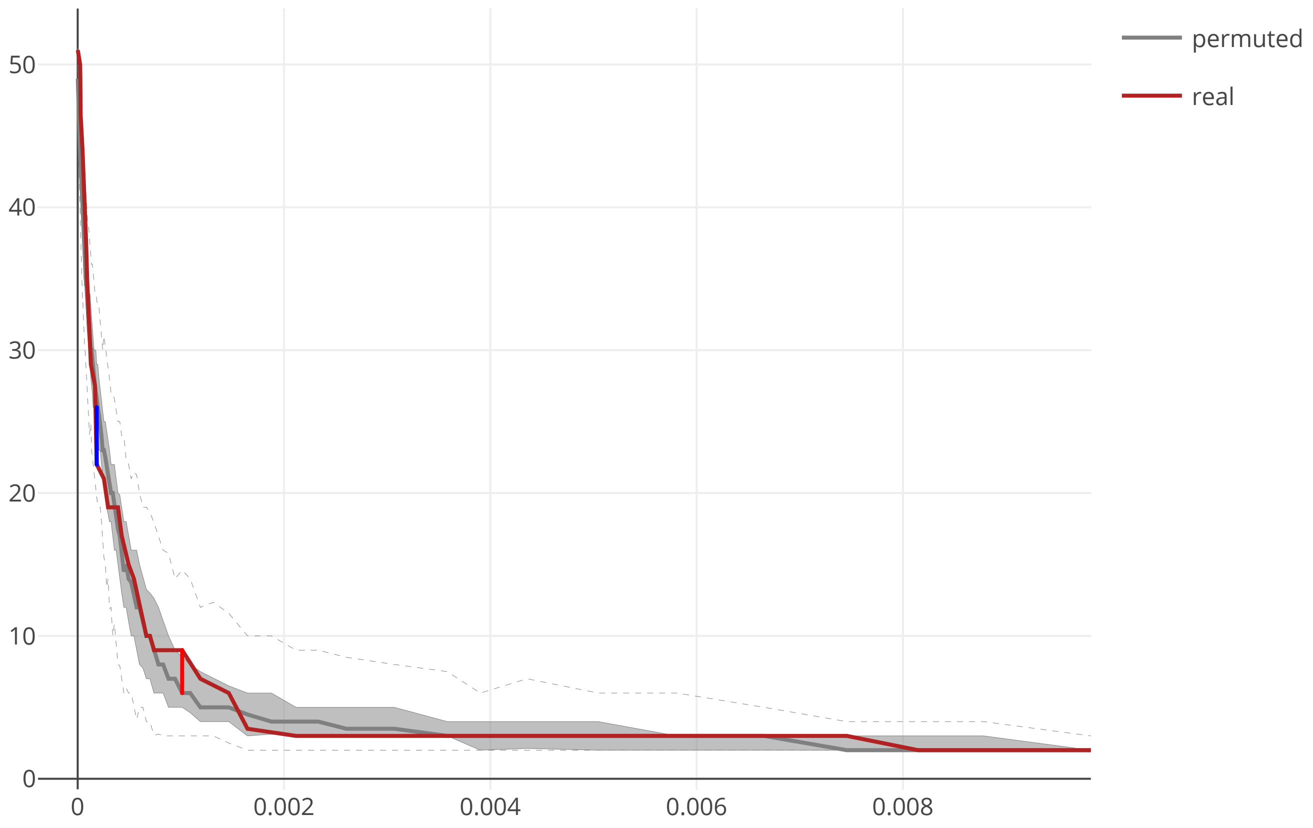

### cov2_SARS_CoV_NSP2_flow_distance_20200609.pdf

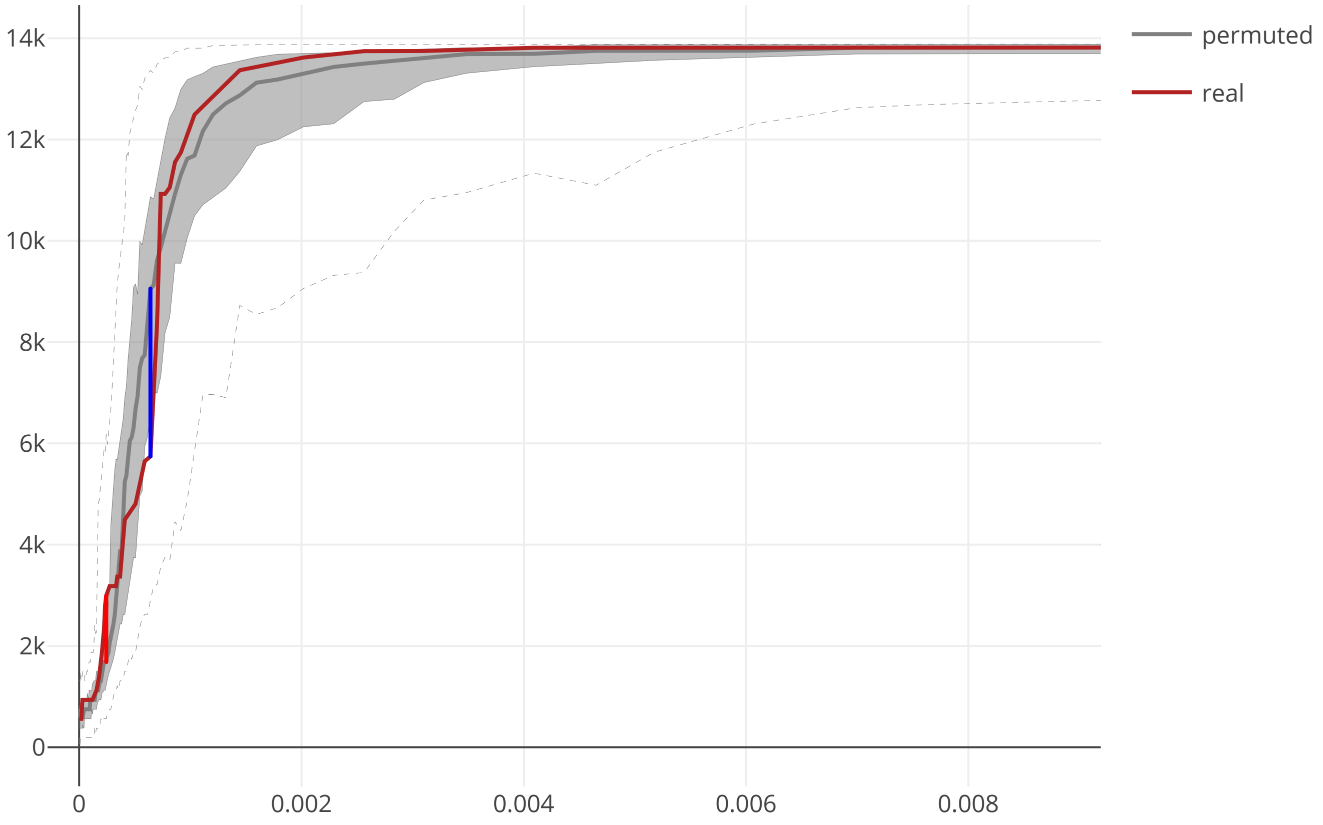

### cov2_SARS_CoV_NSP2_maxcomponent_size_20200609.pdf

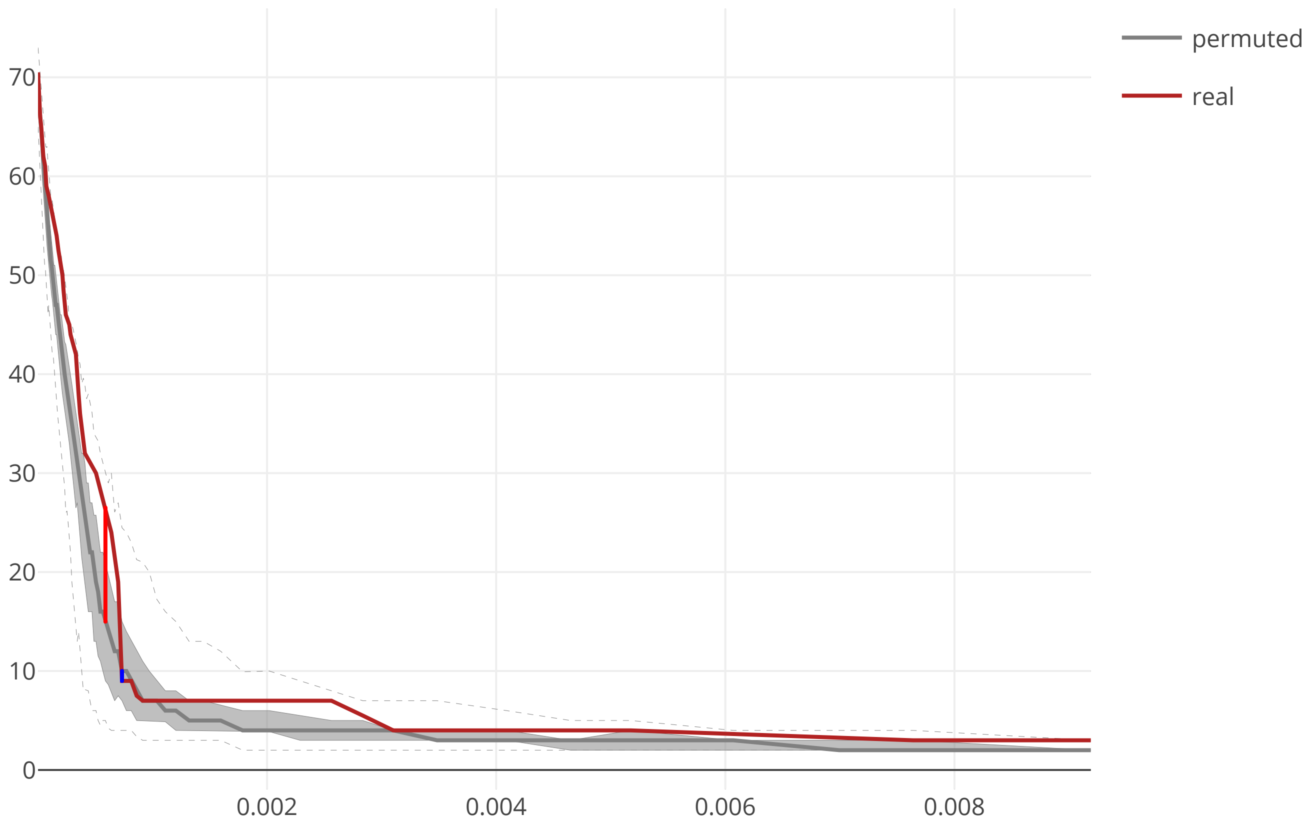

### cov2_SARS_CoV_NSP3_macroD_flow_distance_20200609.pdf

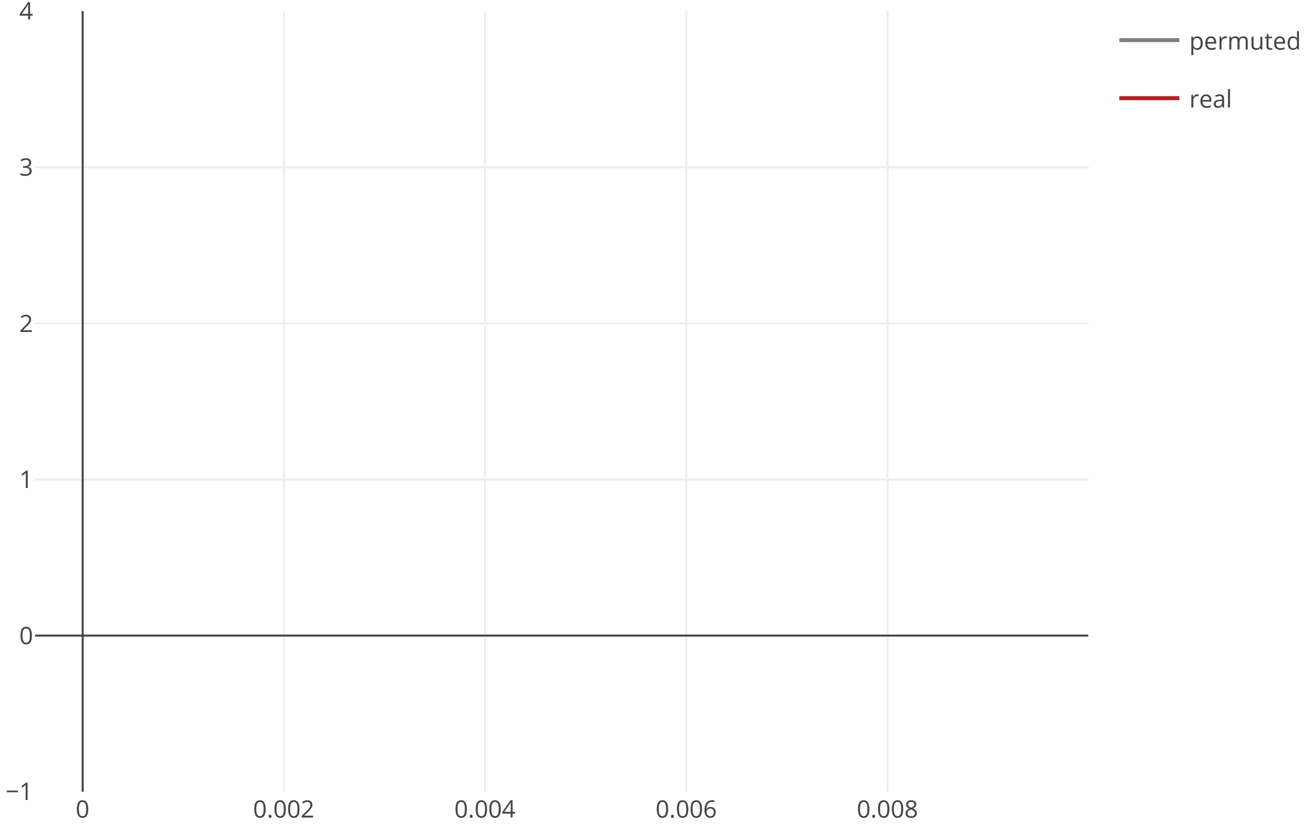

### cov2_SARS_CoV_NSP3_macroD_maxcomponent_size_20200609.pdf

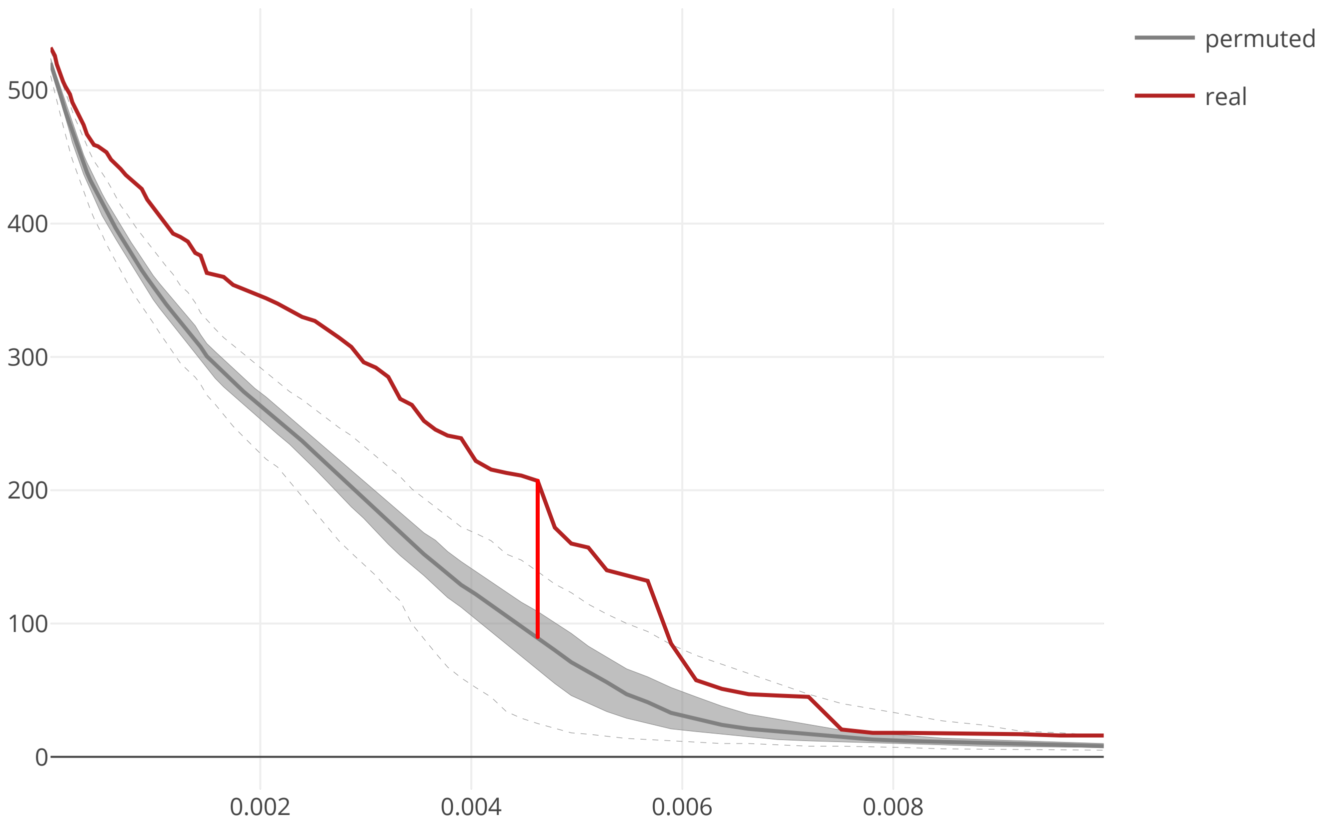

### cov2_SARS_CoV_NSP4_flow_distance_20200609.pdf

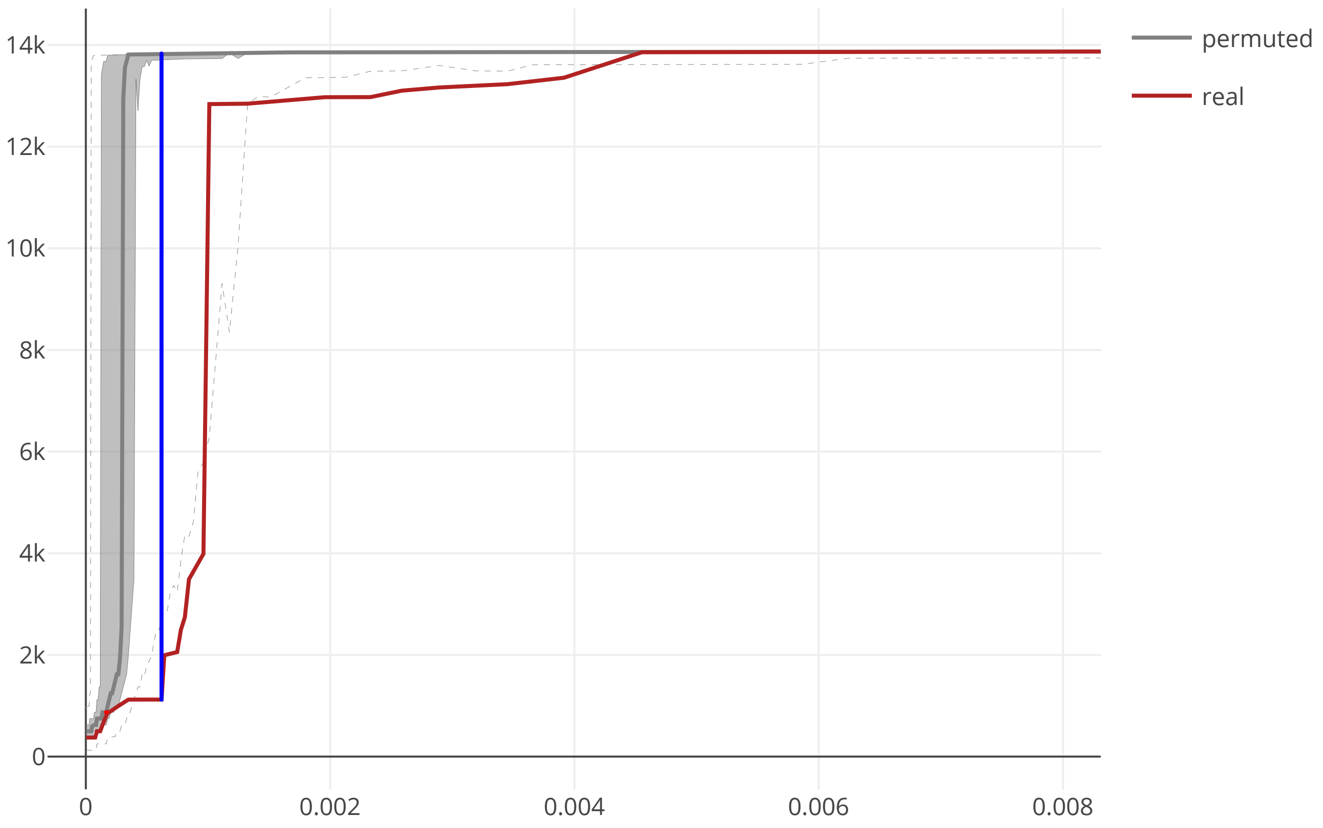

### cov2_SARS_CoV_NSP4_maxcomponent_size_20200609.pdf

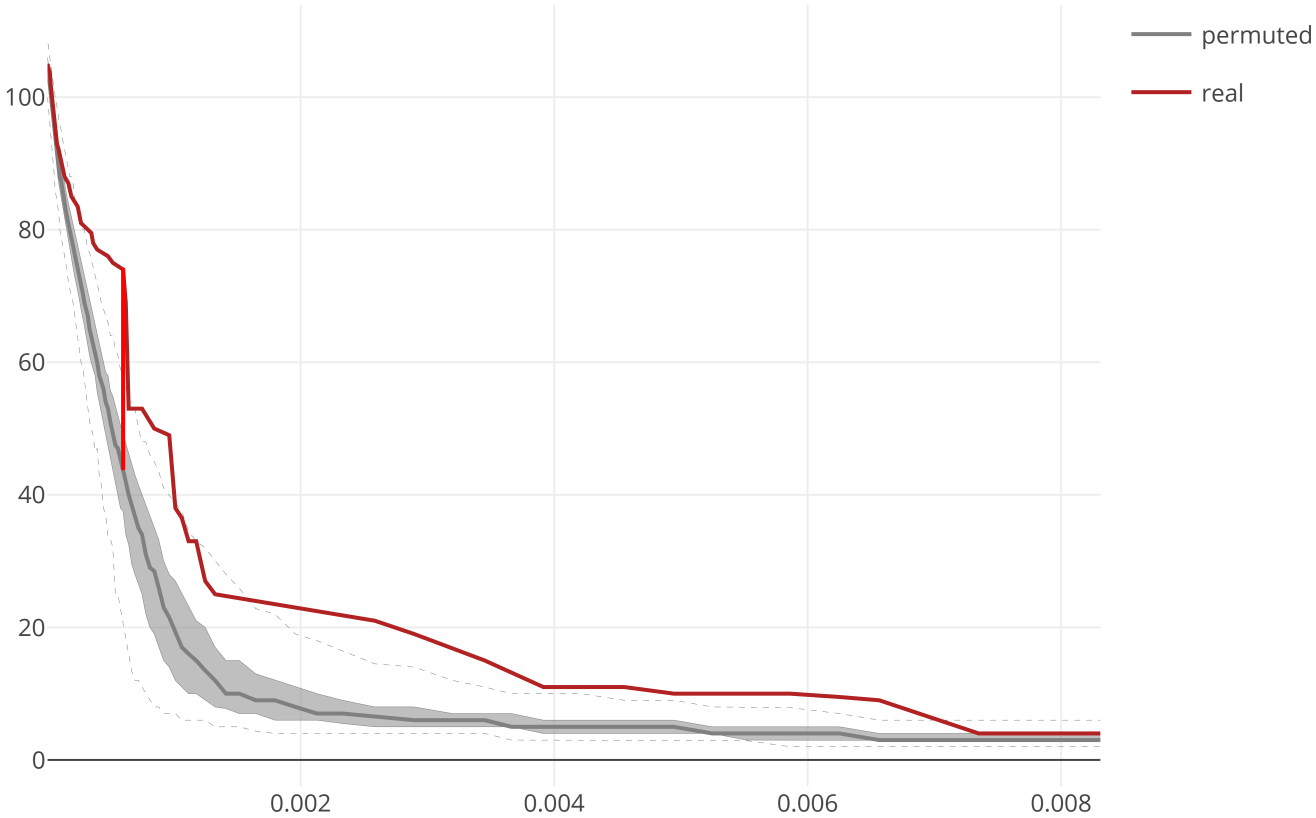

### cov2_SARS_CoV_NSP6_flow_distance_20200609.pdf

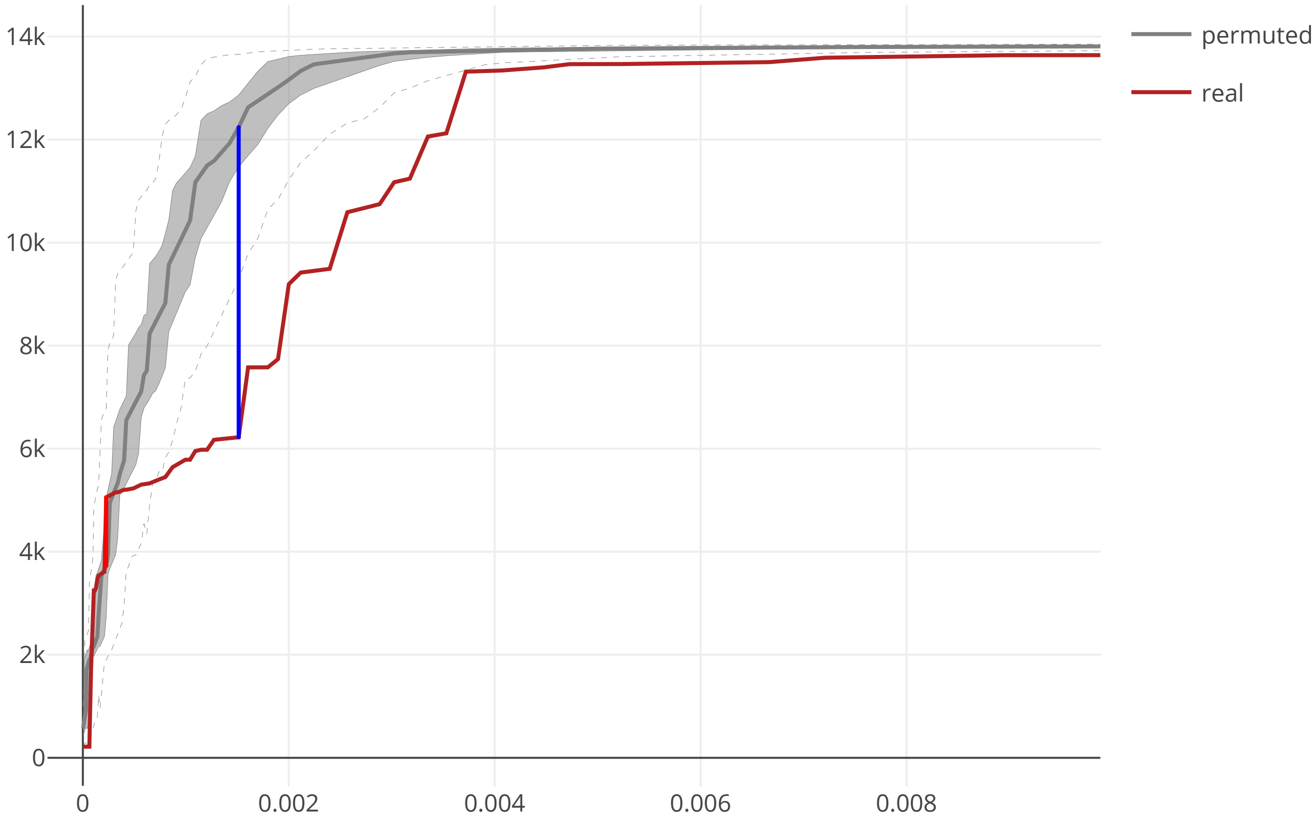

### cov2_SARS_CoV_NSP6_maxcomponent_size_20200609.pdf

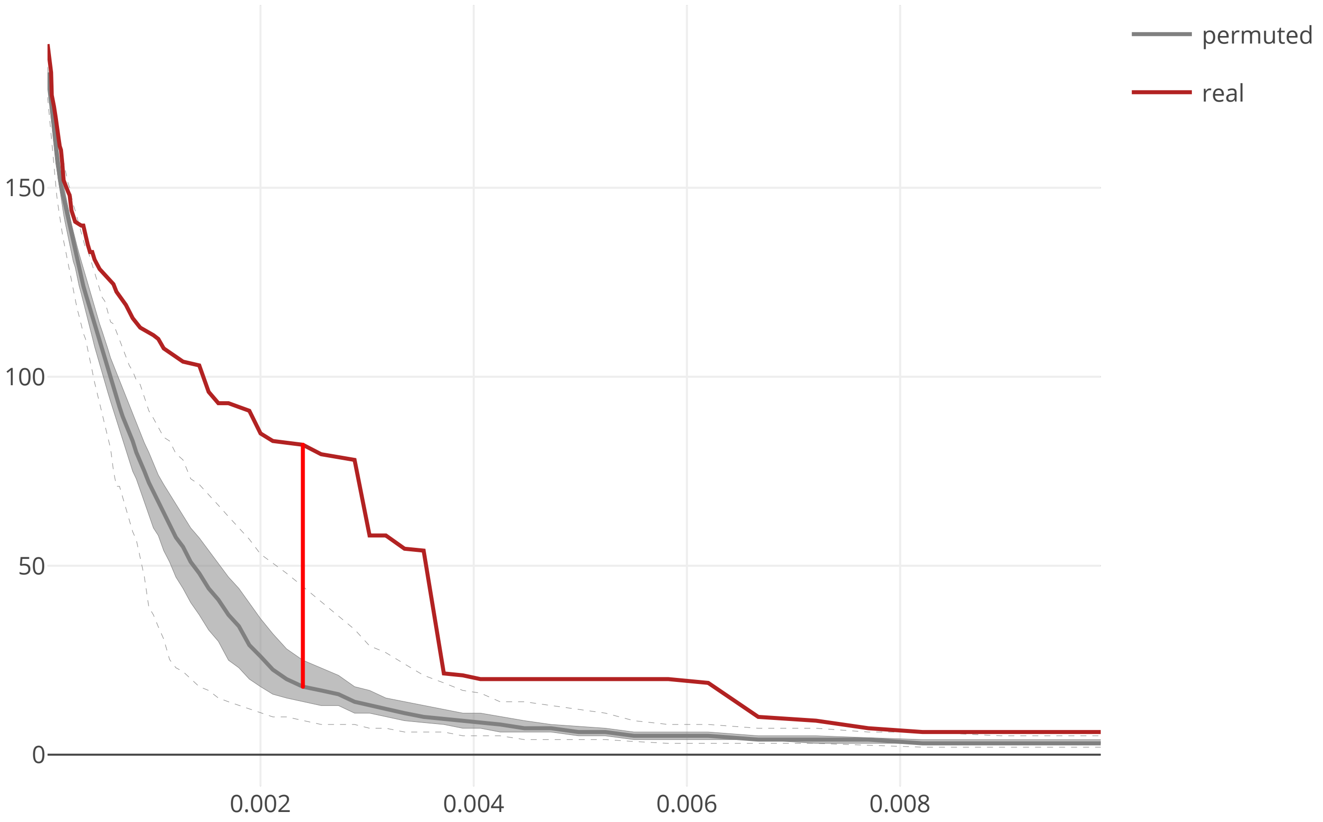

### cov2_SARS_CoV_NSP7_flow_distance_20200609.pdf

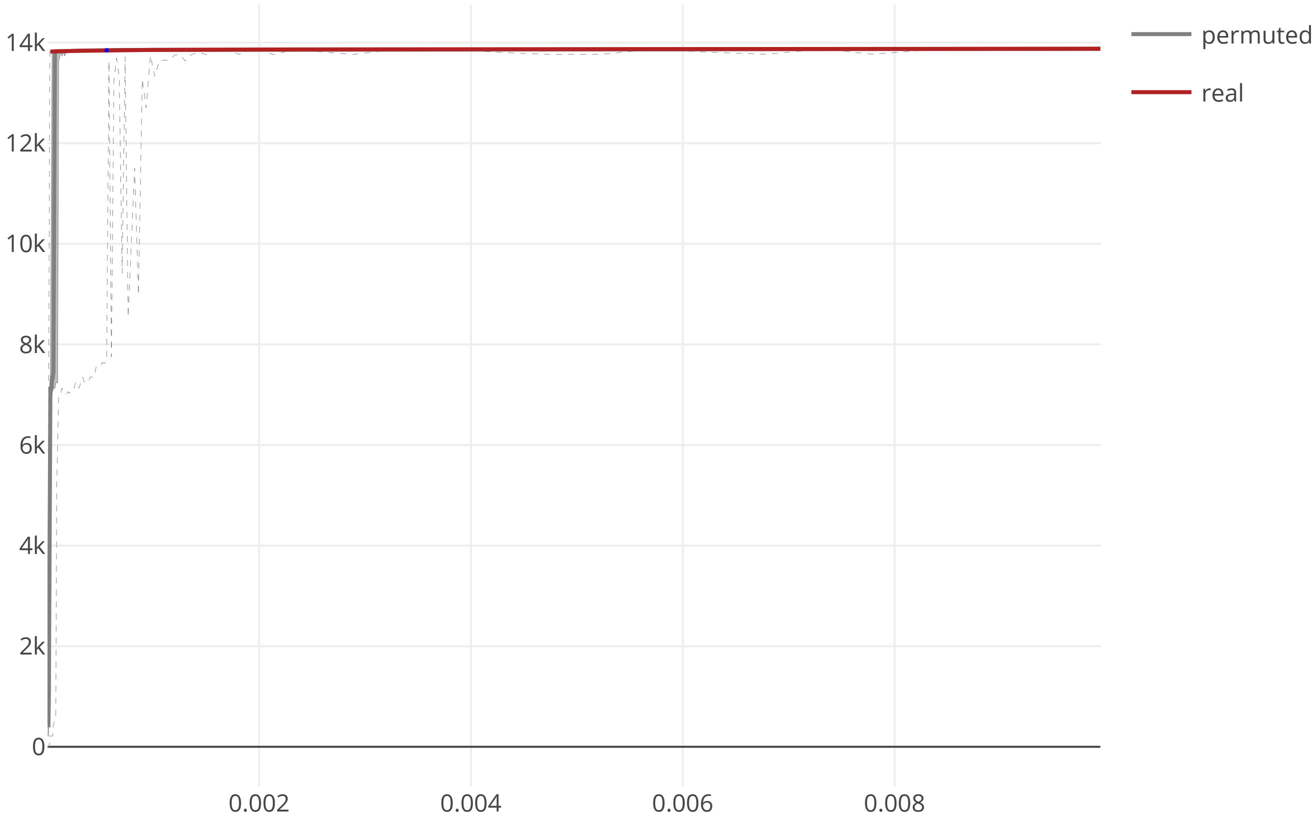

### cov2_SARS_CoV_NSP7_maxcomponent_size_20200609.pdf

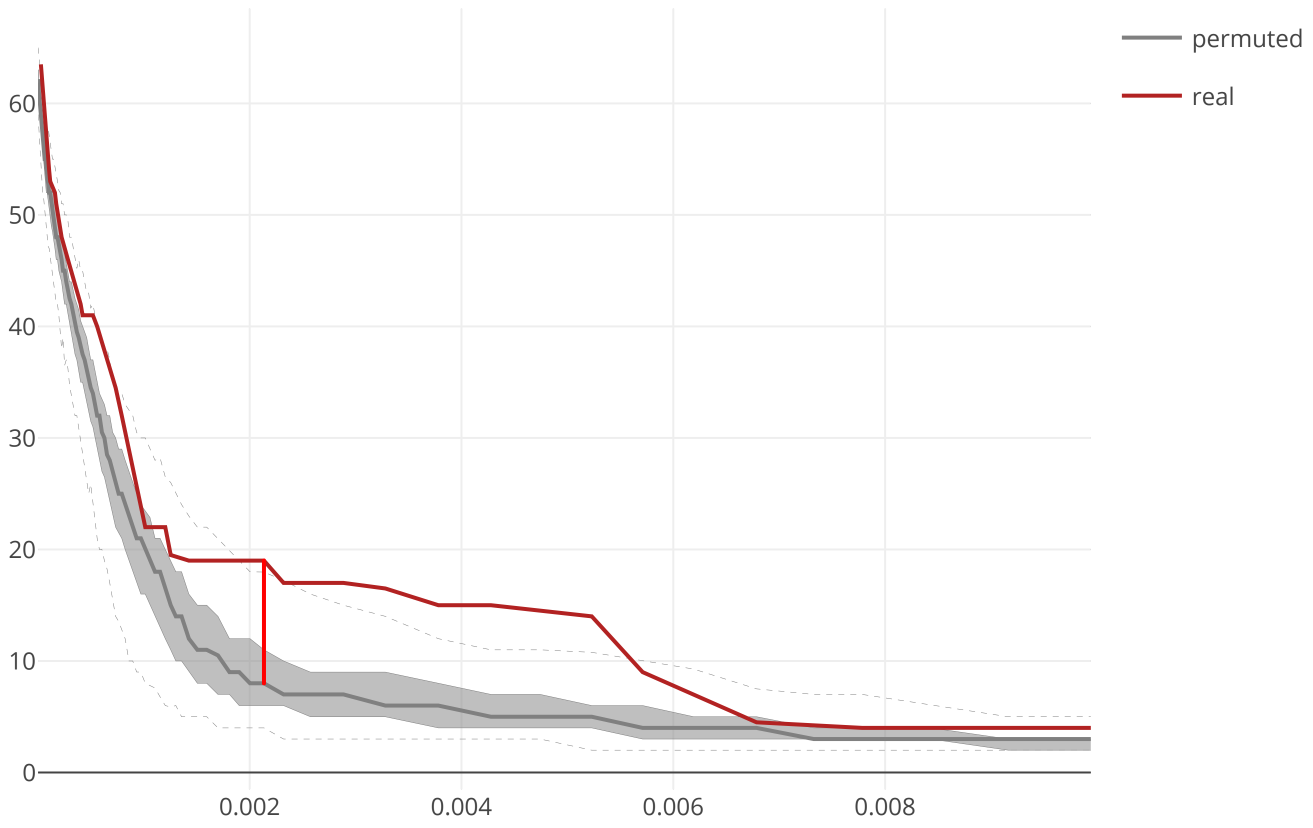

### cov2_SARS_CoV_NSP8_flow_distance_20200609.pdf

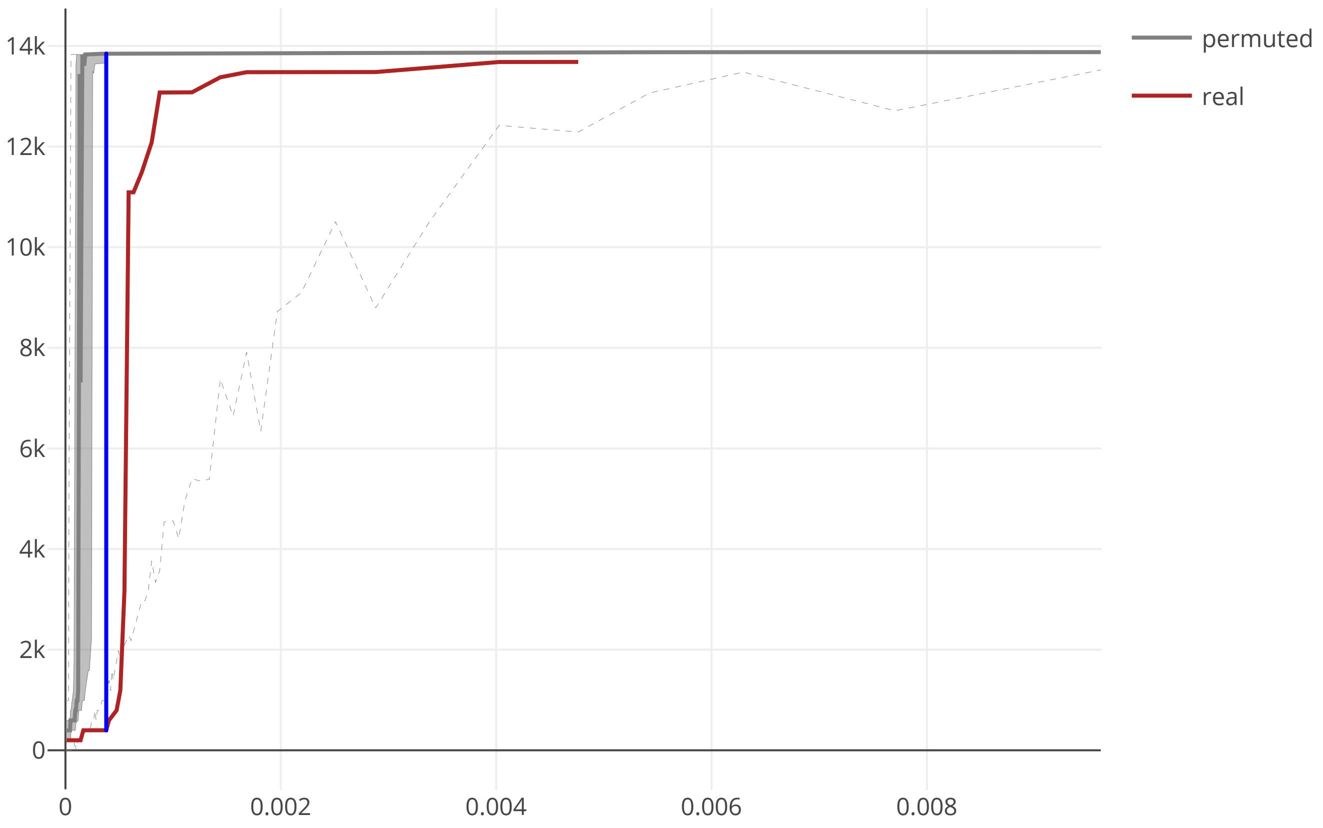

### cov2_SARS_CoV_NSP8_maxcomponent_size_20200609.pdf

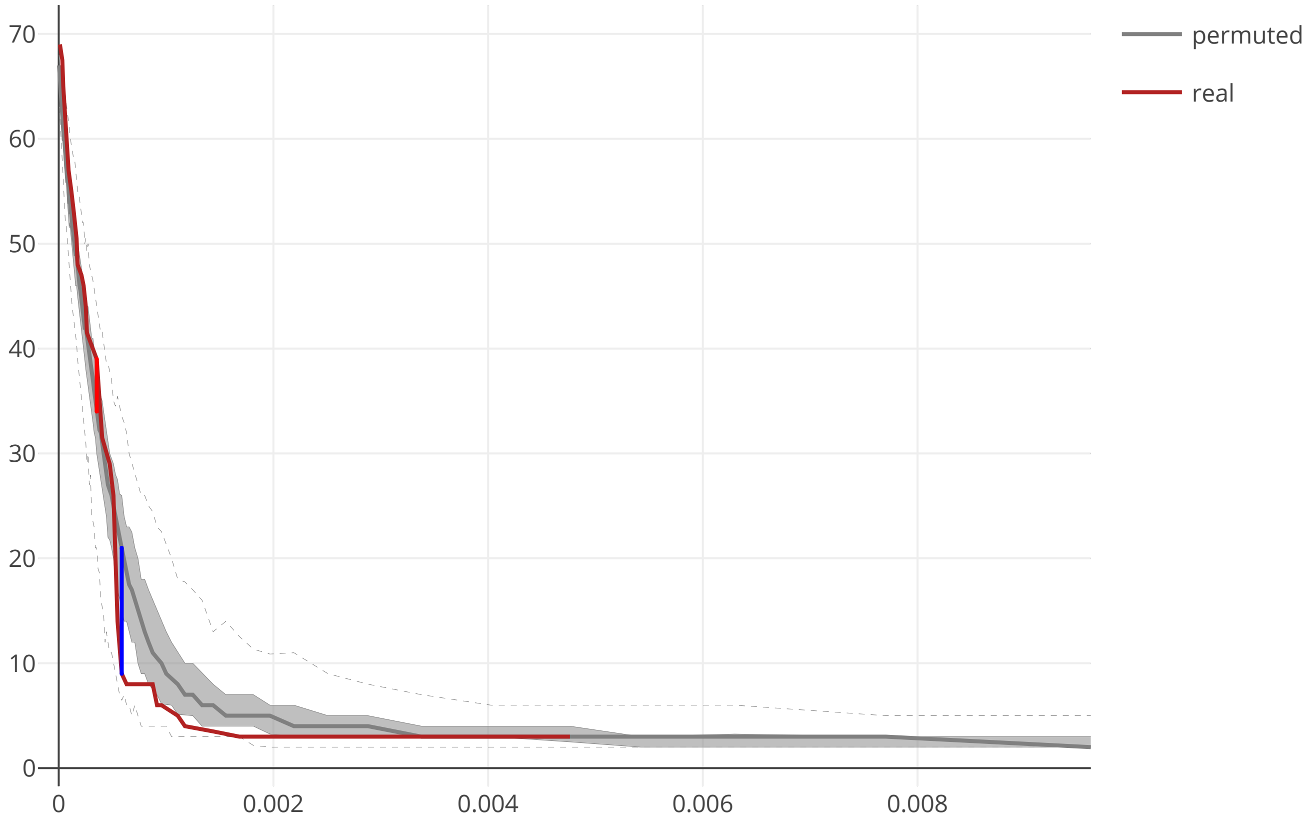

### cov2_SARS_CoV_NSP9_flow_distance_20200609.pdf

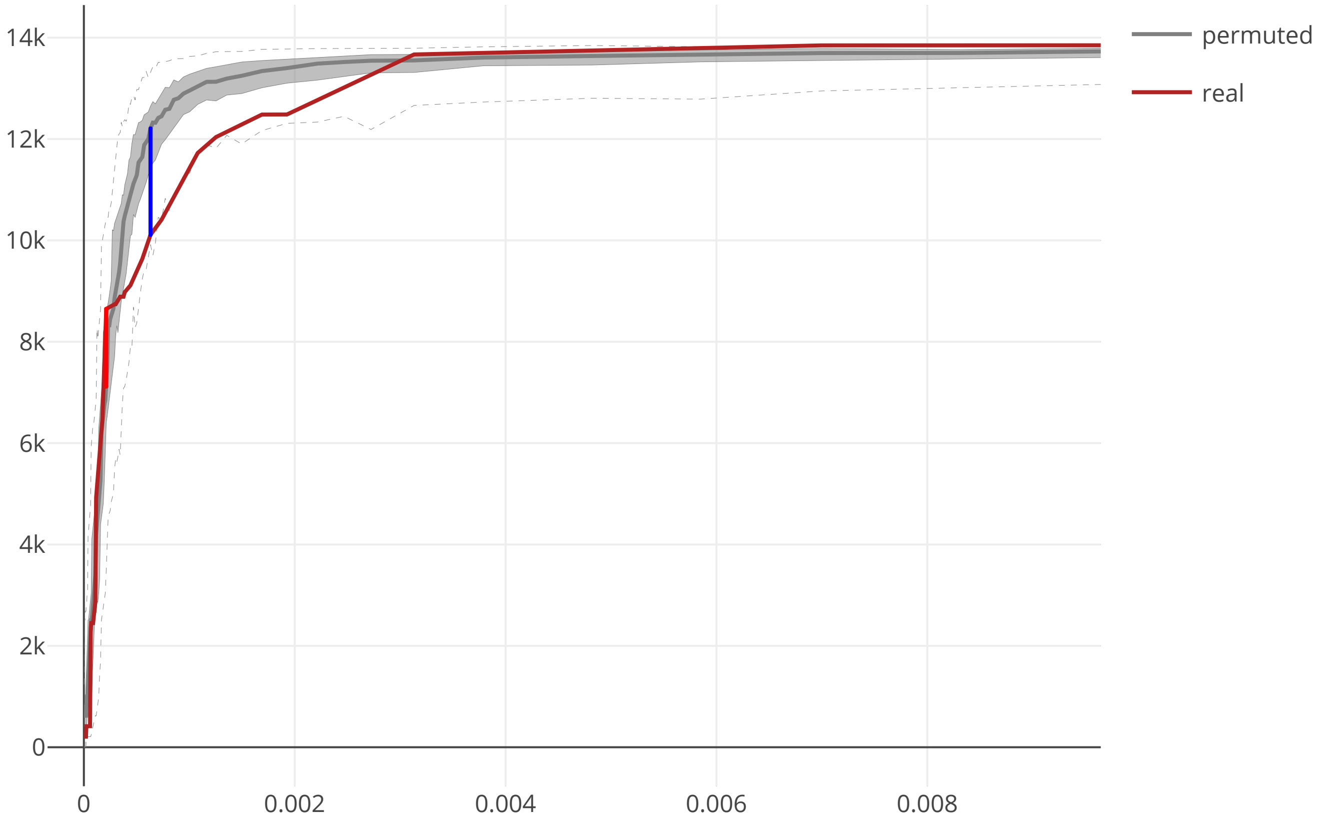

### cov2_SARS_CoV_NSP9_maxcomponent_size_20200609.pdf

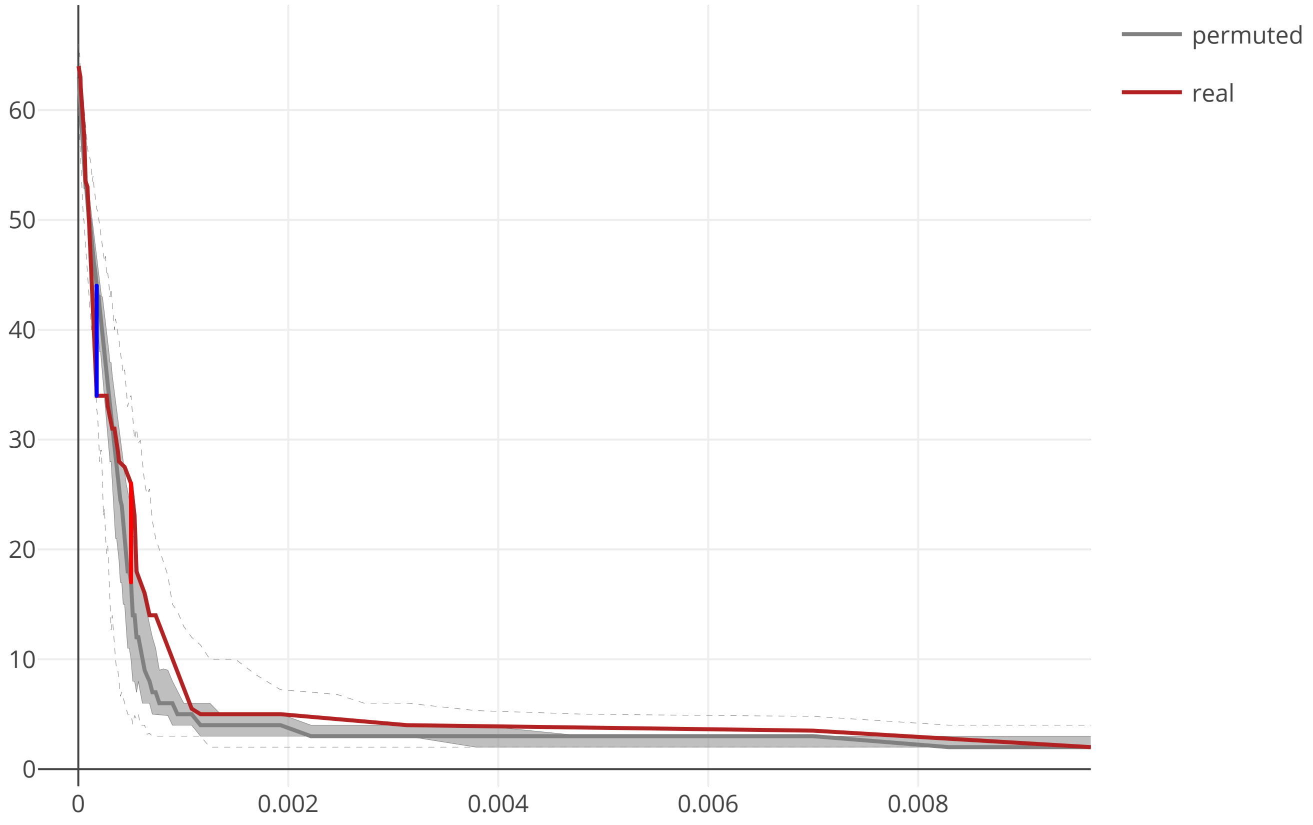

### cov2_SARS_CoV_NSP10_flow_distance_20200609.pdf

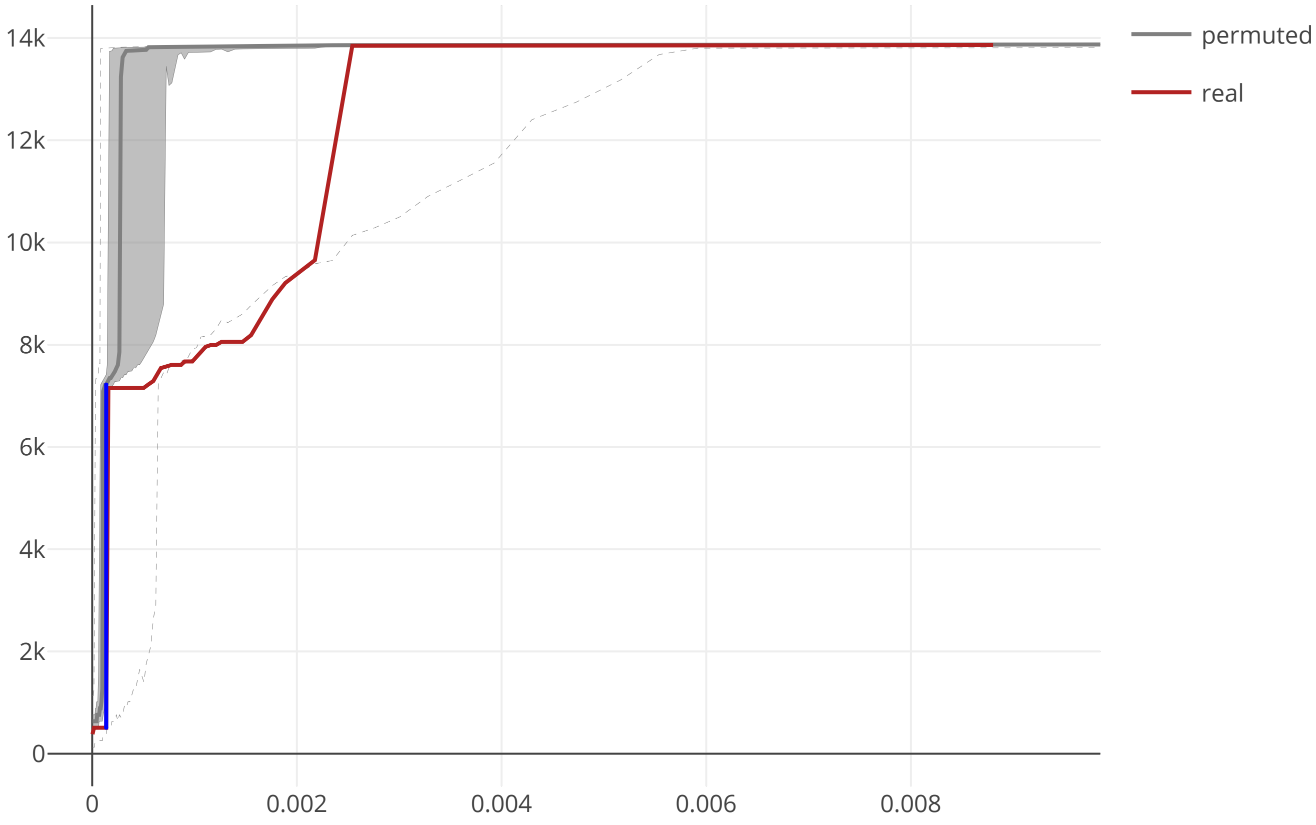

### cov2_SARS_CoV_NSP10_maxcomponent_size_20200609.pdf

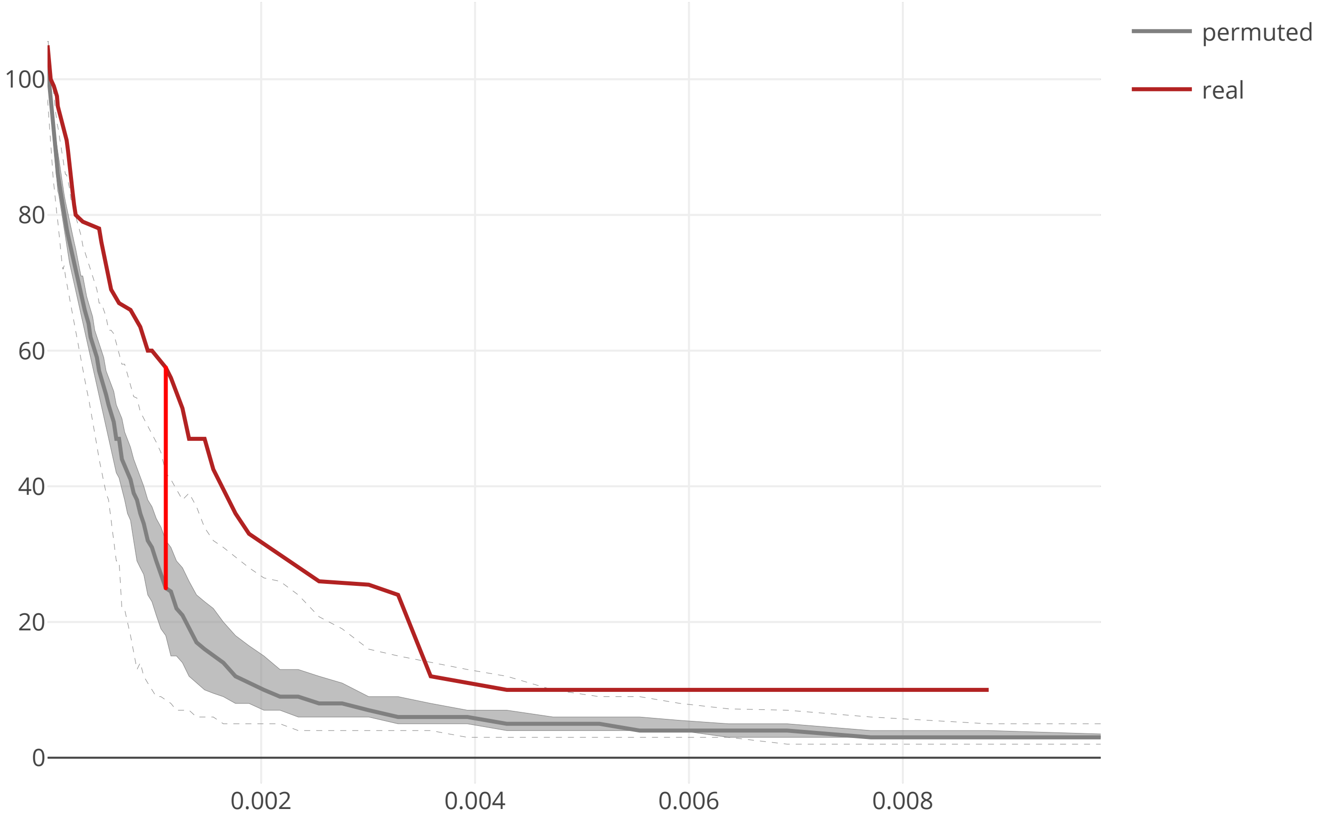

### cov2_SARS_CoV_NSP12_flow_distance_20200609.pdf

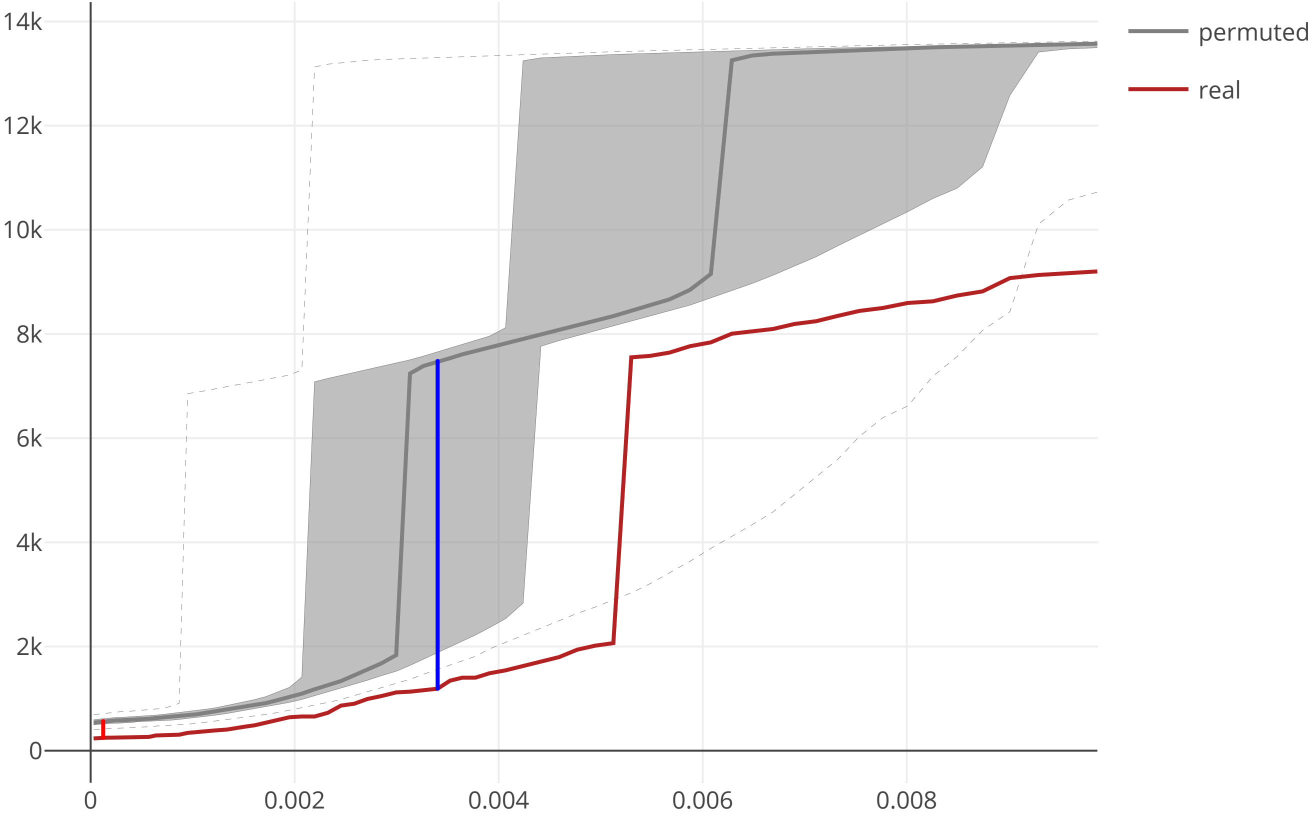

### cov2_SARS_CoV_NSP12_maxcomponent_size_20200609.pdf

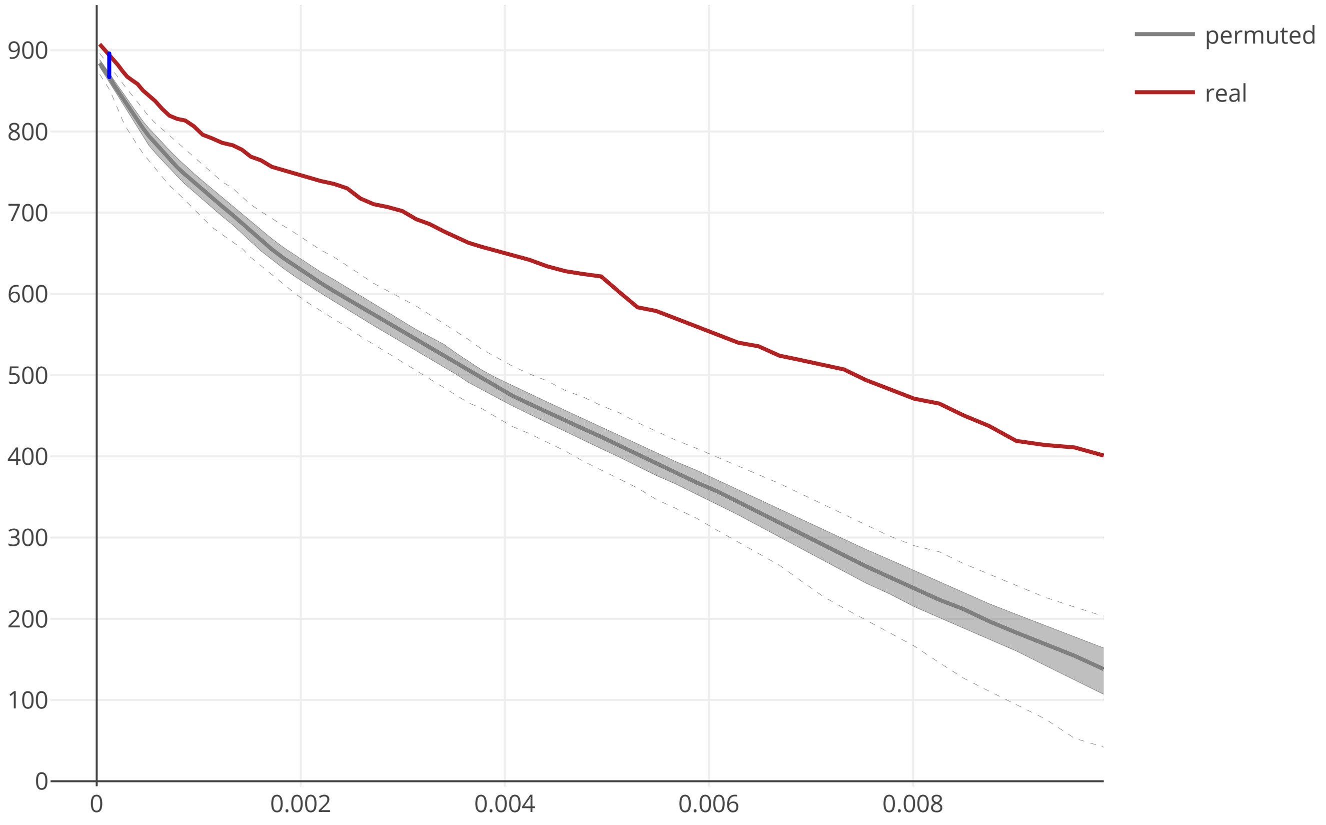

### cov2_SARS_CoV_NSP13_flow_distance_20200609.pdf

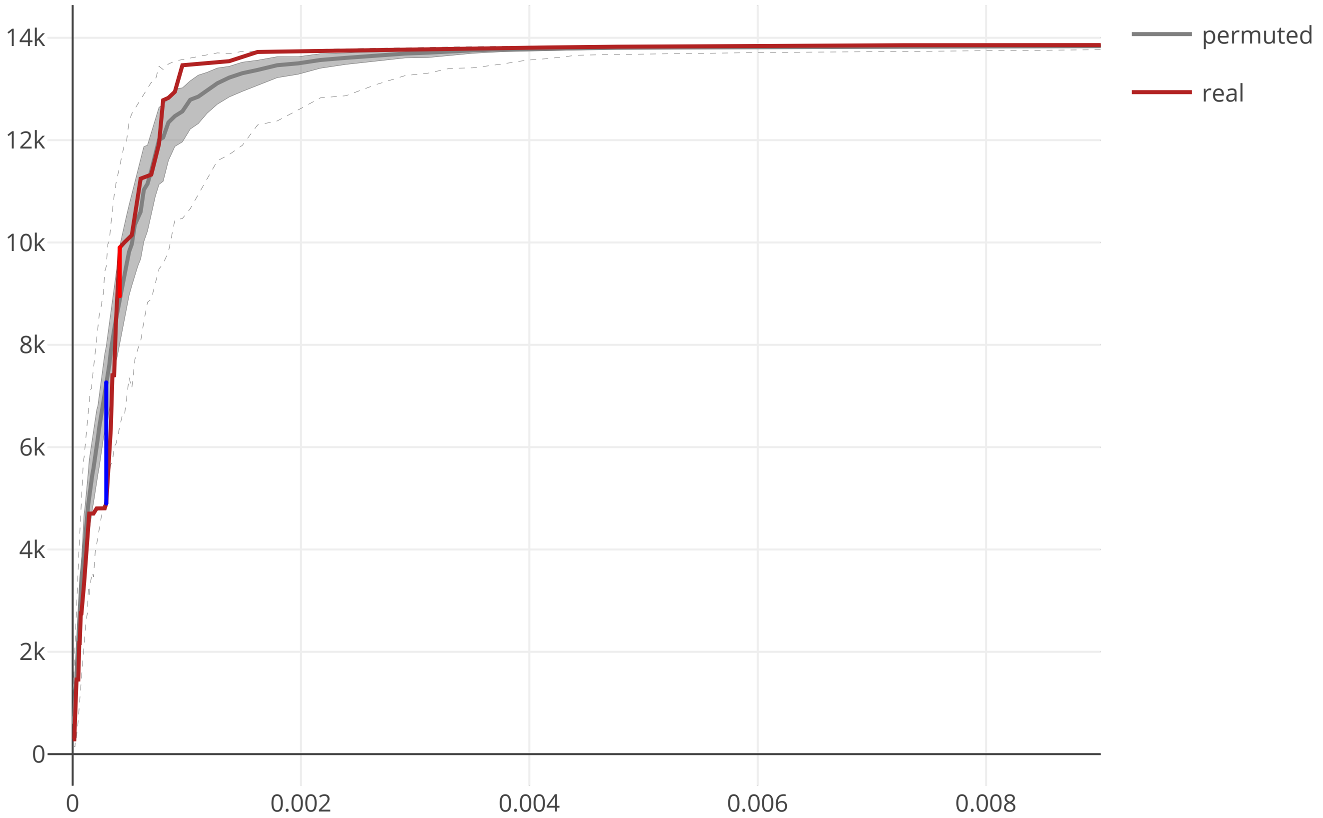

### cov2_SARS_CoV_NSP13_maxcomponent_size_20200609.pdf

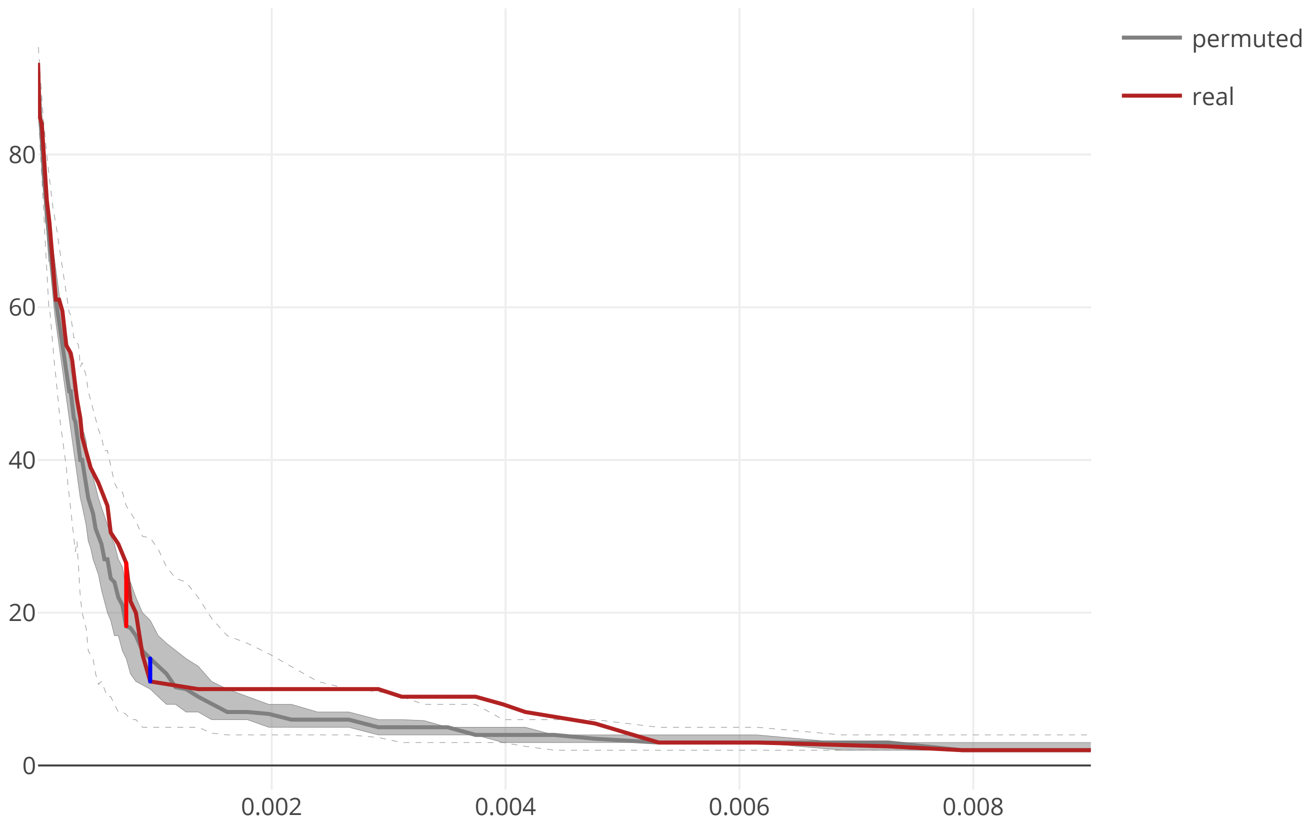

### cov2_SARS_CoV_NSP14_flow_distance_20200609.pdf

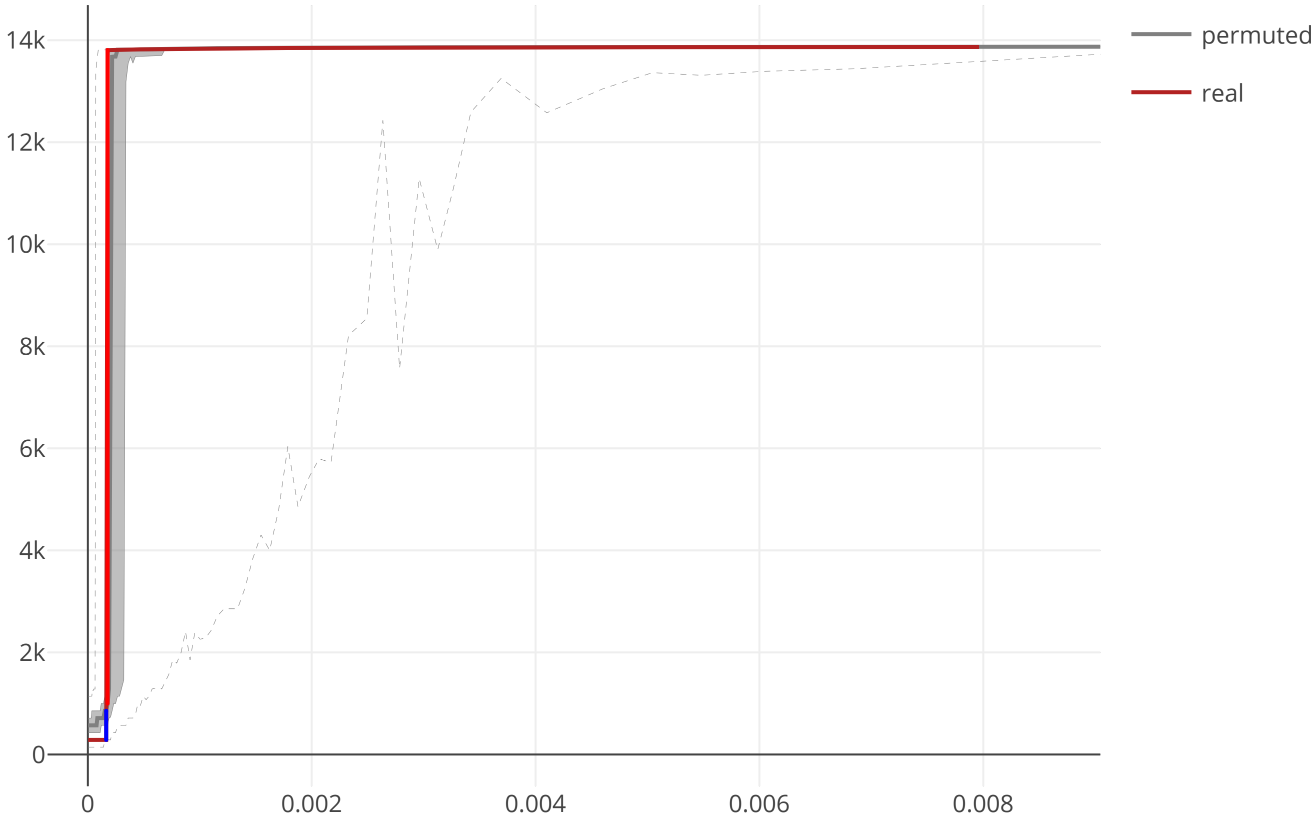

### cov2_SARS_CoV_NSP14_maxcomponent_size_20200609.pdf

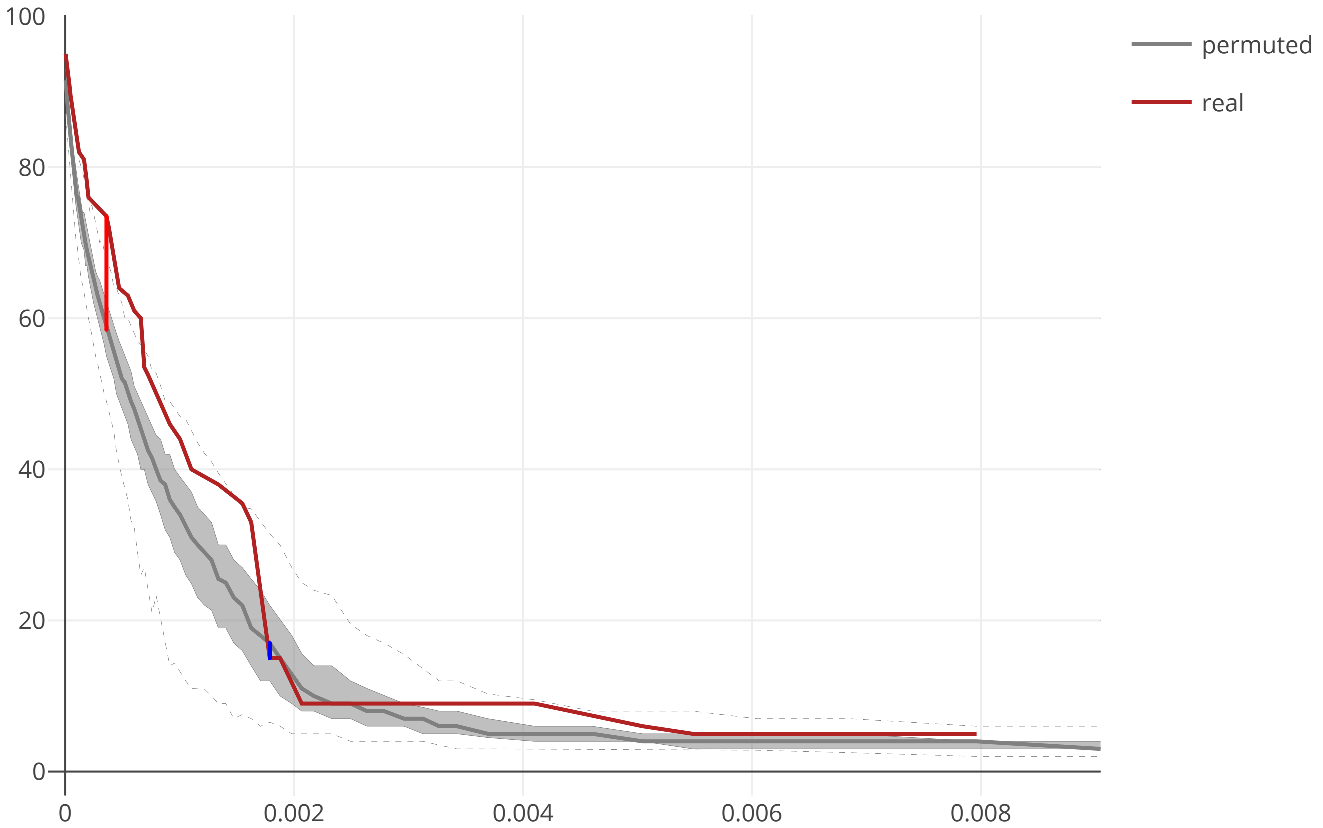

### cov2_SARS_CoV_NSP15_flow_distance_20200609.pdf

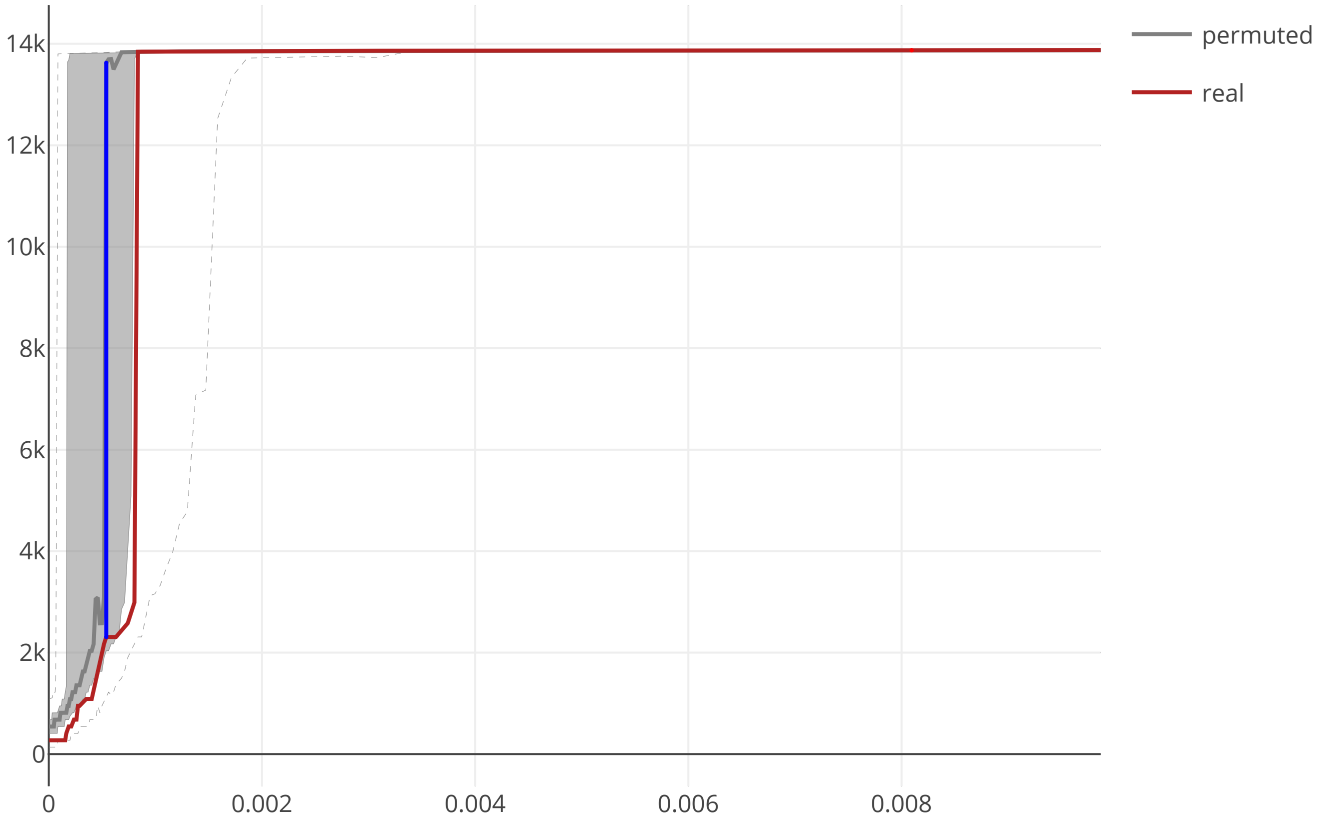

### cov2_SARS_CoV_NSP15_maxcomponent_size_20200609.pdf

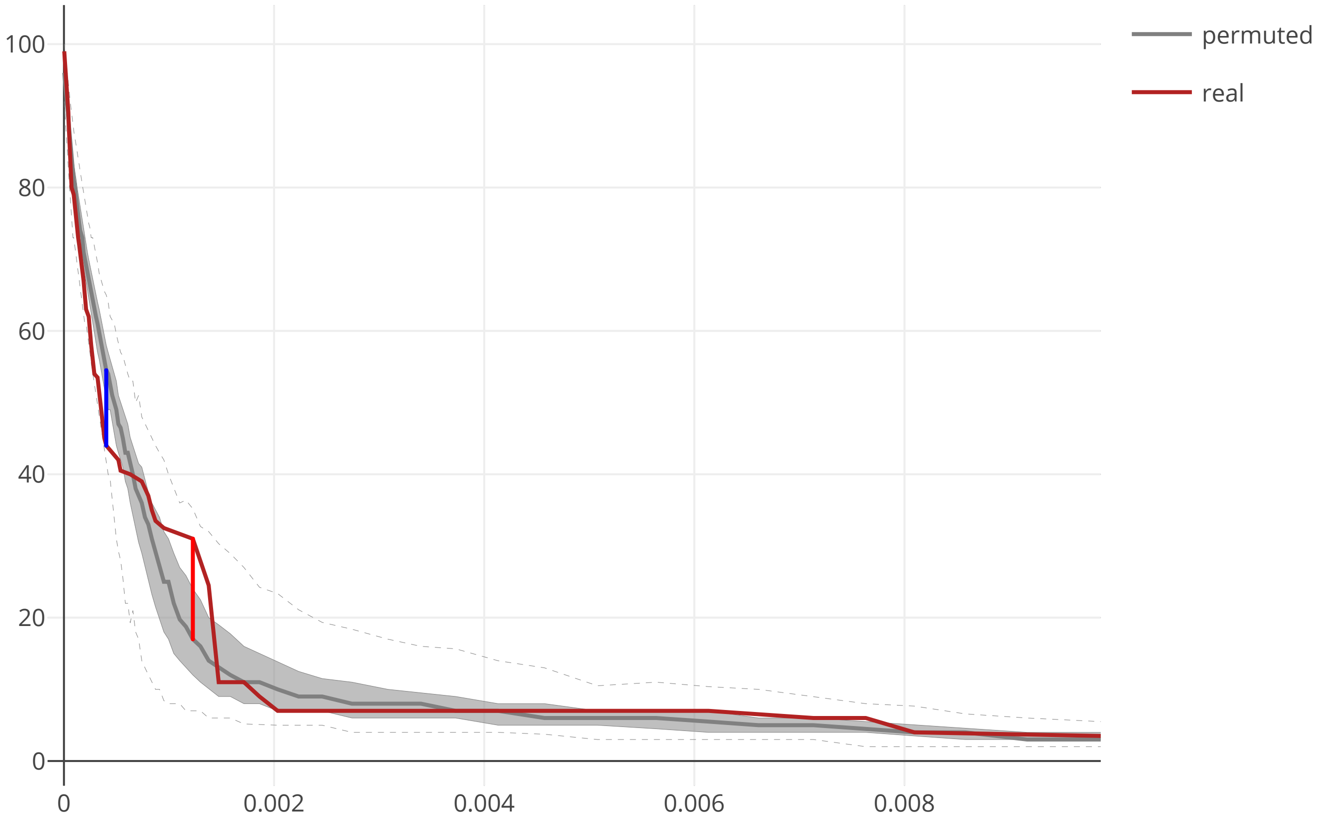

### cov2_SARS_CoV_NSP16_flow_distance_20200609.pdf

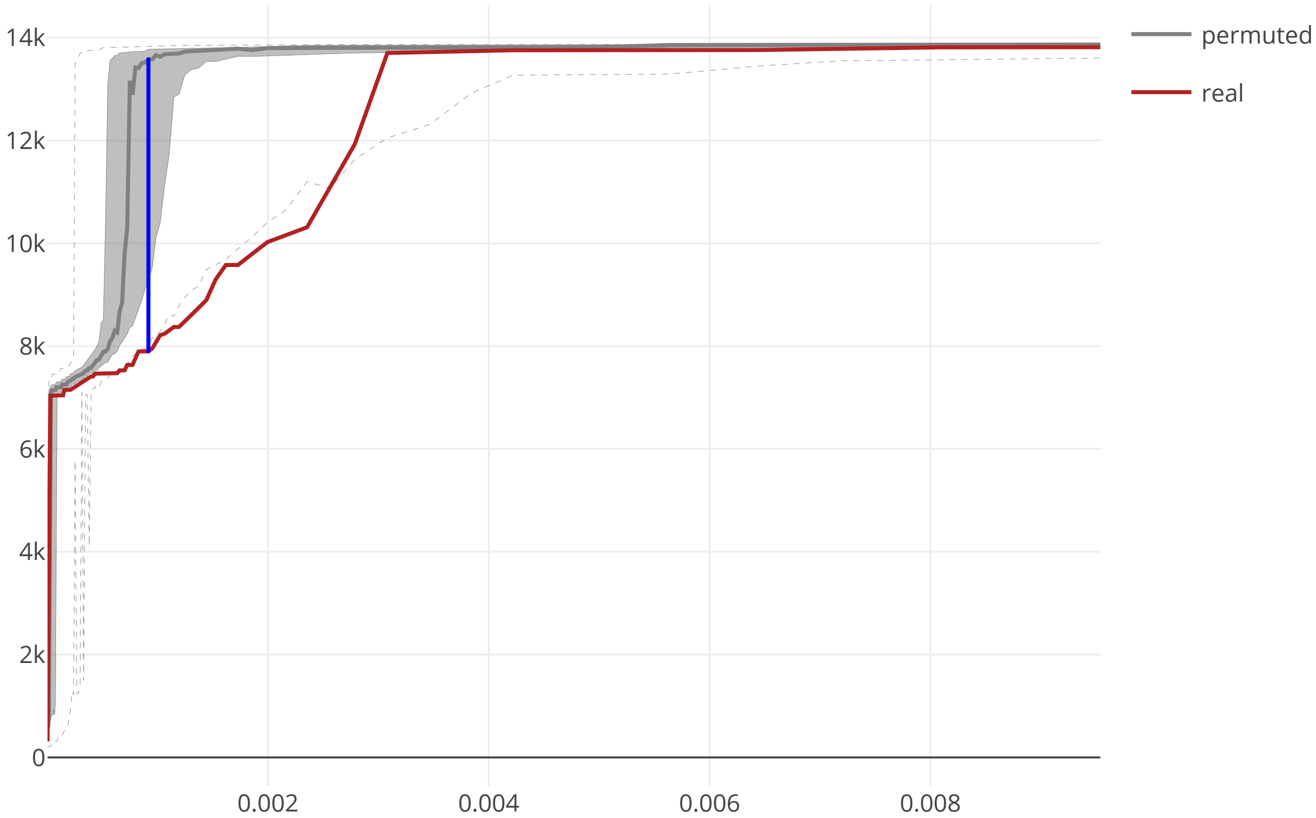

### cov2_SARS_CoV_NSP16_maxcomponent_size_20200609.pdf

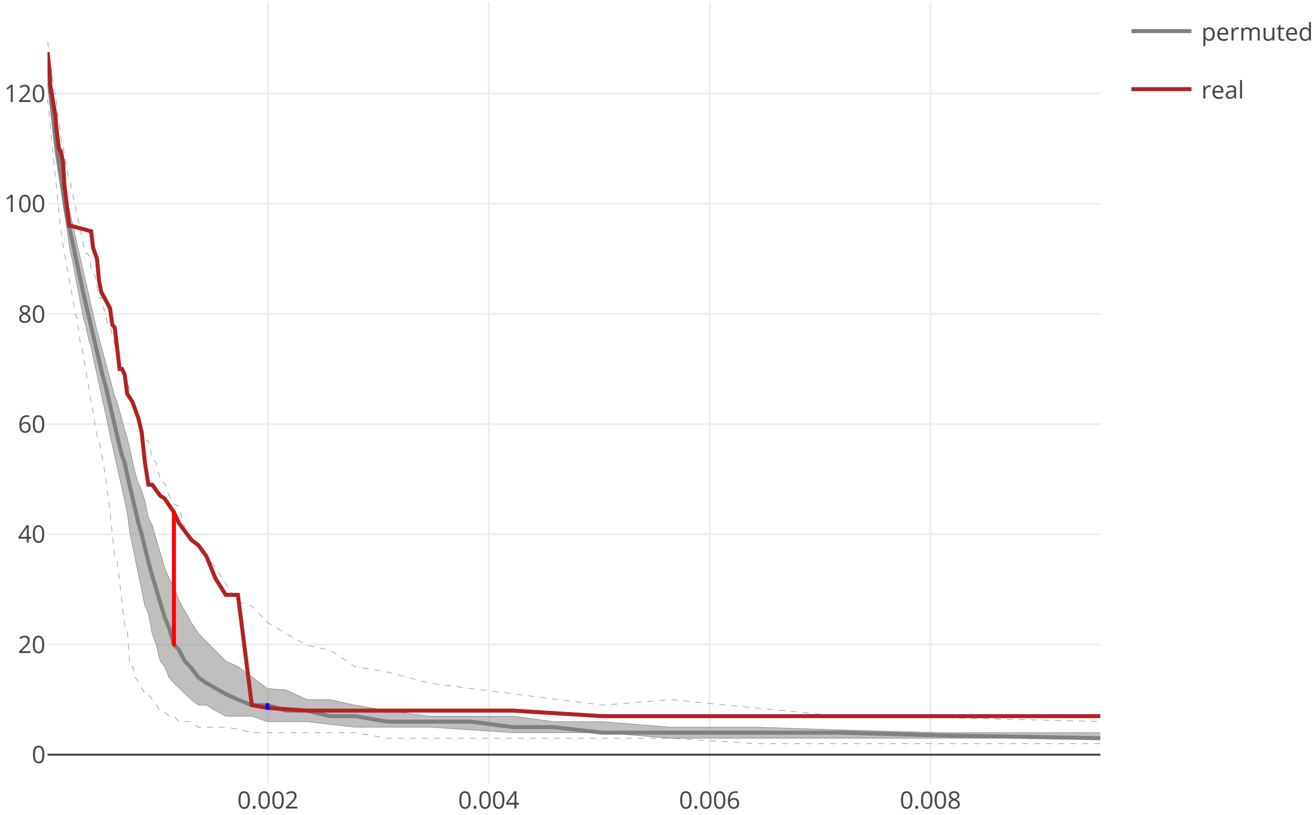

### cov2_SARS_CoV_ORF3a_flow_distance_20200609.pdf

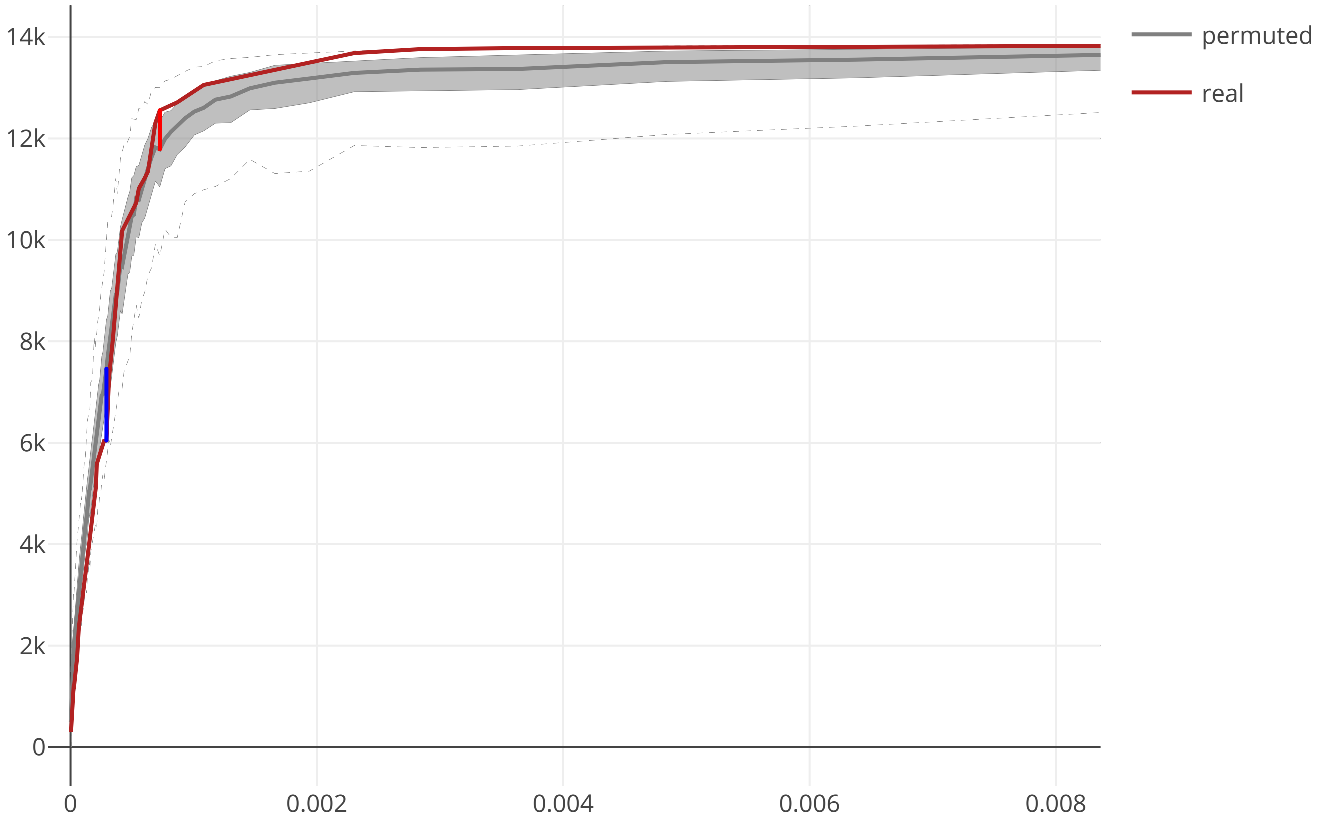

### cov2_SARS_CoV_ORF3a_maxcomponent_size_20200609.pdf

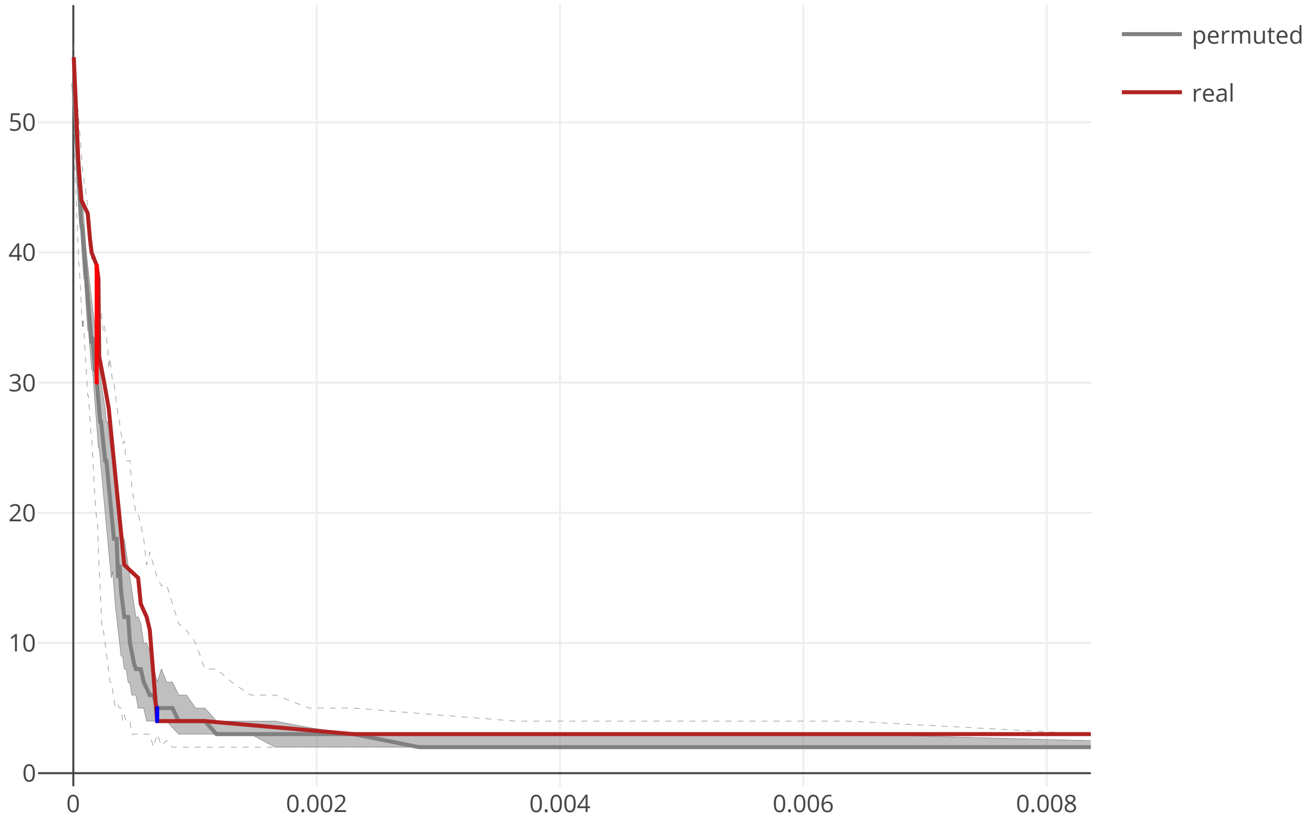

### cov2_SARS_CoV_ORF3b_flow_distance_20200609.pdf

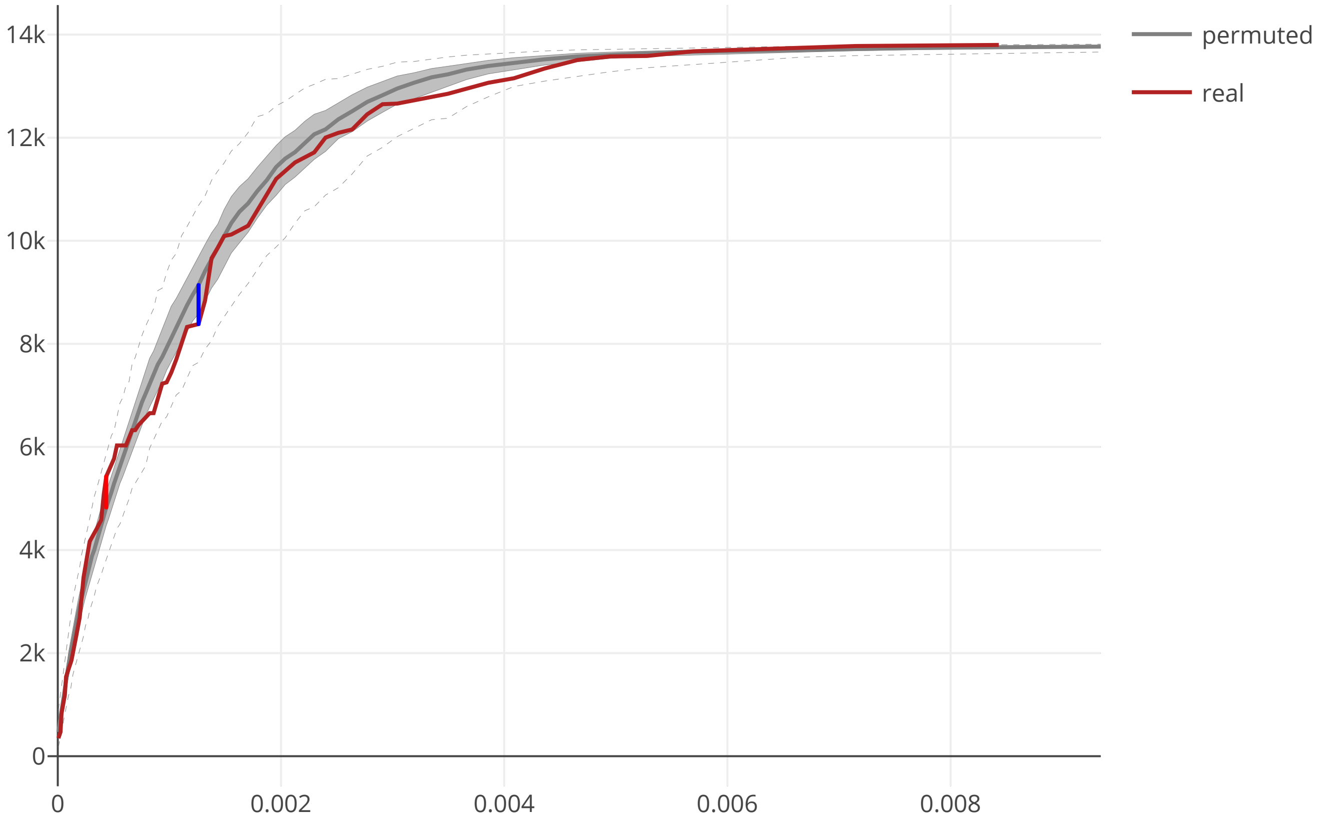

### cov2_SARS_CoV_ORF3b_maxcomponent_size_20200609.pdf

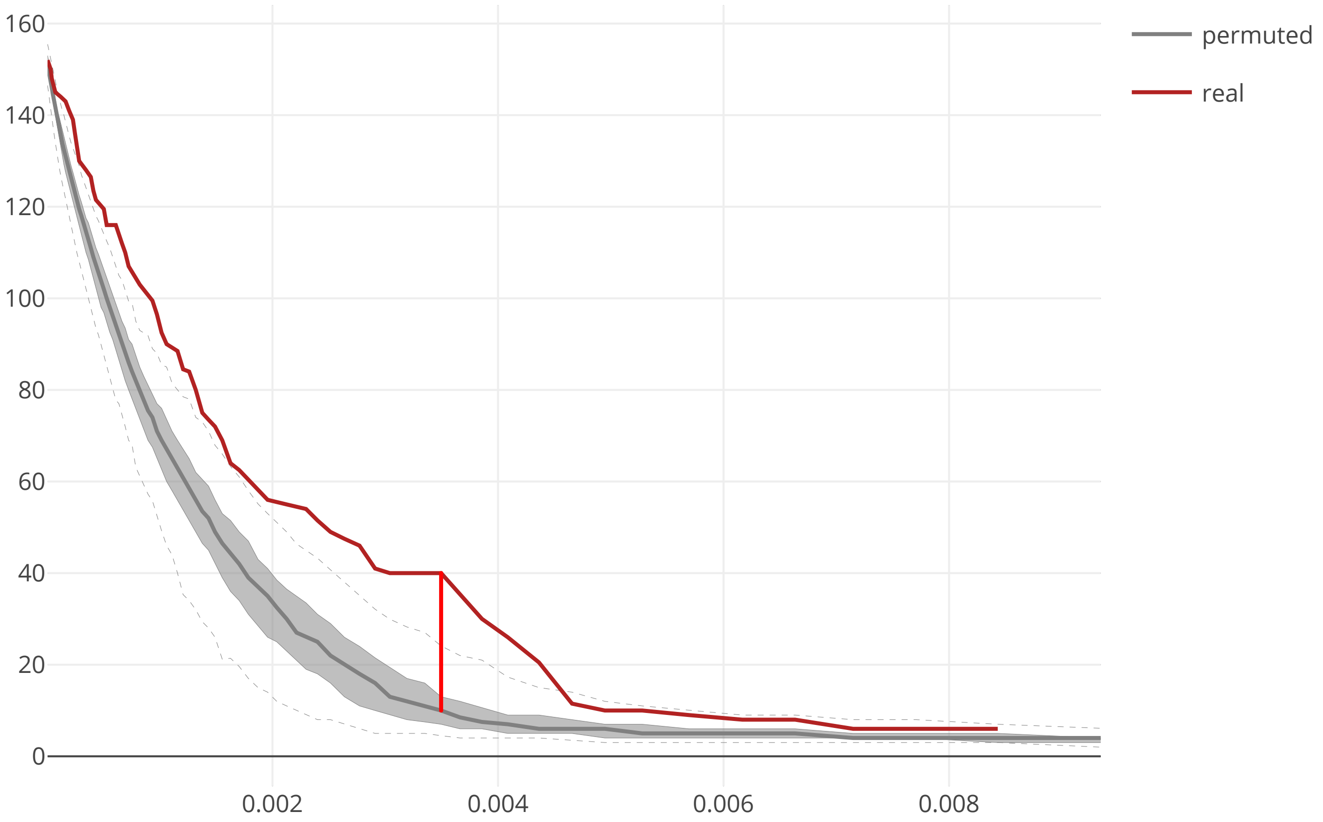

### cov2_SARS_CoV_ORF6_flow_distance_20200609.pdf

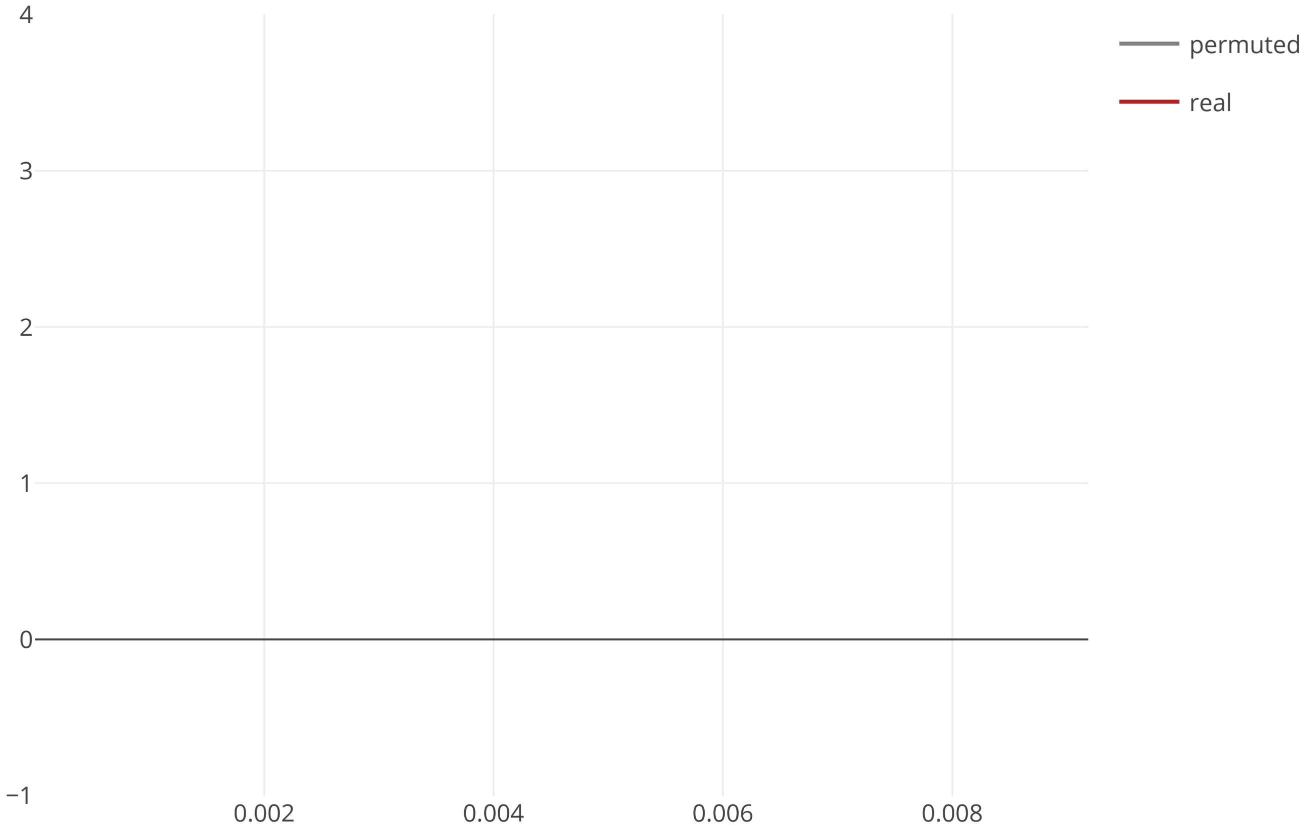

### cov2_SARS_CoV_ORF6_maxcomponent_size_20200609.pdf

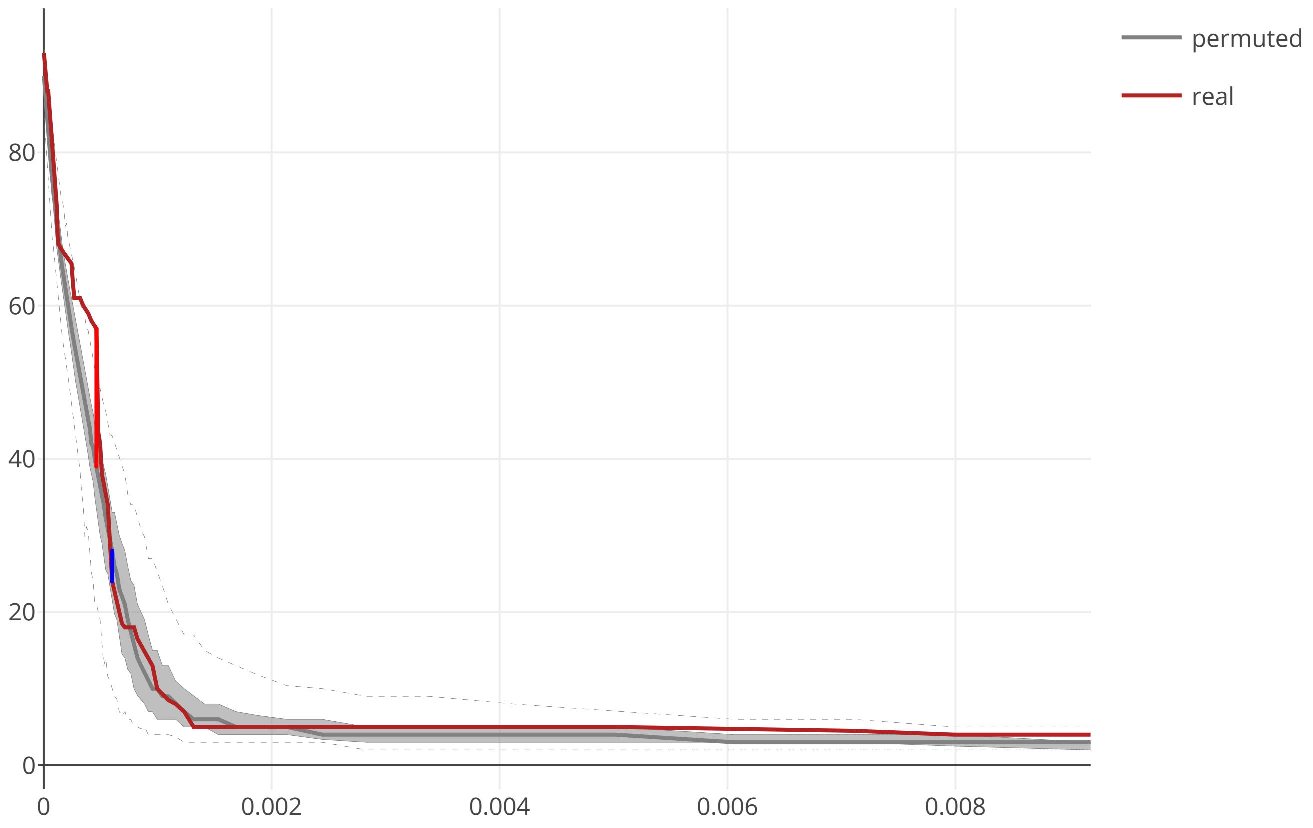

### cov2_SARS_CoV_ORF7a_flow_distance_20200609.pdf

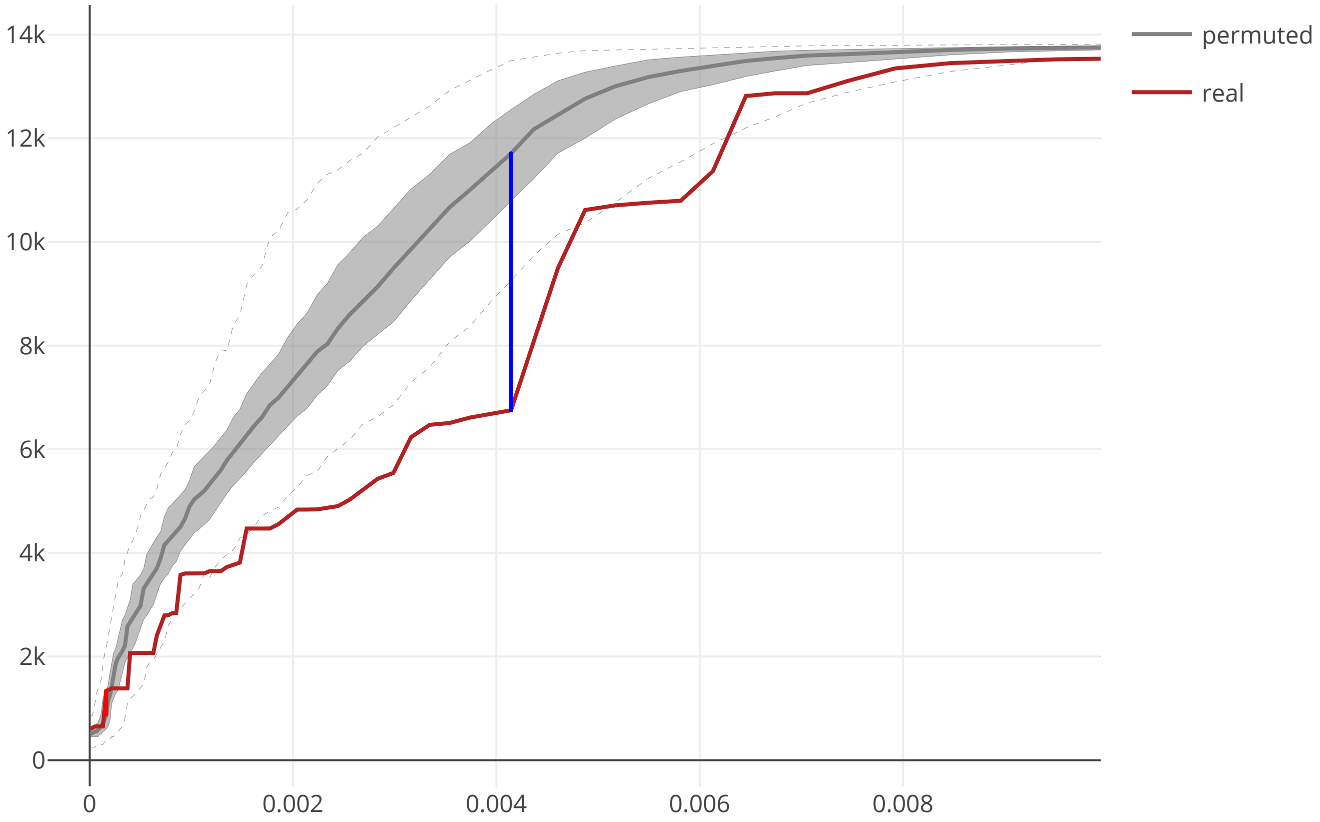

### cov2_SARS_CoV_ORF7a_maxcomponent_size_20200609.pdf

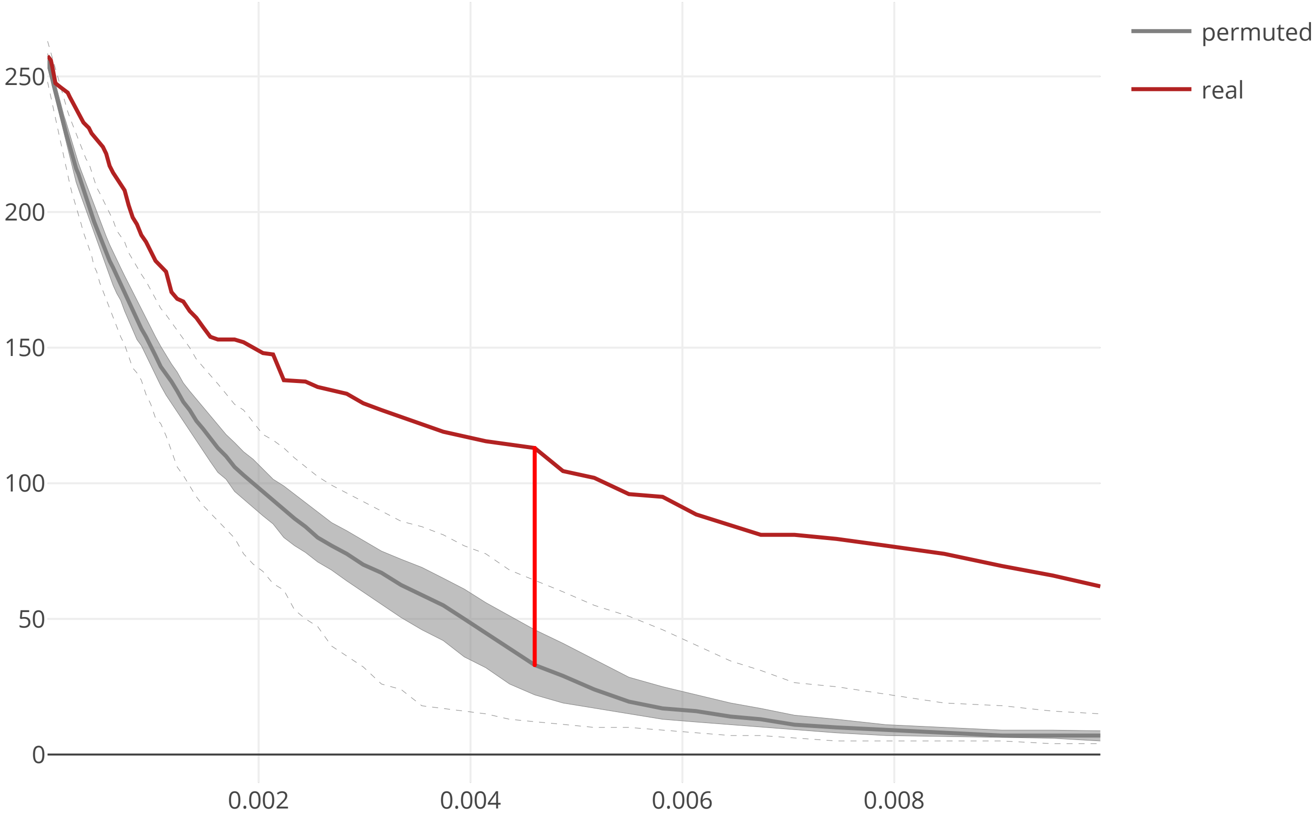

### cov2_SARS_CoV_ORF7b_flow_distance_20200609.pdf

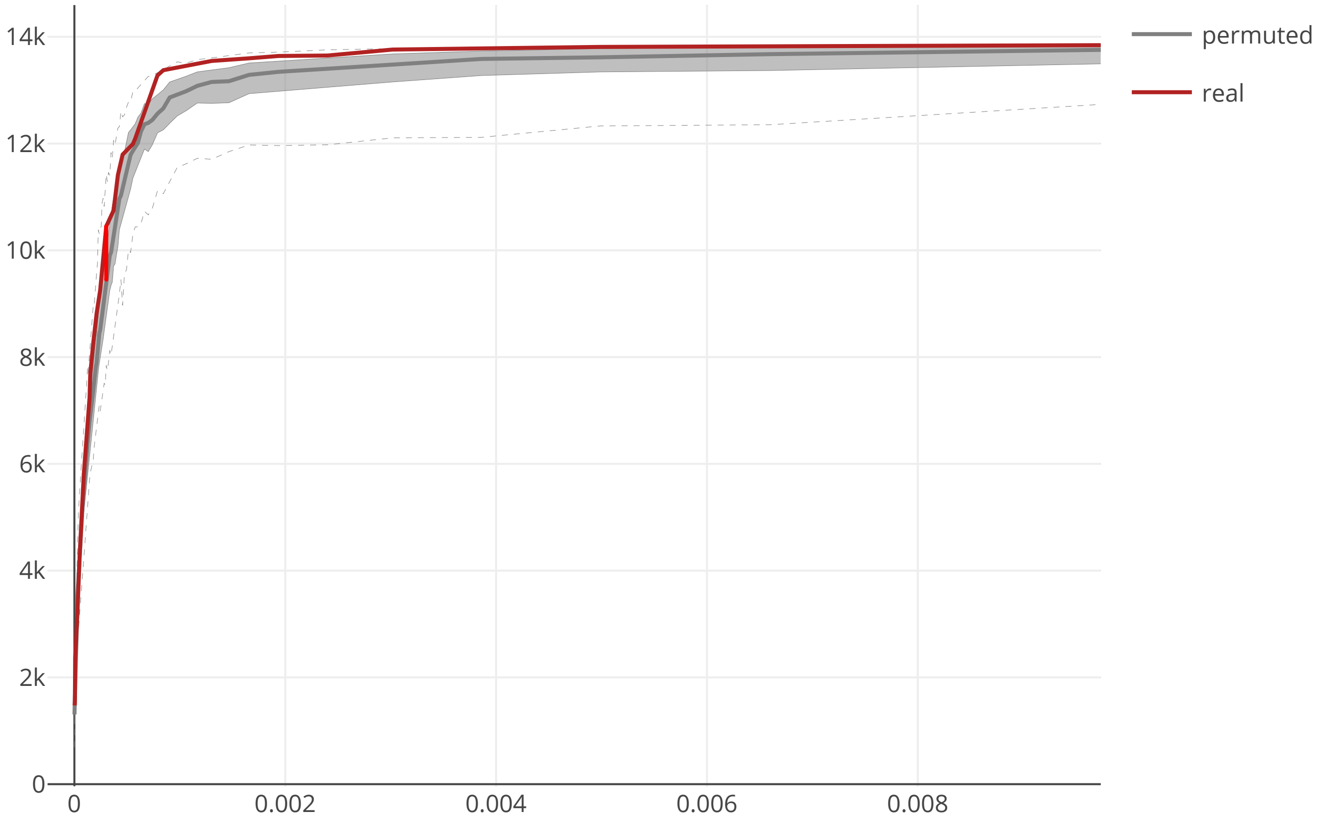

### cov2_SARS_CoV_ORF7b_maxcomponent_size_20200609.pdf

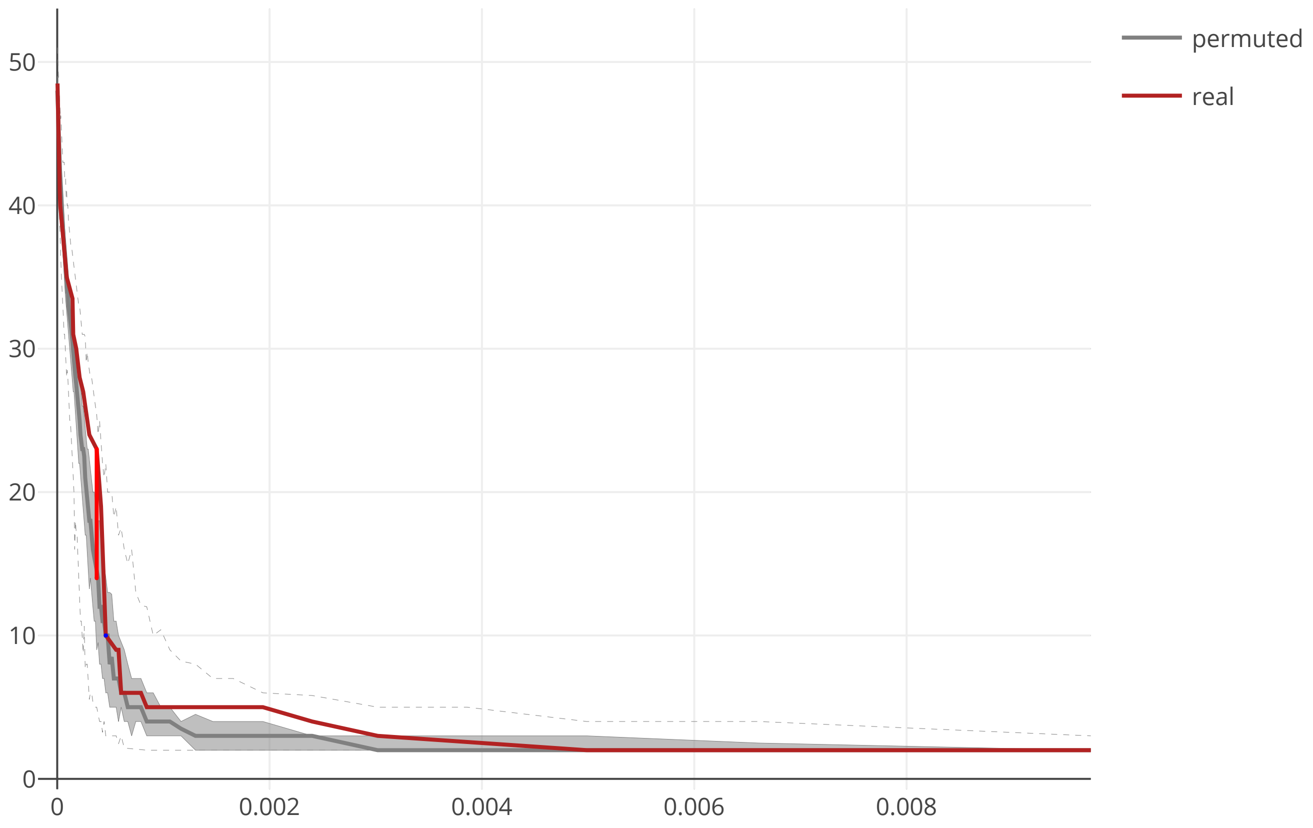

### cov2_SARS_CoV_ORF8a_flow_distance_20200609.pdf

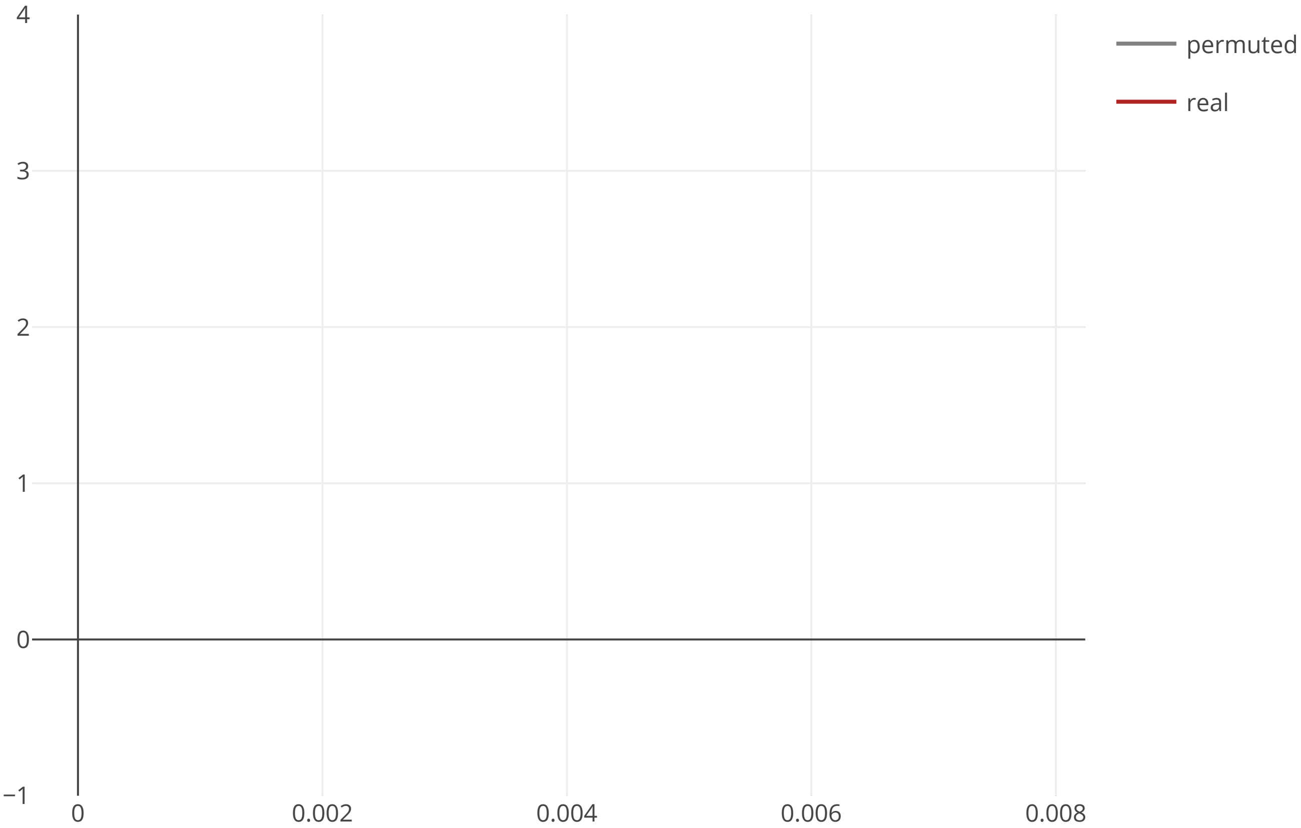

### cov2_SARS_CoV_ORF8a_maxcomponent_size_20200609.pdf

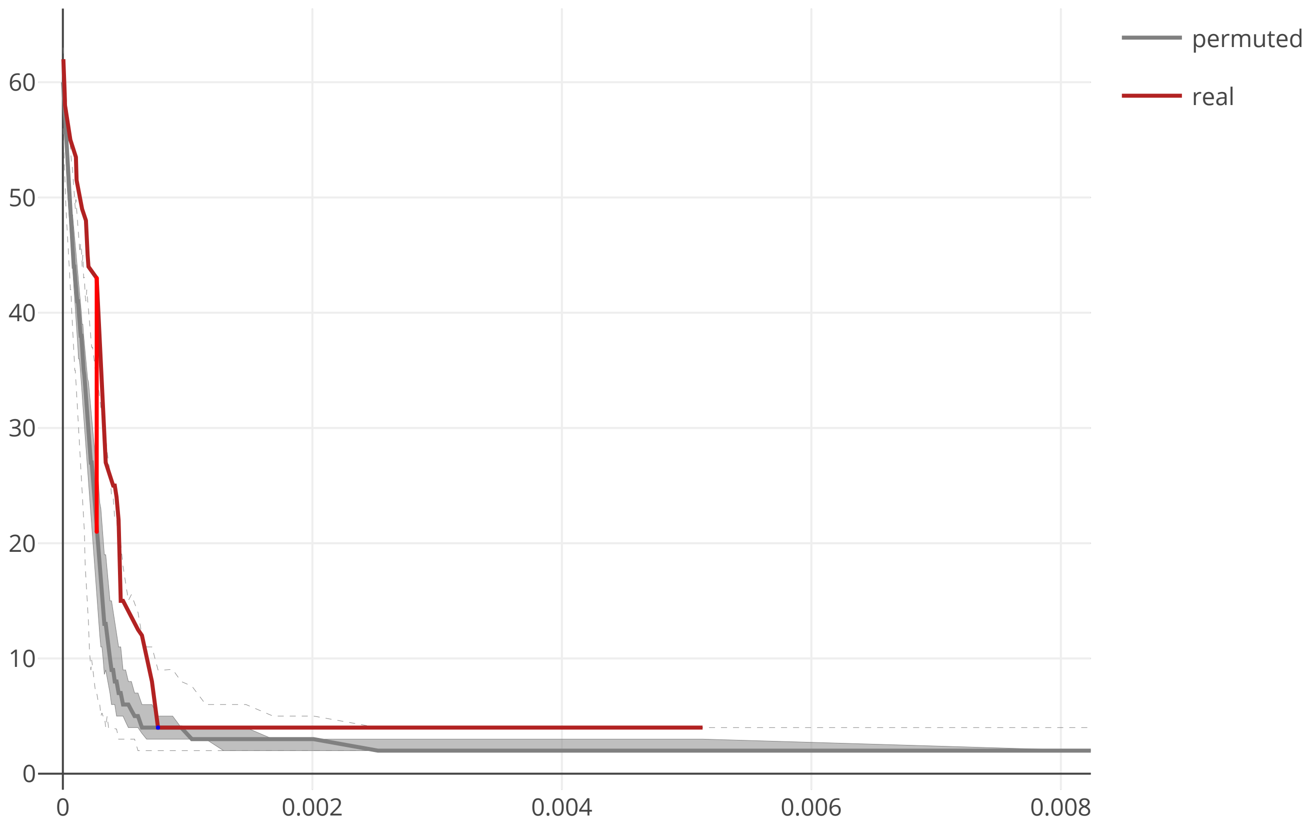

### cov2_SARS_CoV_ORF8b_flow_distance_20200609.pdf

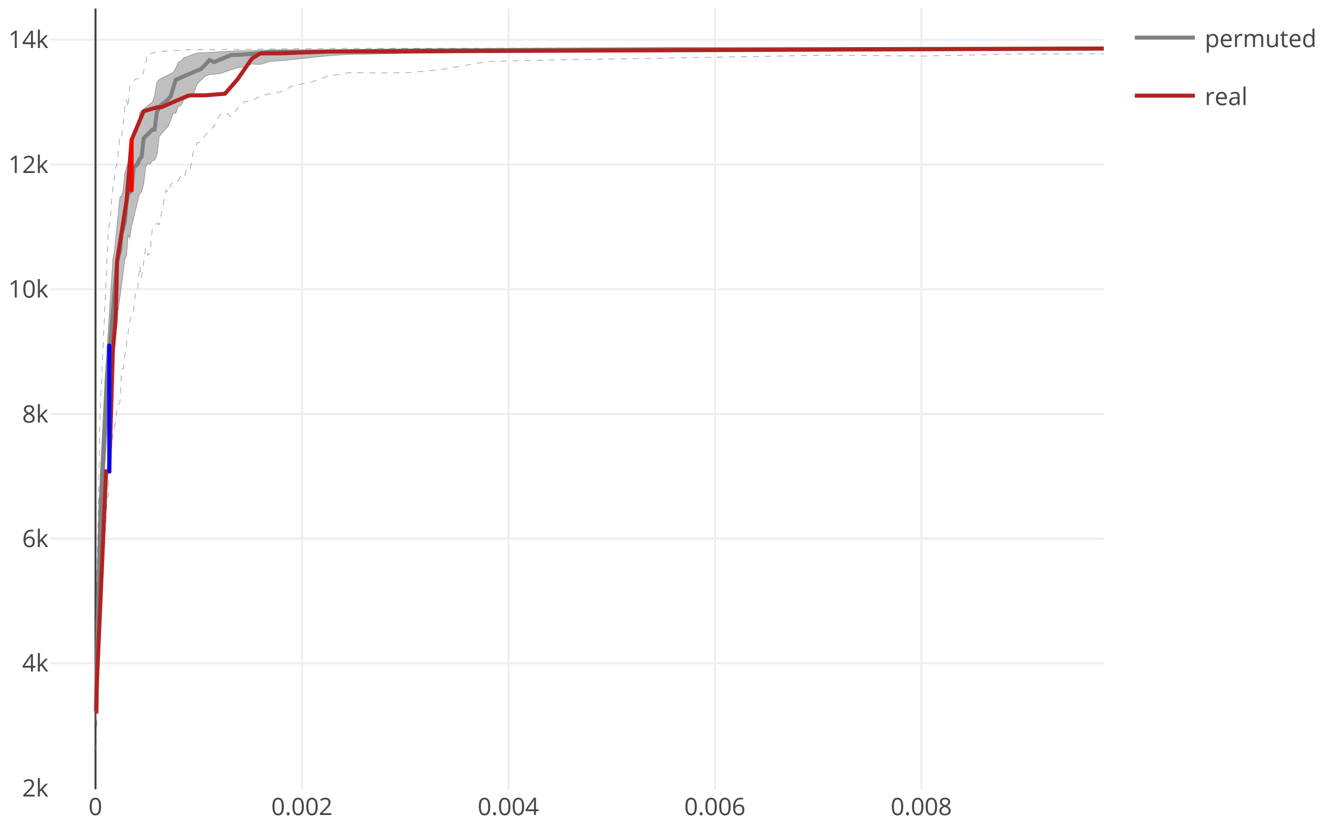

### cov2_SARS_CoV_ORF8b_maxcomponent_size_20200609.pdf

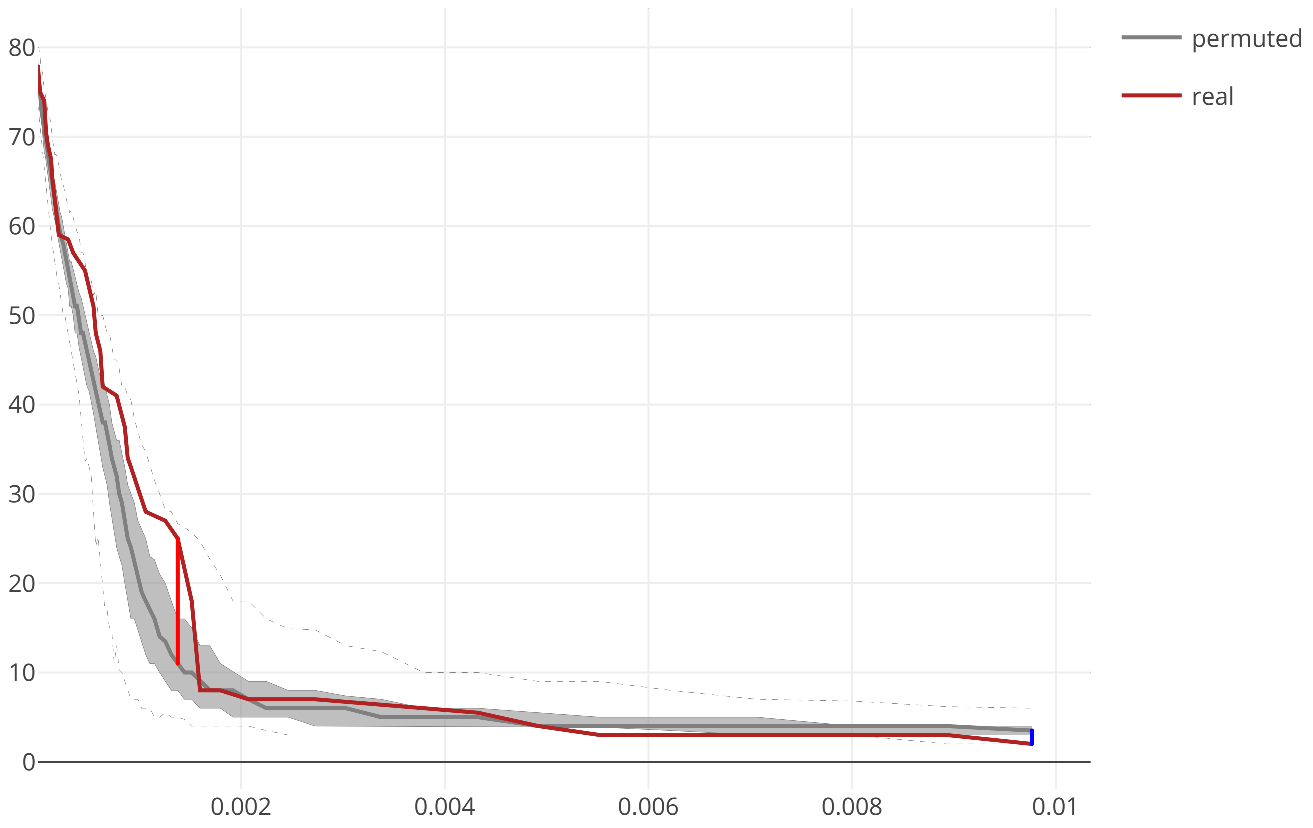
