## Supplementary Discussion for "Multilevel proteomics reveals host perturbations by SARS-CoV-2 and SARS-CoV"

### Supplementary discussion 1: Type-I interferon and NF-κB response in SARS-CoV-2 and SARS-CoV infected A549-ACE2 cells

In line with previous reports[9,22], we did not observe major virus-induced upregulation of type-I interferons (IFN-α/β) and related genes at the mRNA level (e.g. IFNB, IFIT3, MX1), suggesting active viral inhibition of this system. In contrast, both viruses upregulated NF-κB and stress responses, as inferred from the induction of CXCLs (CXCL1, -2, -3, -5, -8), CCL2, NFKBIZ/-A, STAT1/3, IL6 and TNF (Figure 2c, Extended data Fig. 4c-d, Supplementary Table 4) and transcription factor enrichment analysis (Extended data Fig. 4e; Supplementary Tables 4, 8). This was confirmed at the proteome level, where we identified 272 proteins being regulated by SARS-CoV-2 or SARS-CoV (Figure 2a - b, Extended data Fig. 4f). Both viruses induced ACE2 downregulation only at the protein level (Extended data Fig. 4g-h). Similarly, SARS-CoV-2 and SARS-CoV failed to induce a detectable IFN-α/β proteomic signature while activating the pro-inflammatory NF-κB pathway (NFKB2, RELB, JUNB, TNFAIP2) (Extended data Fig. 4f, i; Supplementary Table 5).

**Supplementary discussion 2: Structure-guided interpretation of post-translation modifications functionality on Nucleocapsid protein N**

The localization of K338 at the end of the $\beta$2-strands of the $\beta$1-$\beta$2 hairpin could serve two potential functions (Figure 3c, lower close-up). The side chain of this residue is solvent-exposed, and it is therefore conceivable that its ubiquitination, specific to SARS-CoV-2, may explain divergences in virus-host interactomes (Supplementary Table 2). Furthermore, the SARS-CoV-2 N C-terminal domain (CTD) four-stranded $\beta$-sheet dimeric interface can undergo significant rearrangements upon sliding and departing of the $\beta$2 strands from each other[32], implying possible conformational consequences of SARS-CoV-2 N K338 ubiquitination. Analogous characterization of the S310/311, which is located at the beginning of the α5 helix, revealed that its phosphorylation contributes to a transfer of an inter-chain interaction, from R262/263 to T263/264, thus stabilizing the structure of the CTD dimer (Figure 3c, upper close-up). In addition, S310/311 phosphorylation gives rise to two negatively charged areas on a polar and basic, solvent-exposed, groove involved in RNA binding[32], indicating its possible implication in nucleocapsid formation and viral genome packaging (Extended data Figure 6h).
